# Direct synthesis of EM-visible gold nanoparticles on genetically encoded tags for single-molecule visualization in cells

**DOI:** 10.1101/520999

**Authors:** Zhaodi Jiang, Xiumei Jin, Yuhua Li, Sitong Liu, Xiao-Man Liu, Ying-Ying Wang, Pei Zhao, Xinbin Cai, Ying Liu, Yaqi Tang, Xiaobin Sun, Yan Liu, Yanyong Hu, Ming Li, Gaihong Cai, Xiangbing Qi, She Chen, Li-Lin Du, Wanzhong He

**Affiliations:** PTN Graduate Program, School of Life Sciences, Tsinghua University, Beijing, China; National Institute of Biological Sciences, Beijing, 102206, China

## Abstract

Single-molecule visualization in cells with genetically encoded tags for electron microscopy (EM) has been a long-awaited but unimplemented tool for cell biologists. Here, we report an approach for directly synthesizing EM-visible gold nanoparticles (AuNPs) on cysteine-rich tags for single-molecule visualization in cells. We first uncovered an auto-nucleation suppression mechanism that allows specific synthesis of AuNPs on isolated cysteine-rich tags. We next exploited this mechanism to develop an approach for single-molecule detection of proteins in prokaryotic cells and achieved an unprecedented labeling efficiency. We then expanded it to more complicated eukaryotic cells and successfully detected the proteins targeted to various organelles, including the membranes of endoplasmic reticulum (ER) and nuclear envelope, ER lumen, nuclear pores, spindle pole bodies, and mitochondrial matrix. Thus, our implementation of genetically encoded tags for EM should allow cell biologists to address an enormous range of biological questions at single-molecule level in diverse cellular ultrastructural contexts without using antibodies.

## INTRODUCTION

The development of genetically encoded tags for electron microscopy (EM), analogous to the fluorescent proteins used in light microscopy (LM)^1^, would be highly valuable because it would enable the characterization of the precise localization of individual protein molecules in the cellular ultrastructural context. Such a development would overcome several limitations of traditional antibody-based immuno-EM staining^2,3^, including poor antigen accessibility, steric hindrance, loss of antigenicity after fixation, and sub-optimal antibody performance, all of which result in low labeling efficiency (< 10%). Moreover, genetically encoded tags for EM would also overcome several limitations of light microscopy^4^, including the inability to visualize cellular organelles that are not fluorescently labeled—which often requires experimental verification with a correlated EM image—and the inability of super-resolution light microscopy to distinguish individual molecules at high abundances.

Given the enormous potential utility of the aforementioned genetically encoded tags for EM, it is not surprising that two types of these tags have been explored in recent years^5–21^. However, none of those implementations have realized single-molecule visualization and counting in cells. The first technology developed in this area is the tag-mediated polymerization of 3,3’-diaminobenzenidine into local precipitates for EM contrasting by OsO_4_ staining^5–9^ or silver/gold precipitation^10^. This approach has been applied for staining cellular regions with highly abundant protein accumulation but is unsuitable for single-molecule counting applications^5–10^. The second technology is based on metal-binding tags, including ferritin^21^ and metallothionein (MT)^11–20^, only the latter of which has real potential for use in single-molecule visualization and counting; the iron-loading ferritin complex is considered too big to be a promising tag^21^. MT is a well-known metal-binding protein that can bind various heavy metal ions (e.g., Zn^2+^, Cd^2+^, Cu^+^, Ag^+^, Au^+^, Hg^2+^, etc.) through its 20 cysteine (Cys) residues^22^. MT tags for EM have been tested with purified fusion proteins^11–15^ and complexes^15^, and more recently for tagging fusion proteins in cells^16–20^. These efforts have clearly demonstrated that the gold-binding properties of MT make it a potential candidate to be used as a genetically encoded tag for EM, but none of these approaches has been successfully implemented for single-molecule detection in cells.

To implement cysteine-rich tags for EM detection in cells, several challenges must be overcome: designing fixative-resistant tags for achieving preservation of ultrastructures and high labelling efficiency, delivery of gold precursors across membrane barriers, minimizing physiological disturbances, suppressing nonspecific staining noise, and generating gold agglomerations of sufficient size and density to enable highly sensitive and precise EM detection. To address these challenges, we here explored an approach based on the synthesis of 2-6 nm diameter gold nanoparticles (AuNPs) on specific cysteine-rich tags in chemically fixed and fast-frozen cells. Building on the Brust-Schiffrin principle of thiolate-capped AuNPs synthesis^23^, we designed and implemented a new concept for the reliable and specific synthesis of AuNPs on native MT tags and on a new type of small cysteine-rich tags that we engineered. We developed additional tags, including one type that lacked aldehyde-reactive residues (“MTn”), a smaller variant of MTn (“MTα”), and a disulfide-rich tag engineered from an insect antifreeze protein, tmAFP^24^ (“AFP”) (**Fig.1a-c** and **Supplementary Table 1**).

**Figure 1.**
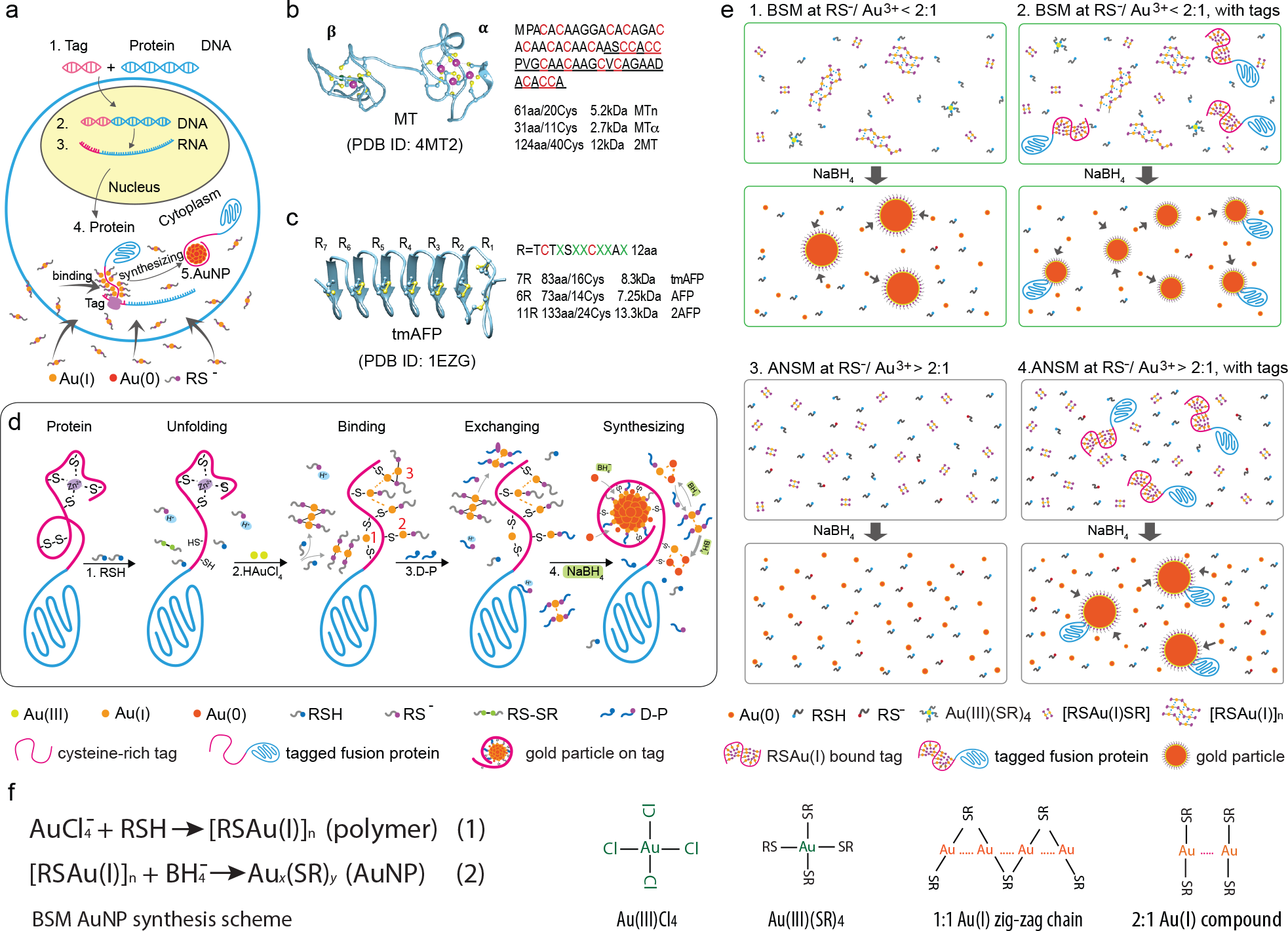
Scheme for the implementation of clonable tags for electron microscopy. **(a)** The Concept of using cysteine-rich tag for protein localization in cells. **(b-c)** Structures and amino acids sequences of the cysteine-rich tags: metallothionein (MT), and antifreeze protein (AFP), MTn and MTα are engineered from MT **(Supplementary Table 1). (d)** Procedures for synthesis of AuNPs on cysteine-rich tag: (i) unfolding the tag by thiol ligands; (ii) Au(I) compounds bind to cysteine residuals on the tag in (1) ligand retention mode (Cys-Au-SR), (2) cysteine-bridged mode (Cys-Au-Cys)^34^ and (3) possible polymer attached mode; (iii) exchanging with stronger D-penicillamine (D-P) ligand for suppressing noises; (iv) reducing the tag bound Au(I) to Au(0) atoms to form a AuNP on tag. (**e**) Schemes for synthesizing AuNPs on the tags with Brust-Schiffrin method (BSM) or the ANSM-based method: (1) At RS^−^/Au(III) < 2:1 conditions (BSM), Au(I)_n_ SR_m_ forming large polymers, which act as auto-nucleation cores for forming AuNPs; (2) When the cysteine-rich tags added to the BSM condition, both the Au(I)_n_ SR_m_ polymers and the tags were acted as nucleation cores for forming AuNPs; (3) At RS^−^ /Au(III)> 2:1 conditions (ANSM), only forms countless tiny Au(I)SR species, there were no EM-visible AuNPs formed due to the absence of large auto-nucleation cores; (4) When the tags added to the ANSM conditions, only individual cysteine-rich tags acted nucleation cores for forming AuNPs. (f) Two steps reducing scheme of AuNPs synthesis with BSM, and the typical four-coordinate Au(III), two-coordinate Au(I) compounds.

For AuNPs synthesis, rather than simply considering the thiol-to-gold ratio (abbreviated as RSH/Au), as is typical in the Brust-Schiffrin method, we systematically investigated the role of thiolate-to-gold (abbreviated as RS^−^/Au) ratio. We were thus able to discover an auto-nucleation suppression mechanism (ANSM) that enables the specific synthesis of AuNPs on cysteine-rich tags, including, surprisingly, even the smallest cysteine-rich tag, MTα (only 2.72 kDa); all of the AuNPs labeled tags are clearly visible under EM. We next implemented the ANSM concept to synthesize AuNPs on cysteine-rich tags expressed in three model organisms, *Escherichia coli* (*E. coli*), the fission yeast *Schizosaccharomyces pombe* (*S. pombe*), and HeLa cells. The application of ANSM in cells required us to implement an oxidization-based strategy to protect the thiol groups of cysteine-rich tags prior to the exposure of an aldehyde-fixative reagent that is commonly used for preservation of cellular ultrastructures. We were thus able to map individual protein molecules in intact cells. This new technology enables the precise localization of fusion proteins in the cellular ultrastructural context without the need for antibodies and should permit cell biologists to quantitatively characterize the roles of proteins of interest in cells via genetic manipulation.

## RESULTS

### Design of fixative-resistant cysteine-rich tags for electron microscopy

An ideal tag for generating an EM-visible electron-dense label in cells would be small, fixative-resistant, and enable precise and sensitive detection. We considered the known limitations of MT in EM tag applications when we designed our tags^12^, primarily the need for multiple copies of MT to form EM-detectable clusters^12–17^ and the reactivity of native MT tags with aldehyde-fixatives. To create more robust tags, first, we engineered mouse MT-1 protein (61 amino acid) into an aldehyde-fixative-resistant variant, MTn, by replacing the 21 aldehyde-reactive residues (*e.g.*, Lys, Ser, Thr, Gln)^25^ with aldehyde-inert residues (*e.g.*, Ala) (**Fig.1b**, **Supplementary Table 1**). We also developed a smaller variant, MTα, that consists of only the alpha domain of MTn. Additionally, to exploit the natural fixative-resistance of disulfide bonds, we employed another cysteine-rich tag, AFP, which we developed from an insect antifreeze protein, tmAFP^24^. AFP tags form highly stable structures via disulfide bonds in oxidizing cellular compartments (**Fig.1c**, **Supplementary Table 1**). The design of these tags should, in theory, ensure the formation of relatively stable structures that protect them from fixative cross-linking denaturation. Moreover, upon exposure to nucleophiles during the synthesis of AuNPs, these tags can unfold readily (**Fig.1d**); this property is superior to tightly-packed cysteine-rich proteins (*e.g.*, BSA and transferrin) that require harsh conditions to allow gold cluster formation^26,27^. To validate whether our engineered tags function as designed in cells, the genes for these tags were synthesized and used to construct tag-fused maltose-binding protein (MBP) that was expressed in *E. coli* (**Supplementary Fig.1**).

### The auto-nucleation suppression mechanism for AuNP synthesis

The Brust-Schiffrin method (BSM)^23^ is widely used for synthesizing thiolate-capped gold nanoclusters. The core mechanism of BSM involves two reduction steps: first, thiol ligands (“RSH”) are used to reduce HAuCl_4_ to form thiolate-Au(I) (abbreviated as [RSAu(I)]_n_) polymers; these polymers are then reduced by NaBH_4_ to form Au(0) nanoparticles (AuNPs) (**Fig.1f**). The size of the AuNPs can be tuned by varying the RSH-to-Au ratio^28^ and by changing the pH^29^. Typical BSM reactions are performed using a thiol-to-gold (RSH/Au) ratios within a relatively narrow range (*e.g*., 1:6 to 6:1), and feature a polymer-mediated auto-nucleation mechanism^28^ (**Fig.1e1**). The cysteine-rich tags can be considered as similar nucleation cores as the [RSAu(I)]_n_ polymers. If the cysteine-rich tags are added into the first step of the regular BSM AuNPs synthesis, both the tags and the [Au(I)SR]_n_ polymers would be expected to act as nucleation cores for AuNPs formation (**Fig.1e2**). Therefore, the classic BSM would not be applicable for the unambiguous identification of the tags by the AuNPs, unless the AuNPs formed via Au(I)SR]_n_ polymers were eliminated.

To suppress the polymer-mediated formation of AuNPs, we here explored the idea of eliminating those [RSAu(I)]_n_ polymers from the BSM process (**Fig.1e3**). It is known that the Au(I) can form two types of Au(I)-to-thiolate compounds: the 1:1 zigzag chain (or cyclic) compounds and the 2:1 compounds^30,31^ (**Fig.1f**). However, no one has clearly characterized the precise role of the thiolate anions (RS^−^) in the BSM. In aqueous solution, RSH dissociates as RS^−^ and H^+^ in a pKa of RSH and pH-dependent manner, the RS^−^ concentration can be calculated using the Henderson-Hasselbalch equation: *RS*^−^ = *RSH*/(1 + 10^(*pka−PH*)^). Thus, the concentration of RS^−^ is tunable by varying the initial concentration of RSH and the pH value. Here we performed BSM AuNPs synthesis experiments by tuning the RS^−^/Au ratio (**Fig2a-d, Supplementary Fig.2**). We discovered an interesting mechanism that can fully suppress the polymer-mediated auto-nucleation formation of AuNPs. For example, when the RS^−^/Au ratio < 2, the typical AuNPs were formed (as occurs in classic BSM). In contrast, no AuNPs were formed at an RS^−^/Au ratio ≥ 2. Building on this discovery, we thus proposed a new strategy for synthesizing tag-specific AuNPs based on performing BSM at an RS^−^/Au ratio ≥ 2; this can suppress off-target (*i.e.*, non-tagged) formation of AuNPs via an auto-nucleation suppression mechanism (ANSM) (**Fig.1d, e4**). The principle of ANSM and its application with genetically encoded tags are presented below.

**Figure 2.**
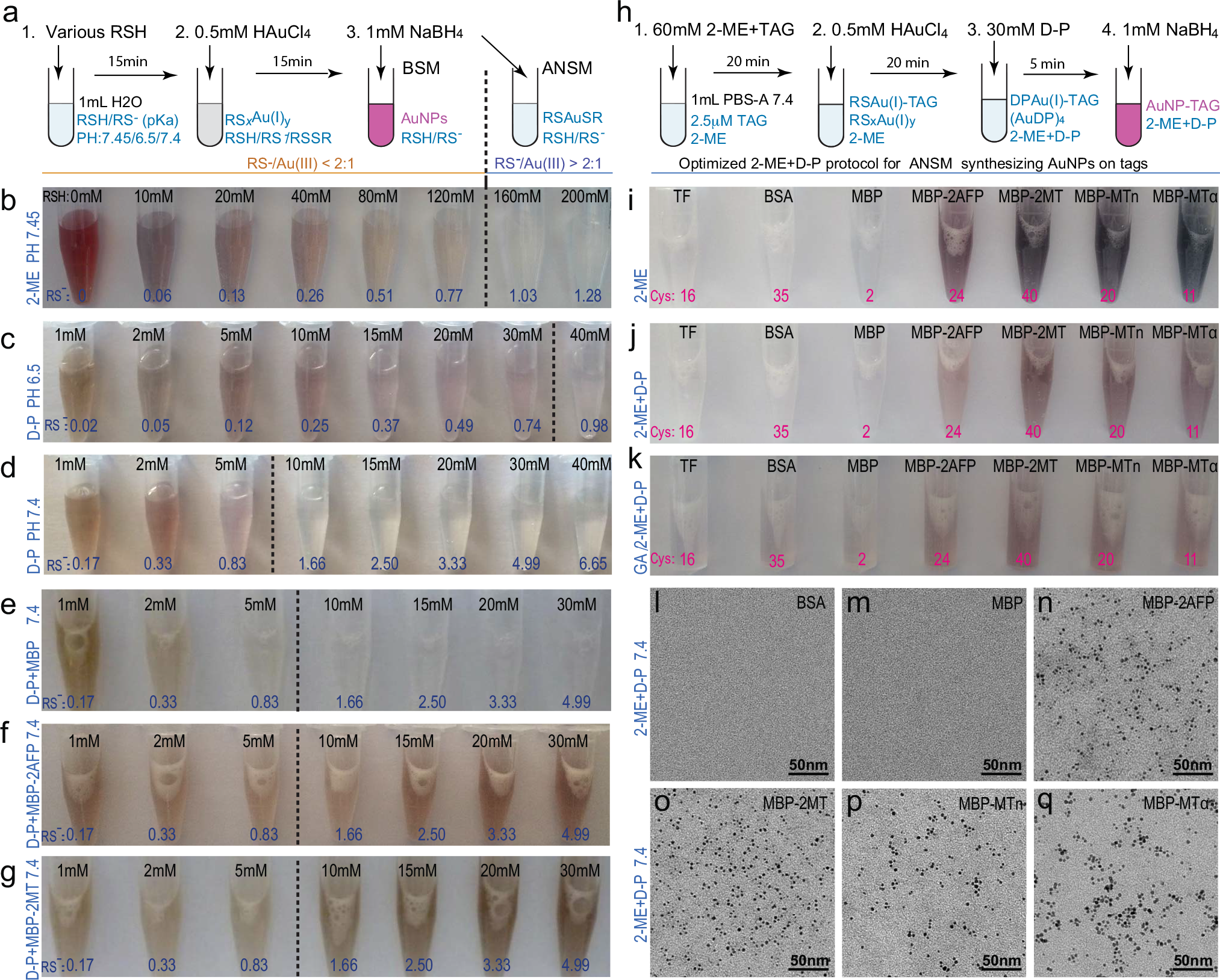
The implementation of ANSM-based synthesis of AuNPs on cysteine-rich tags. **(a)** Scheme for performing BSM at various concentrations of thiol ligands (RSH) and different pH conditions. **(b-d)** BSM for synthesizing thiol-capped AuNPs was conducted: in a 1 mL of the aqueous solution with various concentrations of 2-ME or D-P, by adding 0.5 mM HAuCl_4_ for 15 min followed by 1 mM NaBH_4_ reduction at pH 7.45, 6.5 and 7.4. The initial concentrations of RSH and the corresponding calculated concentrations of RS^−^ were labeled on the top and bottom respectively (pKa for 2-ME: 9.64; pKa for D-P: 8.1). **(e-g)** 2.5 μM of MBP, MBP-2AFP, MBP-2MT proteins added a series of 1 mL aqueous solutions with the same conditions (pH, D-P, HAuCl_4_, and NaBH_4_) as used for (d). **(h)** Scheme for synthesizing AuNPs on tags with optimized 2-ME/D-P protocol used for (j-k). (**i-k**) 2.5 μM of seven proteins, transferrin(TF), BSA, MBP, MBP-2AFP, MBP-2MT, MBP-MTn, MBP-MTα, with different numbers of cysteine residuals (labeled on the bottom), were used for AuNPs synthesis under the pH=7.4, ANSM conditions (RS^−^/Au(III) > 2:1). The experimental conditions are: (i) pH 7.4, 2.5 μM proteins+ 140 mM 2-ME (for 20 min), 0.5 mM HAuCl_4_ (for 20 min), 1 mM NaBH_4_; (j) pH 7.4, 2.5 μM proteins + 60 mM 2-ME, 0.5 mM HAuCl_4_, 30 mM D-P, 2 mM NaBH_4_. (k) 2.5 μM proteins pre-fixed with 1% glutaraldehyde (GA) for 30 min followed by exactly same conditions as (j). (l-q) EM images of the samples in (j), TF is not shown here, but it is similar to BSA. The EM images for (i) and (k) are shown in **Supplementary Fig.6** and **Supplementary Fig.7**, respectively.

To examine the role of RS^−^ in BSM, we performed a series of experiments at neutral pH that tuned the concentrations of two RSH compounds, 2-mercaptoethanol (2-ME) and D-penicillamine (D-P), **(Fig.2a-d)**. These experiments revealed that AuNPs are only formed with RS^−^/Au ratios < 2 (the presence of AuNPs was indicated by solution colors ranging from purple to brown). When the RS^−^/Au ratio ≥ 2, no AuNPs were formed, and the solutions remained colorless. These results imply that a striking transition is occurring at an RS^−^/Au ratio equal to 2; which perfectly fits the calculated RSH concentrations for generating 1 mM RS^−^: 156 mM of 2-ME (pKa 9.64) at pH 7.45, or 40 mM of D-P at pH 6.5, or 6 mM of D-P (pKa 8.1) at pH 7.4 **(Fig.2b-d;)**. In the mixtures of HAuCl_4_ and 2-ME, [RSAu(I)]_*n*_ polymers formed as white precipitates under RS^−^/Au < 2:1 conditions; while a clean solution formed at RS^−^/Au ≥ 2:1 conditions **(Supplementary Fig.2a)**. Similar results were observed in the mixtures of HAuCl_4_ and D-P, except that the color of the precipitates appeared as light brown with an RS^−^/Au ratio < 2:1 **(Supplementary Fig.2b)**. D-P is a stronger gold chelating ligand than 2-ME^12,32^, it can dissolve the precipitates formed in a series of 2-ME and HAuCl_4_ mixtures **(Supplementary Fig.2c)**. The further ligand-competing experiments with fix amino acids for AuNPs synthesis revealed the gold chelating orders (strong to weak): COOH and NH_2_ groups < the thiol group of 2-ME < the thiol of cysteine < the thiol of D-P **(Supplementary Fig.3)**. These results demonstrated that the properties of gold chelating ligands (*e.g*., pKa, chelating strength), and the Au(I)-thiolate polymers formed in the mixture of HAuCl_4_ and RSH under certain pH have essential influences on AuNPs synthesis.

To examine the Au(I)-thiolate species present mixture of HAuCl_4_ and RSH with different RS^−^/Au ratios, we performed a series of MALDI-TOF analyses **(Supplementary Fig.4)**. The MALDI-TOF results demonstrated that the 1:1 gold-to-thiolate compounds^30,31^, [RSAu(I)]]_n_ or [RSAu(I)]_n_SR (where n = 2 to 8) were identified under RS^−^/Au < 2:1 conditions; whereas the 1:2 gold-to-thiolate compounds^30^, [RS-Au(I)-SR]_n_, (where n = 2-6), were identified under RS^−^/Au ≥ 2:1 conditions. The 1:1 compounds are presumably responsible for forming the cloudy precipitates via the aurophilic force between two adjacent Au(I) atoms (**Fig.1e1, f**; **Supplementary Fig.2**). The 1:2 compounds, [RS-Au(I)-SR]_n_, are soluble as a clear solution(**Fig.1e3**; **Supplementary Fig.2)**. This RS^−^/Au = 2:1 transition could be interpreted by the reducing reaction, Au(III) + 2RS^−^ = Au(I) + RSSR. The Au(III) precursors are instantly reduced to form soluble species, [RS-Au(I)-SR]_n_, at RS^−^/Au ≥ 2:1. While the Au(III) precursors are gradually reduced by RS^−^ to form [RS-Au(I)]_n_ cyclic loops or [RS-Au(I)]_n_SR zigzag chains, aurophilically sticking together as large polymers at RS^−^/Au < 2:1 conditions. Thus, our results strongly support the hypothesis that the 1:1 Au(I)-thiolate polymers act as auto-nucleation cores for the formation of AuNPs **(Fig.1e1)**. Suppressing the formation of such polymers inhibits the auto-nucleation **(Fig.1e3)**.

### An ANSM-based approach for the synthesis of AuNPs on purified cysteine-rich tags

We initially tested our ANSM-based approach for the synthesis of AuNPs **(Fig.1d-e4)** by adding 2.5 μM of purified MBP proteins containing tags (MBP-2MT and MBP-2AFP) and the control protein (MBP) into 7 tubes with various concentrations of D-P (ranging from 1 mM to 30 mM) at pH 7.4, which were the same conditions used for experiments depicted **Fig.2d (Fig.2e-g)**. Comparing the colors of each sample in **Fig.2d** with those in **Fig.2e-g**, we found: (i) The patterns for the samples with MBP **(Fig.2e)** were pretty much in the same patterns as those without any protein **(Fig.2d)**, but the patterns for samples with tags were completely different **(Fig.2f-g)**; (ii) In the classic BSM conditions (tubes on the left of the dash lines), all samples were brown in color (indicating the formation of auto-nucleated and/or tag-nucleated AuNPs); (iii) For the samples that have achieved ANSM (the tubes on the right of the dash lines), only those samples with cysteine-rich tags (MBP-2MT and MBP-2AFP) were purple to brown in color (an indication of the formation of tag-based AuNPs) **(Fig.2f-g)**, while those samples without protein **(Fig.2d)** or with the control protein **(Fig.2e)** were colorless (indicating no AuNPs formed). Subsequent EM imaging of the AuNPs formed when using D-P as the gold chelating ligand, confirmed the average diameter of the tag-based AuNPs was approximately 2.3 nm **(Supplementary Fig.5)**. Thus, the results of these experiments match our proposed models **(Fig.1e)**, and collectively confirm that our ANSM-based approach for the specific synthesis of AuNPs on genetically encoded tags is feasible **(Fig.1e4)**.

To optimize our ANSM-based approach for the specific synthesis of stable and large AuNPs on our four tags, we selected 7 model proteins for further tests **(Fig.2h)**. Two tightly-packed cysteine-rich proteins, transferrin (TF, 16 Cys) and bovine serum albumin (BSA, 35 Cys), and a maltose-binding protein (MBP, 2 Cys) were chosen as controls. To evaluate tag-based AuNPs synthesis, we tested four MBP variants that were fused to cysteine-rich tags with 11-40 cysteine residues: MBP-2AFP (24Cys), MBP-2MT (40 Cys), MBP-MTn (20 Cys), and MBP-MTα (11 Cys).

First, we used 140 mM 2-ME at pH 7.45 to achieve the ANSM condition (“2-ME approach”) **(Fig.2i)**. All four tags showed very dark brown colors in solution, while the three control proteins (TF, BSA, MBP) were almost colorless. The EM images showed that no AuNPs formed in the control samples but ~5 nm diameter AuNPs formed on the four proteins with tags **(Supplementary Fig.6)**. Second, we further optimized a protocol for synthesizing stable and large AuNPs on the cysteine-rich tags using the synergistic combination of 2-ME and D-P (“2-ME/D-P approach”) **(Fig.1e; Fig.2h, j, l-q)**. This 2-ME/D-P approach showed similar patterns as those of 2-ME approach, but all had relatively less intense colors **(Fig.2i-j)**. The EM images of these proteins with the 2-ME/D-P approach for AuNPs synthesis **(Fig.2(l-q))** showed that the AuNPs had average diameter ~3-4 nm and were well-dispersed. Finally, we used the optimized 2-ME/D-P protocol to test the aldehyde-resisting abilities of the tags **(Fig.2k)**. The color patterns for fixed and the unfixed groups were quite similar, except the three control proteins showed weaker brown colors that were likely caused by fixatives **(Fig.2k)**. The EM images confirmed that the four tags chemically fixed could form AuNPs; very little background noise was present in the three control samples **(Supplementary Fig.7)**. These results showed that the tags have some aldehyde-resisting capabilities at least in the oxidized states. Interestingly, for a given HAuCl_4_ concentration, we found that the average sizes and size-distributions of the AuNPs depended primarily on the RSH type (2-ME or D-P) **(Supplementary Fig.8)** and the concentration of the tags, while the number of cysteine residues on a tag was relatively less important **(Supplementary Fig.8-9)**.

We have thus validated that the 2-ME/D-P protocol can be used for the synthesis of EM-visible AuNPs on the four tags with high stabilities and good dispersity. Notably, we found that the three untagged control proteins (MBP, TF, BSA) did not form any AuNPs, even if they contained multiple cysteine residues, presumably due to their tightly-packed structures. Therefore, it appears that other cysteine-containing proteins appear unlikely to interfere with the EM-based detection of labeled tags, at least under the relatively mild synthesis conditions used here. It is important to note that water-soluble thiol ligands other than 2-ME can, in principle, be combined with D-P for ANSM synthesis of AuNPs on tags.

### EM visualization of fusion proteins at single-molecule counting level

To assess the oligomerization states of the AuNPs on tags, we performed gel filtration analysis of the tags before and after AuNPs synthesis. The FPLC curves of MBP, MBP-2AFP, and MBP-2AFP with AuNPs suggested that they apparently occur primarily in monomer form in solution **(Supplementary Fig. 10A)**. However, the AuNPs synthesized on isolated fusion tags with the ANSM-based 2ME/D-P protocol tended to stick each other and to proteins, thus making visualization of the AuNPs difficult. Therefore, we optimized another ANSM-based TCEP/D-P protocol for synthesizing more stable AuNPs to avoid such aggregation **(Supplementary Fig. 10B)**. The negative staining EM images of MBP and the MBP-tag fusion proteins clearly confirmed that the majority of AuNPs formed on MBP-tag fusion proteins were in monomer form (*i.e*., one AuNP to one MBP-tag), although a few fusion proteins did not form visible AuNPs; the formation of AuNPs may be hindered by protein aggregates **(Supplementary Fig.10B)**. Therefore, our approach should be valid and ready for single-molecule detection. The sensitivities of single-molecule detection of these tags were further validated with cells in the following sections.

### Screening for conditions suitable for ANSM-based synthesis of AuNPs in *E. coli*

We next tested and optimized our ANSM-based AuNPs synthesis method to visualize tag-fused proteins that were expressed in *E. coli* cells **(Fig.3a)**. We tested *E. coli* cells expressing MBP, MBP-2AFP, MBP-2MT, MBP-MTn, or MBP-MTα and found that it was extremely difficult for gold precursors (*e.g.*, HAuCl_4_, AuCl, sodium gold thiomalate, etc.) to penetrate into live cells. We were aware of a previous report which showed that the incubation of MT-expressing *E. coli* cells with AuCl formed a massive number of AuNPs in periplasmic spaces; we have been perplexed by these results, as they implied that, if the AuNPs were really formed on the tags, then the cytosolic proteins would be mainly localized in extracellular spaces. At the very least, the localization of the AuNPs in the previous study implied that AuCl could not efficiently move across membrane barriers.

**Figure 3.**
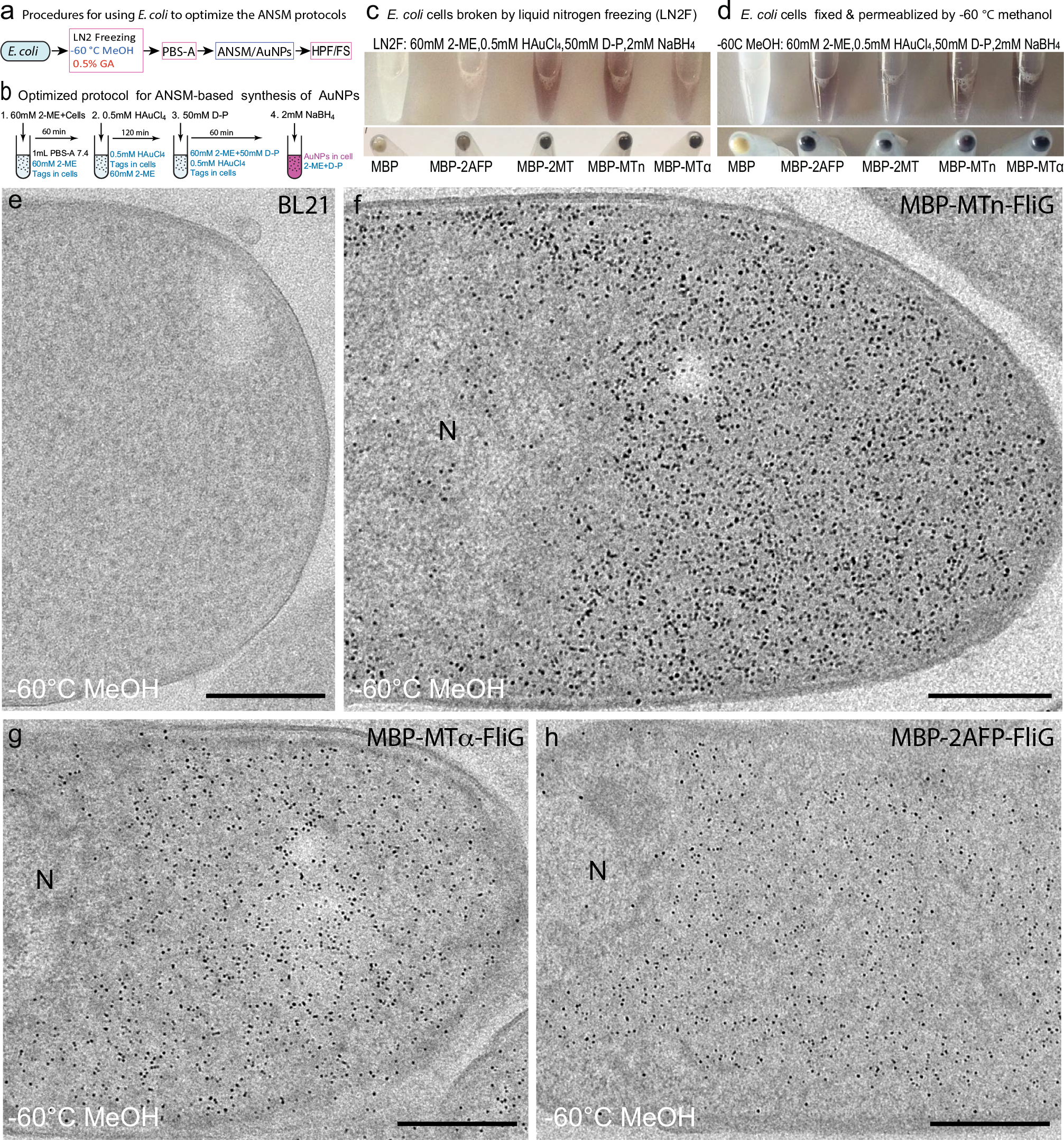
Develop ANSM-based AuNPs synthesis protocols with *E. coli* cells. **(a)** Scheme for developing ANSM-based protocols for synthesizing AuNPs in *E. coli* cells. **(b)** Optimized protocol for ANSM-based synthesizing AuNPs in *E. coli*. **(c)** Liquid nitrogen freezing (LN2F) cracked *E. coli* cells for optimizing the ANSM-based AuNPs synthesizing protocol, the upper row shows the *E. coli* suspension after AuNPs synthesis with an optimized condition as described in **(b)** in 5 mM PBS-A (pH 7.4), the lower row shows the centrifuged pellets, the corresponding EM images were shown in **Supplementary Fig.11. (d)** Using optimized protocol **(b)** for synthesizing AuNPs in the −60 °C methanol fixed and permeabilized *E. coli* cells, the corresponding EM images were shown in **Supplementary Fig.12**. **(e)** the EM image of a cross-section of wild-type *E. coli* (BL21) undergone the same AuNPs synthesizing processes as (d). **(f-h)** the EM images of *E. coli* strains overexpressing MBP-MTn-FliG, MBP-MTα-FliG, and MBP-2AFP-FliG processed with the same procedures as **(d)**.

In an attempt to bypass such penetration barriers, we here simply broke the bacterial cell membranes via liquid nitrogen freezing (LN_2_F), a process that resulted in a marked reduction in the time required to screen optimal conditions for using the ANSM-based AuNP synthesis protocol for *E. coli* cells **(Fig.3a)**. The cell pellets treated with LN_2_F were used to perform a series of AuNPs synthesis reactions by changing the concentrations of thiol ligands and the duration time of incubation based on the optimized ANSM-based AuNP synthesis protocol we had developed for use with purified tags **(Fig.2h)**. Using this screening approach, we obtained an optimized protocol for ANSM-based synthesis of AuNPs on tags expressed in *E. coli* cells **(Fig.3b-c)**. The colors of the five samples in both the solutions and the centrifuged pellets clearly illustrate the striking differences between the MBP (control) and the four tags **(Fig.3c)**. Corresponding EM images of these five samples further confirmed both the formation of AuNPs for the four tags and the near complete lack of any AuNPs in the control cells **(Supplementary Fig.11)**. However, it became obvious that the LN_2_F method was not efficient for breaking cell membranes—the lack of AuNP formation in at least half of the un-cracked cells showed that these membranes remained as barriers for the penetration of gold precursors.

We next applied the optimized ANSM protocol **(Fig.3b)** to *E. coli* cells that were fast fixed and permeabilized with a −60 °C methanol treatment (“−60 °C MeOH”) with the aim of achieving uniform penetration and better preservation of cellular structures (**Fig.3d-h**, **Supplementary Fig.12**, and **Supplementary Fig.13A-E)**. Using this −60 °C MeOH method, there was almost no change in the color of the MBP, while a uniformly dark color was obvious for the four tags **(Supplementary Fig.12)**. EM analysis of control and MBP-MTn-FliG (Note: FliG is the abbreviated form of flagellar motor switch protein), MBP-MTα-FliG, and MBP-2AFP-FliG expressing cells treated with the −60 °C MeOH method showed that there were almost no AuNPs in the control sample **(Fig.3e)** but a great many ~3-6 nm diameter gold particles in the cytosol of the cells expressing each of the three fusion proteins **(Fig.3f-h)**. Notably, almost no AuNPs were formed in the periplasmic spaces of these tag expressing cells **(Fig.3f-h)**, which is strikingly different from the previous report^14^.

Interestingly, the MBP-MTn-FliG, MBP-MTα-FliG, and MBP-2AFP-FliG proteins tended to accumulate near the two poles of some *E. coli* cells; there were many fewer AuNPs in the nucleoid (central) region of cells than at the poles (**Fig.3f-h**, **Supplementary Fig.13B-E)**. Our observation of this specific distribution pattern of AuNPs in the cells we examined suggests that the protocol worked quite well regarding structural preservation, and there was no apparent long-range tag diffusion caused by the treatment. The AuNP distribution patterns of the three fusion proteins in cells were almost the same, although the expression levels differed slightly. The averaged labeling densities of AuNPs (AuNPs/μm^3^) obtained were 44,452, 29,434, and 15,574, respectively, for the MBP-MTn-FliG, MBP-MTα-FliG, and MBP-2AFP-FliG proteins **(Fig.3f-h)**. Thus, we have achieved extremely high labeling densities that never seen before with EM. We further evaluated the efficiency of our AuNPs synthesis protocol by using EM to monitor the number of AuNPs in cells with different expression levels of the GBP-MT fusion protein (altered via different IPTG induction doses), and found that increased expression resulted in increased accumulation of AuNPs **(Supplementary Figure 14)**.

It is thus clear that our protocol is highly specific and efficient for synthesizing AuNPs on the tags of fusion proteins in *E. coli* cells. Importantly, the cellular structures of *E. coli* such as the cell wall, cell membranes, and periplasmic spaces were well preserved by this protocol. We further tested a chemical fixation step prior to the −60 °C MeOH treatment that consisted of a 30 min of fixation with 0.5% glutaraldehyde (GA). We found that this treatment resulted in the similar structural preservation and AuNP distribution patterns as with the −60 °C MeOH protocol alone, with the exception that about 60% fewer AuNPs were formed in samples treated with this GA fixation step **(Supplementary Figure 15)**.

### Visualizing tags expressed in the fission yeast *S. pombe*

After the optimization of our ANSM-based AuNPs synthesis protocol in relatively simple *E. coli* cells, we next extended it to eukaryotic cells in experiments with the fission yeast *S. pombe*. To test whether our protocol is applicable for the specific detection of tag-fused proteins localized to different subcellular locations, we choose three target proteins: Ost4, Nup124, and Sad1, which are expected to be localized, respectively, in the outer membranes of the endoplasmic reticulum (ER) (including the nuclear envelope (NE)), in nuclear pores (NPs), and in the spindle pole body (SPB) **(Fig.4a, b)**. All of these proteins were fused with GFP and GFP-MTn for validating their expression patterns and distributions with fluorescent light microscopy, and all of the fusion proteins showed the expected localization patterns in live *S. pombe* cells **(Fig.4a)**. We initially used Ost4-GFP-MTn expressing cells to screen conditions for optimal protocols for *S. pombe* cells, because the ER is an organelle easily visible in thin sections. For comparing the performance of the three tags (MTn, MTα, and 2AFP) in *S. pombe cells*, we also generated two additional strains expressing Ost4-GFP-MTα and Ost4-GFP-2AFP and used GFP imaging to confirm that these three strains all exhibited similar protein expression patterns (**Fig.4a**, **Supplementary Fig.19A, Supplementary Chart 1)**.

**Figure 4.**
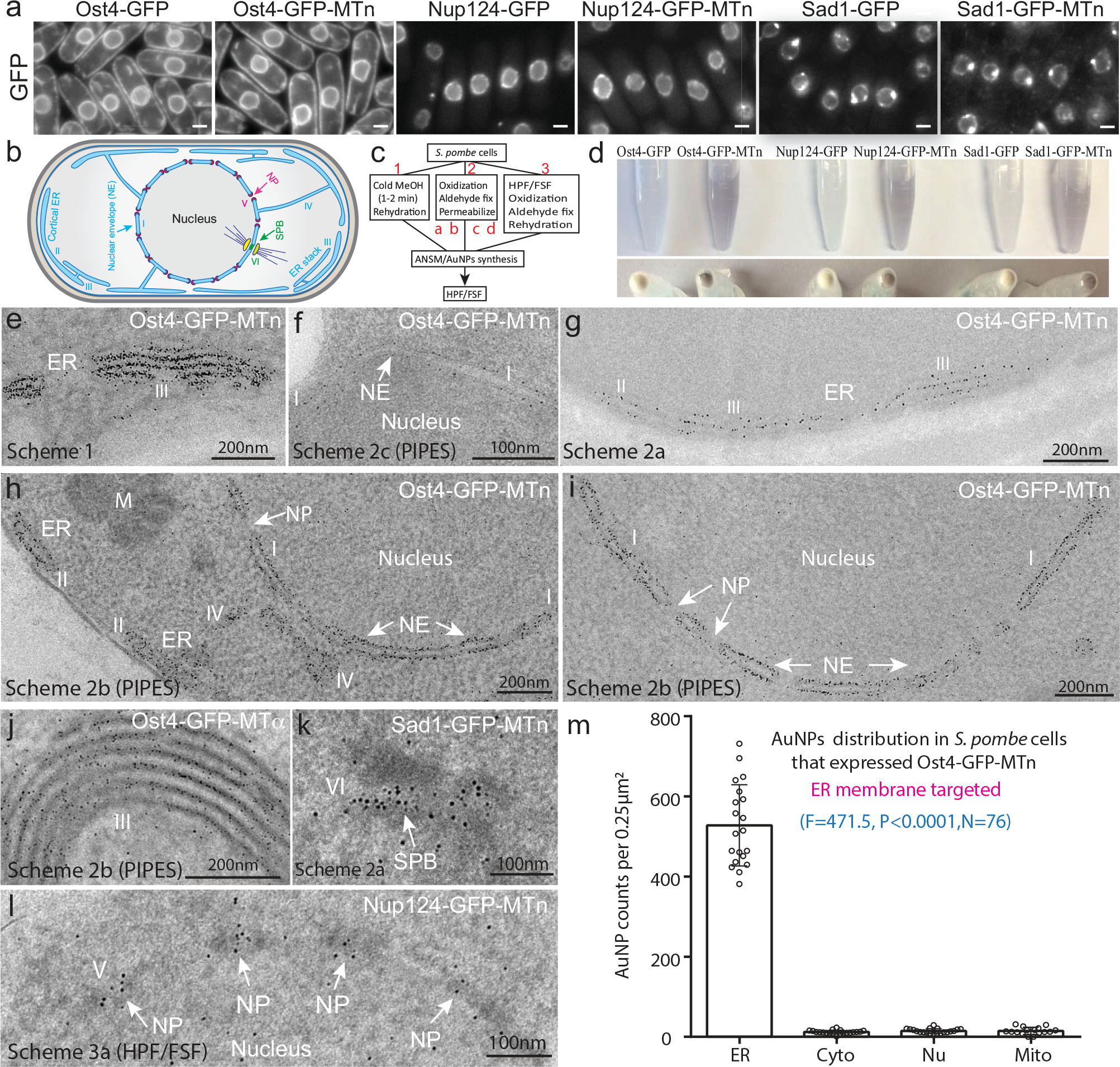
Strategies for achieving single-molecule visualization and cellular structural preservation in *S. pombe* cells. **(a)** The expression patterns of Ost4-GFP, Ost4-GFP-MTn, Nup124-GFP, Nup124-GFP-MTn, Sad1-GFP, and Sad1-GFP-MTn in *S. pombe* observed by live-cell fluorescence imaging (scale bar, 2 μm). The fluorescence images of GFP showed that the fusion proteins were correctly localized to expected subcellular locations: membranes of cortical ER and NE for Ost4, NPs for Nup124, the SPB for Sad1. **(b)** A cartoon shows the NE and cortical ER, NPs, and the SPB in a *S. pombe* cell. The regions of interest are labeled with numbers (I-VI) as references in other figures. **(c)** Procedures for the preparation of *S. pombe* cells for synthesizing AuNPs, three schemes labeled as 1 (“Cold MeOH”), 2 (“Oxidization + Chemical Fixation + Permeabilization”), 3 (“HPF/FSF + Oxidization + Chemical Fixation + Rehydration”) respectively, the Scheme 2 consists of four different sub-schemes (2a-2d), see the **Supplementary Table 3** for details. (d) Synthesis of AuNPs in *S. pombe* cells treated by the “Cold MeOH” protocol with the optimized ANSM-based AuNPs synthesis protocol obtained with *E. coli* cells **(Fig.3b)**, the upper row shows the *S. pombe* cell suspension after AuNPs synthesis and the lower row shows the corresponding centrifuged cell pellets, apparently the colors of those MTn tags expressed cells are much darker than the corresponding control cells. **(e)** A typical EM image showing the unprecedented high labelling densities of AuNPs distributed on the cortical ER (II) and ER stacks (III) in a cell expressing Ost4-GFP-MTn prepared with the Scheme 1. **(f)** EM image showing better membranes preservation of an Ost4-GFP-MTn expressed cell treated with Scheme 2c than that in (e), the AuNPs could be clearly localized to outer membranes of the NE although the labeling densities of AuNPs reduced after the PFA fixation. **(g)** EM image showing the linear distributions of AuNPs along the membranes of cortical ER and ER stacks in a cell expressing Ost4-GFP-MTn treated with the Scheme 2a (with cold MeOH permeabilization), the structural preservation is better than that in (e) but with reduced labelling densities of AuNPs after the GA fixation. **(h-i)** EM images showing the AuNPs distributed along the membranes of NE (I) and ERs (II, IV) in cells expressing Ost4-GFP-MTn treated with Scheme 2b, lack of AuNPs in the nuclear pores (NPs), only a few background AuNPs scattered in cytosols, nuclei and mitochondria (M). (j) high densities of AuNPs formed on ER stacks in cells expressing Ost4-GFP-MTα treated with Scheme 2b (with 0.1% Triton X-100 2 min permeabilization). **(k)** A lot of AuNPs gathered in the SPB (among a dark density region), the specimen was prepared with Scheme 2a. **(l)** AuNPs synthesized in a cell expressing Nup124-GFP-MTn with Scheme 3a, grouped AuNPs formed on the NP (dark regions) with a regular interval along the NE. **(m)** Statistical analysis (one-way ANOVA followed by Brown-Forsythe test) of the AuNPs densities in ER (including NE), cytosol (Cyto), Nucleus (Nu) and mitochondrion (Mito) based on 20 EM images of 90 nm thick sections of *S. pombe* cells expressing Ost4-GFP-MTn (see **Supplementary Chart 1**).

We initially adapted the protocols that we developed with *E. coli* cells **(Fig.3a, b, d)** for *S. pombe* cells. The *S. pombe* cells were treated with −20 °C cold methanol freezing (Scheme 1: “Cold MeOH”) prior to ANSM-based AuNPs synthesis **(Fig.4c-d)**. The cells expressing GFP-MTn tags turned purple, both in solution and in the corresponding centrifuged pellets (an indication for the formation of AuNPs), while the control cells expressing GFP tags did not change color **(Fig.4d)**. An EM image of cells expressing Ost4-GFP-MTn demonstrated that many AuNPs were located, as expected, on ER membranes **(Fig.4e)**. The −20 °C methanol fixation was an effective way to preserve the tags for AuNPs synthesis, as indicated by the unprecedented high labeling density of AuNPs on the ER membranes, although it suffered from some ultrastructural distortions.

We next tried to improve the preservation of the cellular ultrastructures by developing new protocols that are compatible with aldehyde fixation (Scheme 2) or with high-pressure freezing/freeze-substitution fixation (HPF/FSF) that uses aldehyde fixative at low temperature (Scheme 3) (**Supplementary Table 3**, **Fig.4**, **Supplementary Fig.16-19)**. In eukaryotic cells, most cysteine-rich tags in reducing environments such as the cytosol are presumably in unfolded states. The tags will lose their metal-binding ability when their thiol groups react with aldehyde-fixatives. To preserve the tag’s metal-binding activities, we here introduced an oxidizing protection scheme by using 3,3’-dithiodipropionic acid (DTDPA) to oxidize the thiol groups of the tag to aldehyde-inert disulfide bonds prior to the aldehyde fixation; note that the disulfide bonds are reduced back to thiol groups during ANSM-based AuNPs synthesis **(Fig.4c)**.

We conducted a series of experiments to optimize the protocols for *S. pombe* cells, including tests of different DTDPA concentrations, aldehyde fixative types, and fixation strength (both concentration and duration), fixative neutralization, buffer types, permeabilization schemes, and resin types (e.g., SPI-Pon 812, HM20, LR white) (**Fig.4**, **Supplementary Fig.17-18**, **Supplementary Table 3**). Finally, we established the following optimized conditions: 3-5 mM of DTDPA oxidization for 30 min at 4 °C, 0.5% GA or 4% PFA fixation for 30 min at 4 °C, 10% glycine overnight neutralization at 4 °C, 0.1 M PIPES buffer with 0.1 M sorbitol, 1 mM CaCl_2_ and 1 mM MgCl_2_ (PBS also fine but not good for membrane stabilization), 0.1% Triton X-100 2 min permeabilization, and SPI-Pon 812 or HM20 resin

In the EM images of the Ost4-GFP-MTn expressing *S. pombe* cells processed with aldehyde fixation, the AuNPs can be clearly detected on ER stacks and the NE **(Fig.4f-g)** but with a lower density of AuNPs than those without aldehyde fixation **(Fig.4e)**.

However, cellular structures were much better preserved upon aldehyde fixation; AuNPs attached on the outer surfaces of the membranes of the NE can be clearly seen in the specimen prepared with Scheme 2c **(Fig.4f)**. Also, with Scheme 2a, the AuNPs were distributed narrowly along the ER stacks of the specimen **(Fig.4g)**. We also found that for ultrastructural preservation permeabilization treatment with Triton X-100 is superior to treatment with cold methanol.

In the EM images of the Ost4-GFP-MTn and Ost4-GFP-MTα expressing *S. pombe* cells processed with Scheme 2b (PIPES) (5 mM DTDPA oxidization, 0.5% GA fixation, 0.1% Triton X-100 2 min permeabilization), the ER or NE membrane structures were obviously very highly preserved, and the labeling density was quite high **(Fig.4h-j)**. In the EM images of the Ost4-GFP-MTn or Ost4-GFP-MTα expressing *S. pombe* cells processed with Scheme 2b, the AuNPs were very obviously localized to the cytosol-facing outer surface of the ER membranes, and nuclear pores themselves were obviously defined by the tags (**Fig.4h-i**; **Supplementary Fig.16-19)**. In the Ost4-GFP-MTn expressing *S. pombe* cells processed with Scheme 3, the AuNPs were specifically concentrated along ER membranes **(Supplementary Fig.17A(i), 17B(e) & 18c)**. However, the membrane structures were not well preserved on account of the rehydration treatment.

The expression patterns of Ost4-GFP-MTn, Ost4-GFP-MTα, and Ost4-GFP-2AFP in *S. pombe* were confirmed to be quite similar by both live-cell fluorescence (GFP) imaging **(Fig.4a** and **Supplementary Fig.19A)** and EM imaging of the cysteine-rich tags **(Fig.4** and **Supplementary Fig.16-19)**. Protection of thiol groups in the tags by DTDPA oxidization prior to the aldehyde-fixation was found to be critically important to improve the efficiency of AuNPs synthesis in *S. pombe* cells. Compared to samples that were not treated with the tag protection oxidation step, the DTDPA-treated samples had vastly larger numbers of AuNPs **(Supplementary Fig.17A)**. Although a few randomly distributed AuNP background signals were present in both the control (Ost4-GFP) (data not shown) and the Ost4-GFP-MTn cells, the majority of AuNPs were observed in their expected locations. The statistical data (one-way ANOVA followed by Brown-Forsythe test by using the GraphPad Prism 6 Software) of the average AuNPs densities in different organelles from 20 EM images readily confirmed that the AuNPs are highly and specifically localized to the ER (**Fig.4m**, **Supplementary Chart 1)**; while the average background signals of AuNPs in the cytosol, nuclei, and mitochondria each accounted for only about 5% of the total AuNPs present in the sample images (F=471.5; P< 0.0001; N=76).

We then evaluated the performance of MTn tag labeling and the conventional immunogold staining by using two *S. pombe* strains expressing Ost4-GFP or Ost4-GFP-MTn. We first confirmed that these two strains shared similar expression patterns and expression levels using live-cell fluorescence (GFP) imaging **(Supplementary Fig.20(a-b))**. We then conducted three different anti-GFP immunogold staining tests with these two *S. pombe* strains, including thin sections prepared from Tokuyasu cryosectioning, conventional LR white embedded, and lowicryl HM20 embedded specimens **(Supplementary Fig.20)**. Notably, the AuNPs labeling density on the ER and NE membranes of a thin section of Ost4-GFP-MTn expressing cells treated with Scheme 2b (PIPES) was ~10 times higher than the densities observed in the three immunogold staining experiments. It was also notable that the cellular ultrastructures preservation achieved for specimens treated with Scheme 2b (PIPES) was as good as (if not better than) the preservation from either the LR white- or HM20- embedded specimens.

Finally, we validated whether our optimized ANSM-based AuNPs synthesis protocol was effective for the visualization of tags with a more spatially restricted distribution in *S. pombe* cells. Indeed, we were able to visualize the SPB-localized Sad1-GFP-MTn protein **(Fig.4k**; **Supplementary Fig. 18f)** and the NP-localized Nup124-GFP-MTn protein **(Fig 4k**; **Supplementary Fig. 18e)**.

### Specific ER lumen and mitochondrial matrix localization of tags expressed in mammalian cells

To validate the use of our ANSM-based AuNPs synthesis protocol in mammalian cells, we constructed three HeLa cell lines stably-expressing GFP-MTn-KDEL, GFP-2AFP-KDEL, and Mito-acGFP-MTn. The first two cell lines fused GFP-MTn or GFP-2AFP with the classical C-terminal KDEL signal sequence for specific ER lumen localization; the third cell line fused MTn with the known mitochondrial matrix-targeting Mito-tag. We used fluorescent light microscopy to verify that the expression patterns and distributions of the GFP-MTn-KDEL fusion proteins expressed in the HeLa cells were specific to the ER network**(Fig.5a)**. As the KDEL signal sequence should lead to fusion protein accumulation in the ER lumen, we anticipated that successfully synthesized AuNPs would be located mainly inside the cisternae of ER and NE lumens **(Fig.5b)**. We also confirmed that Mito-acGFP-MTn molecules accumulated in mitochondria **(Fig.5c)**. Our design anticipated that the successful synthesized AuNPs would be located mainly in the mitochondrial matrix **(Fig.5d)**, and a series of AuNP synthesis experiments in HeLa cells expressing GFP-MTn-KDEL—using a lightly modified Scheme 2b and 2d (e.g., using 0.1% GA, 3-5 mM DTDPA or 25 mM S-methyl methanethioslfonate (MMTS) as thiol oxidants, 0.7 mM HAuCl_4_)— successfully confirmed that the AuNPs were indeed mainly distributed in the ER (specifically in tubular or expanded cisternae) or NE lumens.

**Figure 5.**
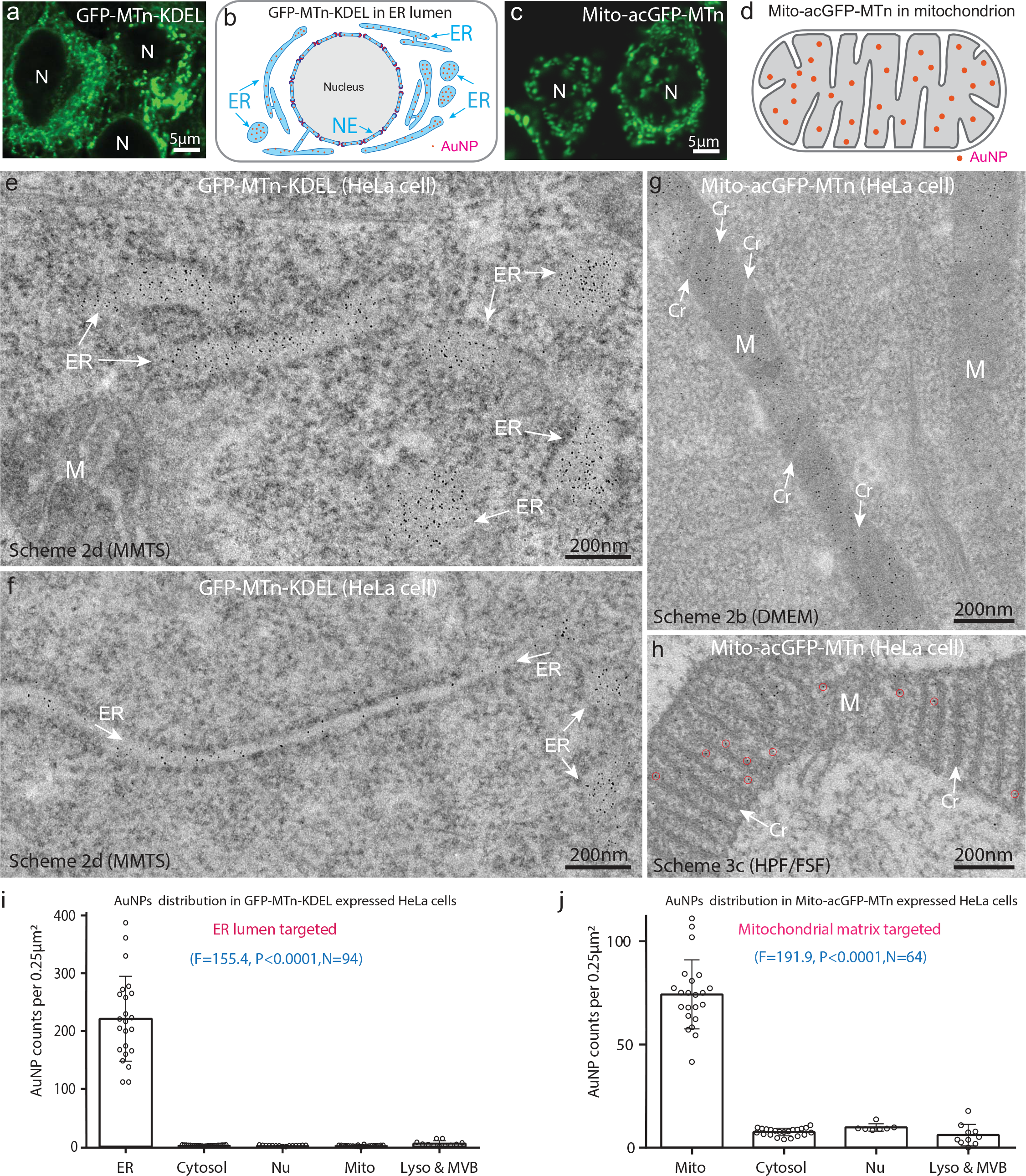
MTn tags specific localized to the lumen of the endoplasmic reticulum and the mitochondrial matrix in mammalian cells. **(a)** The expression patterns of GFP-MTn-KDEL in HeLa cells observed by live-cell fluorescence imaging. **(b)** A cartoon is showing the expected localization of the GFP-MTn-KDEL in a HeLa cell: the corresponding AuNPs synthesized on the tags specific localized to the lumen of ER, NE. **(c)** The expression patterns of Mito-acGFP-MTn in HeLa cells observed by live-cell fluorescence imaging. **(d)** A cartoon is showing the expected localization of the Mito-acGFP-MTn in the mitochondrial matrix in a HeLa cell. **(e-f)** EM image showing the AuNPs specific localized in the lumen of ER network or tubular ER in a GFP-MTn–KDEL expressing cells treated with Scheme 2d (see Supplementary Table 3), and modified ANSM/AuNP synthesis protocol using 0.7 mM HAuCl_4_; hardly seen any AuNPs in the cytosol or mitochondria (M). (g) EM image was showing the AuNPs accumulated in the mitochondrial matrix and almost no AuNPs in the cristae(Cr), in a 90nm thick section of a cell expressing Mito-acGFP-MTn treated with Scheme 2b (DMEM) (see Supplementary Table 3), and the same AuNP synthesis protocol used for (e-f); there are some AuNPs found in cytosol but the AuNPs density is much lower. **(h)** An EM image of a cell expressing Mito-acGFP-MTn treated with Scheme 3c (HPF/FSF) clearly showing AuNPs specific localized in the matrix (*e.g.*, some AuNPs marked with red circles); **(i-j)** Statistical analysis (one-way ANOVA followed by Brown-Forsythe test) of the AuNPs densities in different subcellular organelles, including ER, nuclei (Nu), mitochondria (Mito), lysosomes (Lyso) and multivesicular body (MVB), based on 22-23 EM images of 90 nm thick sections prepared from cells expressing GFP-MTn-KDEL or Mito-acGFP-MTn (see **Supplementary Chart 1**).

Encouragingly, we rarely detected AuNPs in the cytosol or in other organelles like mitochondria (**Fig.5e-f**, **Supplementary Fig.21A-B)**. We found that the nonspecific background signals in cells in suspension were relative higher than such signals in cells cultured on sapphire discs, most likely resulting from preservation during centrifugation **(Supplementary Fig.21E)**. We next optimized a protocol for AuNP synthesis with Scheme 3, which rapidly fixed cells using high-pressure freezing and found that AuNPs localization was highly specific in the ER lumen in the specimen prepared with Scheme 3, although the membrane preservation was substantially degraded, likely from the rehydration treatment **(Supplementary Fig.21C)**. We further tested Scheme 2b (HEPES) for improving membrane preservation by using 0.2 M HEPES buffer with 1 mM CaCl_2_ and 1 mM MgCl_2_ (pH 7.4). The AuNPs were specific localized to ER lumens and it was notable that the membranes of vesicles and mitochondrial cristae were better preserved in these samples **(Supplementary Fig.21D)** than in those preserved using PBS buffer (**Fig.5e-f**, **Supplementary Fig.21A-C)**.

In addition, we tested another HeLa cell line expressing a GFP-2AFP-KDEL fusion with Scheme 2d and Scheme 2b(PBS) and found that the AuNPs exhibited similar distribution patterns as those in GFP-MTn-KDEL expressing cells **(Supplementary Fig.22)**. Finally, we determined the average densities of AuNPs in the ER, cytosol, nuclei, mitochondria, lysosomes, and multi-vesicular bodies in 23 EM images of the GFP-MTn-KDEL expressing cells treated with Scheme 2d and performed the one-way ANOVA statistical analysis (**Fig.5i**, **Supplementary Chart 1)**. The specific AuNPs distribution patterns (P<0.0001) strongly supported that the AuNPs formed in ER lumen resulted from the expected accumulation of GFP-MTn-KDEL molecules. These results clearly demonstrate that the tags and the protocols are suitable for use with mammalian cells.

We further validated whether the above Scheme 2 optimized for ER lumen targeting tags could be applied to visualize tags targeted to other cellular organelles, for instance the mitochondrial matrix. We initially tested Scheme 2d (oxidization with 25 mM MMTS for 5 min at RT) with Mito-acGFP-MTn expressing cells, but found that only a limited number of AuNPs were present in their mitochondrial matrix **(Supplementary Fig.23B(c))**. We next tested a modified Scheme 2b (DMEM) (3 mM DTDPA in DMEM for 30 min oxidization at 4 °C) which allowed better oxidant penetration and oxidization of the tags accumulated in mitochondrial matrix, which did lead to more AuNPs (**Fig.5g**, **Supplementary Fig.23A-B)**: the averaged densities of AuNPs in the mitochondrial matrix were much higher than those in the cytosol, nuclei, lysosomes or multivesicular bodies (MVBs) **(Fig.5j)**.

These results together confirmed that the Mito-acGFP-MTn molecules were translocated from cytosol to mitochondrial matrix as expected, but we found that the membrane preservation achieved using Scheme 2b(DMEM) was still suboptimal. We therefore explored Scheme 2b (HEPES) and Scheme 2b (PIPES) for better membrane preservation and found that although both schemes actually provided excellent mitochondrial membrane and cristae preservation; however, only a very limited number of AuNPs formed in mitochondrial matrix of cells treated with these schemes (most likely owing to poor penetration across two layers of PIPES- or HEPES-stabilized mitochondrial membranes) **(Supplementary Fig.23C)**. Finally, we explored new schemes based on high-pressure freezing for rapid fixation of cells, with the aim of achieving better mitochondrial membrane preservation and AuNP labeling efficiency. The AuNPs in samples treated with these schemes were highly specifically localized in the matrix, obviously partitioned by the cristae (**Fig.5h**, **Supplementary Fig.24A-B)**. Thus, we confirmed that these additional schemes enable fine-tuning to achieve visualization of tags in other organelles like mitochondria.

## DISCUSSION

In summary, we have developed a general approach for synthesizing 2-6 nm diameter AuNPs directly on individual cysteine-rich tags as electron-dense labels for single-molecule visualization for cells with electron microscopy. We first discovered an Auto-nucleation suppression mechanism (ANSM) that can be profitably exploited for the highly specific synthesis of AuNPs on the cysteine-rich tags (in principle, it should be easily expanded to other synthetic thiol-rich macromolecules). We then adapted our ANSM AuNP synthesis protocol for use in single-molecule detection applications in *E. coli*, *S. pombe*, and HeLa cells. We have demonstrated that our AuNP synthesis protocol can achieve remarkably higher labeling efficiency than traditional immuno-EM.

By basing our protocol on the previously unknown ANSM that we discovered in the course of our research, we were able to achieve the specific synthesis of large AuNPs on individual cysteine-rich tags that could be easily detected under regular EM magnification. It is known that MT can bind up to 20 Au(I) atoms via its 20 cysteine residues to form gold clusters **(Fig.1d)**^33,34^; these are not visible under regular EM imaging magnifications. Mecogliano and DeRosier first confirmed that concatenated, 3-copy versions of MT tags were visible with EM at high magnification, and they hypothesized that one MT could, in theory, bind up to ~40 gold atoms via an auto-reduction mechanism to form some Au(0) atoms^12,13^. Nevertheless, if one interprets their mass spectrometry data based on the ligand retention binding mode that we present in **Fig.1d**, the number of gold atoms bound to MT is still fewer than 20^33,34^.

Incubation of the MT tag with AuCl salt could lead to the formation of larger AuNPs because of the auto-reduction of the AuCl to Au(0); however, such an approach suffers from auto-nucleation and the generation of many nonspecific background signals^15^, so it is not applicable to cells. Although a study based on that method reported the formation of many AuNPs in the periplasmic spaces of cells, we are of the view that these AuNPs were unlikely to have been formed on the tags^15,17^ (our unpublished data indicated that there were many gold clusters formed in the periplasmic spaces of all of the ampicillin-resistant strains that we tested (*e.g.*, pMAL-C2X) either with or without the expression of MT tags, but not in the wildtype or kanamycin-resistant strains, even with MT tags). Direct incubation of AuCl_3_ with unfixed permeabilized cells could also lead to the formation of some ~1 nm AuNPs in cells^18,19^. However, nonspecific noise, physiological disturbances, and ultrastructure distortions have heavily hindered the practical application of this approach. Morphew *et al.* (2015) tried to use the HAuCl_4_ staining the MT tags in the budding yeast *S. cerevisiae* cells in organic solvents. However, this approach could only contrast the MT array in spindles; it failed for other structures^20^. Hence, it is quite clear that none of the previous approaches for single-molecule visualization in cells are ready for widespread implementation.

Building on the fundamental principle of ANSM that we discovered, we developed an ANSM-based AuNPs synthesis method derived from BSM that can directly synthesize large AuNPs on individual tags. The sizes of these AuNPs can be tuned by altering the tag/Au ratio, the choice of thiolate ligands, and pH values. We estimate that the ~3 nm diameter tag-localized AuNPs are composed of ~100-200 gold atoms (2.5 μM tags mixed with 0.5 mM HAuCl_4_) **(Fig.2j)**. These tag-based AuNPs seem to have a relatively smaller number of gold atoms than classic colloidal AuNPs of the same diameter, which implies that the gold atoms might be packed differently. The 2-6 nm diameter AuNPs from our synthesis method are large enough to enable the identification of the single fusion proteins in cell sections with EM at regular magnifications. For easier visualization of the ~3 nm AuNPs in thin sections of cells, we simply positively stained the sections with 2% UA for contrast membranes, or completely avoided positive staining.

There are several technical challenges that limit our current approach, and that will need to be overcome in follow-up research and development efforts. First, the cysteine-rich tags expressed in the reducing compartments are not in well-folded states and are thus sensitive to the exposure of aldehyde-fixatives. As a result, the cells cannot tolerate high-strength of chemical fixation. Our current solution to the preservation of the thiol groups is the addition of 0.04-0.1 mM of ZnSO_4_ to the culture medium to enhance zinc-mediated folding and/or oxidizing the thiol groups into disulfide bonds with DTDPA or MMTS prior to the aldehyde fixation step. Second, the penetration of the gold precursors across the cell membrane barrier is still inefficient; it requires the permeabilization of the membranes, either via the use of detergents or organic solvents, both of which can cause some ultrastructural distortions. Third, the high-pressure freezing/freeze-substitution fixation (HPF/FSF) treated samples need to be rehydrated for ANSM-based AuNPs synthesis, although additional UA for fixation in combination with an instant rehydration scheme with HEPES buffer could achieve better preservation of membrane structures. Indeed, the rehydration step of our current protocol downgrades some high-resolution structural information. To solve these problems, we might need to design more highly fixative-resistant tags or develop a more efficient thiol-protection strategy that can tolerate high-strength of chemical fixation. We may also need to develop better approaches for preserving ultrastructures that are fully compatible with HPF/FSF.

The background noise in the model systems that we used was relatively low (e.g., < 5%), which was well supported by the statistical data (P<0.0001) resulted from statistical analysis (one-way ANOVA followed by Brown-Forsythe test) by using GraphPad Prism 6 Software (GraphPad, CA, USA) **(Supplementary Chart 1)**. However, there were still some notable nonspecific background signals observed in some samples, for example a small proportion of the AuNPs scattered in the cytosol, nuclei, and mitochondria of Ost4-GFP-MTn expressing *S. pombe* cells. These background signals may represent AuNPs that formed on some unknown endogenous cysteine-rich proteins, a few large and undissolved [Au(I)SR]_n_ polymers, tags that have relocated or undergone diffusion caused by insufficient fixation and/or the specimen processing (e.g., centrifugation). In contrast to the suspension *S. pombe* or HeLa cells collected by centrifugation, we found that the GFP-MTn-KDEL expressing HeLa cells, which were directly grown on sapphire discs without any centrifugation, had much lower background signals for AuNPs in cytosol, nuclei, and mitochondria (Supplementary **Fig.21E, Supplementary Chart 1**). We will in the future need to systematically evaluate and validate the factors leading to the background signals in mammalian cells and other biological model systems. We anticipate that the continued development of this approach will promote the use of these genetically engineered tags for EM as favorite tools for cell biologists.

In conclusion, we have developed a novel reliable method to directly synthesize 2-6 nm AuNPs on individual protein molecules fused to small cysteine-rich tags, enabling the unambiguous EM-based identification of these genetically tagged proteins in cells at the single-molecule level. The basis of this achievement was our discovery of an ANSM and our use of this mechanism to extend BSM methods. This implementation of genetically engineered tags for EM should allow cell biologists to address an enormous range of biological questions at an unprecedentedly high resolution.

## MATERIALS AND METHODS (online)

## Supporting information

Lists of supplementary information

Supplementary Chart 1

## ACKNOWLEDGMENTS

We thank Dr. X-D. Wang for his vision and long-term support for this high-risk exploring project. W.H. would like to thank his wife R.H. constant standing by him for this time-consuming adventure. W. H. would like to thank Dr. David DeRosier for helpful discussions. We are grateful to Drs. X-C. Wang, D-F. Zhao, X-G. Lei, N. Huang, Y. Cai, W. Hunziker, Y-M. Yuan, K. Ye, F-C. Wang, G-S Ou, G-H. Liu, P-Y. Xu., F. Shao, and Y. Rao for their helps and discussions. We thank Dr. D.R. Winge for providing mouse MT-1 plasmid. We are grateful to Dr. J.H. Snyder for his professional editing service. L. Du was supported by grant from a MOST (973 Program: 2014CB849901); W.H. was supported by grants from the MOST (973 Programs: 2011CB812502 and 2014CB849902), and by funding support from Beijing Municipal Government.

## Author Contributions

W.H. conceived the project, designed the experiments and supervised research, initially discovered and conceptualized the ANSM for AuNP synthesis, analyzed the data and wrote the paper; X.J. performed the experiments for developing the ANSM/AuNP synthesis method, optimizing the AuNP synthesizing conditions for both isolated tags and tags expressed in *E. coli* cells, and also the early efforts on developing AuNP synthesis in *S. pombe* cells with “cold MeOH” approach; Z.J. implemented and optimized the protocols for eukaryotic cells and the related wonderful EM work, also perform some the pre-embedding staining work; Y-H.L. implemented helped to design the tags, established HeLa cell lines, performed the molecular biology and immunogold labeling work, prepared GFP-tag-KDEL HeLa cell lines, and the in vitro protein experiments. S.L. conducted the initial development of a freeze-substitution protocol for synthesizing gold particles in *S. pombe* cells and helped to design the cartoon figures. P.Z., X.C., and Y-H. L. performed the molecular biology work with *E. coli* system; L-L.D. established and supervised the *S. pombe* cells and GBP related collaborations; X-M.L. and Y-Y. W. created the *S. pombe* strains (Ost4, Nup124, Sad1) and confocal took the images; Y.L., X.S., Y.T., Y.H., M.L. performed a series of EM-related work; X-B. Q. for part of chemistry related support; S.C. and G. C. for the MADITOF analysis. All authors participated in discussions and data interpretation.

## Instructions for AuNPs synthesis in *E. coli*, *S. pombe* and HeLa cells

### Scheme LN_2_F for *E. coli*

1. Transfer 1.5 mL of *E. coli* cell culture (OD_600_ of ~1.4) to a 1.5mL microtube
2. Centrifuge at 3000 rpm for 2 min and suck away the supernatant
3. Fill another 1.5 mL of *E. coli* cell culture into the same microtube
4. Centrifuge at 3000 rpm for 5 min and suck away the supernatant
5. Immerse the microtube containing the cell pellet into liquid nitrogen (LN_2_) for 30 s
6. Thaw immediately at RT for 5 min
7. Disperse with 1 mL of PBS-A
8. AuNPs synthesis in of PBS-A

Add 4.28 μL 2-ME in the microtube at RT for 1 h
Add 50 μL 10 mM HAuCl_4_, vortex, incubate at 4 °C for 2 h
Add 100 μL 500 mM D-P, vortex, incubate at 4 °C for 1 h
Add 20 μL 100 mM NaBH_4_, vortex, incubate for 5 min
9. Centrifuge 3000 rpm for 5 min to obtain cell pellet
10. HPF/FSF

### Scheme 1 (Cold MeOH) (Note: also cited as “-60 °C MeOH”, “-60°C methanol” etc.)

1. Transfer 1.5 mL of *E. coli* cell culture (OD_600_ of ~1.4) to a 1.5mL microtube
2. Centrifuge at 3000 rpm for 2 min and suck away the supernatant
3. Fill another 1.5 mL of *E. coli* cell culture into the same microtube
4. Centrifuge at 3000 rpm for 5 min and suck away the supernatant
5. Fast inject 1 mL −60 °C methanol into the microtube, disperse instantly by pipetting for 1 min (Note: or using −80 °C, −20 °C, 4 °C methanol instead)
6. Centrifuged at 3000 rpm for 1 min to remove the supernatant
7. Disperse with 1 mL of PBS-A
8. AuNPs synthesis in PBS-A

Add 4.28 μL 2-ME in 1 mL of PBS-A at RT for 1 h
Add 50 μL 10 mM HAuCl_4_, vortex, incubate at 4 °C for 2 h
Add 100 μL 500 mM D-P, vortex, incubate at 4 °C for 1 h
Add 20 μL 100 mM NaBH_4_, vortex, incubate for 5 min
9. Centrifuge to obtain cell pellet
10. HPF/FSF

### Scheme 2a (4 °C methanol) for *S. pombe* and *E. coli*

1. Centrifuge at 2000 rpm × 2 min (*S. pombe*) or 3000 rpm × 5 min (*E. coli*) to obtain cell pellet
2. Oxidize in 1 mL of 3 or 5 mM DTDPA in PBS in 1.5mL microtube for 30 min at 4 °C
3. Add 20 μL of 25% GA (0.5% final concentration) to the tube for 30 min fixation at 4 °C
4. Centrifuge to remove the supernatant
5. Neutralize excess aldehyde with 1 mL of 10% glycine in PBS at 4 °C overnight
6. Centrifuge to obtain cell pellet
7. Permeabilized with 4 °C methanol for 1 min (see Scheme 1 Step 5-6)
8. Wash with 1 mL of PBS-A for 5 min × 3
9. Disperse with 1 mL of PBS-A
10. AuNPs synthesis in PBS-A

For *E. coli* cells:

Add 4.28 μL 2-ME in 1 mL of PBS-A at RT for 1 h
Add 50 μL 10 mM HAuCl_4_, vortex, incubate at 4 °C for 2 h
Add 100 μL 500 mM D-P, vortex, incubate at 4 °C for 1 h
Add 20 μL 100 mM NaBH_4_, vortex, incubate for 5 min
For *S. pombe* cells:

Add 4.28 μL 2-ME in 1 mL of PBS-A at RT for 1 h
Add 50 μL 10 mM HAuCl_4_, vortex; add 100 μL 500 mM D-P, vortex, incubate at 4 °C for 2 h (Note: Adding D-P immediately after HAuCl_4_ is better for penetration)
Add 20 μL 100 mM NaBH_4_, vortex, incubate for 5 min
11. Centrifuge to obtain cell pellet
12. HPF/FSF

### Scheme 2b (PBS or PIPES) for *S. pombe*

1. Centrifuge at 2000 rpm × 2 min to obtain cell pellet
2. Add 1 mL of 3 or 5 mM DTDPA in selected buffers (PBS or PIPES) in 1.5 mL microtube containing cell pellet for 30 min oxidizing at 4 °C
3. Add 20 μL of 25% GA (0.5% final concentration) to the tube for 30 min fixation at 4 °C
4. Centrifuge to remove the supernatant (at 2000 rpm for 2 min)
5. Add 1 mL of 10% glycine in PBS for neutralization at 4 °C overnight
6. Centrifuge to remove the supernatant
7. Wash with PBS 5 min × 3
8. Centrifuge to remove the supernatant
9. Incubate with 1 mL of 0.2 mg/ml zymolyase-20T in PBS for removing cell wall at RT for ~30 min (check under microscope to make sure cells properly digested as dull gray)
10. Permeabilized with 1 mL of 0.1% triton X-100 in PBS for 2 min at RT
11. Wash with 1 mL PBS-A for 5 min × 3
12. Disperse cell pellet with 1 mL PBS-A
13. AuNPs synthesis in PBS-A

Add 4.28 μL 2-ME in 1 mL of PBS-A at RT for 1 h
Add 50 μL 10 mM HAuCl_4_, vortex; add 100 μL 500 mM D-P, vortex, incubate at 4 °C for 2 h
Add 20 μL 100 mM NaBH_4_, vortex, incubate for 5 min
14. Centrifuge to obtain cell pellet
15. HPF/FSF

### Scheme 2b (PBS, PIPES, HEPES) for HeLa cells

1. For cells in suspension, centrifuge at 800 rpm × 2 min to obtain cell pellet;
  For cells cultured on sapphire disc, transferred it to another microtube with forceps

1. Add 1 mL of 3 or 5 mM DTDPA in selected buffers in a 1.5 mL microtube containing cell pellet or sapphire discs for 30 min oxidizing at 4 °C
2. Add 4 μL of 25% GA (0.1% final conc.) to the tube for 5-30 min fixation at 4 °C
3. For cells in suspension: centrifuge to remove the supernatant (at 800 rpm for 2 min); For cells on sapphire discs: simply suck away the solution
4. Add 1 mL of 10% glycine in PBS to the microtube for neutralization at 4 °C overnight
5. For cells in suspension: centrifuge to remove the supernatant; Simply suck away the solution for cells on sapphire discs
6. Permeabilized with 1 mL of 0.1% triton X-100 in PBS (or the selected buffer) at RT for 2 min
7. Wash with PBS-A for 5 min × 3
8. Disperse in 1 mL of PBS-A for cell pellet; For sapphire discs simply replaced with 1 mL of PBS-A in a 1.5 mL microtube
9. AuNPs synthesis in PBS-A

Add 4.28 μL 2-ME in 1 mL of PBS-A at RT for 1 h
Add 80 μL 10 mM HAuCl_4_, vortex; add 80 μL 500 mM D-P, vortex, incubate at 4 °C for 2 h
Add 20 μL 100 mM NaBH_4_, vortex, incubate for 5 min
10. Centrifuge to obtain cell pellet and resuspend to 1 mL of PBS-A, or change solution to 1 mL of PBS-A for cells on sapphire discs
11. HPF/FSF

### Scheme 2b (DMEM + 10% FBS) for HeLa cells

1. For cells in suspension, centrifuge at 800 rpm × 2 min to obtain cell pellet; For cells cultured on sapphire disc, transferred it to another microtube with forceps
2. Add 1 mL of 3 or 5 mM DTDPA in culture medium in a 1.5 mL microtube containing cells on sapphire discs for 30 min oxidizing at 4 °C
3. Add 4 μL of 25% GA (0.1% final conc.) to the tube for 30 min fixation at 4 °C
4. Replace the solution with 1 mL of 10% glycine in PBS for neutralization at 4 °C overnight
5. Permeabilized with 0.1% triton X-100 in PBS at RT for 2 min
6. Wash with PBS-A for 5 min × 3
7. Keep the sapphire discs in a 1.5mL microtube filled with 1 mL of PBS-A
8. AuNPs synthesis in PBS-A

Add 4.28 μL 2-ME in 1 mL of PBS-A at RT for 1 h
Add 80 μL 10 mM HAuCl_4_, vortex, add 80 μL 500 mM D-P, vortex, incubate at 4 °C for 2 h
Add 20 μL 100 mM NaBH_4_, vortex, incubate for 5 min
9. Change the solution to 1 mL of PBS-A
10. HPF/FSF

### Scheme 2c (PIPES/PFA) for *S. pombe*

1. Centrifuge at 2000 rpm × 2 min to obtain cell pellet
2. Add 1 mL of 3 or 5 mM DTDPA in PIPES buffer in a 1.5 mL microtube containing cell pellet 30 min oxidizing at 4 °C
3. Centrifuge to remove the supernatant (at 2000 rpm for 2 min)
4. Add 1 mL of ice-cold 3 or 5 mM DTDPA and 4% PFA in PIPES buffer at 4 °C
5. Centrifuge to remove the supernatant
6. Add 1 mL of 10% glycine in PBS to the microtube for neutralization at 4 °C overnight
7. Centrifuge to remove the supernatant
8. Wash with PBS 5 min × 3
9. Centrifuge to remove the supernatant
10. Remove cell wall with 1 mL of 0.2 mg/ml zymolyase-20T in PBS at RT for about 30 min (check under microscope to make sure cells properly digested as dull gray)
11. Centrifuge to remove the supernatant
12. Permeabilized with 1mL 0.1% Triton X-100 in PBS for 2 min at RT
13. Wash with PBS-A for 5 min × 3
14. Centrifuge to remove the supernatant
15. Disperse in 1 mL of PBS-A in a 1.5mL microtube
16. AuNPs synthesis in PBS-A

Add 4.28 μL 2-ME in 1 mL of PBS-A at RT for 1 h
Add 50 μL 10 mM HAuCl_4_, vortex; add 100 μL 500 mM D-P, vortex, incubate at 4 °C for 2 h
Add 20 μL 100 mM NaBH_4_, vortex, incubate for 5 min
17. Centrifuge to obtain cell pellet
18. HPF/FSF

### Scheme 2d (MMTS) for HeLa cells cultured on sapphire disc

1. Transfer the cells cultured on sapphire discs to a 1.5 mL microtube containing 1 mL of 25 mM MMTS in PBS for 5 min at RT
2. Add 4 μL of 25% GA (0.1 % final conc.) to the microtube for 5 min fixation at RT
3. Replace the solution with 1 mL of 10% glycine in PBS for aldehyde neutralization overnight at 4 °C
4. Replaced the solution with 1 mL 0.1% Triton X-100 in PBS for 2 min permeabilization at RT
5. Wash with PBS-A for 5 min × 3
6. Keep the specimens in 1 mL PBS-A
7. AuNPs synthesis in PBS-A:

Add 4.28 μL 2-ME in 1 mL of PBS-A at RT for 1 h
Add 80 μL 10 mM HAuCl_4_, vortex, add 80 μL 500 mM D-P, vortex, incubate at 4 °C for 2 h
Add 20 μL 100 mM NaBH_4_, vortex, incubate for 5 min
8. Change solution to PBS-A
9. HPF/FSF

### Scheme 3a (HPF/FSF) for *S. pombe*

1. Centrifuge at 2000 rpm × 2 min to obtain cell pellet
2. Remove cell wall with 1 mL of 0.2 mg/ml zymolyase-20T in PBS at RT for about 30 min (check under microscope to make sure cells properly digested as dull gray)
3. Centrifuge at 2000 rpm × 2 min to obtain cell pellet and perform HPF
4. FSF-Rehydration

Load 10-12 HPF specimens (cells frozen in a 0.1 mm deep carrier) into a 2mL polypropylene microtube with 1 mL of frozen acetone containing 3 or 5 mM DTDPA + 3% ddH_2_O (under LN_2_), then perform following FSF procedures:

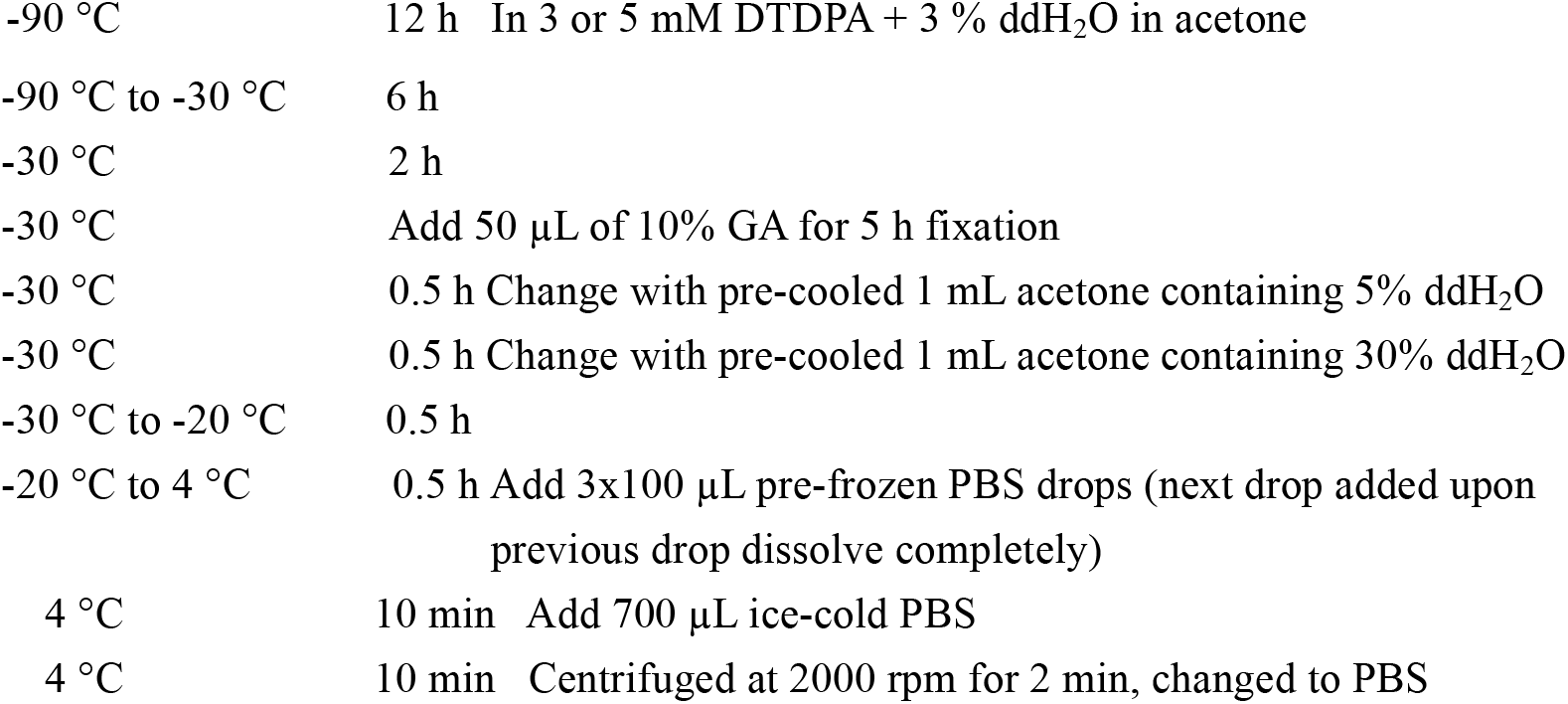
5. Neutralize excess aldehyde with 1 mL of 10% glycine in PBS at 4 °C overnight
6. Wash with 1 mL PBS-A for 5 min × 3
7. Keep the specimens in 1 mL PBS-A
8. AuNPs synthesis in PBS-A

Add 4.28 μL 2-ME in 1 mL of PBS-A at RT for 1 h
Add 50 μL 10 mM HAuCl_4_, vortex, add 100 μL 500 mM D-P, vortex, incubate at 4 °C for 2 h
Add 20 μL 100 mM NaBH_4_, vortex, incubate for 5 min
9. Centrifuge to obtain cell pellet
10. HPF/FSF

### Scheme 3b (HPF/FSF) for HeLa cells cultured on sapphire discs

1. HPF HeLa cells grown on sapphire discs
2. FSF-Rehydration

1-4 specimens loaded into a 2mL polypropylene microtube with 1 mL of frozen acetone containing 10 mM DTDPA + 5% ddH_2_O + 1% methanol under LN_2_

3. Neutralize excess aldehyde with 1 mL of 10% glycine in PBS at 4 °C overnight
4. Wash with PBS-A for 5 min × 3
5. AuNPs synthesis in PBS-A

|  |  |
| --- | --- |
| -90 °C | 12 h |
| -90 °C to -20 °C | 8 h |
| -20 °C | 2h |
| -20 °C to -30 °C | 1 h |
| -30 °C | Add 10 µL of 10% GA (in acetone) for 5 h fixation |
| -30°C | Change to 30% ddH<sub>2</sub>O in acetone, gradually warm up to -20°C within 30 min |
| -20 °C | Change to 50% PBS in acetone, gradually warm up to 4 °C within 1 h |
| 4 °C | Change to 70% PBS in acetone for 10 min |
| 4 °C | Change to PBS for 10 min |

Add 4.28 μL 2-ME in 1 mL of PBS-A at RT for 1 h
Add 80 μL 10 mM HAuCl_4_, vortex, add 80 μL 500 mM D-P, vortex, incubate at 4 °C for 2 h
Add 20 μL 100 mM NaBH_4_, vortex, incubate for 5 min
6. Change solution to PBS-A
7. HPF/FSF

### Scheme 3c (HPF/FSF) for HeLa cells cultured on sapphire discs

1. HPF HeLa cells grown on sapphire discs
2. FSF-Rehydration

1-4 specimens loaded into a 2mL polypropylene microtube with 1 mL of frozen acetone containing 20 mM DTDPA + 5% ddH_2_O + 1% methanol under LN_2_

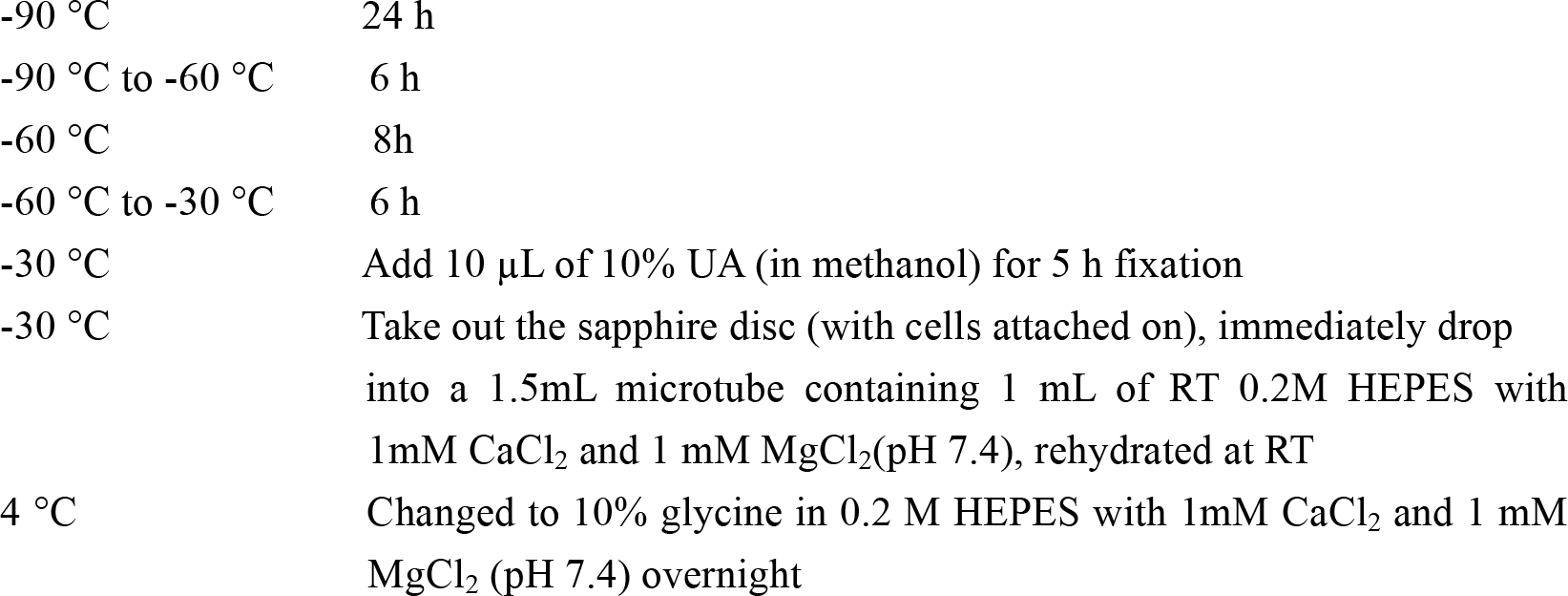
3. Wash with 1 mL PBS-A for 5 min × 3 at 4 °C
4. Keep in a 2mL microtube with 1 mL PBS-A
5. AuNPs synthesis in PBS-A

|  |  |
| --- | --- |
| -90 °C | 24 h |
| -90 °C to -60 °C | 6 h |
| -60 °C | 8h |
| -60 °C to -30 °C | 6 h |
| -30 °C | Add 10 µL of 10% UA (in methanol) for 5 h fixation |
| -30 °C | Take out the sapphire disc (with cells attached on), immediately drop into a 1.5mL microtube containing 1 mL of RT 0.2M HEPES with 1mM CaCl<sub>2</sub> and 1 mM MgCl<sub>2</sub>(pH 7.4), rehydrated at RT |
| 4 °C | Changed to 10% glycine in 0.2 M HEPES with 1mM CaCl<sub>2</sub> and 1 mM MgCl<sub>2</sub> (pH 7.4) overnight |

Add 4.28 μL 2-ME in 1 mL of PBS-A at RT for 1 h
Add 80 μL 10 mM HAuCl_4_, vortex, add 80 μL 500 mM D-P, vortex, incubate at 4 °C for 2 h
Add 20 μL 100 mM NaBH_4_, vortex, incubate for 5 min
6. Change solution to PBS-A
7. HPF/FSF

### Scheme 3d (HPF/FSF) for HeLa cells cultured on sapphire discs

1. HPF HeLa cells grown on sapphire discs
2. FSF-Rehydration

1-4 specimens loaded into a 2mL polypropylene microtube with 1 mL of frozen acetone containing 20 mM DTDPA + 5% ddH_2_O + 1% methanol under LN_2_

3. Wash with 1 mL PBS-A for 5 min × 3 at 4 °C
4. Keep in a 2mL microtube with 1 mL PBS-A
5. AuNPs synthesis in PBS-A

|  |  |
| --- | --- |
| -90 °C | 24 h |
| -90 °C to -60 °C | 6 h |
| -60 °C | 8h |
| -60 °C to -30 °C | 6 h |
| -30 °C | Add 10 µL of 10% GA (in acetone) + 1 µL of 10 % UA (in methanol) for 5h fixation; |
| -30 °C | Take out the sapphire disc (with cells attached on), immediately drop into a 1.5mL microtube containing 1 mL of RT 0.2M HEPES with 1mM CaCl<sub>2</sub> and 1 mM MgCl<sub>2</sub> (pH 7.4), rehydrated at RT |
| 4 °C | Changed to 10% glycine in 0.2 M HEPES with 1mM CaCl<sub>2</sub> and 1 mM MgCl<sub>2</sub> (pH 7.4) overnight |

Add 4.28 μL 2-ME in 1 mL of PBS-A at RT for 1 h
Add 80 μL 10 mM HAuCl_4_, vortex, add 80 μL 500 mM D-P, vortex, incubate at 4 °C for 2 h
Add 20 μL 100 mM NaBH_4_, vortex, incubate for 5 min
6. Change solution to PBS-A
7. HPF/FSF

## ONLINE METHODS

### Plasmid construction and protein expression in *E. coli*

#### MBP and MBP fusion proteins

The MT gene was amplified from pET-3d-MT (a gift from Dr. Dennis R. Winge, University of Utah), which contains the mouse MT-1 gene within the pET-3d vector (Novagen). All the engineered tags, including the MT-derived variants (MTn, MTα, and MTβ), and the tmAFP-derived AFP variants (as listed **Supplementary Table 1**) were synthesized by Beijing Aoke Dingsheng Biotechnology Co. Ltd. The gene of FliG (flagellar motor with protein) was extracted from *E. coli* cells. The ampicillin resistance gene in the pMAL-C2X vector (New England Biolabs) was replaced with a kanamycin resistance gene to make a new vector that we named pMAL-C2X^dAmp/Kana+^ (for avoiding the interference of the beta-lactamase on gold particle synthesis (refer to the discussion section); this vector was used to generate the MBP (maltose binding protein) fusion proteins with the MT or AFP tags. The genes for the MT and AFP variants (**Supplementary Table 1**) were sub-cloned into pMAL-C2X^dAmp/Kana+^ between the *EcoRI* and *HindIII* sites. The 2MT gene was constructed by linking two copies of MT at the *XbaI* site and sub-cloning into pMAL-C2X^dAmp/Kana+^ between the *EcoRI* and *HindIII* sites. For MBP-MTn-FliG, MBP-MTα-FliG and MBP-2AFP-FliG cases, the MTn-FliG, MTα-FliG and 2AFP-FliG genes were sub-cloned into the original pMAL-C2X vector respectively. For cutting off the MBP and affinity purification, a PreScission Protease recognition sequence and a 6xHis tag was added to the N-terminus and C-terminus of those MT and AFP tags respectively. The details of DNA constructs can be found in the **Supplementary Table 2**.

The correctly fused constructs were transformed into BL21 (DE3) competent cells. Bacteria were cultured in LB medium with kanamycin or ampicillin selection at 37 °C; When an OD_600_ of was 0.8 reached, the growth temperature was lowered to 22 °C, and the culture medium was supplemented with 0.04 mM ZnSO_4_ and 0.5 mM isopropyl-b-D-thiogalactopyranoside (IPTG, Sigma) to induce protein expression. After 2.5 h of induction, cells were collected and used for gold particle synthesis. For protein purification, the induction time was adjusted to overnight.

#### Protein purification

MBP and all MBP fusion proteins (including MBP-2MT, MBP-MT, MBP-MTn, MBP-MTα, MBP-MTβ, MBP-2AFP, MBP-MTn-FliG) were expressed in the *E. coli* strain BL21 (DE3) and purified using a standard MBP purification protocol (New England Biolabs), the proteins fused with 6xHis tags were further purified with a Ni-NTA column (Qiagen). For further analysis, purified proteins were concentrated by ultrafiltration and diluted in standard phosphate buffered saline (PBS) (137 mM NaCl, 2.7 mM KCl, 10 mM, Na_2_HPO_4_, 2 mM KH_2_PO_4_, pH 7.4).

### Mammalian cell lines and cell culture

The detailed information of the constructed plasmids for preparing the stably transfected HeLa cell lines were listed in **Supplementary Table 2**. We established three HeLa cell lines for stable expression of GFP-MTn-KDEL, GFP-2AFP-KDEL, and Mito-acGFP-MTn. The pcDNA3.1 plasmids were linearized by *Hind III* (NEB) and *Xho I* (NEB), and then integrated with EGFP and MTn-KDEL or 2AFP-KDEL by Gibson Assembly Master Mix (NEB). The Mito-acGFP sequence was cloned from pAcGFP-Mito plasmid (Clontech), and was added to the 5’-end of MTn gene to construct Mito-acGFP-MTn. The Mito-acGFP-MTn construct was inserted into the *Nhe I* and *Not I* sites of the pcDNA 3.1 plasmid. The recombinant plasmids, pcDNA3.1-EGFP-MTn-KDEL, pcDNA3.1-EGFP-2AFP-KDEL, and pcDNA3.1-Mito-acGFP-MTn were transfected into HeLa cells by Lipofectamine 3000 (Thermo Fisher Scientific). The transfected cells were selected by supplemented with 500 μg/mL G418 (GIBCO) after maintaining in standard cell cultural medium for 24 h. After 1-2 weeks of high G418 selection, the positive colonies were obtained. The selected stable expression cells were cultured under selective growth conditions with 200 μg/mL G418.

The HeLa cells stably expressing GFP-MTn-KDEL, GFP-2AFP-KDEL, Mito-acGFP-MTn were maintained in Dulbecco’s modified Eagle medium (DMEM) supplemented with 10% fetal bovine serum (FBS) and 200 μg/mL G418 at 37°C 5% CO_2_ incubator. All the above reagents were purchased from GIBCO.

For EM experiments, cells were grown either directly in 35 mm cell culture dishes, or on 3 mm × 0.16 mm sapphire discs (Beijing Wulundes Biotech Ltd., or Engineering Office of M. Wohlwend GmbH) placed on the bottom of a 35mm cell culture dish or a well of 6-well cell culture plates. For light microscopy experiments, cells were grown in a 35mm diameter glass bottom dish (No. D141400, Matsunami). Cells on the sapphire discs or harvested from a 35mm cell culture dishes by 0.25% trypsin-EDTA (GIBCO) were used for further EM experiment. It should be noted that all the sapphire discs prior to use were subjected to following treatments: 5 min ultrasonic cleaning in a 2mL polypropylene screw cap microtubes (Cat. 81-0204, Biologix) with 1 mL ethanol, then burnt in the flame of an alcohol lamp inside a laminar flow cabinet, and followed by sterilization under UV light for 30 min.

### *S. pombe* plasmids and cell culture

The fission yeast *Schizosaccharomyces pombe* (*S. pombe*) strains used in this study are listed in **Supplementary Table 2**. Genetic methods for strain construction and composition of EMM medium were as previously described^35^. For the construction of plasmids expressing fluorescent-protein-tagged Ost4, the coding sequence of Ost4 was amplified by PCR using genomic DNA as a template and inserted into modified a pDUAL vector that contains the P1nmt1 promoter and a GFP sequence.

The constructed *P1nmt1*-ost4-GFP plasmid was then digested with *BglII* and ligated with the MTn PCR product to obtain the *P1nmt*1-ost4-GFP-MTn plasmid. *P1nmt*1-ost4-GFP-MTα and *P1nmt*1-ost4-GFP-2AFP constructed similar to the *P1nmt*1-ost4-GFP-MTn. The plasmids were linearized with *NotI* and integrated at the leu1 locus in *S. pombe*. Cells were grown to the mid-log phase in liquid EMM medium in the absence of thiamine at 30 °C.

Other strains that used for expression of Nup124 or Sad1, fused with GFP or GFP-MTn, were prepared similarly to the above Ost4 case (**Supplementary Table 2**).

### Fluorescence Microscopy

*S. pombe* cells were grown to the mid-log phase in liquid EMM medium. Microscopy was performed using a DeltaVision PersonalDV system (Applied Precision) equipped with a YFP filter set (Chroma 89006 set) and a Photometrics CoolSNAP HQ2 camera. Images were acquired with a 1003 1.4 NA objective lens and were analyzed with SoftWoRx software. HeLa cells were grown on the bottom of 35 mm glass dish (No. D141400, Matsunami) maintained with DMEM. Live-cell imaging performed using a Nikon A1-R confocal microscope with a 60x oil immersion objective lens.

### MALDI-TOF (Matrix-Assisted Light Desorption/Ionization Time of Flight) analysis

The samples (*e.g.*, the mixtures of HAuCl_4_ and RSH as described in **Supplementary Fig.4**) were analyzed on an Autoflex II MALDI-TOF/TOF mass spectrometer (Bruker) equipped with a nitrogen pulsed laser. In brief: 1 μL of sample was mixed with 1 μL of 2,5-Dihydroxybenzoic acid matrix solution (Agilent) and spotted on a Bruker MTP 384 massive stainless steel MALDI target. The matrix spots were allowed to dry at room temperature (RT). The spectra were acquired in positive reflector mode with pulsed ion extraction.

### Synthesizing gold nanoparticles on isolated fusion proteins

#### 140 mM 2-ME protocol

2.5 μM concentrations of purified proteins were prepared in ddH_2_O (pH 7.45). To 1 mL protein solution in a 1.5 mL MaxyClear snaplock microtube (MCT150C, Axygen Scientific) was added 10 μL of 2-mercaptoethanol (2-ME, Cat.0482-250ML, Amresco); samples were incubated for 30 min at RT. 50 μL of 10 mM HAuCl_4_ (Cat.4022-1G, Sigma) was then added to the same microcentrifuge tube with immediate rigorous shaking, followed by an additional incubation for 30 min. Finally, 20 μL of 100 mM NaBH_4_ (Cat.452882-25G) (which must be freshly prepared at 4 °C) was added to the tube with immediate gentle shaking for gold nanoparticle synthesis. It should be noted that all the ANSM-based AuNPs synthesis experiments (including this and the following protocols) were simply performed under normal illumination conditions (but avoiding UV light).

#### 2-ME/D-P protocol

2.5 μM concentrations of purified proteins were prepared in PBS-A buffer (1.125 mM NaH_2_PO_4_, 3.867 mM Na_2_HPO_4_, 100 mM NaCl, pH 7.4). To 1 mL protein solution in a 1.5 mL microtube was added 4.28 μL of 2-ME, followed by incubation for 30 min at RT; next, 50 μL of 10 mM HAuCl_4_ was added to the tube, which was immediately shaken well, followed by an additional incubation for 30 min; then, 60 μL 500 mM D-penicillamine (D-P) (Cat.P0147, TCI) was added to the tube and shaken well; finally, 20 μL of 100 mM NaBH_4_ (which must be freshly prepared at 4 °C) was added to the tube with immediate gentle shaking for gold nanoparticle synthesis.

#### TCEP/D-P protocol

1 μM concentrations of purified proteins were prepared in PBS-A buffer. To 0.8 mL protein solution in a 1.5 mL microtube was added 8 μL of 50 mM tris(2-carboxyethyl)phosphine hydrochloride (TCEP·HCl) (Cat.646547, Sigma), followed by incubation for 20 min at RT. Next, 174 μL ddH_2_O, 6 μL of 500 mM D-P and 20 μL of 50 mM HAuCl_4_ was in turn added to another 1.5 mL microtube and vortexed before being transferred into the aforementioned microtube containing the protein that was, after shaking to mix well, incubated for 20 min at 4 °C with very slow rotation on a rotator. Then, 20 μL of 500 mM D-P was added into the microtube, vortexed to mix well immediately. Finally, 20 μL of 100 mM NaBH_4_ (which must be freshly prepared at 4 °C) was added to the tube with immediate gentle shaking for gold nanoparticle synthesis.

### Preparation of cells for AuNPs synthesis

#### Collecting cell pellets for experiments

After the above mentioned 2.5 h IPTG induction, 3 mL of *E. coli* cell culture (OD_600_ of ~1.4) was harvested for each experiment. The first 1.5 mL of cell culture in a 1.5 mL microtube was centrifuged at 3000 rpm for 2 min for removing the supernatant followed by adding the second 1.5 mL culture into the same microtube centrifuged at 3000 rpm for 5 min to obtain the final cell pellet for each sample. The *S. pombe* cells were grown in the EMM to an OD_600_ of ~1.0, and 10 mL of the culture was harvested into a 15mL centrifuge tube for centrifugation at 2000 rpm for 2 min to obtain a cell pellet. The pellet was then dispersed with 1 mL PBS and transferred to a 1.5 mL microtube for centrifugation at 3000 rpm for 5 min to obtain a final cell pellet for each sample.

#### Cracking E. coli cells by liquid nitrogen freezing (LN_2_F)

After the IPTG induction, 3 mL of *E. coli* cell culture (OD_600_ of ~1.4) was harvested as cell pellet for each experiment (described above). The microtube containing the cell pellet was immersed into liquid nitrogen for 30 s prior to immediately thawing at RT for 5 min. Then the cell pellet was dispersed in 1 mL of PBS-A buffer for AuNPs synthesis.

#### Digestion of cell wall of S. pombe cells

10 mL *S. pombe* cell culture (OD_600_ of ~1.0) was harvested as cell pellet. Cell pellet was dispersed into 1 mL PBS buffer and transferred to a 1.5 mL microcentrifuge tube, to which was added zymolyase-20T (Cat.320921, MP) to a 0.2 mg/mL final concentration, followed by incubation for about 30 min at RT for cell wall digestion (Note: stop digestion upon the cells appeared as dull gray color under microscope). Finally, the tube was centrifuged to collect a cell pellet at 2000 rpm for 2 min and then placed on ice for further experiments (*e.g.*, oxidizing, chemical fixation, cold methanol fixation, or high pressure freezing).

#### Oxidization of S. pombe and HeLa cells by 3,3′-dithiodipropionic acid (DTDPA)

2 mL of the selected buffer for specific scheme (see **Supplementary Table 3**; *e.g.*, DMEM, PBS, 0.1 M PIPES or 0.2 M HEPES with 1 mM MgCl_2_ and 1 mM CaCl_2_) was added into a 2 mL microcentrifuge tube (Cat. 81-0204, Biologix) containing 1.2 or 2 mg of DTDPA powder (Cat.D0947, TCI), vortexed to prepare 3 or 5 mM DTDPA oxidization solution. 1 mL of 3 or 5mM DTDPA in selected buffer was added into a 1.5 mL microtube containing the aforementioned cell pellet, dispersed and placed on ice for 30 min. These were centrifuged at 2000 rpm for 2-5 min, and the supernatant was discarded. For cells grown directly on sapphire discs, the disc was simply immersed in 1 mL of 3 or 5 mM DTDPA in PBS buffer (pH 7.4) or in the selected buffer (See **Supplementary Table 3**) in a 2 mL microcentrifuge tube on ice for 30 min, before being picked out with forceps. Alternatively, the culture medium for the sapphire discs was replaced with 2 mL of the DTDPA oxidizing solution in the cell culture dish. For Mito-acGFP-MTn expressing HeLa cells, the PBS buffer used for the previously described oxidization processes was replaced with culture medium (DMEM with 10% FBS). Note that two additional oxidization schemes were explored with the aim of achieving better cell membrane preservation; these were both conducted for 30 min at 4 °C, and used 3 or 5 mM of DTDPA in 0.1 M PIPES buffer with 1 mM MgCl_2_ and 1 mM CaCl_2_ (adjusted to pH 7.2 for HeLa cells; for *S. pombe* cells, an additional 0.1 M sorbitol was added and the pH was adjusted to 6.8) or 0.2 M HEPES buffer with 1 mM MgCl_2_ and 1 mM CaCl_2_ (adjusted to pH 7.4 for HeLa cells).

#### Oxidization of HeLa cells by S-methyl methanethioslfonate (MMTS)

HeLa cells cultured on sapphire discs were simply immersed in 1 mL 25 mM S-methyl methanethioslfonate (MMTS) (Cat.401398, J&K scientific) in PBS buffer (pH 7.4) in a 2 mL microcentrifuge tube for 5 min at RT. Subsequently, an additional of 4 μL of 25% GA (Cat.16320, Electron Microscopy Science) was added for another 5 min at RT. Then, the disc was picked out and immersed into 10% glycine in PBS for neutralization overnight at 4 °C. Another oxidization scheme employed 25 mM MMTS in PBS buffer; this was performed at 4 °C for 50 min and was followed by fixation with 0.5% GA for 30 min at 4 °C.

#### Chemical fixation of E. coli and S. pombe cells

The cell pellet was dispersed and fixed with 0.5% glutaraldehyde (Cat.16220, Electron Microscopy Sciences) in 1 mL of PBS buffer in a 1.5 mL microtube at 4 °C for 15-30 min, and the cells were then washed with PBS 3X for 5 min on ice. Finally, samples were incubated with 10% glycine in PBS for about 1 h or overnight to quench the unreacted aldehydes at 4 °C. Centrifugation at 2000 or 3000 rpm for 2-5 min (Note: 2000 rpm for *S. pombe* cells, 3000 rpm for *E. coli* cells) was used for each buffer change. The final cell pellet was stored on ice for further processing, *e.g.*, cold methanol fixation, high-pressure freezing, or 0.1% Triton X-100 permeabilization. For comparison, another commonly used fixative, paraformaldehyde (PFA) (Cat.A11313, Alfa Aesar), was tested with Ost4-GFP-MTn expressing *S. pombe* cells by replacing the 0.5% GA with 4% PFA and 3 or 5mM DTDPA (freshly prepared) in 100 mM PIPES buffer with 1 mM MgCl_2_, 1 mM CaCl_2_ and 0.1 M sorbitol (pH 6.8) for 1 h fixation at 4 °C.

#### Cold methanol fixation and permeabilization of cells

1 mL of pre-cooled methanol (4 °C, or −20 °C or −60 °C or −80 °C) was rapidly injected into a 1.5 mL microtube containing *S. pombe* cell pellet, or *E. coli* cell. The mixture was pipetted in and out of the tip for about 60 s. The samples were then immediately centrifuged (Note: 2000 rpm for *S. pombe* cells, 3000 rpm for *E. coli* cells); the supernatant was removed and 1 mL of ice-cold PBS buffer was added, and the cells were re-suspended with pipetting. These samples were kept on ice before gold nanoparticle synthesis.

#### 0.1% Triton X-100 permeabilization

1 mL of 0.1% Triton X-100 (Sigma) in PBS buffer (pH 7.4) was quickly mixed with *S. pombe* or *E. coli* cell pellet in a 1.5 mL microcentrifuge tube and dispersed well for 2-5 min, then centrifuged (Note: 2000 rpm × 2 min for *S. pombe* cells, 3000 rpm × 5 min for *E. coli* cells) to remove the solution. The specimen was washed with PBS twice and the cell pellet was placed on ice.

### ANSM-based AuNP synthesis in *S. pombe* or *E. coli* cells

4.28 μL of 2-ME was added to a 1.5 mL microtube containing 1 mL of PBS-A and the pre-permeabilized cell pellet (cold methanol treated, or Triton X-100 permeabilized, or freeze-substituted, etc.). This was vortexed (on a gentle setting) and incubated for 60 min at 4 °C. Next, the gold binding and ligand exchanging was conducted for E. coli and S. pombe cells with slightly different schemes. For *E. coli* cells: 50 μL of 10 mM HAuCl_4_ was added to the microcentrifuge tube, which was gently vortexed, followed by incubation for 120 min at 4 °C; 100 μL of 500 mM D-P was then added to the tube and incubated for 60 min at 4 °C. For *S. pombe* cells: 50 μL of 10 mM HAuCl_4_ was added to the microcentrifuge tube, which was gently vortexed, followed by adding 100 μL of 500 mM D-P to the tube for 120 min of incubation at 4 °C. Finally, 20 μL of 100 mM NaBH_4_ (freshly prepared at 4 °C using pre-chilled ddH_2_O, which must be prepared immediately prior to use) was added to the tube, which was immediately gently shaken for gold nanoparticle synthesis; after a 2-5 min incubation at RT with shaking, samples were centrifuged (Note: 2000 rpm × 2 min for *S. pombe* cells, 3000 rpm × 5 min for *E. coli* cells) to separate the cells from the solution. The cell pellets were then rapidly frozen via high-pressure freezing (below) before further freeze-substitution fixation processing (It should be noted that other conventional fixation and dehydration protocol could also be used if HPF-FSF devices are not available).

### ANSM-based AuNP synthesis in mammalian cells

The protocol we developed for *S. pombe* or *E. coli* cells (as described above) was fine-tuned for AuNP synthesis in HeLa cells. Briefly, 4.28 μL of 2-ME was added into the tube with 1 mL of PBS-A and the cells (cells in suspension or on grown sapphire discs) for 60 min reduction at RT. Then, 80 μL of 10 mM HAuCl_4_ was added and the solution was vortexed rapidly, 80 μL of 500 mM D-P was added and the solution was vortexed rapidly again; the specimen was then placed on a rotor for 2h incubation at 4 °C. Next, 20 μL of 100 mM NaBH_4_ (which must be freshly prepared at 4 °C) was added into the tube and vortexed gently for 3-5 min. Then the cells in suspension were centrifuged at 800 rpm for 2 min for cell pellet collection; the sapphire discs were simply transferred to another microtube filled with ice-cold PBS by forceps. Finally, the cell pellets or sapphire discs with cells were then rapidly frozen via high-pressure freezing (below) before further freeze-substitution fixation processing (It should be noted that other conventional fixation and dehydration protocol could also be used if HPF-FSF devices are not available).

### Optimized protocol for AuNP synthesis in chemically-fixed *S. pombe* cells

10 mL of *S. pombe* cell culture (OD_600_ of ~1.0) was harvested as cell pellet for cell wall digestion. The cell pellet was dispersed by either adding 1 mL of ice-cold PBS buffer with 3 or 5 mM DTDPA or adding 1 mL of ice-cold pH 6.8 0.1 M PIPES buffer containing 3 or 5 mM DTDPA, 0.1 M sorbitol, 1 mM MgCl_2_, and 1 mM CaCl_2_. Samples were then transferred to a 1.5 mL microcentrifuge tube and were oxidized for 30 min at 4 °C on a rotor, before fixation was initiated by directly adding 20 μL 25% GA into the tubes followed by incubation at 30 min at 4 °C (final 0.5% GA). The samples were then centrifuged, and the cell pellet was immediately dispersed into 1 mL of 10% glycine in PBS buffer, which was incubated overnight to quench the unreacted aldehydes. The cells were then washed with PBS buffer (pH 7.4) 3X for 2 min on ice. 1 mL of PBS buffer containing 0.2 mg zymolyase-20T was then added for cell wall digestion for 30 min at RT. The cells were then centrifuged and placed on ice for 2 min during which they underwent 0.1% Triton X-100 permeabilization. After being permeabilized, the cells were washed with 1 mL of PBS buffer 3X for 2 min on ice prior to gold nanoparticle synthesis. Noted that 2000 rpm centrifugation for 2 min was used for each buffer change.

### Freeze-substitution fixation (FSF) based AuNPs synthesis in cells

#### Procedures for FSF-based AuNP synthesis in S. pombe cells (Scheme 3a)

The samples rapidly frozen by high-pressure freezing (see below) were transferred into 2 mL polypropylene screw cap microtubes (Cat. 81-0204, Biologix) with 1 mL of frozen acetone containing 3% ddH_2_O and 3 or 5 mM DTDPA under liquid nitrogen. The tubes were transferred into the specimen chamber of a Leica AFS2 freeze-substitution machine for freeze-substitution processing. The samples were processed by the following FSF-based oxidization, fixation, and rehydration protocol. The temperature of the chamber was kept at −90 °C for 12 h; then gradually warmed up from −90 °C to −30 °C within 6 h; Stayed at - 30 °C for 2 h. Then, 50 μL of 10% GA (Cat.16530, Electron Microscopy Sciences) was added at −30 °C for 5 h fixation (Scheme 3a). The medium was changed with −30 °C pre-cooled 1 mL of 95% acetone containing 5% ddH_2_O and stayed at −30 °C for 30 min.

Then a modified gradual rehydration^36^ procedures were conducted by changing the medium to 700 μL of −30 °C pre-cooled acetone containing 30% ddH_2_O, stayed at −30 °C for 30 min; warmed up to −20 °C within 30 min; Then gradually warmed up to 4 °C within 30 min, during warming processes 3x of 100 μL of PBS frozen drop was added (the drops were pre-frozen on the surface of an aluminum foil in freeze-substitution chamber) into the medium (Note: Next drop was added when the previous drop was dissolved completely). The tubes were transferred to an icebox for further rehydration by adding 700 μL of PBS into the tube for 10 min. Then the tube was centrifuged at 2000 rpm for 2 min to obtain cell pellet at 4 °C. The specimen was rinsed with 1 mL of PBS prior to 1 mL of 10% glycine in PBS overnight incubation on ice. The specimen was rinsed with 3x (at 5 min intervals) PBS-A on ice. Finally, the standard ANSM-based AuNP synthesis in *S. pombe* cells was conducted (described above). Note that typically 10-12 HPF-FSF samples were combined as cell pellet at a time for AuNPs synthesis; *S. pombe* cells were detached from the carriers and took out the carriers from the tube at −30 °C; Other temperature lower than −30 °C for 0.5% GA fixation was also tried, however, the ultrastructure preservation seemed to be inferior to those performed at −30 °C.

#### Procedures for FSF-based AuNP synthesis in mammalian cells

The specimens (cells grown on sapphire discs) rapidly frozen by high-pressure freezing (see below) were transferred into 2 mL polypropylene microtubes with 1 mL of frozen acetone containing 5% ddH_2_O, 1% methanol and 10 mM (Scheme 3a, 3b) or 20 mM (Scheme 3c) DTDPA under liquid nitrogen (see **Supplementary Table 3**). The tubes were transferred into a Leica AFS2 freeze-substitution machine for freeze-substitution processing. The samples were processed by the following FSF-based oxidization, fixation, and rehydration protocols.

Scheme 3b was derived from the Scheme 3a, the FSF-rehydration procedures were performed in following sequences: Stayed at −90 °C for 12 h; Gradually warmed up from - 90 °C to −20 °C within 8 h; Stayed at −20 °C for 2h; Cooled back from −20 °C to −30 °C within 1 h; Added 10 μL of 10% GA at −30 °C for 5 h fixation; Changed the medium to 1mL pre-cooled 30% ddH_2_O in acetone at −30 °C; Warmed up to −20 °C within 30 min and then changed to 1mL pre-cooled 50% ddH_2_O in acetone for 10 min; Gradually warmed up from −20 °C to 4 °C within 1 h; Changed to ice-cold 1mL 70% ddH_2_O in acetone for 10 min at 4 °C; Changed to 1mL ice-cold PBS at 4 °C for 10 min; Neutralized with 10% glycine overnight at 4 °C; Washed with 1 mL PBS-A 3 × 5min at 4 °C. Finally, the standard ANSM-based AuNP synthesis for mammalian cells was conducted for these specimens.

To achieve better membrane ultrastructural preservation, we developed two slightly different Scheme 3c and 3d by introducing 0.1% GA + 0.01% or 0.1% UA as fixative, and using 0.2 M HEPES buffer with 1 mM CaCl_2_ and 1 mM MaCl_2_ (pH 7.4) for one-step instant rehydration. The FSF-rehydration procedures were performed in following orders: Stayed at −90 °C for 12 h; Gradually warmed up from −90 °C to −60 °C within 6 h; Stayed at −60 °C for 8 h; Gradually warmed up from −60 °C to −30 °C within 6 h; Stayed at −30 °C: added 10 μL of 10% UA (pre-prepared in methanol) (Scheme 3c) or 10 μL of 10% GA and 1 μL of 10% UA (Scheme 3d) for 5 h fixation; Immediately transferred the sapphire discs to 1.5mL microtube containing 1 mL of RT 0.2 M HEPES buffer with 1 mM CaCl_2_ and 1 mM MaCl_2_ (pH 7.4), for instant rehydration; Replaced the solution with 1 mL of ice-cold 10% glycine in 0.2 M HEPES buffer with 1 mM CaCl_2_ and 1 mM MgCl_2_ (pH 7.4) for overnight neutralization at 4 °C. Rinsed the specimen with 1 mL of PBS-A for 3 × 5 min. Finally, the standard ANSM-based AuNP synthesis in mammalian cells was performed for these specimens.

### AuNP synthesis on cysteine-rich tags expressed in mammalian cells

#### Protocol for mammalian cells attached to sapphire discs

The HeLa cells expressing GFP-MTn-KDEL, GFP-2AFP-KDEL, and Mito-acGFP-MTn were directly cultured on 3 mm × 0.16 mm sapphire discs to 90% confluence in the culture plate. Then, the cell culture plate was cooled on ice. The cell culture medium in each well was replaced with 2 mL 3 or 5 mM DTDPA in the scheme-specific oxidizing solution (*e.g.*, in cell culture medium, DMEM, PBS, 0.1 M PIPES or 0.2 M HEPES with 1 mM MgCl_2_ and 1 mM CaCl_2_) to oxidize cells at 4 °C on a shaker for 30 min. In some experiments, 25 mM MMTS in PBS buffer was used for oxidization at RT on a shaker for 5 min, or at 4 °C for 50 min. Then, the cells were further fixed by 0.1% GA by directly adding 8 μL of 25% GA into the well with 2 mL oxidizing medium and incubating for 30 min at 4 °C (for the MMTS case, 5 min fixation at RT was used instead) on a shaker. The medium was replaced with 2 mL of 10% glycine in PBS (or the same buffer used for oxidization) for overnight to quench the unreacted aldehydes. Next, the medium in the well was replaced with 2 mL of 0.05-0.1% Triton X-100 in PBS buffer (or the same buffer used for oxidizing) at RT for 1-2 min permeabilization. Then the medium was sucked off and the well was rinsed 3x2 min with 1 mL of PBS-A at 4 °C. Finally, the cells were ready for ANSM/AuNP synthesis in mammalian cells.

#### Protocol for AuNP synthesis in mammalian cells in suspension

The HeLa cells directly grown in a 35mm dish, were harvested with 1x trypsin-EDTA and centrifuged at 800 rpm for 2 min to obtain a cell pellet. Immediately, the cell pellet was cooled on ice and re-suspended with 1 mL of cell culture medium containing 5mM DTDPA for oxidization 30 min at 4 °C on a rotator. Then the cells were fixed by 0.1% GA by directly adding 4 μL of 25% GA into the tube at 4 °C for 30 min. The fixative was replaced with 1 mL of 10% glycine in PBS at 4 °C on rotator overnight for quenching the unreacted aldehydes. The cells were permeabilized with 1 mL of 0.1% Triton X-100 in PBS at RT for 1-2 min, then rinsed with 1 mL of PBS-A for 3x 2 min at 4 °C. Finally, the cells were ready for ANSM/AuNP synthesis in mammalian cells.

### Rapid fixation of cells by high pressure freezing (HPF)

Cell pellet was collected by centrifugation in 1.5 mL microtubes (Note: 2 min at 800 rpm for the HeLa cells in suspension; 2 min at 2000 rpm for *S pombe* cells; 5 min at 3000 rpm for *E. coli* cells). Cell pellet was then used to fill the 0.1 mm or 0.05 mm deep well of an aluminum carrier (Engineering Office of M. Wohlwend GmbH), quickly capped with a 3 mm × 0.16 mm sapphire disc, and loaded into a customized 3.05 mm × 0.66 mm HPF specimen holder for rapid freezing at ~2050 bar pressure for 340-360 milliseconds with a Wohlwend HPF Compact-01 high pressure freezing machine. For cells grown on a 3 mm × 0.16 mm sapphire disc, the disc (with the cells facing the cavity) was quickly capped onto a 0.025 mm deep aluminum carrier that was pre-filled with 1-Hexadecene; this carrier was loaded into the HPF specimen holder for high pressure freezing. After freezing, carriers were unloaded under liquid nitrogen and stored in 2 mL polypropylene microcentrifuge tubes in liquid nitrogen.

### Freeze-substitution fixation (FSF) of the frozen cells

1-3 specimen(s) frozen by HPF were transferred into a 2mL polypropylene microcentrifuge tube containing 1 mL of pre-frozen 1% OsO_4_ (Cat.19110, Electron Microscopy Sciences) and 0.1% UA solution (Cat.22400, Electron Microscopy Sciences) in acetone under liquid nitrogen in a cryobox. The tubes were then transferred into a Leica AFS2 freeze-substitution machine for freeze-substitution processing. The samples were processed using a standard FSF protocol: kept at −90 °C for 24 h; warmed to −60 °C within 5 h, and held at −60 °C for 6 h; then warmed to −20 °C within 3 h and held at −20 °C for 18 h; warmed to 4 °C within 2 h; acetone was changed 3 times (5 min intervals) at 4 °C.

### Resin infiltration, embedding, polymerization, and thin sectioning

The freeze-substituted fixed specimens were infiltrated with SPI-Pon 812 resin (SPI Inc.); then the specimens were embedded in a flat mold or flat-tip BEEM embedding capsule (Tedpella Inc.), and polymerized at 45 °C for 18 h followed by 60 °C for 24-48 h. 90 nm sections were prepared with a Leica UC7 ultramicrotome. The sections were picked on a 200-mesh hexagonal copper grid (G200HEX, Tedpella Inc.) that coated with/without formvar. The sections were used for EM imaging. For evaluation of the visibility of AuNPs in cells in different resins, HM20 and LR White resin infiltration and embedding was also performed for *S. pombe* specimen cells expressing Ost4-GFP-MTn (the procedures were the same as those described in “*Immuno-EM staining plastic sections of S. pombe cells embedded in LR white or HM20 resin*”).

### Immuno-EM staining of GFP expressed in cells

#### Immuno-EM staining cryosection of S. pombe cells prepared with Tokuyasu technique

Preparation of Tokuyasu sections from *S. pombe* cells stably expressing Ost4-GFP was done according to the protocol as previously described^37,38^. 10 mL of *S. pombe* cell culture (OD_600_ of ~1.0) was mixed with 10 mL of double-strength fixative (4% PFA, 0.4% GA with 1.5 M sorbitol in 2x PHEM buffer (pH 6.9, 20 mM PIPES, 50 mM HEPES, 4 mM MgCl_2_, 20 mM EGTA)) in a 50mL centrifuge tube for 15 min initial fixation at 30°C. The initial fixative was then removed by centrifugation at 2000 rpm for 2min. The cell pellet was immediately dispersed with fresh standard strength fixative for 1 h fixation at 30 °C. Cell pellet was collected by centrifugation. The cell pellet was dispersed and transferred into a 1.5 mL microtube containing 1.2 mL PHEM buffer for 3x5 min buffer changes. Then the cell pellet was suspended into 1 mL of HIO_4_ in ddH_2_O for 30 min quenching, followed with 3x5 min PHEM buffer washes. The cell pellet was further quenched with 1 mL of 5% glycine in 1x PHEM for 10 min. The tube was centrifuged to obtained cell pellet, and incubated 1 mL of 12% gelatin in PHEM at 37 °C for 30 min. Next the cells were centrifuged at 3000 rpm for 2 min at 37 °C to collect the cell pellet. Leaving a small volume of gelatin on top of the cell pellet, and mixed the cell pellet well with the small volume of gelatin at 37 °C for a few minutes. The tube was placed on ice and allowed to solidify for 30 min. After gelation, blocks of ~1 mm^3^ were prepared on an ice-cold metal plate under a stereo-microscope. The blocks were transferred to ice-cold 2.3 M sucrose in PHEM buffer and placed on a rotator at 4 °C. After overnight infiltration, the blocks were mounted on specimen holders and plunge frozen in liquid nitrogen. The specimens were collected for the standard *Tokuyasu* cryofixation and cryosectioning with a Leica UC7/FC7 ultramicrotome at −120 °C with 1.4 mm/s cutting speed for preparing 60 nm thin sections. The thin sections were retrieved with 2.3 M sucrose and placed on 200 mesh carbon film coated nickel grids, and stored on 2% gelatin plate on ice.

Further, immuno-EM staining was performed. The gelatin plate was melt at 37 °C for 10 min, then the grids were incubated with droplets of 5% glycine in PBS for 2 × 2 min and blocked for 2 min on droplets of 1% BSA in PBS. After blocking, the sections were incubated for 1.5 h on droplets of 5 μL anti-GFP (goat) antibody (Cat.No.600-101-215, Rockland) 1:100 dilution in block solution. After eight washes of 2 min on droplets of 0.1% BSA in PBS buffer the grids with sections were incubated for 30 min on droplets of 7 μL of Protein-A-Gold 10 nm solution in 1:50 dilution (Cat.No.25285, EMS), washed 6 × 2 min on droplets of PBS and fixed for 5 min with 1% GA in PBS. After 10 washes of 2 min on droplets of ddH_2_O, the grids with sections were transferred to droplets of embedding solution containing 2% methyl cellulose (Sigma) and 0.4% uranyl acetate (Cat. 22400, Electron Microscopy Science) in ddH_2_O on Parafilm on an ice-cold metal plate. After 5 min of incubation, the grids were picked up. Most of the solution on the grids was blotted away with filter paper after which the grids with sections were air-dried for EM imaging.

#### Immuno-EM staining plastic sections of S. pombe cells embedded in LR white or HM20 resin

10 mL of *S. pombe* cell culture (OD_600_ of ~1.0) was harvested as cell pellet for high pressure freezing and freeze-substitution fixation (described above). Briefly, 1-3 specimen(s) frozen by HPF were then transferred into a 2mL polypropylene microcentrifuge tube containing 1 mL of pre-frozen 0.1% UA with 5% ddH_2_O in acetone under liquid nitrogen in a cryobox. The tubes were then transferred into a Leica AFS2 freeze-substitution machine for freeze-substitution processing. The samples were processed using a standard FSF protocol and then embedded in either lowicryl HM20 or LR White resin. A standard FSF protocol was used: kept at −90 °C for 24 h; warmed to −30 °C within 6 h; acetone was changed 3 times (30 min intervals) at −30 °C; Warmed from −30 °C to RT within 10 h, and changed fresh acetone at RT for another 2 h. Infiltrated with a graded series ratio of resin/acetone at 1:2, 1:1, 2:1 and 1:1 for 60 min each, and another 2x pure resin changes for 30 min each. Kept in pure resin and stayed at RT for 24 h. Transferred the specimens with fresh resin to BEEM capsules. For lowicryl HM20 resin embedding case, the capsules were cooled from RT to - 50°C within 1 h; then the specimens were first polymerized with UV light for 24 h; next the specimens were further polymerized with UV light and warmed up from −50 °C to RT within 14 h, maintained for another 10 h. For the LR White resin embedding case, the specimens in capsules were transferred to a vacuum oven polymerized at 45 °C for 24 h. 90 nm sections were prepared with a Leica UC7 ultramicrotome. The sections were picked on a 200-mesh hexagonal nickel grid (Tedpella Inc.) that coated with/without formvar. The sections were then processed using standard immuno-EM staining.

### Electron microscopy imaging

#### EM imaging of AuNPs on isolated tags

The copper grids (G200HEX, Tedpella Inc.) coated with formvar film and a thin layer of carbon film (~10 nm), were glow-discharged for 1 min. Then, 5 μL of the solution containing purified proteins with freshly synthesized AuNPs was deposited on copper grids. After 10 min, the excess solution was blotted away, and the gird was washed twice with ddH_2_O and air dried for EM imaging on an FEI Spirit electron microscope (FEI Corp.) operating at 120kV. The images were recorded with a 4k × 4k 895 CCD camera (Gatan Corp.) under proper defocus values to ensure 2-3 nm AuNPs visible.

#### EM imaging of isolated protein prepared by negative staining

The copper grids (G200HEX, Tedpella Inc.) coated with formvar film and a thin layer of carbon film (~10 nm), were glow-discharged for 1 min. Then, 5 μL of the solution containing purified proteins with freshly synthesized AuNPs was deposited on copper grids. After 10 s, the excess solution was blotted away with a piece of filter paper. Three drops of 10 μL of 2% uranyl acetate (UA) were added on a clean piece of Parafilm placed on a petri dish. The grid (with specimen facing the solution) was floated on first two drop for 5 seconds and the excess solution was blotted away. Then the grid was floated on the third drop for 1 min before the excess staining solution was blotted away and air dried for 10 min. EM images were recorded on an FEI Spirit electron microscope (FEI Corp.) operating at 120kV. The images were recorded with a 4k × 4k 895 CCD camera (Gatan Corp.) under proper defocus values to ensure 2-3 nm AuNPs visible.

#### EM imaging of cells

90 nm thin section on copper grids positive stained with 2% UA for 6 min or without positive staining were directly used for EM imaging on an FEI Spirit electron microscope (FEI Corp.) operating at 120 kV. The images were recorded with a 4k × 4k 895 CCD camera (Gatan Corp.) under proper defocus values. Usually at magnification range from 11 kX to 30 kX are suitable for visualizing AuNPs in cells with proper defocus values to ensure 2-3 nm AuNPs visible.

### Statistical analysis of AuNPs distribution in cells

The number of AuNPs in certain organelle (*e.g.*, ER, cytosol, nuclei, mitochrondria, lysosomes, MVB) and the corresponding area was determined using the particle analysis module of a free software ImageJ (version 1.52g; http://imagej.nih.gov/ij). Then the average density (AuNP counts per 0.25μm^2^) for each organelle was derived accordingly. The average densities of AuNPs in different organelles were determined from 20-23 EM images of the specimens of cells expressing the tags (*e.g.*, *S. pombe* cells expressing Ost4-GFP-MTn, HeLa cells expressing Mito-acGFP-MTn, GFP-MTN-KDEL, and GFP-2AFP-KDEL) (see **Supplementary Chart 1**). Then, ordinary one-way analysis of variance (ANOVA) followed by Brown-Forsythe test was used to determine whether there were any statically significant differences between the means of those average AuNPs densities in different organelles; all the statistical analysis was done with GraphPad Prism 6 software (GraphPad, CA, USA). In all experiments, data were expressed as means ± standard deviation. The details were described in **Supplementary Chart 1**. Values of P lower than 0.05 were considered as significant.

## Information for Supplementary Tables, Chart and Figures

**Supplementary Table 1.**
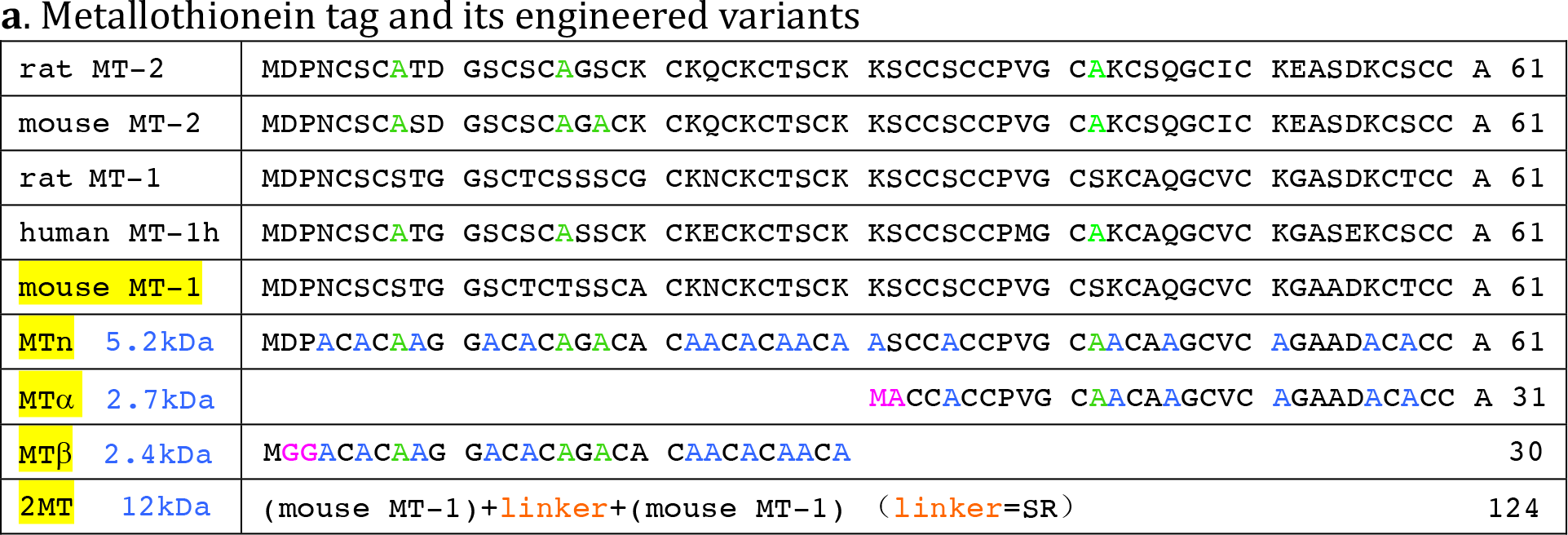
Cysteine-rich tags and their engineered variants. **(a)** *Metallothionein tag (MT) and its engineered variants (MTn, MTα, MTβ and 2MT)* The protein sequences of rat and mouse MT-1, MT-2, and human MT-1h were first aligned for comparison, then the structure of rat MT-2 (PDB code: 4MT2) and the sequence of mouse MT-1 were used for rational design with UCSF Chimera (http://www.rbvi.ucsf.edu/chimera); By keeping the main folding-structure of mouse MT-1 largely unchanged by following the prediction from UCSF Chimera and the sequence variations of those five MT-1 and MT-2, we replaced these aldehyde-reactive residues (K,S,Q,T,N) with structural conservative aldehyde-inert alanine (A) and designed MTn, MTα and MTβ which might help to improve their aldehyde-fixative resistance properties. The expression behaviors of four variants with *E. coli* system, 2MT (two copies of mouse MT-1), MTn, MTα and MTβ were confirmed to be normal (**Supplementary Fig.1**).

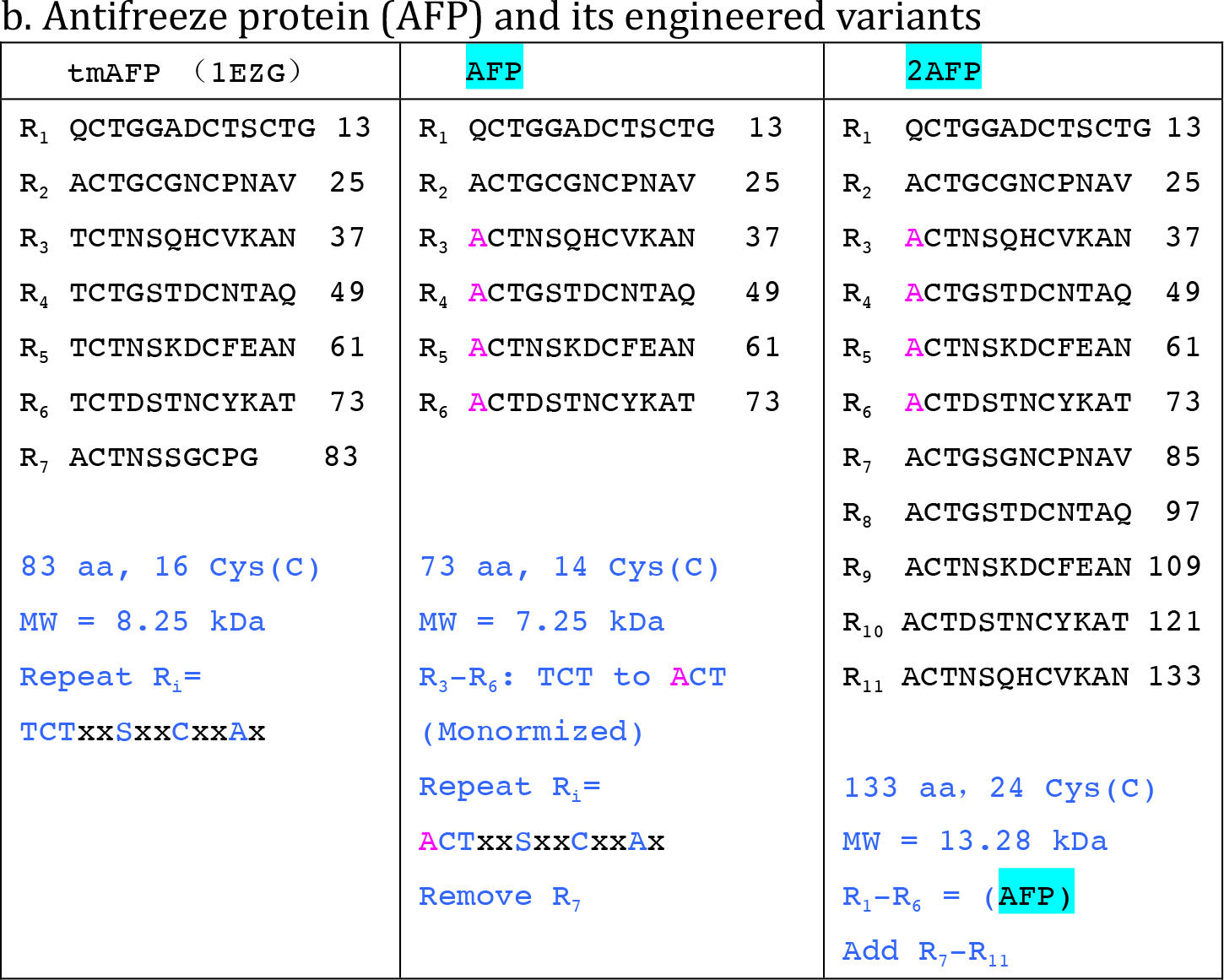
**(b)** *tmAFP and its engineered variants (AFP, 2AFP)* The original 7 repeats of tmAFP (PDB code:1EZG) was used for the design of the AFP and 2AFP engineered variants. The tmAFP is featured by its seven 12 amino acids (12-aa) repeat (e.g.: R_i_ = TCTxxSxxCxxAx) (**Fig.1c**). We first engineered the tmAFP to a monomer variant (AFP) by replacing four residues (T) in R_3_-R_6_ with alanine (A) and remove the last repeat R_7_. Then, we extended AFP to a new variant, named as 2AFP, with eleven R_i_ (12-aa) repeats, by appending extra 5 R_i_ repeats to the 6-repeats AFP tag. And finally confirmed their normal expression behaviors with *E. coli* (**Supplementary Fig.1**).

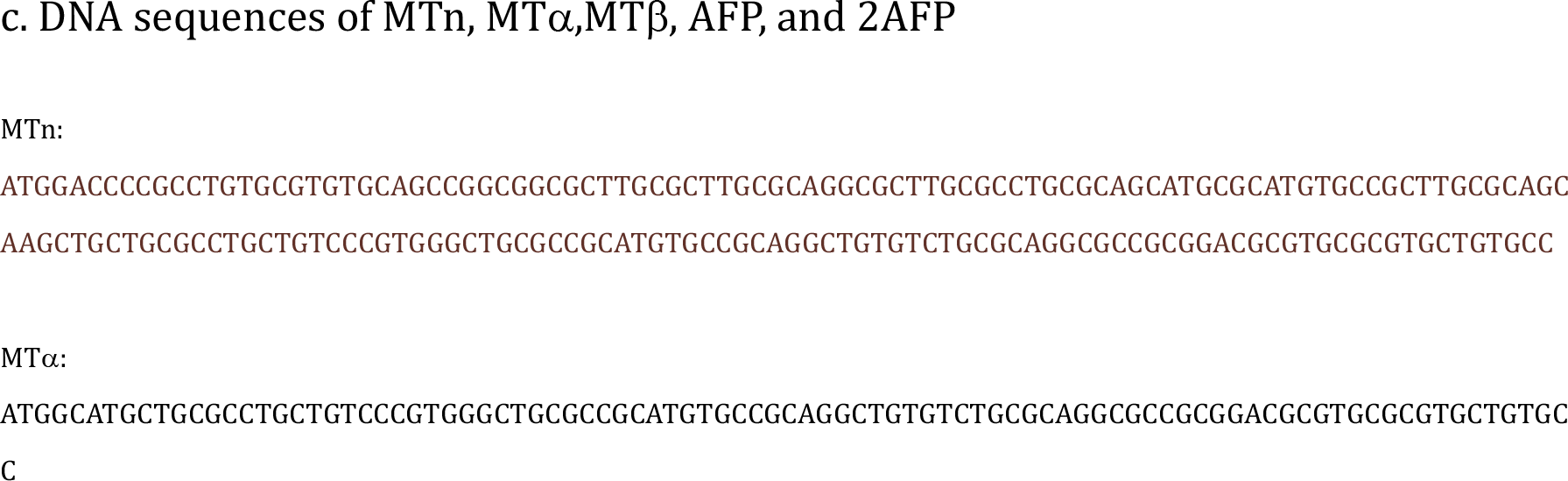
**(c)** DNA sequences of engineered cysteine-rich tags: MTn, MTα,MTβ, AFP, and 2AFP.

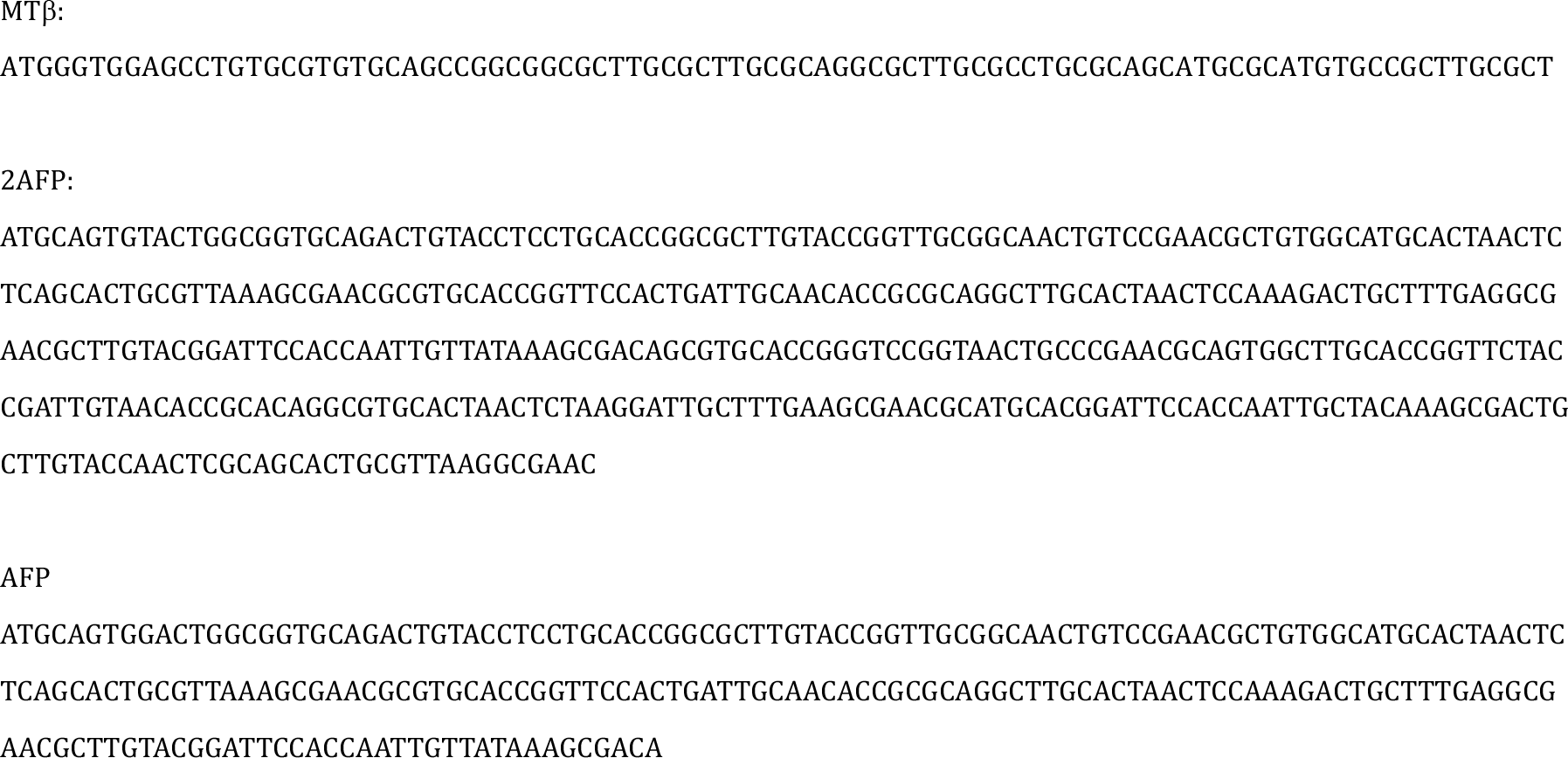

**Table S2.**
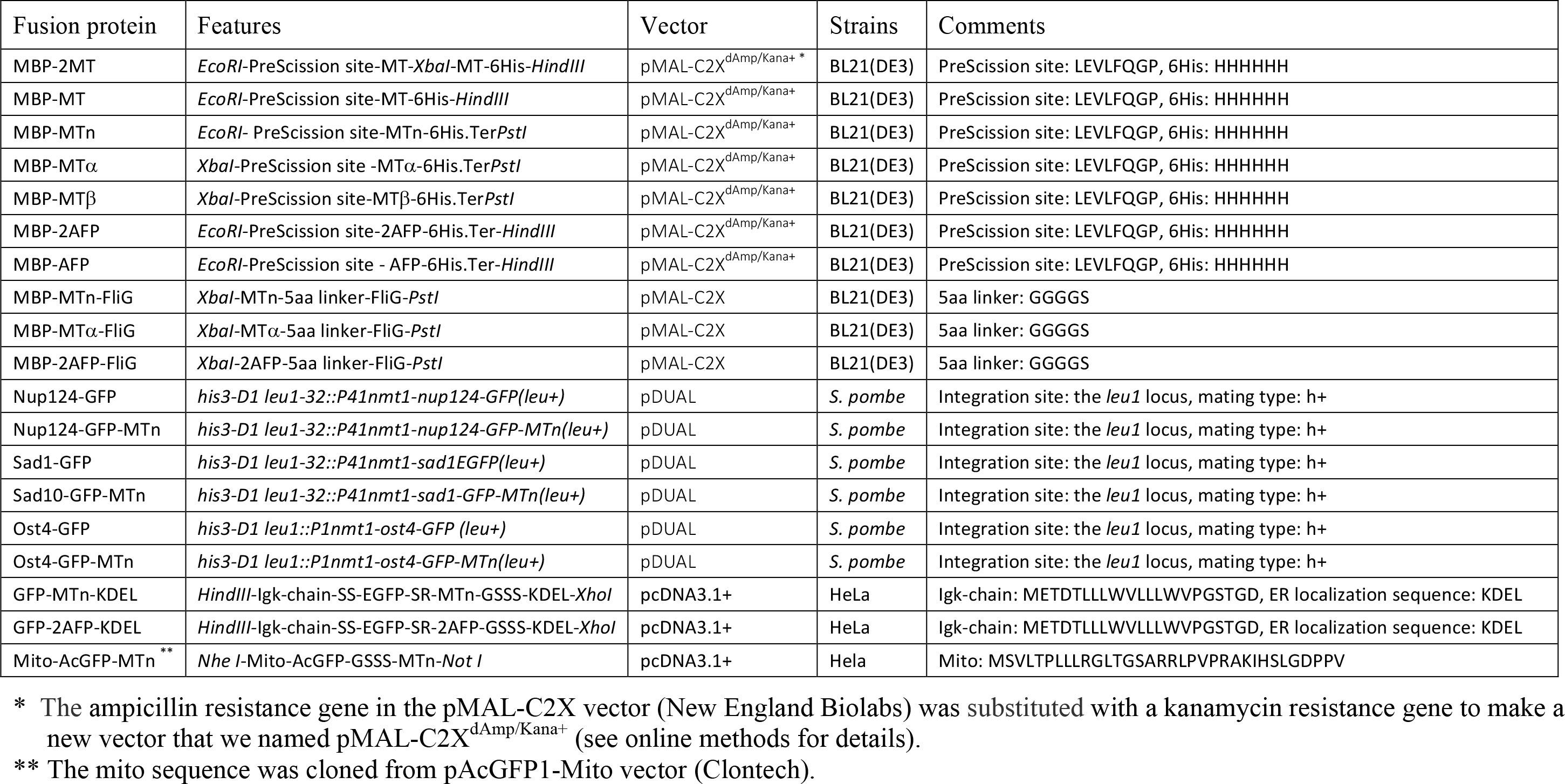
Genetic constructs for prokaryotic (*E. coli*) and eukaryotic (*S. pombe* and HeLa) expression

**Table S3.** Schemes for AuNPs synthesis in cells

| Scheme | LN<sub>2</sub> freezing | Scheme 1<br>Cold Methanol | Scheme 2<br>Oxidization + Chemical Fixation + Permeabilization |  |  |  |  | Scheme 3<br>HPF/FSF + Oxidization + Chemical Fixation + Rehydration |  |  |  |
| --- | --- | --- | --- | --- | --- | --- | --- | --- | --- | --- | --- |
| Scheme index<sup>1</sup> | LN<sub>2</sub>F | Scheme 1<br>Cold MeOH | Scheme 2a | Scheme 2b<br>(PBS, PIPES) | Scheme 2b<br>(PBS, PIPES, HEPES, DMEM<sup>2</sup>) | Scheme 2c<br>(PIPES/PFA) | Scheme 2d<br>(MMTS) | Scheme 3a<br>(HPF/FSF) | Scheme 3b<br>(HPF/FSF) | Scheme 3c<br>(HPF/FSF) | Scheme 3d<br>(HPF/FSF) |
| Cells states | *E. coli* cell pellet | *E. coli* or *S. pombe* cell pellet | *E. coli* or *S. pombe* cell pellet | *S. pombe* cell pellet | HeLa cells in suspension or on sapphire discs | *S. pombe* cell pellet | cell cultured on sapphire disc | HPF fixed *S. pombe* cell pellet | High pressure freezing (HPF) fixed HeLa cells cultured on sapphire discs |  |  |
| Oxidization<sup>3</sup> | n/a | n/a | 5mM DTDPA in PBS 4°C 30min | 5mM DTDPA in PBS or PIPES 4°C 30min | 5mM DTDPA in selected buffers 4°C 30min | 5mM DTDPA in PIPES 4°C 30min | 25mM MMTS in PBS at RT 5min | Oxidized under FSF in 3 or 5mM DTDPA + 3% ddH<sub>2</sub>O in acetone | Oxidized under FSF with 10mM DTDPA + 5% H<sub>2</sub>O + 1% methanol in acetone | Oxidized under FSF with 20mM DTDPA + 5% ddH<sub>2</sub>O + 1% methanol in acetone | Oxidized under FSF with 20mM DTDPA + 5% ddH<sub>2</sub>O + 1% methanol in acetone |
| Fixation<sup>4,5</sup> | n/a | -60°C methanol (or -80°C or -20°C or 4°C) 1min | 0.5% GA in PBS 4°C 30min | 0.5% GA in PBS or PIPES 4°C 30min | 0.1% GA in selected buffer 4 °C 5-30min | 4% PFA in PIPES 4°C 60min | 0.1% GA in PBS at RT 5min | Fixed under FSF with 0.5% GA in acetone at -30°C 5h | Fixed under FSF with 0.1% GA in acetone at -30°C 5h | Fixed under FSF with 0.1% UA in acetone at -30°C 5h | Fixed under FSF with 0.1% GA + 0.01% UA in acetone at -30°C 5h |
| Rehydration | Thawed at RT | changed to 4°C PBS-A | n/a | n/a | n/a | n/a | n/a | Yes, gradual rehydration | Yes, gradual rehydration | Yes, instant rehydration | Yes, instant rehydration |
| Neutralization | n/a | n/a | 10% glycine in PBS overnight at 4°C |  | 10% glycine in selected buffer overnight at 4°C | 10% glycine in PBS overnight at 4°C |  | 10% glycine in PBS overnight at 4°C |  | 10% glycine in HEPES overnight at 4°C |  |
| Permeabilization | n/a, broken already | n/a, permeabilized already | 4°C methanol 1 min | 0.1% Triton X-100 in PBS (or in selected buffer) 2 min |  |  |  | n/a, permeabilized already |  |  |  |
| PBS-A buffer | Changed to PBS-A |  |  |  |  |  |  |  |  |  |  |
| AuNPs synthesis | ANSM-based AuNP synthesis protocols applied (See online methods for details: *for E. coli*, *S. pombe* and HeLa cells, the AuNPs synthesis protocols are slightly different). |  |  |  |  |  |  |  |  |  |  |
| HPF/FSF | After AuNP synthesis, cell pellets or cell cultured on sapphire discs were collected for high pressure freezing and standard freeze-substitution fixation (HPF/FSF) with 1%Os+0.1%UA in acetone |  |  |  |  |  |  |  |  |  |  |
| Resin<sup>6</sup> | SPI-Pon 812 resin |  |  |  | HM20 resin |  | SPI-Pon 812 resin |  |  |  |  |
| Sectioning | 90 nm thin sections were cut for TEM imaging (For clearly visualization of AuNPs, no further UA-lead positive staining applied<sup>7</sup>. Only for some sections of HeLa cells were stained with 2% UA for 6 min, which made the cell membrane slightly better visualized). |  |  |  |  |  |  |  |  |  |  |
| EM imaging | Thin sections of cells were imaged with FEI spirit operating at 120kV, usually at 6500X-21kX magnification. |  |  |  |  |  |  |  |  |  |  |
| Notes | The step by step instructions see the following list, and the actual procedures could be found in the online methods.<br>For HeLa cells expressing Mito-GFP-MTn case, the cells were oxidized and fixed in culture medium (DMEM + 10% FBS).<br>Recommend 5 mM DTDPA (Using 3 mM DTDPA or 5 mM DTDPA did not make notable difference, for some experiments we used 3mM DTDPA).<br>During the GA or PFA fixation, the oxidants (DTDPA or MMTS) were kept in fixation solution; the fixation duration might be adjusted from 5 min to 30 min; 0.1% GA for HeLa cells, 0.5%GA for *S. pombe* or *E. coli*.<br>In the early exploration of the *S. pombe* fixation, 0.5% GA was diluted from 25% GA which was not freshly opened, the effective concentration of GA might be slightly lower than 0.5%.<br>SPI-Pon 812 resin, HM20, LR White resins tested for *S. pombe* cells: SPI-Pon 812 resin and HM20 were confirmed good (HM20 preserving cell membrane better), but all the AuNPs were not visible in LR White resin.<br>Standard UA-lead positive staining can be applied to achieve better image contrast, but it requires high magnification to distinguish the AuNPs from the lead staining background. |  |  |  |  |  |  |  |  |  |  |

**Supplementary Chart 1**

Statistical analysis (one-way ANOVA) of subcellular distribution of AuNPs synthesized in cells expressing cysteine-rich tags targeted to ER or mitochondria

**Supplementary Figure 1**

Expression and purification of MBP and MBP-tag fusion proteins

**Supplementary Figure 2**

Polymeric Au(I)-thiolate precipitates formed in the mixtures of HAuCl_4_ and RSH (2- ME or D-P) at RS^−^/Au(III) < 2:1 conditions

**Supplementary Figure 3**

Gold chelating orders of 6 amino acids, 2-mercaptoethanol (2-ME) and d-penicillamine (D-P)

**Supplementary Figure 4**

Identification of gold-thiolate species in the mixtures of HAuCl4 and RSH by MALDI-TOF analysis

**Supplementary Figure 5**

EM images of AuNPs synthesized on MBP, MBP-2MT and MBP-2AFP with Brush-Schiffrin method performed at ANSM conditions achieved with 30 mM d-penicillamine

**Supplementary Figure 6**

EM images of AuNPs synthesized on 7 proteins with the 2-ME protocol

**Supplementary Figure 7**

EM images of AuNPs synthesized on 1% GA pre-fixed proteins with the ME/D-P protocol

**Supplementary Figure 8**

Diameter distributions of AuNPs synthesized on 4 cysteine-rich tags (2AFP, 2MT, MTn and MTα) with the 2-ME protocol and the 2-ME/D-P protocol

**Supplementary Figure 9**

Diameter distributions of AuNPs synthesized at various concentrations of MBP-MTn at pH 7.45, with 2-ME/D-P protocol

**Supplementary Figure 10A**

AuNPs formed on MBP-2AFP might be mainly in monomer forms and in similar sizes

**Supplementary Figure 10B**

Single-molecule level EM imaging of MBP-tag fusion proteins (MBP-2MT, MBP-MTn, MBP-MTα and MBP-2AFP) subjected to AuNP synthesis followed by negative staining

**Supplementary Figure 11**

Optimization of the conditions for ANSM-based AuNPs synthesis in *E. coli* cells by the disruption of cell membranes with liquid nitrogen freezing (LN2F)

**Supplementary Figure 12**

ANSM-based synthesis of AuNPs on cysteine-rich tags expressed in *E. coli* cells treated by −60 °C methanol

**Supplementary Figure 13**

AuNPs specific distributions *E. coli* cells that expressing MBP-MTn-FliG, MBP-MTα-FliG and MBP-2AFP-FliG

A. AuNPs specific distributions in *the E. coli* cells overexpressing MBP-MTn-FliG
B. AuNPs specific distributions in the *E. coli cells* overexpressing MBP-MTn-FliG
C. AuNPs specific distributions in the *E. coli cells* overexpressing MBP-MTn-FliG
D. AuNPs specific distributions in the *E. coli cells* overexpressing MBP-2AFP-FliG
E. AuNPs specific distributions in the *E. coli* cells overexpressing MBP-MTα-FliG

**Supplementary Figure 14**

The number of AuNPs formed in cells expressing GBP-MT increasing with the induction concentration of IPTG for 4 hours

**Supplementary Figure 15**

ANSM-based synthesis of AuNPs on 0.5% Glutaraldehyde fixed *E. coli* cells

**Supplementary Figure 16**

Synthesis of gold nanoparticles (AuNPs) in *S. pombe* cells processed with 3 mM DTDPA oxidization, 0.5% GA fixation, 10% glycine neutralization, and 4°C methanol permeabilization (Scheme 2a, see Fig.4c and Supplementary Table 3)

**Supplementary Figure 17**

A. Optimization of specimen preparation protocols for visualizing the MTn tags and fine structures in *S. pombe* cells expressing Os4-GFP-MTn: thiol oxidization is essential for protecting the tags for AuNP synthesis.
B. Optimization of specimen preparation protocols for visualizing the MTn tags and fine structures in *S. pombe* cells expressing Ost4-GFP-MTn: fixatives, fixation strength, buffers, resin types.

**Supplementary Figure 18** Localization of GFP-MTn tagged Ost4, Nup124, and Sad1 fusion proteins in *S. pombe* cells by synthesis of AuNPs on MTn tags

**Supplementary Figure 19**

A. Expression patterns of *S. Pombe* strains expressing Ost4-GFP-MTα or Ost4-GFP-2AFP
B. Expression patterns of *S. pombe* strains that expressed Ost4-GFP-MTα

**Supplementary Figure 20** A comparison of the performance of metallothionein tag labeling (ANSM), immunogold staining with Tokuyasu technique and conventional immuno-EM methods

**Supplementary Figure 21**

A. Distributions of AuNPs synthesized on GFP-MTn-KDEL that expressed in HeLa cells
B. Evaluation of the performance of oxidants and the strength of glutaraldehyde fixation for preserving the activities of the MTn tags and fine structures in HeLa cells
C. Distributions of AuNPs synthesized on GFP-MTn-KDEL that expressed in HeLa cells fixed with high pressure freezing
D. HeLa cells expressing GFP-MTn-KDEL oxidized and fixed in HEPES buffer preserved membrane structures
E. Distributions of AuNPs synthesized on GFP-MTn-KDEL that expressed in HeLa cells: the high nonspecific background signals of AuNPs might be caused by the centrifugation

**Supplementary Figure 22**

Distributions of AuNPs synthesized on GFP-2AFP-KDEL that expressed in HeLa cells

**Supplementary Figure 23**

A. Distributions of AuNPs synthesized on mito-GFP-MTn that expressed in HeLa cells
B. Distributions of AuNPs synthesized on mito-GFP-MTn that expressed in HeLa cells
C. Distributions of AuNPs synthesized on mito-acGFP-MTn that expressed in HeLa cells

**Supplementary Figure 24**

A. Distributions of AuNPs synthesized in HeLa cells expressing Mito-acGFP-MTn rapidly fixed with high pressure freezing
B. Distributions of AuNPs synthesized in HeLa cells expressing Mito-acGFP-MTn rapidly fixed with high pressure freezing

## REFERENCES

1 Shaner, N. C., Patterson, G. H. & Davidson, M. W. Advances in fluorescent protein technology. J Cell Sci 120, 4247–4260, doi:10.1242/jcs.005801 (2007).

2 Griffiths, G. & Hoppeler, H. Quantitation in immunocytochemistry: correlation of immunogold labeling to absolute number of membrane antigens. The journal of histochemistry and cytochemistry : official journal of the Histochemistry Society 34, 1389–1398, doi:10.1177/34.11.3534077 (1986).

3 Tokuyasu, K. T. A technique for ultracryotomy of cell suspensions and tissues. The Journal of cell biology 57, 551–565 (1973).

4 de Boer, P., Hoogenboom, J. P. & Giepmans, B. N. Correlated light and electron microscopy: ultrastructure lights up! Nature methods 12, 503–513, doi:10.1038/nmeth.3400 (2015).

5 Connolly, C. N., Futter, C. E., Gibson, A., Hopkins, C. R. & Cutler, D. F. Transport into and out of the Golgi complex studied by transfecting cells with cDNAs encoding horseradish peroxidase. The Journal of cell biology 127, 641–652 (1994).

6 Gaietta, G. et al. Multicolor and electron microscopic imaging of connexin trafficking. Science 296, 503–507, doi:10.1126/science.1068793 (2002).

7 Shu, X. et al. A genetically encoded tag for correlated light and electron microscopy of intact cells, tissues, and organisms. PLoS biology 9, e1001041, doi:10.1371/journal.pbio.1001041 (2011).

8 Martell, J. D. et al. Engineered ascorbate peroxidase as a genetically encoded reporter for electron microscopy. Nature biotechnology 30, 1143–1148, doi:10.1038/nbt.2375 (2012).

9 Lam, S. S. et al. Directed evolution of APEX2 for electron microscopy and proximity labeling. Nature methods 12, 51–54, doi:10.1038/nmeth.3179 (2015).

10 Mavlyutov, T. A. et al. APEX2-enhanced electron microscopy distinguishes sigma-1 receptor localization in the nucleoplasmic reticulum. Oncotarget 8, 51317–51330, doi:10.18632/oncotarget.17906 (2017).

11 Sano, T., Glazer, A. N. & Cantor, C. R. A streptavidin-metallothionein chimera that allows specific labeling of biological materials with many different heavy metal ions. Proceedings of the National Academy of Sciences of the United States of America 89, 1534–1538 (1992).

12 Mercogliano, C. P. & DeRosier, D. J. Gold nanocluster formation using metallothionein: mass spectrometry and electron microscopy. Journal of molecular biology 355, 211–223, doi:10.1016/j.jmb.2005.10.026 (2006).

13 Mercogliano, C. P. & DeRosier, D. J. Concatenated metallothionein as a clonable gold label for electron microscopy. Journal of structural biology 160, 70–82, doi:10.1016/j.jsb.2007.06.010 (2007).

14 Nishino, Y., Yasunaga, T. & Miyazawa, A. A genetically encoded metallothionein tag enabling efficient protein detection by electron microscopy. Journal of electron microscopy 56, 93–101, doi:10.1093/jmicro/dfm008 (2007).

15 Bouchet-Marquis, C., Pagratis, M., Kirmse, R. & Hoenger, A. Metallothionein as a clonable high-density marker for cryo-electron microscopy. Journal of structural biology 177, 119–127, doi:10.1016/j.jsb.2011.10.007 (2012).

16 Fukunaga, Y. et al. Electron microscopic analysis of a fusion protein of postsynaptic density-95 and metallothionein in cultured hippocampal neurons. Journal of electron microscopy 56, 119–129, doi:10.1093/jmicro/dfm027 (2007).

17 Diestra, E., Fontana, J., Guichard, P., Marco, S. & Risco, C. Visualization of proteins in intact cells with a clonable tag for electron microscopy. Journal of structural biology 165, 157–168, doi:10.1016/j.jsb.2008.11.009 (2009).

18 Risco, C. et al. Specific, sensitive, high-resolution detection of protein molecules in eukaryotic cells using metal-tagging transmission electron microscopy. Structure 20, 759–766, doi:10.1016/j.str.2012.04.001 (2012).

19 Barajas, D., Martin, I. F., Pogany, J., Risco, C. & Nagy, P. D. Noncanonical role for the host Vps4 AAA+ ATPase ESCRT protein in the formation of Tomato bushy stunt virus replicase. PLoS Pathog 10, e1004087, doi:10.1371/journal.ppat.1004087 (2014).

20 Morphew, M. K. et al. Metallothionein as a clonable tag for protein localization by electron microscopy of cells. J Microsc 260, 20–29, doi:10.1111/jmi.12262 (2015).

21 Wang, Q., Mercogliano, C. P. & Lowe, J. A ferritin-based label for cellular electron cryotomography. Structure 19, 147–154, doi:10.1016/j.str.2010.12.002 (2011).

22 Nielson, K. B., Atkin, C. L. & Winge, D. R. Distinct metal-binding configurations in metallothionein. The Journal of biological chemistry 260, 5342–5350 (1985).

23 Brust, M., Walker, M., Bethell, D., Schiffrin, D. J. & Whyman, R. Synthesis of thiol-derivatised gold nanoparticles in a two-phase Liquid-Liquid system. J. Chem. Soc., Chem. Commun., 7:801–802, doi:10.1039/C39940000801 (1994).

24 Liou, Y. C., Tocilj, A., Davies, P. L. & Jia, Z. Mimicry of ice structure by surface hydroxyls and water of a beta-helix antifreeze protein. Nature 406, 322–324, doi:10.1038/35018604 (2000).

25 Toews, J., Rogalski, J. C., Clark, T. J. & Kast, J. Mass spectrometric identification of formaldehyde-induced peptide modifications under in vivo protein cross-linking conditions. Anal Chim Acta 618, 168–183, doi:10.1016/j.aca.2008.04.049 (2008).

26 Xavier, P. L., Chaudhari, K., Verma, P. K., Pal, S. K. & Pradeep, T. Luminescent quantum clusters of gold in transferrin family protein, lactoferrin exhibiting FRET. Nanoscale 2, 2769–2776, doi:10.1039/c0nr00377h (2010).

27 Xie, J., Zheng, Y. & Ying, J. Y. Protein-directed synthesis of highly fluorescent gold nanoclusters. J Am Chem Soc 131, 888–889, doi:10.1021/ja806804u (2009).

28 Frenkel, A. I. et al. Size-controlled synthesis and characterization of thiol-stabilized gold nanoparticles. J Chem Phys 123, 184701, doi:10.1063/1.2126666 (2005).

29 Brinas, R. P., Hu, M., Qian, L., Lymar, E. S. & Hainfeld, J. F. Gold nanoparticle size controlled by polymeric Au(I) thiolate precursor size. J Am Chem Soc 130, 975–982, doi:10.1021/ja076333e (2008).

30 LeBlanc, D. J., Smith, R. W., Wang, Z., Howard-Lock, H. E. & Lock, C. J. L. Thiomalate complexes of gold(I): preparation, characterization and crystal structures of 1:2 gold to thiomalate complexes. J. Chem. Soc., Dalton Trans., 3263–3268, doi:10.1039/A700827I (1997).

31 Bau, R. Crystal structure of the antiarthritic drug gold thiomalate (myochrysine): a double-helical geometry in the solid state. Journal of the American Chemical Society 120, 9380–9381 (1998).

32 Schaeffer, N., Shaw, C. F., Thompson, H. & Satre, R. In vitro penicillamine competition for protein-bound gold (i). Arthritis & Rheumatology 23, 165–171 (1980).

33 Schmitz, G., Minkel, D. T., Gingrich, D. & Shaw, C. F., 3rd. The binding of Gold(I) to metallothionein. J Inorg Biochem 12, 293–306 (1980).

34 Laib, J. E. et al. Formation and characterization of aurothioneins: Au,Zn,Cd-thionein, Au,Cd-thionein, and (thiomalato-Au)chi-thionein. Biochemistry 24, 1977–1986 (1985).

## References

35 Forsburg, S. L. & Rhind, N. Basic methods for fission yeast. Yeast 23, 173–183, doi:10.1002/yea.1347 (2006).

36 van Donselaar, E., Posthuma, G., Zeuschner, D., Humbel, B. M. & Slot, J. W. Immunogold labeling of cryosections from high-pressure frozen cells. Traffic 8, 471–485, doi:10.1111/j.1600-0854.2007.00552.x (2007).

37 Tokuyasu, K. T. A technique for ultracryotomy of cell suspensions and tissues. The Journal of cell biology 57, 551–565 (1973).

38 Slot, J. W. G., H.J. Cryosectioning and immunolabeling. Nature Protocols 2, 2480–2491, doi:doi:10.1038/nprot.2007.365 (2007).

