## Supplementary Chart 1 for "Direct synthesis of EM-visible gold nanoparticles on genetically encoded tags for single-molecule visualization in cells"

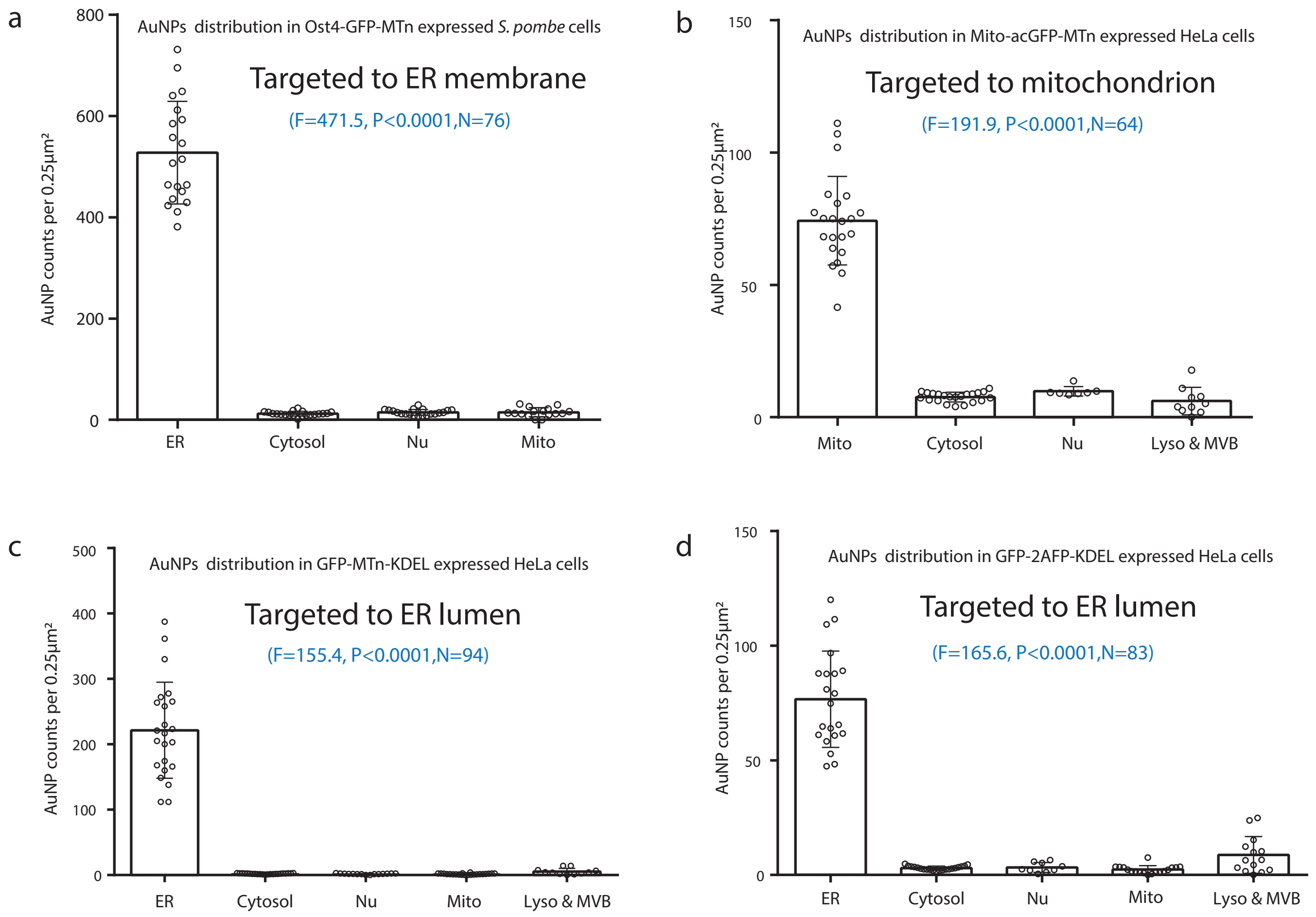


**Supplementry Chart 1. Statistical analysis (one-way ANOVA) of subcellular distribution of AuNPs synthesized in cells expressing cysteine-rich tags targeted to ER or mitochondria**

(a) Statistical chart of AuNPs distribution pattern in *S. pombe* cells expressing Ost4-GFP-MTn ($F=471.5$ ; $P<0.0001$; $N=76$), using GraphPad Prism 6 Software (GraphPad, San Diego, USA) for one-way analysis of variance (ANOVA); the chart showed the average densities of AuNPs (AuNP counts per 0.25 $\mu m^2$) and standard deviation in different organelles: endoplasmic reticulum (ER), cytosol, nuclei (Nu), and mitochondria (Mito) based on 20 EM images from 90 nm thick sections; the specimens prepared with Scheme 2b (PIPES) (See **Supplementary Table 3**). (b) Statistical chart of AuNPs distribution pattern in HeLa cells expressing Mito-acGFP-MTn ($F=191.9$; $P<0.0001$; $N=64$); the chart showed the average densities of AuNPs and standard deviation in different organelles: mitochondria (Mito), cytosol, nuclei (Nu), lysosomes and MVBs based on 22 EM images from 90 nm thick sections; the specimens prepared with Scheme 2b (DMEM). (c) Statistical chart of AuNPs distribution pattern in HeLa cells expressing GFP-MTn-KDEL ($F=155.4$ ; $P<0.0001$; $N=94$); the chart showed the average densities of AuNPs and standard deviation in ER, cytosol, nuclei (Nu),mitochondria (Mito), lysosomes and MVBs based on one-way ANOVA of 23 EM images from 90 nm thick sections; the specimens prepared with Scheme 2d. (d) Statistical chart of AuNPs distribution pattern in HeLa cells expressing GFP-2AFP-KDEL ($F=165.6$ ; $P<0.0001$; $N=83$); the chart showed the average densities of AuNPs and standard deviation in ER, cytosol, nuclei (Nu),mitochondria (Mito), lysosomes and MVBs based on one-way ANOVA of 21 EM images from 90 nm thick sections; the specimens prepared with Scheme 2d. The averaged densities of AuNPs in different organelles clearly demonstrated that the tags were localized to the targeted organelles vey specifically, while the background noises were quite low. Only in the Mito-AcGFP-MTn case (b), the background AuNP density is relative higher, it might be caused by the cytosol overexpressed tags were not efficiently delivered into mitochondria, or perhaps also cuased by the specimen preparation artifact. The higher AuNP density in lysosomes and MVBs were observed some tags dlivered to degradation pathways (d). The statistical analysis is based on the TEM images listed in the following tables, and one typical example image was selected from these 4 groups to demonstrated the processes for obtaining the average AuNPs densities in different organelles (see the following summary of the average densities of AuNPs in cells, images (a-d), and Table (A-D) ).

| Ost4-GFP-MTn (S. pombe cells) | | |
|---|---|---|
| Organelle | Mean<br>(AuNP counts per 0.25μm$^2$) | Standard deviation |
| ER | 527.7 | 101.5 |
| Cyto | 12.14 | 4.535 |
| Nu | 15.89 | 5.768 |
| Mito | 14.97 | 9.062 |
| Ordinary one-way ANOVA : P < 0.0001<br>Brown-Forsythe test: F(3,72) = 471.5; P<0.0001<br>Number of images: 20<br>Number of measurements: N=76 | | |
| Relative to ER: Cyto/ER = 2.3%; Nu/ER = 3.01%; Mito/ER = 2.83% | | |

| Mito-acGFP-MTn (HeLa cells) | | |
|---|---|---|
| Organelle | Mean<br>(AuNP counts per 0.25μm$^2$) | Standard Deviation |
| Mito | 74.26 | 16.7 |
| Cyto | 7.63 | 1.846 |
| Nu | 9.81 | 1.788 |
| Lyso & MVB | 6.14 | 5.197 |
| Ordinary one-way ANOVA : P < 0.0001<br>Brown-Forsythe test: F(3,57) = 191.9; P<0.0001<br>Number of images: 22<br>Number of measurements: N=83 | | |
| Relative to Mito: Cyto/Mito = 10.27%; Nu/Mito = 13.21%;<br>Lyso & MVB/Mito = 8.27% | | |

| GFP-MTn-KDEL (HeLa cells) | | |
|---|---|---|
| Organelle | Mean<br>(AuNP counts per 0.25μm$^2$) | Standard Deviation |
| ER | 221.48 | 73.59 |
| Cyto | 2.05 | 0.662 |
| Nu | 1.74 | 0.821 |
| Mito | 1.57 | 0.978 |
| Lyso & MVB | 5.49 | 4.838 |
| Ordinary one-way ANOVA: P < 0.0001<br>Brown-Forsythe test: F(4,89) = 155.4; P<0.0001<br>Number of images: 23<br>Number of measurements: N=94 | | |
| Relative to ER: Cyto/ER = 0.92%; Nu/ER = 0.79%; Mito/ER = 0.71%<br>Lyso & MVB/ ER = 2.48% | | |

| GFP-2AFP-KDEL (HeLa cells) | | |
|---|---|---|
| Organelle | Mean<br>(AuNP counts per 0.25μm$^2$) | Standard Deviation |
| ER | 76.66 | 21.00 |
| Cyto | 2.99 | 0.885 |
| Nu | 3.29 | 2.136 |
| Mito | 2.35 | 1.732 |
| Lyso & MVB | 8.71 | 8.021 |
| Ordinary one-way ANOVA: P < 0.0001<br>Brown-Forsythe test: F(4,78) = 165.6; P<0.0001<br>Number of images: 21<br>Number of measurements: N=83 | | |
| Relative to ER: Cyto/ER = 3.9%; Nu/ER = 4.3%; Mito/ER = 3.1%<br>Lyso & MVB/ ER = 11.4% | | |

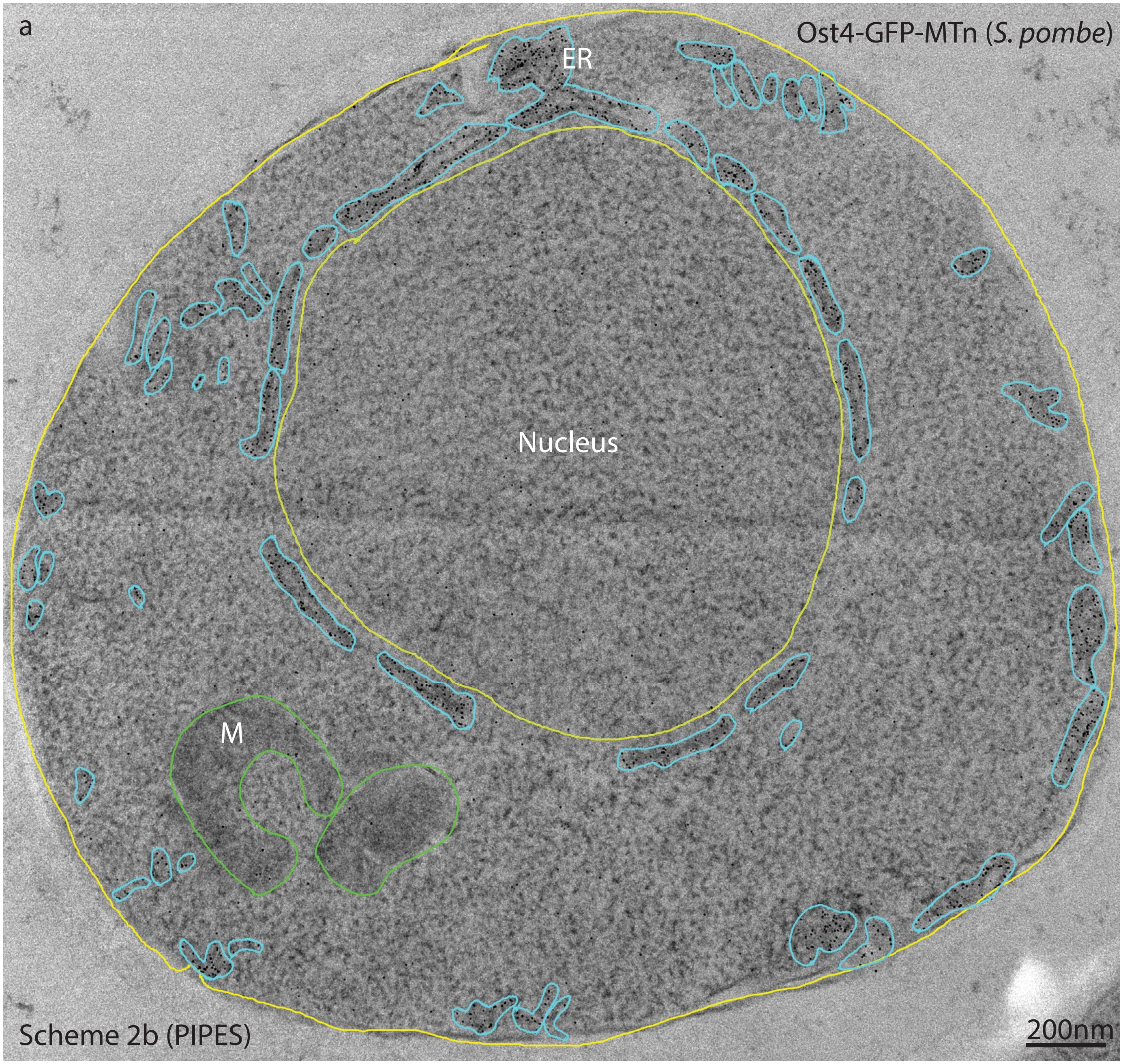


**a. Statistical analysis of the distribution of AuNPs in S. pombe cell expressing Ost4-GFP-MTn**
*S. pombe* cell expressed Ost4-GFP-MTn was processed with Scheme 2b (PIPES), embedded in SPI-Pon 812 resin, an 90 nm thin section was used for TEM imaging. The ER including nuclear envelope (NE) regions were selected (marked with cyan color) for counting AuNPs and caculating the corresponding area, then obtained the the averaged density of AuNPs in ER: 546.41 counts per 0.25μm² ( total 953 counts in 436484.6 nm²). The averaged density of AuNPs in cytosol, mitochondrion (M) and nucleus were also obtained as 9.51, 17.53, 13.84 counts per 0.25μm² ,respectively. The AuNPs counts and areas of interest were obtianed either mannualy or semiautomatically by using a free software called ImageJ (http://imagej.nih.gov/ij/). The image ID is 202-0008. 20 images of *S. pombe* cell expressed Ost4-GFP-MTn were used for statistical analysis (see following Table A, and also the above statistical chart (a)).

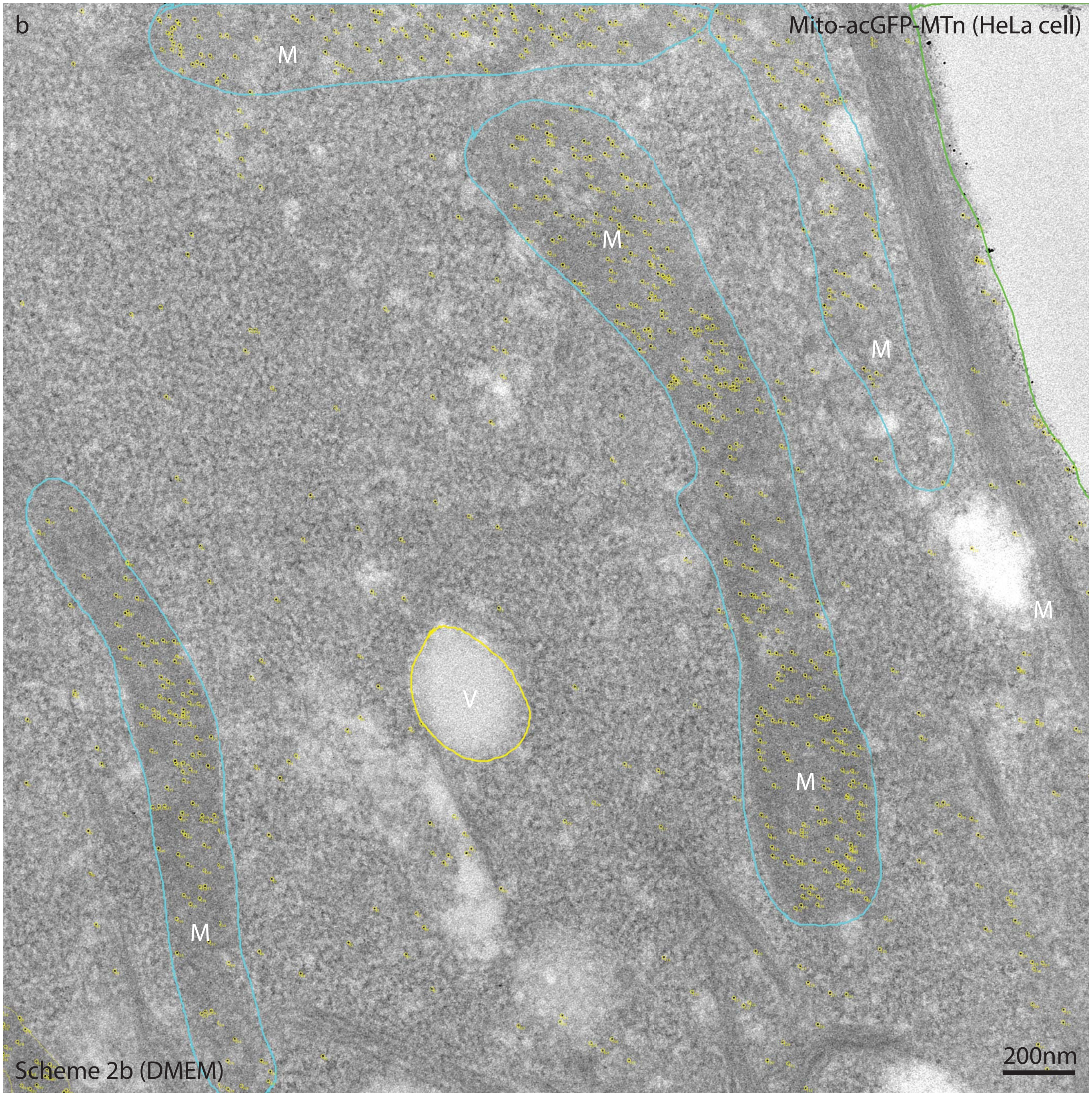


**b. Statistical analysis of the distribution of AuNPs in HeLa cell expressing Mito-acGFP-MTn**
*HeLa cell* expressed Mito-acGFP-MTn was processed with Scheme 2b (DMEM (see Supplementary Table 3, adapted from Scheme 2b (PBS)), embedded in SPI-Pon 812 resin, an 90 nm thin section was used for TEM imaging. The mitochondria were selected (marked with cyan) for counting AuNPs and caculating the corresponding area, then obtained the the averaged density of AuNPs in mitochondria: 84.22 counts per 0.25 $\mu m^2$. The averaged density of AuNPs in cytosol was also obtained as 7.63 counts per 0.25$\mu m^2$ . The AuNPs counts and areas of interest were obtianed esemiautomatically by using a free software called ImageJ (http://imagej.nih.gov/ij/).The image ID is Mito-MTn-stained-0056. 22 images were used for statistical analysis, the statistical data and chart(see following Table B, and also the above statistical chart (b)) .

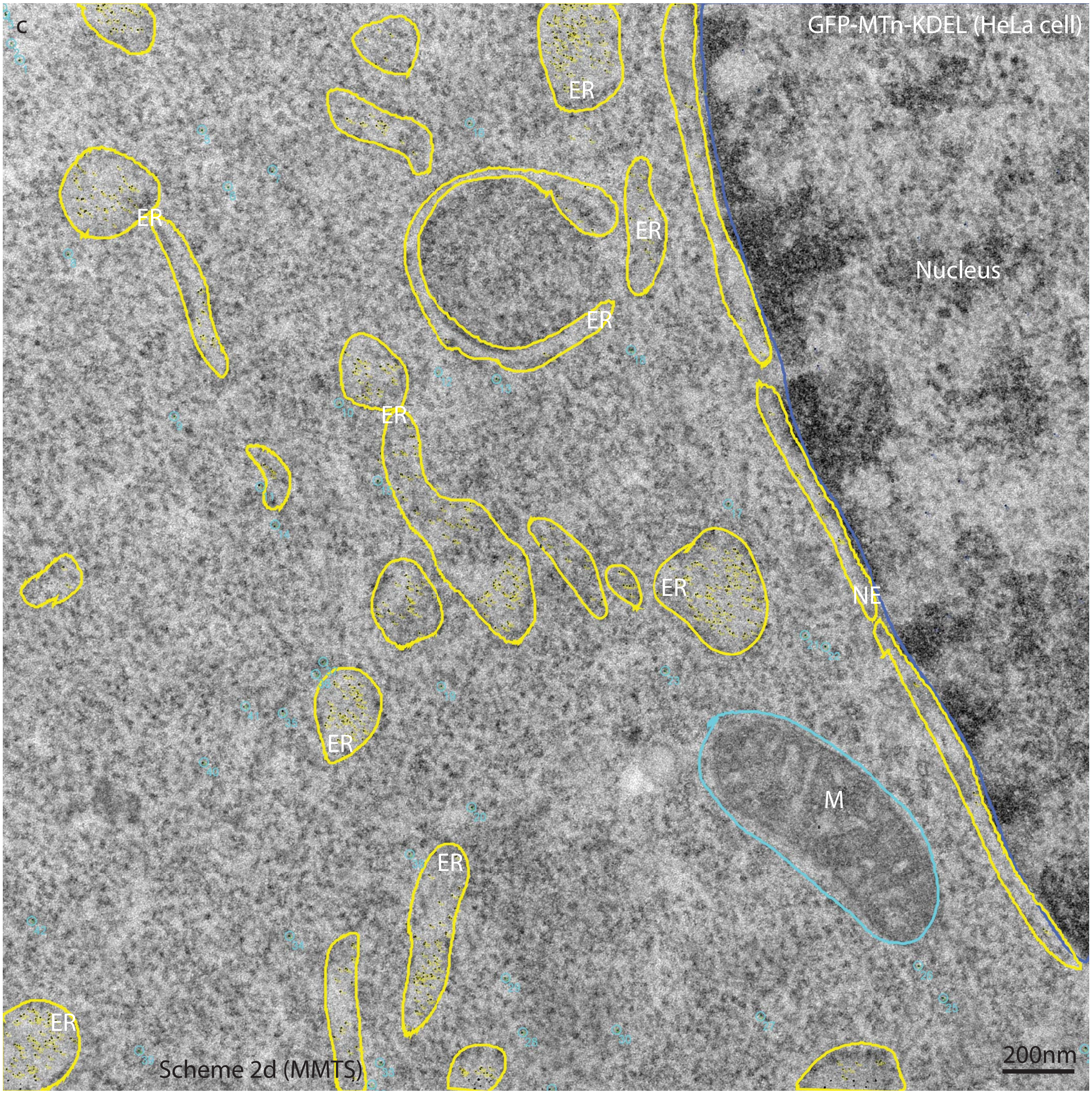


**c. Statistical analysis of the distribution of AuNPs in HeLa cell expressing GFP-MTn-KDEL**

*HeLa cell* expressed GFP-MT-KDEL was processed with Scheme 2d (MMTS) (see Supplementary Table 3), embedded in SPI-Pon 812 resin, an 90 nm thin section was used for TEM imaging. The ER lumens were selected (marked with yellow) for counting AuNPs and caculating the corresponding area, then obtained the the averaged density of AuNPs in ER: 174.34 counts per 0.25μm². The averaged density of AuNPs in cytosol, mitochondrion (M) and nucleus were also obtained as 1.71, 0.00, 2.47 counts per 0.25μm² ,respectively. The AuNPs counts and areas of interest were obtianed either mannualy or semiautomatically by using a free software called ImageJ (http://imagej.nih.gov/ij/).The image ID is MMTS-MTn-KDEL-2-stain UA6min-0003. 23 images were used for statistical analysis, the statistical data and chart (see following Table C, and also the above statistical chart (c)) .

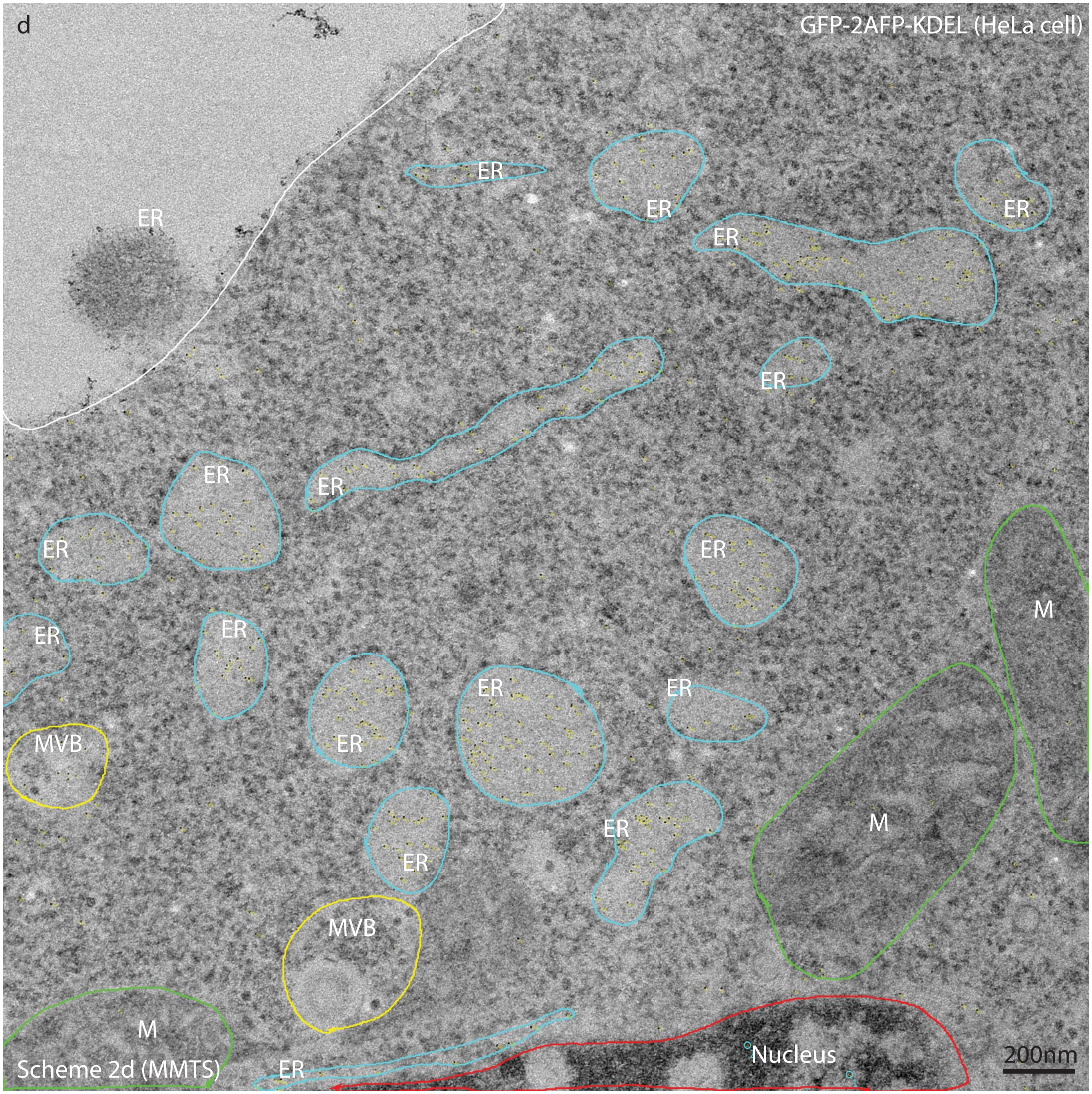


**d. Statistical analysis of the distribution of AuNPs in HeLa cell expressing GFP-2AFP-KDEL**

*HeLa cell* expressed GFP-2AFP-KDEL was processed with Scheme 2d (MMTS) (see Supplementary Table 3), embedded in SPI-Pon 812 resin, an 90 nm thin section was used for TEM imaging. The ER lumens were selected (marked with cyan) for counting AuNPs and caculating the corresponding area, then obtained the the averaged density of AuNPs in ER: 96.77 counts per 0.25μm². The averaged density of AuNPs in cytosol, mitochondrion (M) ,nucleus and MVB were also obtained as 4.54, 3.15, 2.03,15.29 counts per 0.25μm² ,respectively. The AuNPs counts and areas of interest were obtianed either mannualy or semiautomatically by using a free software called ImageJ (http://imagej.nih.gov/ij/).The image ID is MMTS-AFP-KDEL-2-0006. 21 images were used for statistical analysis, the statistical data and chart (see following Table D, and also the above statistical chart (d)) .

### A. AuNPs in S.pombe cells expressing Ost4-GFP-MTn

ER

| Image ID | counts (AuNPs in ROI) | ROI Area ($nm^2$) | particles in $0.04\mu m^2$ | particles in $0.25\mu m^2$ |
|---|---|---|---|---|
| 202-0002 | 1038 | 594608.858 | 69.83 | 436.42 |
| 202-0003 | 1700 | 655264.519 | 103.77 | 648.59 |
| 202-0004 | 1562 | 700035.313 | 89.25 | 557.83 |
| 202-0005 | 1235 | 809409.333 | 61.03 | 381.45 |
| 202-0006 | 1381 | 749417.008 | 73.71 | 460.69 |
| 202-0007 | 1211 | 596989.388 | 81.14 | 507.13 |
| 202-0008 | 1019 | 564315.954 | 72.23 | 451.43 |
| 202-0009 | 1462 | 787712.299 | 74.24 | 464.00 |
| 202-0010 | 1176 | 715160.038 | 65.78 | 411.10 |
| 202-0011 | 648 | 382515.08 | 67.76 | 423.51 |
| 202-0012 | 1238 | 601265.321 | 82.36 | 514.75 |
| 202-0013 | 992 | 387169.434 | 102.49 | 640.55 |
| 202-0014 | 1405 | 480249.915 | 117.02 | 731.39 |
| 202-0015 | 1024 | 596359.053 | 68.68 | 429.27 |
| 202-0016 | 1244 | 531659.572 | 93.59 | 584.96 |
| 202-0017 | 1325 | 476402.393 | 111.25 | 695.32 |
| 202-0018 | 878 | 370149.488 | 94.88 | 593.00 |
| 202-0019 | 1793 | 732321.525 | 97.94 | 612.09 |
| 202-0020 | 1346 | 725224.488 | 74.24 | 463.99 |
| 202-0021 | 954 | 436484.576 | 87.43 | 546.41 |

Cyto

| Image ID | counts (AuNPs in ROI) | ROI Area ($nm^2$) | particles in $0.04\mu m^2$ | particles in $0.25\mu m^2$ |
|---|---|---|---|---|
| 202-0002 | 145 | 3481317.134 | 1.67 | 10.41 |
| 202-0004 | 220 | 2371227.288 | 3.71 | 23.19 |
| 202-0006 | 131 | 2821397.161 | 1.86 | 11.61 |
| 202-0007 | 63 | 1719064.389 | 1.47 | 9.16 |
| 202-0009 | 221 | 4526484.188 | 1.95 | 12.21 |
| 202-0010 | 177 | 3674032.805 | 1.93 | 12.04 |
| 202-0011 | 162 | 3450202.487 | 1.88 | 11.74 |
| 202-0012 | 158 | 3588230.32 | 1.76 | 11.01 |
| 202-0013 | 217 | 4287386.136 | 2.02 | 12.65 |
| 202-0014 | 238 | 49000425.95 | 0.19 | 1.21 |
| 202-0015 | 233 | 3157084.886 | 2.95 | 18.45 |
| 202-0017 | 83 | 2197316.046 | 1.51 | 9.44 |
| 202-0019 | 144 | 2419374.167 | 2.38 | 14.88 |
| 202-0020 | 153 | 5030118.628 | 1.22 | 7.60 |
| 1-15K | 278 | 6965647.883 | 1.60 | 9.98 |
| 3-21K | 147 | 2440787.733 | 2.41 | 15.06 |
| 4-21k | 169 | 2520223.303 | 2.68 | 16.76 |
| 8-15K | 278 | 4366870.424 | 2.55 | 15.92 |
| 9-15K | 169 | 4199164.069 | 1.61 | 10.06 |
| 202-0008 | 126 | 3311996.443 | 1.52 | 9.51 |

Mito

| Image ID | counts (AuNPs in ROI) | ROI Area ($nm^2$) | particles in $0.04\mu m^2$ | particles in $0.25\mu m^2$ |
|---|---|---|---|---|
| 202-0002 | 2 | 29421.106 | 2.72 | 16.99 |
| 202-0006 | 15 | 142802.645 | 4.20 | 26.26 |
| 202-0007 | 12 | 138280.848 | 3.47 | 21.69 |
| 202-0009 | 11 | 214882.98 | 2.05 | 12.80 |
| 202-0010 | 14 | 259637.823 | 2.16 | 13.48 |

| | | | | |
|---|---|---|---|---|
| 202-0011 | 0 | 19262.051 | 0.00 | 0.00 |
| 202-0012 | 12 | 262911.602 | 1.83 | 11.41 |
| 202-0013 | 9 | 72003.33 | 5.00 | 31.25 |
| 202-0014 | 36 | 721131.714 | 2.00 | 12.48 |
| 202-0019 | 0 | 6389.149 | 0.00 | 0.00 |
| 202-0020 | 9 | 303526.369 | 1.19 | 7.41 |
| 1-15K | 5 | 100786.402 | 1.98 | 12.40 |
| 4-21k | 5 | 41974.593 | 4.76 | 29.78 |
| 8-15K | 5 | 77781.396 | 2.57 | 16.07 |
| 9-15K | 3 | 75268.309 | 1.59 | 9.96 |
| 202-0008 | 13 | 185407.971 | 2.80 | 17.53 |

| Nu | | | | |
|---|---|---|---|---|
| Image ID | counts (AuNPs in ROI) | ROI Area ($nm^2$) | particles in 0.04μ$m^2$ | particles in 0.25μ$m^2$ |
| 202-0002 | 39 | 1465762.912 | 1.06 | 6.65 |
| 202-0004 | 72 | 922952.031 | 3.12 | 19.50 |
| 202-0006 | 112 | 1934630.506 | 2.32 | 14.47 |
| 202-0007 | 83 | 1850115.642 | 1.79 | 11.22 |
| 202-0009 | 48 | 807959.454 | 2.38 | 14.85 |
| 202-0010 | 69 | 1400924.822 | 1.97 | 12.31 |
| 202-0011 | 100 | 2144837.13 | 1.86 | 11.66 |
| 202-0012 | 83 | 1804420.248 | 1.84 | 11.50 |
| 202-0013 | 47 | 1448944.644 | 1.30 | 8.11 |
| 202-0015 | 72 | 963319.285 | 2.99 | 18.69 |
| 202-0016 | 219 | 1880033.425 | 4.66 | 29.12 |
| 202-0017 | 97 | 2119647.955 | 1.83 | 11.44 |
| 202-0019 | 155 | 1886511.129 | 3.29 | 20.54 |
| 202-0020 | 68 | 2272330.24 | 1.20 | 7.48 |
| 1-15K | 119 | 1629878.67 | 2.92 | 18.25 |
| 3-21K | 109 | 1307699.077 | 3.33 | 20.84 |
| 4-21k | 114 | 1649110.895 | 2.77 | 17.28 |
| 8-15K | 107 | 1240818.233 | 3.45 | 21.56 |
| 9-15K | 53 | 1563651.205 | 1.36 | 8.47 |
| 202-0008 | 91 | 1644261.405 | 2.21 | 13.84 |

**B. AuNPs in HeLa cells expressing Mito-GFP-MTn**

| Mito | | | | |
|---|---|---|---|---|
| Image ID | Counts (AuNPs in ROI) | ROI Area ($nm^2$) | Particles in $0.04\mu m^2$ | Particles in $0.25\mu m^2$ |
| mito-MTn-0.1GA 20min-0001 | 216 | 779498.442 | 11.08405038 | 69.28 |
| mito-MTn-0.1GA 30min-0005 | 280 | 1026733.882 | 10.90837674 | 68.18 |
| mito-MTn-0.1GA 30min-0010 | 485 | 2228261.155 | 8.706340348 | 54.41 |
| mito-MTn-0.1GA 30min-0012 | 609 | 2383189.76 | 10.22159478 | 63.88 |
| mito-MTn-0.1GA 30min-0028 | 456 | 1829639.675 | 9.969176035 | 62.31 |
| Mito-MTn-stained-0031 | 326 | 1961542.925 | 6.64782801 | 41.55 |
| Mito-MTn-stained-0034 | 341 | 1103321.644 | 12.36266874 | 77.27 |
| Mito-MTn-stained-0035 | 347 | 1156413.299 | 12.00262917 | 75.02 |
| Mito-MTn-stained-0040 | 556 | 1797356.762 | 12.37372595 | 77.34 |
| Mito-MTn-stained-0054 | 353 | 865479.09 | 16.31466336 | 101.97 |
| Mito-MTn-stained-0055 | 472 | 1102158.414 | 17.13002392 | 107.06 |
| Mito-MTn-stained-0056 | 592 | 1757316.777 | 13.47508902 | 84.22 |
| Mito-MTn-stained-0057 | 368 | 828028.214 | 17.77717202 | 111.11 |
| Mito-MTn-stained-0058 | 320 | 956786.611 | 13.37811363 | 83.61 |
| Mito-MTn-stained-0059 | 419 | 1415220.139 | 11.84268054 | 74.02 |
| Mito-MTn-stained-0061 | 650 | 2162604.2 | 12.02254208 | 75.14 |
| Mito-MTn-stained-0062 | 716 | 3069862.225 | 9.329408912 | 58.31 |
| Mito-MTn-stained-0064 | 659 | 2038563.629 | 12.93067316 | 80.82 |
| Mito-MTn-stained-0075 | 253 | 930694.699 | 10.87359798 | 67.96 |
| Mito-MTn-stained-0076 | 525 | 1927128.646 | 10.89704107 | 68.11 |
| Mito-MTn-stained-0077 | 586 | 1954230.826 | 11.99448893 | 74.97 |
| Mito-MTn-stained-0080 | 519 | 2271320.385 | 9.140057976 | 57.13 |

| Cyto | | | | |
|---|---|---|---|---|
| Image ID | Counts (AuNPs in ROI) | ROI Area ($nm^2$) | Particles in $0.04\mu m^2$ | Particles in $0.25\mu m^2$ |
| mito-MTn-0.1GA 20min-0001 | 92 | 3694339.995 | 0.996118388 | 6.23 |
| mito-MTn-0.1GA 30min-0005 | 64 | 3592316.709 | 0.712632044 | 4.45 |
| mito-MTn-0.1GA 30min-0010 | 62 | 2479888.699 | 1.000044881 | 6.25 |
| mito-MTn-0.1GA 30min-0012 | 72 | 2324960.094 | 1.238730939 | 7.74 |
| mito-MTn-0.1GA 30min-0028 | 168 | 4839854.992 | 1.388471351 | 8.68 |
| Mito-MTn-stained-0031 | 154 | 4154625.388 | 1.482684821 | 9.27 |
| Mito-MTn-stained-0034 | 91 | 2332175.591 | 1.560774418 | 9.75 |
| Mito-MTn-stained-0035 | 65 | 2209055.003 | 1.176973863 | 7.36 |
| Mito-MTn-stained-0040 | 134 | 3478761.695 | 1.540778147 | 9.63 |
| Mito-MTn-stained-0054 | 118 | 3743686.955 | 1.260789178 | 7.88 |
| Mito-MTn-stained-0055 | 118 | 7368013.546 | 0.640606857 | 4.00 |
| Mito-MTn-stained-0056 | 213 | 6983066.234 | 1.220094399 | 7.63 |
| Mito-MTn-stained-0057 | 136 | 3880121.64 | 1.402017902 | 8.76 |
| Mito-MTn-stained-0058 | 85 | 3245513.473 | 1.047600026 | 6.55 |
| Mito-MTn-stained-0059 | 83 | 2847687.369 | 1.165858316 | 7.29 |
| Mito-MTn-stained-0061 | 232 | 7065328.367 | 1.313456292 | 8.21 |
| Mito-MTn-stained-0062 | 101 | 5340357.013 | 0.75650373 | 4.73 |
| Mito-MTn-stained-0064 | 160 | 7189408.898 | 0.890198359 | 5.56 |
| Mito-MTn-stained-0075 | 84 | 2339313.635 | 1.436318735 | 8.98 |
| Mito-MTn-stained-0076 | 217 | 5870446.25 | 1.478592875 | 9.24 |
| Mito-MTn-stained-0077 | 214 | 6152479.645 | 1.391308951 | 8.70 |
| Mito-MTn-stained-0080 | 289 | 6613420.206 | 1.74796091 | 10.92 |

| Nu | | | | |
|---|---|---|---|---|
| Image ID | Counts (AuNPs in ROI) | ROI Area ($nm^2$) | Particles in $0.04\mu m^2$ | Particles in $0.25\mu m^2$ |
| mito-MTn-0.1GA 30min-0005 | 44 | 1115833.145 | 1.577296756 | 9.86 |
| mito-MTn-0.1GA 30min-0028 | 70 | 1935898.875 | 1.446356541 | 9.04 |
| Mito-MTn-stained-0031 | 97 | 2707172.976 | 1.433229437 | 8.96 |
| Mito-MTn-stained-0034 | 47 | 1272652.619 | 1.477229506 | 9.23 |
| Mito-MTn-stained-0035 | 60 | 1090944.283 | 2.199929032 | 13.75 |
| Mito-MTn-stained-0040 | 133 | 3544875.304 | 1.500758008 | 9.38 |
| Mito-MTn-stained-0075 | 40 | 1182345.815 | 1.353241987 | 8.46 |

| Lyso & MVB | | | | |
|---|---|---|---|---|
| Image ID | Counts (AuNPs in ROI) | ROI Area （$nm^2$） | Particles in $0.04\mu m^2$ | Particles in $0.25\mu m^2$ |
| mito-MTn-0.1GA 30min-0028 | 13 | 622578.985 | 0.835235388 | 5.22 |
| Mito-MTn-stained-0054 | 1 | 98983.809 | 0.404106494 | 2.53 |
| Mito-MTn-stained-0055 | 6 | 390395.361 | 0.614761403 | 3.84 |
| Mito-MTn-stained-0056 | 0 | 89034.46 | 0 | 0.00 |
| Mito-MTn-stained-0031 | 3 | 404631.238 | 0.296566327 | 1.85 |
| Mito-MTn-stained-0035 | 4 | 251737.269 | 0.635583283 | 3.97 |
| Mito-MTn-stained-0058 | 36 | 505849.77 | 2.846694978 | 17.79 |
| Mito-MTn-stained-0076 | 34 | 1140362.949 | 1.19260276 | 7.45 |
| Mito-MTn-stained-0077 | 35 | 1121262.056 | 1.248593041 | 7.80 |
| Mito-MTn-stained-0080 | 15 | 343231.936 | 1.748089082 | 10.93 |

### C. AuNPs in Hela cells expressing GFP-MTn-KDEL

| ER | | | | |
|---|---|---|---|---|
| Image ID | counts (AuNPs in ROI) | ROI Area ($nm^2$) | Particles in $0.04\mu m^2$ | Particles in $0.25\mu m^2$ |
| MMTS-MTn-kdel-2-0004 | 509 | 766770.456 | 26.55 | 165.96 |
| MMTS-MTn-kdel-2-0008 | 843 | 1315955.084 | 25.62 | 160.15 |
| MMTS-MTn-kdel-2-0010 | 419 | 935629.673 | 17.91 | 111.96 |
| MMTS-MTn-kdel-2-0015 | 458 | 682349.647 | 26.85 | 167.80 |
| MMTS-MTn-kdel-2-0018 | 828 | 1500165.64 | 22.08 | 137.98 |
| MMTS-MTn-kdel-2-0020 | 768 | 1715726.806 | 17.90 | 111.91 |
| MMTS-MTn-kdel-2-stain UA6min-0002 | 5051 | 3825853.192 | 52.81 | 330.06 |
| MMTS-MTn-kdel-2-stain UA6min-0003 | 705 | 1010979.936 | 27.89 | 174.34 |
| MMTS-MTn-kdel-2-stain UA6min-0004 | 827 | 900207.19 | 36.75 | 229.67 |
| MMTS-MTn-kdel-2-stain UA6min-0005 | 642 | 782979.841 | 32.80 | 204.99 |
| MMTS-MTn-kdel-2-stain UA6min-0006 | 1154 | 1059683.445 | 43.56 | 272.25 |
| MMTS-MTn-kdel-2-stain UA6min-0007 | 1211 | 1512552.316 | 32.03 | 200.16 |
| MMTS-MTn-kdel-2-stain UA6min-0009 | 870 | 973631.798 | 35.74 | 223.39 |
| MMTS-MTn-kdel-2-stain UA6min-0010 | 1207 | 1169117.419 | 41.30 | 258.10 |
| MMTS-MTn-kdel-2-stain UA6min-0011 | 1224 | 1507619.096 | 32.48 | 202.97 |
| MMTS-MTn-kdel-2-stain UA6min-0012 | 1128 | 1275316.116 | 35.38 | 221.12 |
| MMTS-MTn-kdel-2-stain UA6min-0013 | 756 | 1270605.109 | 23.80 | 148.75 |
| MMTS-MTn-kdel-2-stain UA6min-0014 | 1572 | 1415551.221 | 44.42 | 277.63 |
| MMTS-MTn-kdel-2-stain UA6min-0015 | 836 | 787368.53 | 42.47 | 265.44 |
| MMTS-MTn-kdel-2-stain UA6min-0016 | 738 | 699749.297 | 42.19 | 263.67 |
| MMTS-MTn-kdel-2-stain UA6min-0017 | 1263 | 1456828.774 | 34.68 | 216.74 |
| MMTS-MTn-kdel-2-stain UA6min-0019 | 6820 | 4399488.855 | 62.01 | 387.55 |
| MMTS-MTn-kdel-2-stain UA6min-0020 | 1927 | 1332412.532 | 57.85 | 361.56 |

| Mito | | | | |
|---|---|---|---|---|
| Image ID | counts (AuNPs in ROI) | ROI Area ($nm^2$) | Particles in $0.04\mu m^2$ | Particles in $0.25\mu m^2$ |
| MMTS-MTn-kdel-2-0004 | 5 | 643840.945 | 0.31 | 1.94 |
| MMTS-MTn-kdel-2-0008 | 12 | 1427562.776 | 0.34 | 2.10 |
| MMTS-MTn-kdel-2-0010 | 2 | 341177.024 | 0.23 | 1.47 |
| MMTS-MTn-kdel-2-0015 | 2 | 962415.585 | 0.08 | 0.52 |
| MMTS-MTn-kdel-2-0018 | 3 | 556678.237 | 0.22 | 1.35 |
| MMTS-MTn-kdel-2-0020 | 6 | 608386.01 | 0.39 | 2.47 |
| MMTS-MTn-kdel-2-stain UA6min-0002 | 2 | 124199.53 | 0.64 | 4.03 |
| MMTS-MTn-kdel-2-stain UA6min-0003 | 0 | 236595.415 | 0.00 | 0.00 |
| MMTS-MTn-kdel-2-stain UA6min-0004 | 2 | 208789.747 | 0.38 | 2.39 |
| MMTS-MTn-kdel-2-stain UA6min-0005 | 1 | 108150.204 | 0.37 | 2.31 |
| MMTS-MTn-kdel-2-stain UA6min-0006 | 9 | 822163.429 | 0.44 | 2.74 |
| MMTS-MTn-kdel-2-stain UA6min-0007 | 6 | 589167.411 | 0.41 | 2.55 |
| MMTS-MTn-kdel-2-stain UA6min-0009 | 7 | 1266333.575 | 0.22 | 1.38 |
| MMTS-MTn-kdel-2-stain UA6min-0010 | 9 | 2097343.138 | 0.17 | 1.07 |
| MMTS-MTn-kdel-2-stain UA6min-0011 | 4 | 1109377.558 | 0.14 | 0.90 |
| MMTS-MTn-kdel-2-stain UA6min-0012 | 8 | 1285340.963 | 0.25 | 1.56 |
| MMTS-MTn-kdel-2-stain UA6min-0013 | 2 | 1152981.738 | 0.07 | 0.43 |
| MMTS-MTn-kdel-2-stain UA6min-0015 | 4 | 845609.009 | 0.19 | 1.18 |
| MMTS-MTn-kdel-2-stain UA6min-0016 | 0 | 308413.935 | 0.00 | 0.00 |
| MMTS-MTn-kdel-2-stain UA6min-0017 | 7 | 1404680.977 | 0.20 | 1.25 |
| MMTS-MTn-kdel-2-stain UA6min-0019 | 4 | 1269630.455 | 0.13 | 0.79 |
| MMTS-MTn-kdel-2-stain UA6min-0020 | 12 | 1433695.611 | 0.33 | 2.09 |

| Cyto | | | | |
|---|---|---|---|---|
| Image ID | counts (AuNPs in ROI) | ROI Area ($nm^2$) | Particles in $0.04\mu m^2$ | Particles in $0.25\mu m^2$ |
| MMTS-MTn-kdel-2-0004 | 46 | 7817374.417 | 0.24 | 1.47 |
| MMTS-MTn-kdel-2-0008 | 36 | 5159337.823 | 0.28 | 1.74 |

| | | | | |
|---|---|---|---|---|
| MMTS-MTn-kdel-2-0010 | 22 | 4784137.041 | 0.18 | 1.15 |
| MMTS-MTn-kdel-2-0015 | 40 | 5449571.616 | 0.29 | 1.84 |
| MMTS-MTn-kdel-2-0018 | 45 | 7171141.941 | 0.25 | 1.57 |
| MMTS-MTn-kdel-2-0020 | 18 | 6535749.457 | 0.11 | 0.69 |
| MMTS-MTn-kdel-2-stain UA6min-0002 | 28 | 4009929.079 | 0.28 | 1.75 |
| MMTS-MTn-kdel-2-stain UA6min-0003 | 42 | 6156848.624 | 0.27 | 1.71 |
| MMTS-MTn-kdel-2-stain UA6min-0004 | 47 | 4653479.609 | 0.40 | 2.52 |
| MMTS-MTn-kdel-2-stain UA6min-0005 | 42 | 3880235.85 | 0.43 | 2.71 |
| MMTS-MTn-kdel-2-stain UA6min-0006 | 32 | 5780271.668 | 0.22 | 1.38 |
| MMTS-MTn-kdel-2-stain UA6min-0007 | 62 | 6673399.42 | 0.37 | 2.32 |
| MMTS-MTn-kdel-2-stain UA6min-0009 | 56 | 6278159.396 | 0.36 | 2.23 |
| MMTS-MTn-kdel-2-stain UA6min-0010 | 64 | 5961525.261 | 0.43 | 2.68 |
| MMTS-MTn-kdel-2-stain UA6min-0011 | 44 | 4711779.492 | 0.37 | 2.33 |
| MMTS-MTn-kdel-2-stain UA6min-0012 | 32 | 6479331.777 | 0.20 | 1.23 |
| MMTS-MTn-kdel-2-stain UA6min-0013 | 59 | 4569134.155 | 0.52 | 3.23 |
| MMTS-MTn-kdel-2-stain UA6min-0014 | 68 | 6105491.22 | 0.45 | 2.78 |
| MMTS-MTn-kdel-2-stain UA6min-0015 | 37 | 4171456.398 | 0.35 | 2.22 |
| MMTS-MTn-kdel-2-stain UA6min-0016 | 44 | 3919701.055 | 0.45 | 2.81 |
| MMTS-MTn-kdel-2-stain UA6min-0017 | 49 | 5212740.238 | 0.38 | 2.35 |
| MMTS-MTn-kdel-2-stain UA6min-0019 | 19 | 3144152.67 | 0.24 | 1.51 |
| MMTS-MTn-kdel-2-stain UA6min-0020 | 76 | 6461877.675 | 0.47 | 2.94 |

Nu

| Image ID | counts (AuNPs in ROI) | ROI Area ($nm^2$) | Particles in 0.04μm$^2$ | Particles in 0.25μm$^2$ |
|---|---|---|---|---|
| MMTS-MTn-kdel-2-0008 | 0 | 47813.56 | 0.00 | 0.00 |
| MMTS-MTn-kdel-2-0010 | 6 | 963706.506 | 0.25 | 1.56 |
| MMTS-MTn-kdel-2-0020 | 0 | 70064.475 | 0.00 | 0.00 |
| MMTS-MTn-kdel-2-stain UA6min-0003 | 18 | 1823561.843 | 0.39 | 2.47 |
| MMTS-MTn-kdel-2-stain UA6min-0004 | 22 | 2335952.213 | 0.38 | 2.35 |
| MMTS-MTn-kdel-2-stain UA6min-0005 | 28 | 3002440.644 | 0.37 | 2.33 |
| MMTS-MTn-kdel-2-stain UA6min-0006 | 12 | 1565867.276 | 0.31 | 1.92 |
| MMTS-MTn-kdel-2-stain UA6min-0007 | 3 | 452866.671 | 0.26 | 1.66 |
| MMTS-MTn-kdel-2-stain UA6min-0011 | 13 | 1748010.819 | 0.30 | 1.86 |
| MMTS-MTn-kdel-2-stain UA6min-0013 | 11 | 1816869.627 | 0.24 | 1.51 |
| MMTS-MTn-kdel-2-stain UA6min-0014 | 11 | 1706943.377 | 0.26 | 1.61 |
| MMTS-MTn-kdel-2-stain UA6min-0015 | 21 | 2306298.994 | 0.36 | 2.28 |
| MMTS-MTn-kdel-2-stain UA6min-0016 | 17 | 2145125.346 | 0.32 | 1.98 |
| MMTS-MTn-kdel-2-stain UA6min-0017 | 7 | 1087268.551 | 0.26 | 1.61 |
| MMTS-MTn-kdel-2-stain UA6min-0019 | 5 | 414713.838 | 0.48 | 3.01 |
| MMTS-MTn-kdel-2-stain UA6min-0020 | | 0 | | |

lyso & MVB

| Image ID | counts (AuNPs in ROI) | ROI Area ($nm^2$) | Particles in 0.04μm$^2$ | Particles in 0.25μm$^2$ |
|---|---|---|---|---|
| MMTS-MTn-kdel-2-0004 | | 0 | | |
| MMTS-MTn-kdel-2-0008 | 59 | 1041699.113 | 2.27 | 14.16 |
| MMTS-MTn-kdel-2-0010 | 26 | 1457261.648 | 0.71 | 4.46 |
| MMTS-MTn-kdel-2-0015 | 2 | 2133648.97 | 0.04 | 0.23 |
| MMTS-MTn-kdel-2-0020 | 3 | 298059.07 | 0.40 | 2.52 |
| MMTS-MTn-kdel-2-stain UA6min-0002 | 5 | 1268004.017 | 0.16 | 0.99 |
| MMTS-MTn-kdel-2-stain UA6min-0009 | 19 | 709861.049 | 1.07 | 6.69 |
| MMTS-MTn-kdel-2-stain UA6min-0011 | 1 | 151198.853 | 0.26 | 1.65 |
| MMTS-MTn-kdel-2-stain UA6min-0012 | 3 | 187996.962 | 0.64 | 3.99 |
| MMTS-MTn-kdel-2-stain UA6min-0013 | 13 | 418395.189 | 1.24 | 7.77 |
| MMTS-MTn-kdel-2-stain UA6min-0016 | 7 | 123972.367 | 2.26 | 14.12 |
| MMTS-MTn-kdel-2-stain UA6min-0017 | 1 | 66467.278 | 0.60 | 3.76 |

### D. AuNPs in Hela cells expressing GFP-2AFP-KDEL

**ER**

| Image ID | Counts (AuNPs in ROI) | ROI Area (nm$^2$) | Particles in 0.04μm$^2$ | Particles in 0.25μm$^2$ |
|---|---|---|---|---|
| MMTS-AFP-KDEL-1-0001 | 265.00 | 551899.58 | 19.21 | 120.04 |
| MMTS-AFP-KDEL-1-0002 | 244.00 | 558009.31 | 17.49 | 109.32 |
| MMTS-AFP-KDEL-2-0001 | 427.00 | 957364.11 | 17.84 | 111.50 |
| MMTS-AFP-KDEL-2-0002 | 394.00 | 1122485.87 | 14.04 | 87.75 |
| MMTS-AFP-KDEL-2-0003 | 250.00 | 701869.66 | 14.25 | 89.05 |
| MMTS-AFP-KDEL-2-0004 | 414.00 | 1382909.14 | 11.97 | 74.84 |
| MMTS-AFP-KDEL-2-0005 | 303.00 | 956184.85 | 12.68 | 79.22 |
| MMTS-AFP-KDEL-2-0006 | 412.00 | 1064359.25 | 15.48 | 96.77 |
| MMTS-AFP-KDEL-2-0007 | 235.00 | 668695.15 | 14.06 | 87.86 |
| MMTS-AFP-KDEL-2-0008 | 273.00 | 843428.14 | 12.95 | 80.92 |
| MMTS-AFP-KDEL-2-0009 | 222.00 | 632295.27 | 14.04 | 87.78 |
| MMTS-AFP-KDEL-2-0010 | 276.00 | 1117496.00 | 9.88 | 61.75 |
| MMTS-AFP-KDEL-2-0013 | 346.00 | 1335712.16 | 10.36 | 64.76 |
| MMTS-AFP-KDEL-2-0014 | 250.00 | 953945.67 | 10.48 | 65.52 |
| MMTS-AFP-KDEL-2-0015 | 217.00 | 1026322.47 | 8.46 | 52.86 |
| MMTS-AFP-KDEL-2-0016 | 146.00 | 755872.71 | 7.73 | 48.29 |
| MMTS-AFP-KDEL-2-0018 | 184.00 | 756104.82 | 9.73 | 60.84 |
| MMTS-AFP-KDEL-2-0019 | 101.00 | 532152.95 | 7.59 | 47.45 |
| MMTS-AFP-KDEL-2-0020 | 118.00 | 506049.30 | 9.33 | 58.29 |
| MMTS-AFP-KDEL-3-0003 | 241.00 | 985773.16 | 9.78 | 61.12 |
| MMTS-AFP-KDEL-3-0004 | 274.00 | 1070943.66 | 10.23 | 63.96 |

**Nu**

| Image ID | counts (AuNPs in ROI) | ROI Area (nm$^2$) | Particles in 0.04μm$^2$ | Particles in 0.25μm$^2$ |
|---|---|---|---|---|
| MMTS-AFP-KDEL-1-0001 | 28.00 | 1347233.64 | 0.83 | 5.20 |
| MMTS-AFP-KDEL-2-0003 | 1.00 | 586067.99 | 0.07 | 0.43 |
| MMTS-AFP-KDEL-2-0006 | 2.00 | 246654.36 | 0.32 | 2.03 |
| MMTS-AFP-KDEL-2-0007 | 6.00 | 399262.95 | 0.60 | 3.76 |
| MMTS-AFP-KDEL-2-0008 | 1.00 | 38531.25 | 1.04 | 6.49 |
| MMTS-AFP-KDEL-2-0009 | 2.00 | 85776.08 | 0.93 | 5.83 |
| MMTS-AFP-KDEL-2-0018 | 8.00 | 1623869.04 | 0.20 | 1.23 |
| MMTS-AFP-KDEL-3-0003 | 3.00 | 306054.31 | 0.39 | 2.45 |
| MMTS-AFP-KDEL-3-0004 | 4.00 | 458723.94 | 0.35 | 2.18 |

**Cyto**

| Image ID | counts (AuNPs in ROI) | ROI Area (nm$^2$) | Particles in 0.04μm$^2$ | Particles in 0.25μm$^2$ |
|---|---|---|---|---|
| MMTS-AFP-KDEL-1-0001 | 88.00 | 6343047.42 | 0.55 | 3.47 |
| MMTS-AFP-KDEL-1-0002 | 95.00 | 6427147.22 | 0.59 | 3.70 |
| MMTS-AFP-KDEL-2-0001 | 115.00 | 7305921.76 | 0.63 | 3.94 |
| MMTS-AFP-KDEL-2-0002 | 100.00 | 7516043.68 | 0.53 | 3.33 |
| MMTS-AFP-KDEL-2-0003 | 104.00 | 7127889.69 | 0.58 | 3.65 |
| MMTS-AFP-KDEL-2-0004 | 89.00 | 7456347.27 | 0.48 | 2.98 |
| MMTS-AFP-KDEL-2-0005 | 67.00 | 6910163.61 | 0.39 | 2.42 |
| MMTS-AFP-KDEL-2-0006 | 111.00 | 6108524.55 | 0.73 | 4.54 |
| MMTS-AFP-KDEL-2-0007 | 116.00 | 5997746.85 | 0.77 | 4.84 |
| MMTS-AFP-KDEL-2-0008 | 89.00 | 7925311.12 | 0.45 | 2.81 |
| MMTS-AFP-KDEL-2-0009 | 72.00 | 7475508.67 | 0.39 | 2.41 |
| MMTS-AFP-KDEL-2-0010 | 61.00 | 7905633.79 | 0.31 | 1.93 |
| MMTS-AFP-KDEL-2-0013 | 68.00 | 6194558.04 | 0.44 | 2.74 |
| MMTS-AFP-KDEL-2-0014 | 57.00 | 6882394.24 | 0.33 | 2.07 |
| MMTS-AFP-KDEL-2-0015 | 66.00 | 6507931.05 | 0.41 | 2.54 |

| | | | | |
|---|---|---|---|---|
| MMTS-AFP-KDEL-2-0016 | 59.00 | 7806545.34 | 0.30 | 1.89 |
| MMTS-AFP-KDEL-2-0018 | 63.00 | 6808242.49 | 0.37 | 2.31 |
| MMTS-AFP-KDEL-2-0019 | 49.00 | 7581014.87 | 0.26 | 1.62 |
| MMTS-AFP-KDEL-2-0020 | 53.00 | 4151502.56 | 0.51 | 3.19 |
| MMTS-AFP-KDEL-3-0003 | 66.00 | 6674422.39 | 0.40 | 2.47 |
| MMTS-AFP-KDEL-3-0004 | 111.00 | 7075673.69 | 0.63 | 3.92 |

| Mito | | | | |
|---|---|---|---|---|
| Image ID | counts (AuNPs in ROI) | ROI Area (nm$^2$) | Particles in 0.04µm$^2$ | Particles in 0.25µm$^2$ |
| MMTS-AFP-KDEL-1-0001 | 30.00 | 985804.51 | 1.22 | 7.61 |
| MMTS-AFP-KDEL-1-0002 | 29.00 | 2242816.00 | 0.52 | 3.23 |
| MMTS-AFP-KDEL-2-0001 | 7.00 | 500264.96 | 0.56 | 3.50 |
| MMTS-AFP-KDEL-2-0002 | 2.00 | 332020.67 | 0.24 | 1.51 |
| MMTS-AFP-KDEL-2-0003 | 4.00 | 707075.70 | 0.23 | 1.41 |
| MMTS-AFP-KDEL-2-0005 | 1.00 | 538528.90 | 0.07 | 0.46 |
| MMTS-AFP-KDEL-2-0006 | 9.00 | 714215.64 | 0.50 | 3.15 |
| MMTS-AFP-KDEL-2-0007 | 2.00 | 316801.89 | 0.25 | 1.58 |
| MMTS-AFP-KDEL-2-0008 | 4.00 | 304919.05 | 0.52 | 3.28 |
| MMTS-AFP-KDEL-2-0009 | 0.00 | 18518.41 | 0.00 | 0.00 |
| MMTS-AFP-KDEL-2-0010 | 1.00 | 112974.52 | 0.35 | 2.21 |
| MMTS-AFP-KDEL-2-0013 | 4.00 | 808292.77 | 0.20 | 1.24 |
| MMTS-AFP-KDEL-2-0014 | 2.00 | 625122.87 | 0.13 | 0.80 |
| MMTS-AFP-KDEL-2-0015 | 3.00 | 594196.34 | 0.20 | 1.26 |
| MMTS-AFP-KDEL-2-0016 | 4.00 | 438769.95 | 0.36 | 2.28 |
| MMTS-AFP-KDEL-2-0019 | 6.00 | 926227.46 | 0.26 | 1.62 |
| MMTS-AFP-KDEL-3-0003 | 18.00 | 1261722.67 | 0.57 | 3.57 |
| MMTS-AFP-KDEL-3-0004 | 9.00 | 622631.24 | 0.58 | 3.61 |

| Lyso & MVB | | | | |
|---|---|---|---|---|
| Image ID | counts (AuNPs in ROI) | ROI Area (nm$^2$) | Particles in 0.04µm$^2$ | Particles in 0.25µm$^2$ |
| MMTS-AFP-KDEL-2-0001 | 8.00 | 464421.70 | 0.69 | 4.31 |
| MMTS-AFP-KDEL-2-0002 | 1.00 | 79351.73 | 0.50 | 3.15 |
| MMTS-AFP-KDEL-2-0003 | 10.00 | 105069.49 | 3.81 | 23.79 |
| MMTS-AFP-KDEL-2-0004 | 16.00 | 388716.12 | 1.65 | 10.29 |
| MMTS-AFP-KDEL-2-0005 | 82.00 | 823095.18 | 3.98 | 24.91 |
| MMTS-AFP-KDEL-2-0006 | 10.00 | 163542.63 | 2.45 | 15.29 |
| MMTS-AFP-KDEL-2-0007 | 5.00 | 100871.11 | 1.98 | 12.39 |
| MMTS-AFP-KDEL-2-0013 | 17.00 | 667076.96 | 1.02 | 6.37 |
| MMTS-AFP-KDEL-2-0014 | 5.00 | 766509.74 | 0.26 | 1.63 |
| MMTS-AFP-KDEL-2-0015 | 5.00 | 1099522.67 | 0.18 | 1.14 |
| MMTS-AFP-KDEL-2-0016 | 2.00 | 226784.53 | 0.35 | 2.20 |
| MMTS-AFP-KDEL-2-0018 | 0.00 | 39756.17 | 0.00 | 0.00 |
| MMTS-AFP-KDEL-2-0019 | 5.00 | 188577.24 | 1.06 | 6.63 |
| MMTS-AFP-KDEL-2-0020 | 2.00 | 50598.00 | 1.58 | 9.88 |
