## Supplementary Figure 1-10 for "Direct synthesis of EM-visible gold nanoparticles on genetically encoded tags for single-molecule visualization in cells"

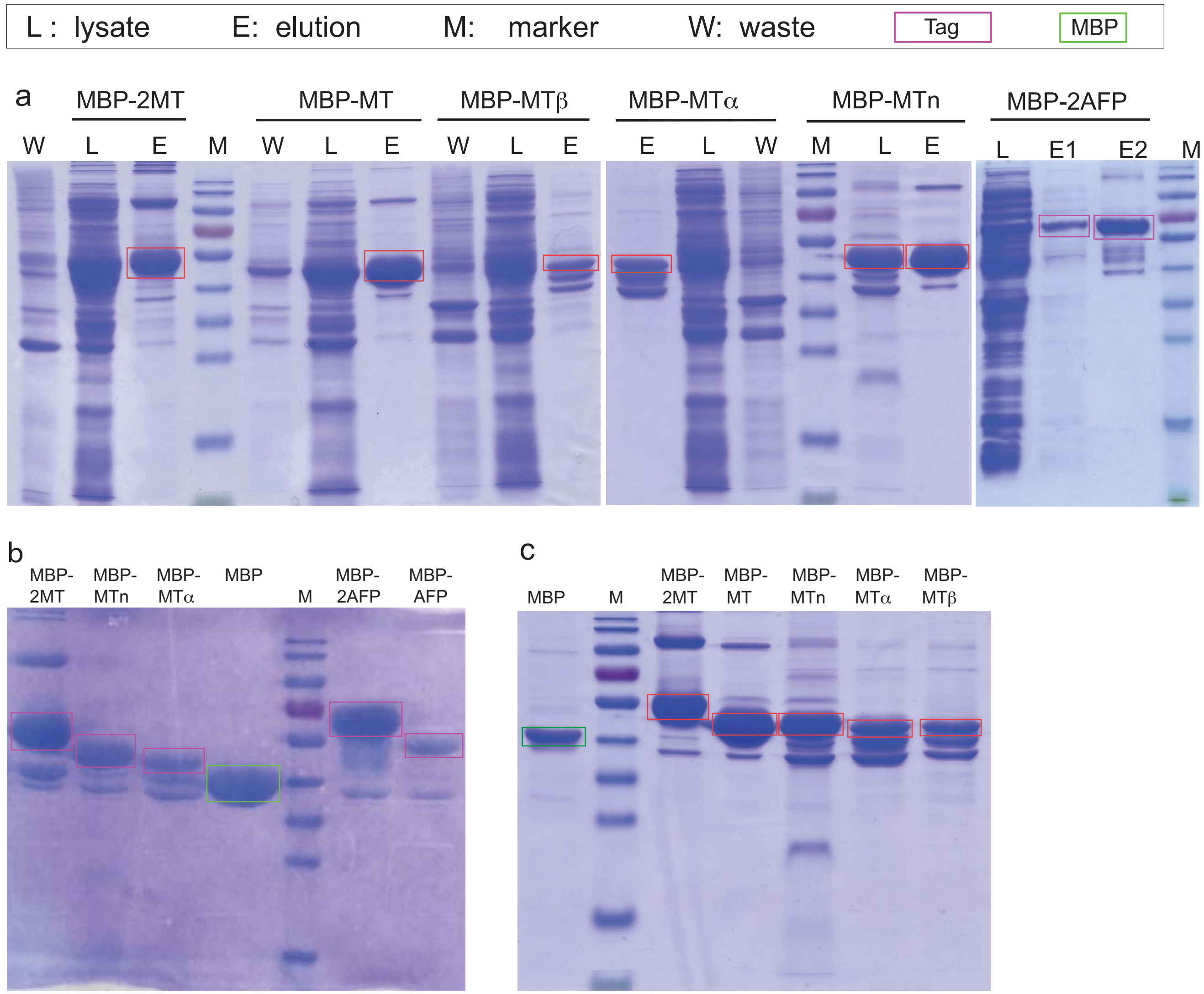


**Supplementary Figure 1 Expression and purification of MBP and MBP-tag fusion proteins**

(**a**) MBP-MT,MBP-2MT, MBP-MTβ,MBP-MTα, MBP-2AFP can be normally expressed in E coli cells (BL21), the SDS PAGE demonstrated the expression levels of those proteins.

(**b**) SDS-PAGE of MBP-2MT, MBP-MTn, MBP-MTα, MBP, MBP-2AFP, MBP-AFP.

(**c**) SDS-PAGE of MBP,MBP-2MT,MBP-MT,MBP-MTn, MBP-MTa,MBP-MTb.

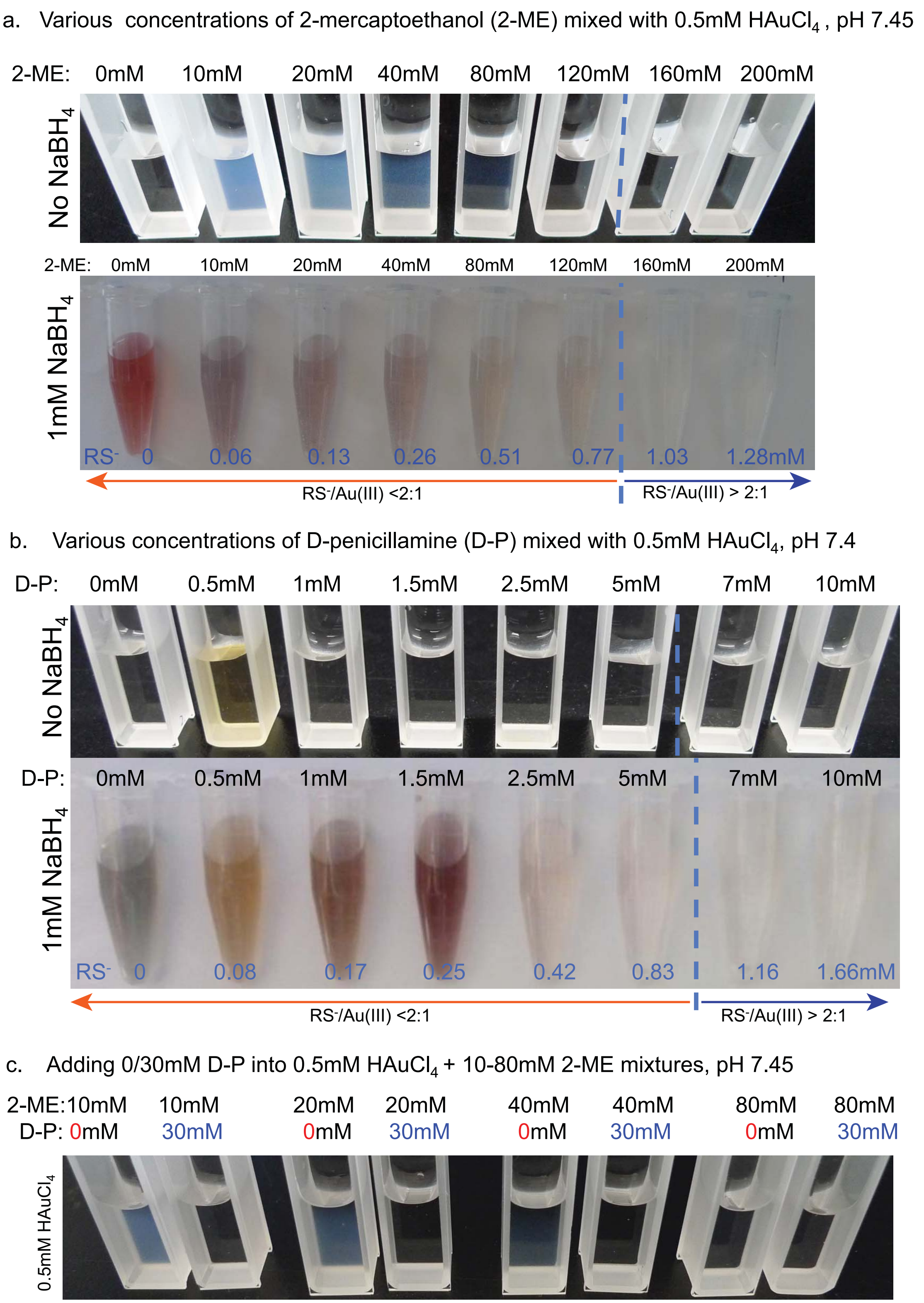


**Supplementary Figure 2 Polymeric Au(I)-thiolate precipitates formed in the mixtures of $HAuCl_4$ and RSH (2-ME or D-P) at RS-/Au(III) < 2:1 conditions.** **(a)** In 2-ME and $HAuCl_4$ mixtures: when RS-/Au(III) < 2:1 , white precipitates of Au(I)-thiolate polymers formed in the mixtures; but no detectable precipitates formed in the RS-/Au(III) >2:1 conditions, the solution is clear. When the Au(I)-thiolates in the mixtures were reduced by 1mM $NaBH_4$ , all those mixtures under the RS-/Au(III) < 2:1 conditions were turned the color to purple or dark brown (indicating the formation of AuNPs); while those mixture under RS-/Au(III) > 2:1 conditions, were still colorless and clear. (b) In D-P and HAuCl4 mixtures: when RS-/Au(III) = 1:1, the solution turned to light brown due to the formation of the unstable (D-P)Au(I) species, but no detectable colors in other condtions; When reduced by 1mM $NaBH_4$, the solutions of those RS-/Au(III) <2:1 cases turned to brown in color, only those RS-/Au(III) > 2:1 cases were still colorless and clear. (c) In a series of 0.5mM $HAuCl_4$ and 10,20,40,80mM of 2-ME mixtures: when added additional 30mM D-P into the 2-ME and 0.5mM $HAuCl_4$ mixtures, the cloudy Au(I)(2-ME) polymers (while precipitates) were dissolved completely and form a colorless clear solution.

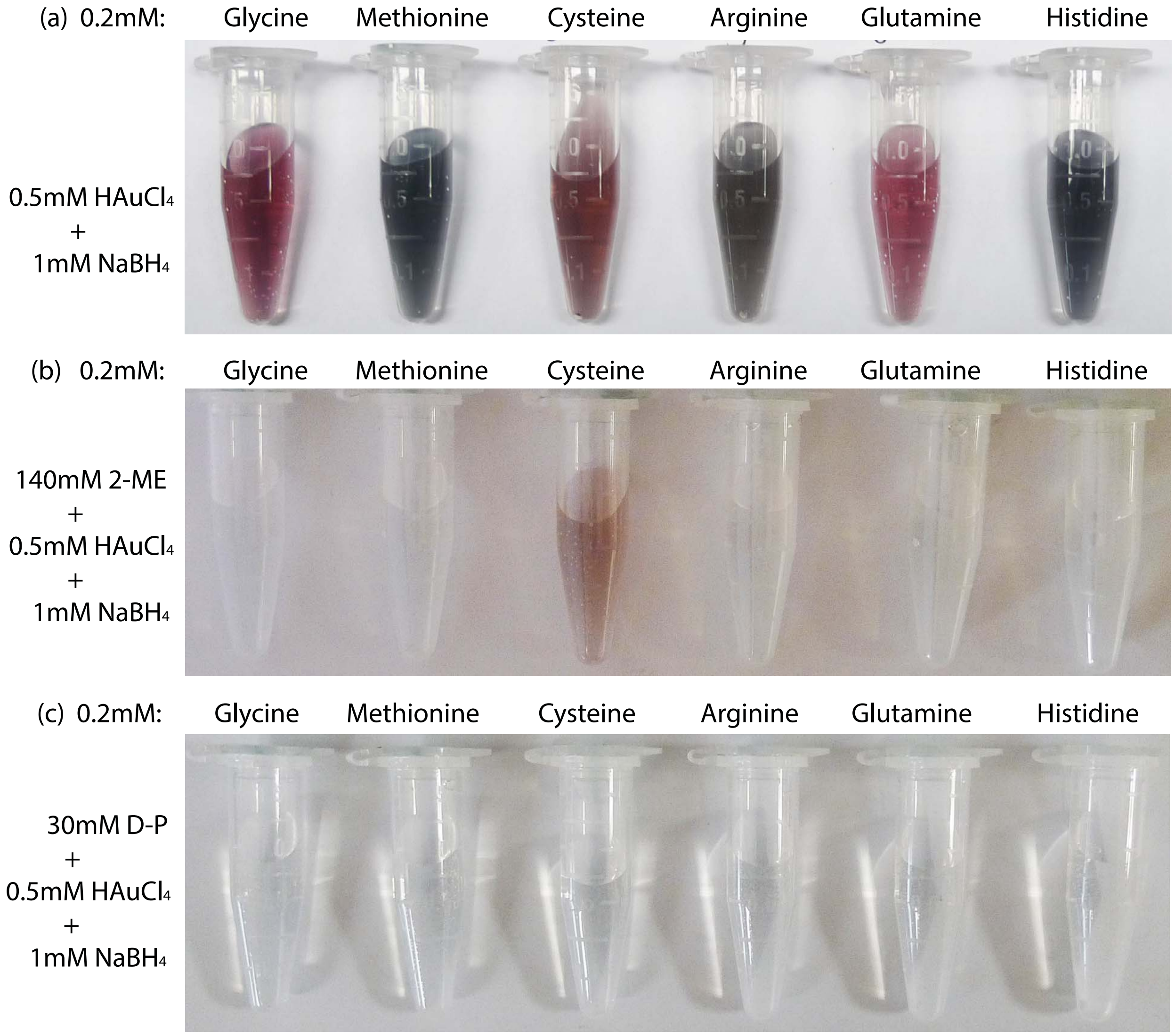


**Supplementary Figure 3 Gold chelating orders of 6 amino acids, 2-mercaptoethanol (2-ME) and d-penicillamine (D-P)**

(a) 0.2mM of 6 typical amino acids and 0.5mM $HAuCl_4$ mixtures reduced by 1mM $NaBH_4$, the solutions all changed to purple or dark brown in color, indicating the formation of AuNPs. (b) 0.2mM of 6 typical amino acids first mixed with 140mM 2-ME and 0.5mM $HAuCl_4$, then reduced by 1mM $NaBH_4$, only the solution with cysteine turned to brown color, the rests are in colorless. (c) 0.2mM of 6 typical amino acids first mixed with 30mM D-P and 0.5mM $HAuCl_4$, then reduced by 1mM $NaBH_4$, all the solution are clear and colorless. These experiments implied: the strongest gold chelating ligand, thiol group of D-P, suppressed all the weaker ligands, thiol group of cysteine residue and the -$NH_2$ or -COOH groups in other amino acids; the thiol group of 2-ME ligand effectively suppressed the -$NH_2$ or -COOH group in other amino acids, but could not suppress the stronger thiol group of cysteine residue. Therefore,these experiments implied the gold chelating orders (from weak to strong): -COOH,-$NH_2$ group of amino acid< thiol group of 2-ME < thiol group of cysteine < thiol group of D-P. The (b) and (c) demonstrated the capabilites of 2-ME and D-P for auto-nucleation suppression.

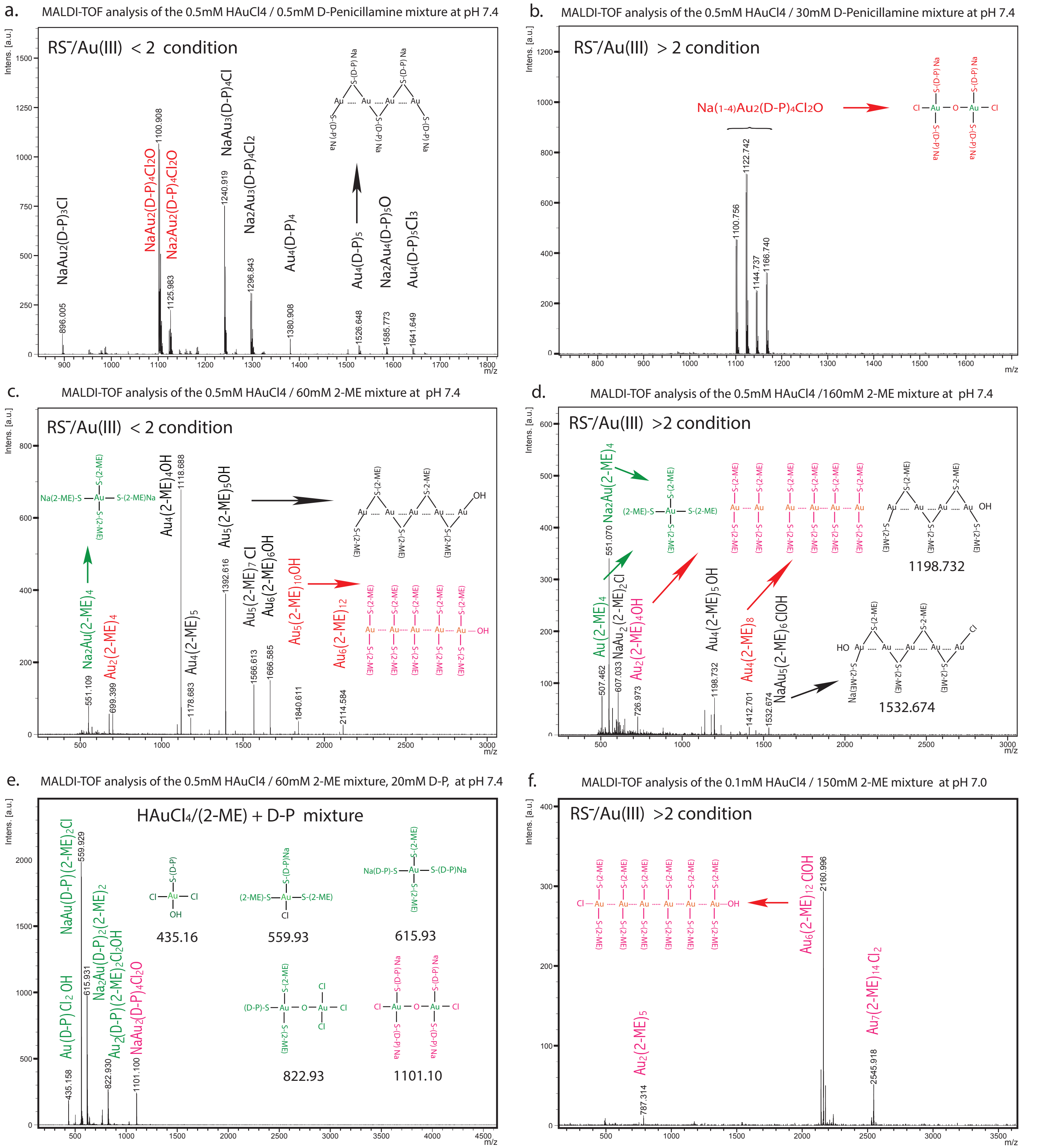


**Supplementary Figure 4 Identification of gold-thiolate species in the mixtures of $HAuCl_4$ and RSH by MALDI-TOF analysis**
(**a**) Two types of gold thiolate compounds identied in the 1:1 Au(III)/(D-P) mixture (at RS-/Au(III) <2 :1 condition): $Au(I)_n(D\text{-}P)_{n+1}$, (n=2,3,4) in zigzag chain forms (labelled in black), and the $Na_{(1\text{-}2)}Au_2(D\text{-}P)_4Cl_2O$ compounds (labeled in red). (**b**) Only the 1:2 Au(III)/(D-P) compounds (in red), $Na_{(1\text{-}2)}Au_2(D\text{-}P)_4Cl_2O$, found at RS-/Au(III) > 2 :1 condition. (**c**) In the mixture of 0.5mM HAuCl4 +60mM 2-ME (RS-/Au(III) <2 :1 condition), the major Au(I) thiolate species were zigzag chain-like 1:1 Au(I)/(2-ME) compouds (in black), mixed with a little $Na_2Au(III)(2\text{-}ME)_4$ (in green) and 1:2 Au(I)/(2-ME) compounds (in red). (**d**) In the mixture of 0.5mM HAuCl4+160mM 2-ME (RS-/Au(III) >2 :1 condition), the mount of zigzag chain-like 1:1 Au(I)/(2-ME) compounds (in back) were largely reduced compared with (c), and $Na_{(0\text{-}2)}Au(III)(2\text{-}ME)_4$ ( in green) became the major spieces, and also some 1:2 Au(I)/(2-ME) compounds (in red). (**e**) Added 20mM D-P into the 0.5mM $HAuCl_4$ + 60mM 2-ME mixture, compared with (c), all the chain-like 1:1 Au(I)/(2-ME) compounds (in black) were disappeared, only the small Au(III) compounds (in green) and a little $NaAu_2(D\text{-}P)_4Cl_2O$ (red) found in the mixture. (**f**) In the 0.1mM $HAuCl_4$ and 150mM 2-ME mixture, only three 1:2 Au(I)/(2-ME) compounds (red) formed in the mixture at RS-/Au(III) > 2 :1 condition.

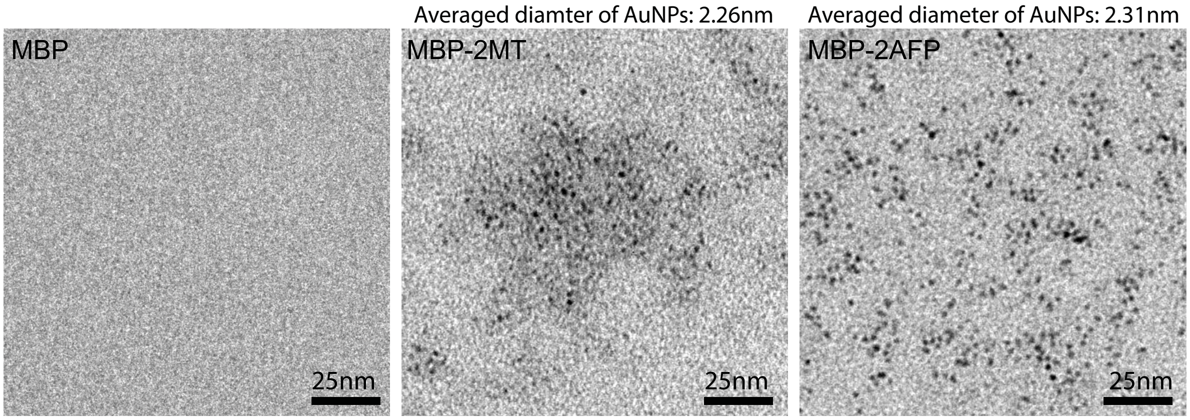


**Supplementary Figure 5 EM image of AuNPs synthesized on MBP, MBP-2MT and MBP-2AFP with Brush-Schiffrin method performed at ANSM conditions achieved with 30 mM D-penicillamine**

The experiments performed under conditions: 2.5μM protein, 30mM D-P, pH 7.4, reduced by 1mM $NaBH_4$. The corresponding solutions are shown in those tubes labeled as 30mM D-P (the most right ones) in Figure 2 (e-g).

The average diameter ~2.3nm AuNPs found in MBP-2MT and MBP-2AFP samples, but no AuNPs detected in MBP samples.

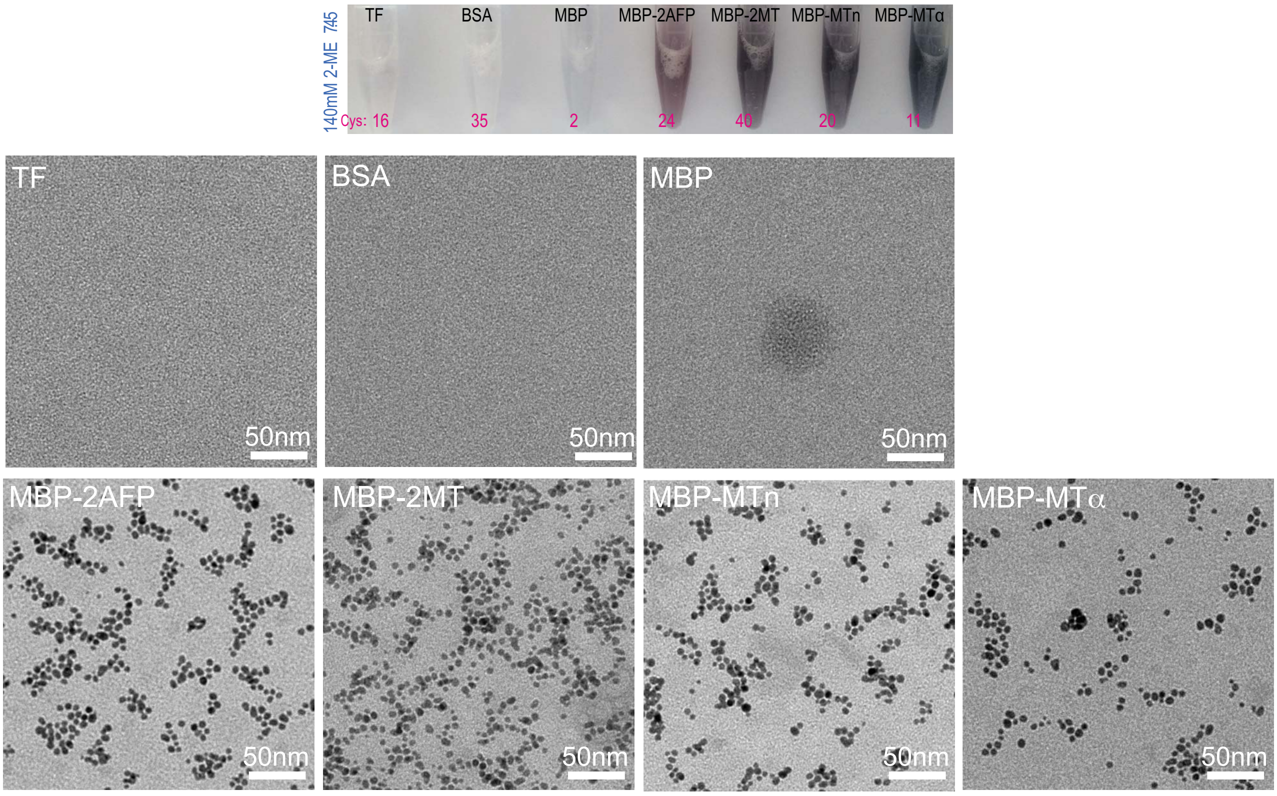


**Supplementary Figure 6**. **TEM images of ANSM synthesizing AuNPs on 7 proteins with the 2-ME scheme**. 2.5uM of corresponding protein and 10uL (140mM) 2-ME were added into 1mL DDH$_2$O (pre-adjusted PH to 7.45) for 30 minutes at room temperature, then reduced by 1mM NaBH$_4$ for AuNPs synthesis. No AuNPs formed in the 3 control proteins: transferrin (TF), BSA and MBP, but in the MBP case found some blurred blobs. The 4 tag-MBP fusion proteins all formed 4-7nm AuNPs, the AuNPs tends to aggregate slightly.

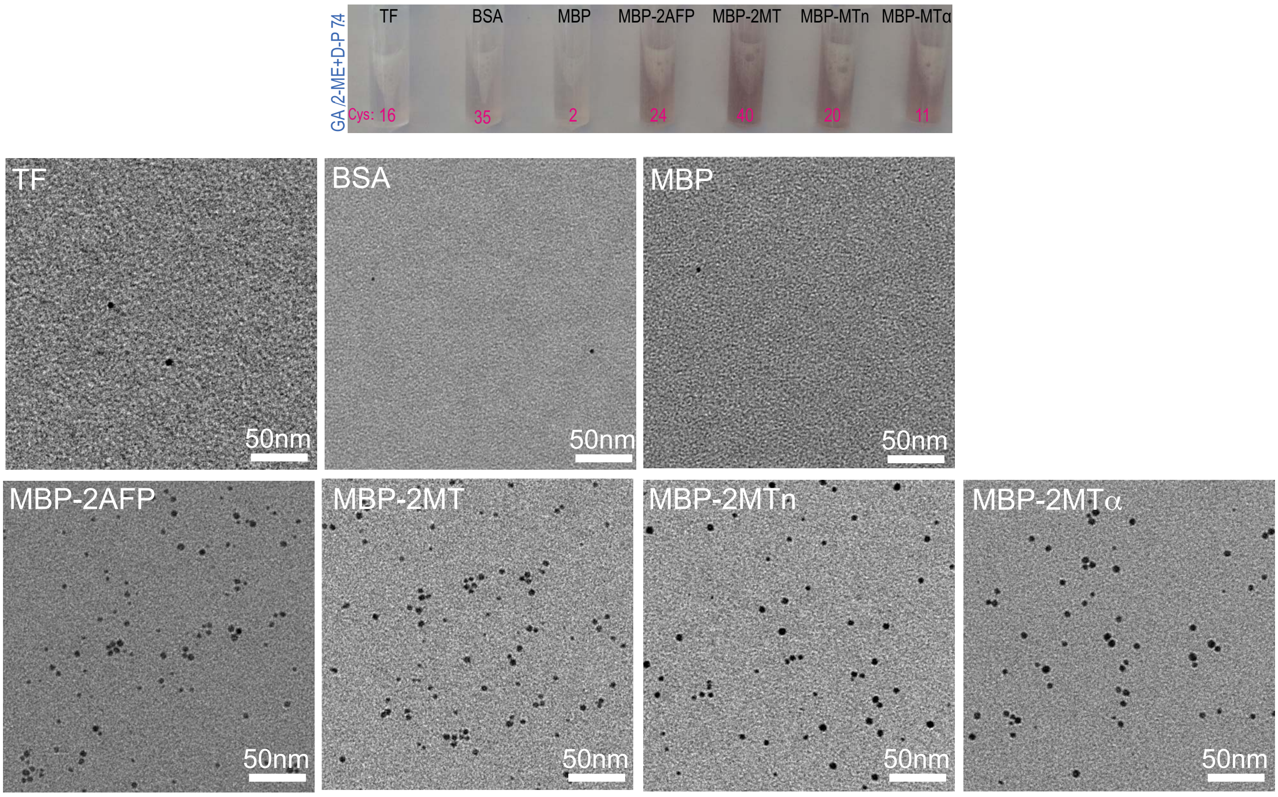


**Supplementary Figure 7 EM images of AuNPs synthesized on 1% GA pre-fixed proteins with the 2-ME/D-P protocol.** 2.5uM of corresponding proteins pre-fixed with 1% GA (for 30min) and neutralized with 5% glycine (30min), then filtered for AuNPs synthesis with the 2-ME/D-P protocol (**Fig.2h,k**) . Only occasionly found a few AuNPs in the 3 control proteins (TF,BSA and MBP), perhaps caused by the unneutralized fixatives.The 4 tag-MBP fusion proteins all formed 3-4nm AuNPs.

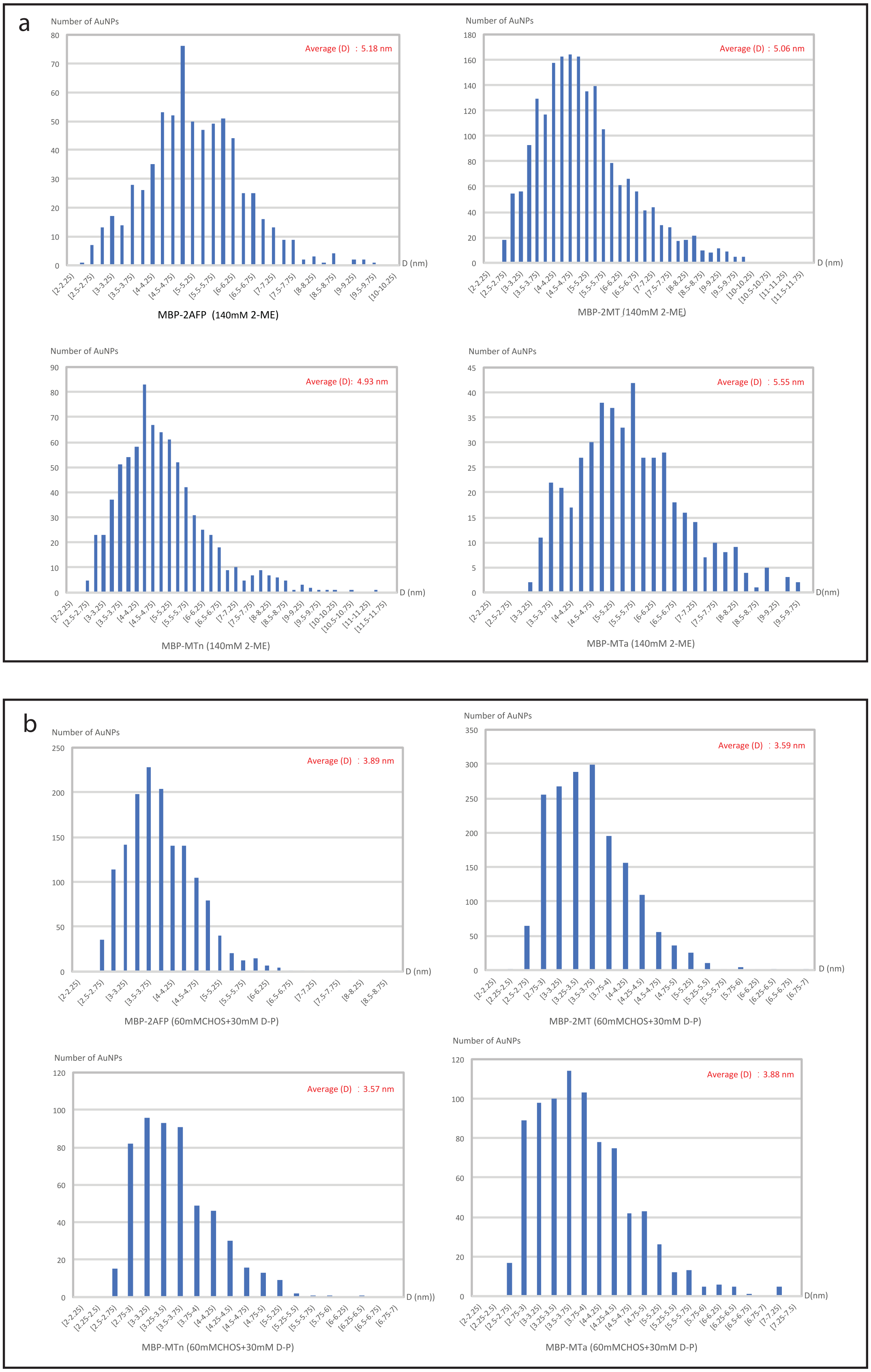


**Supplementary Figure 8 Diameter distributions of AuNPs synthesized on 4 cysteine-rich tags (2AFP, 2MT,MTn and MT$\alpha$) with the 2-ME protocol and the 2-ME/D-P protocol.**

(a) The average diamter of AuNPs synthesized on 4 tags by the 2-ME protocol (2.5 uM proteins, 140 mM 2-ME, 0.5mM $HAuCl_4$, 1mM $NaBH_4$, pH 7.45). (b) The average diameter of AuNPs synthesized on 4 tags by the 2-ME/D-P protocol (2.5 uM proteins, 60 mM 2-ME, 0.5mM $HAuCl_4$, 30mM D-P, 1 mM NaBH4, pH 7.4).

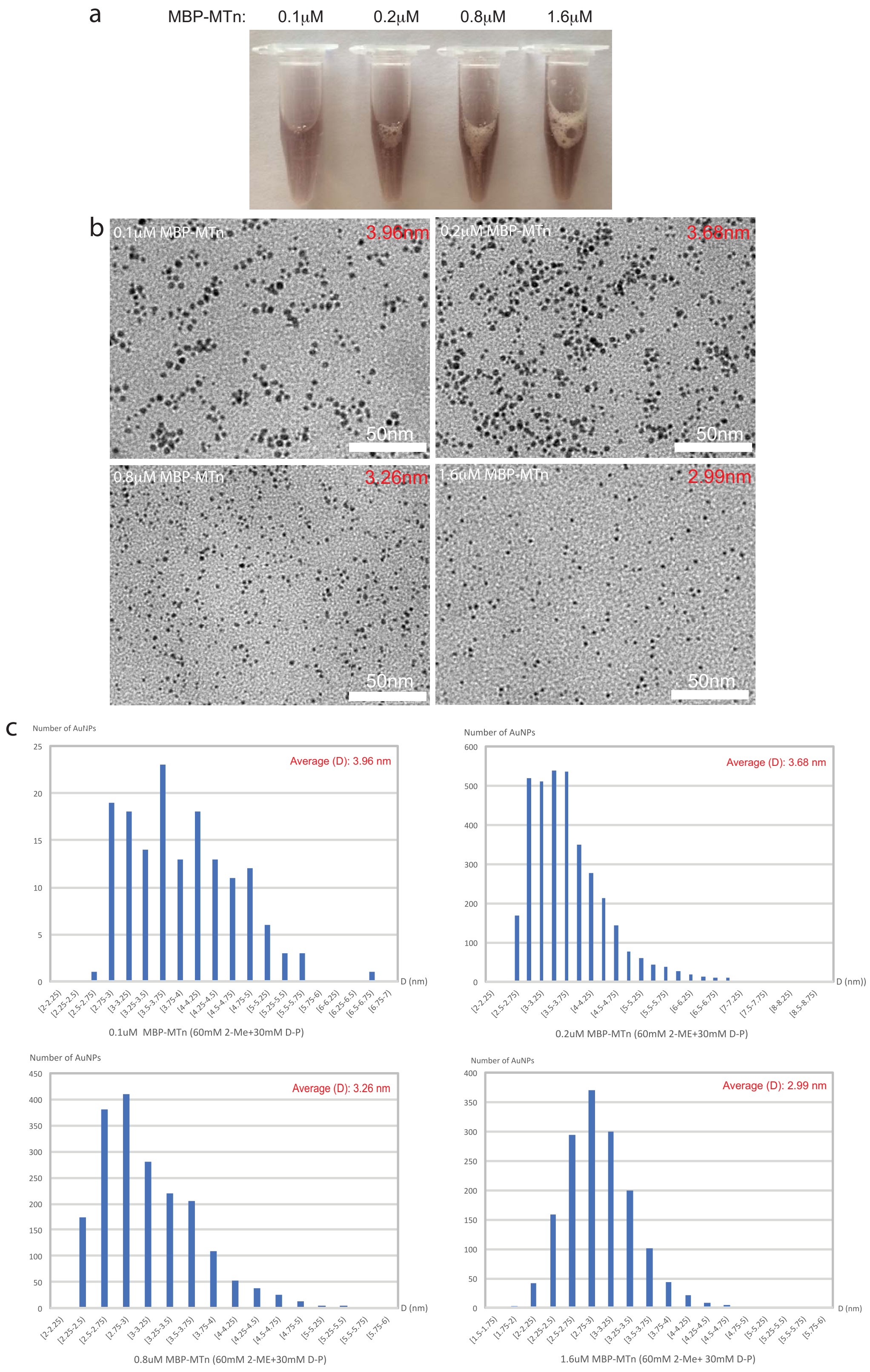


**Supplementary Figure 9 Diameter distributions of AuNPs synthesized at various cencontrations of MBP-MTn at PH 7.45, with 2-ME/D-P protocol**. The average diameters for 0.1μM,0.2μM,0.8μM, 1.6μM are : 3.96nm, 3.68nm, 3.26nm and 2.99nm respectively. (a) The AuNPs formed in solutions as indicated by the dark brown colors. (b) The EM images of the MBP-MTn concentration series, showing the diameters of AuNPs. (c) The corresponding diameter distributions of AuNPs formed in the MBP-MTn concentraton series.

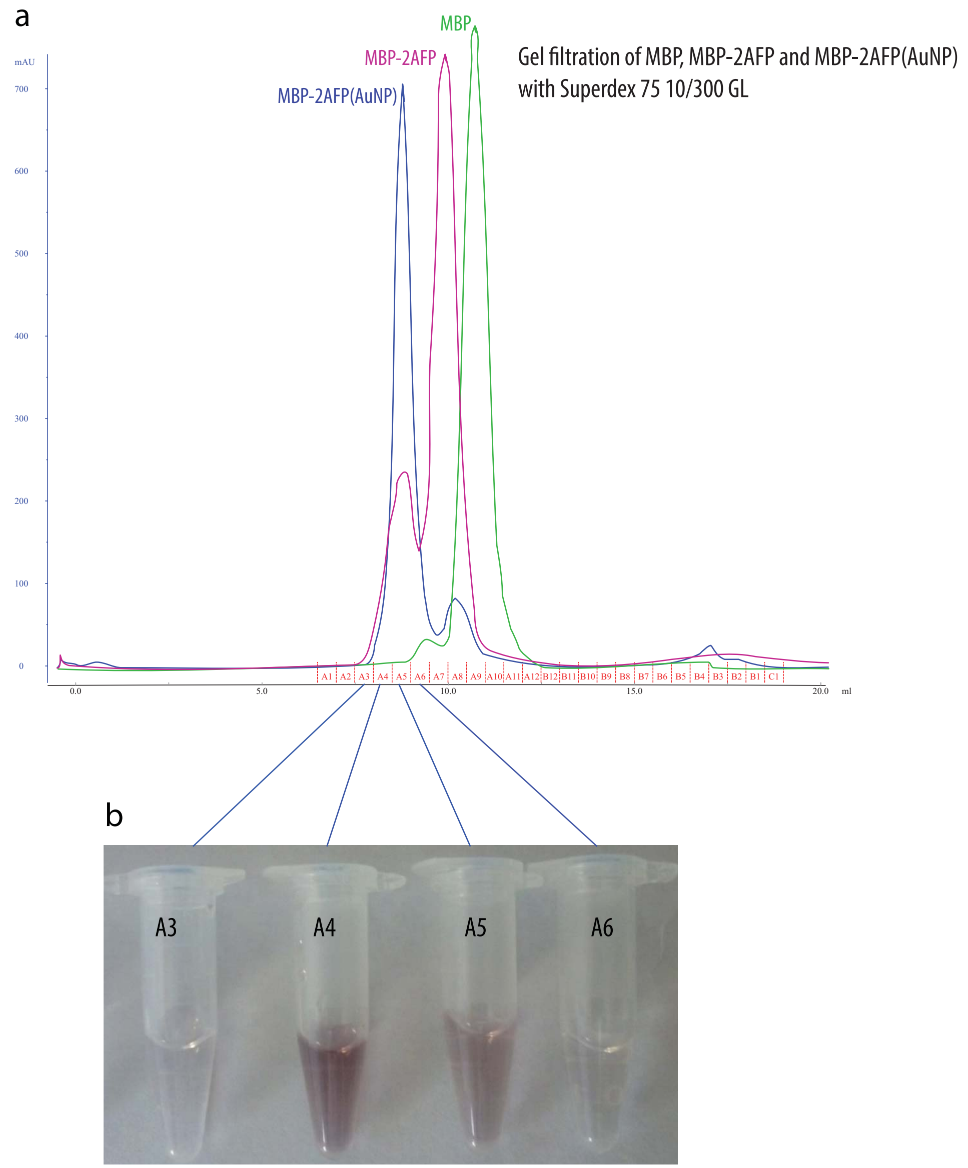


**Supplementary Figure 10A AuNPs formed on MBP-2AFP might be mainly in monomer forms and in similar sizes**
(a) Gel filtration chromatograophy of MBP (42.7kDa), MBP-2AFP(57.2kDa) and MBP-2AFP with AuNPs: (i) MBP and MBP-2AFP were mainly in monomer forms as indicated by a major narrow ans sharp peak, only a small portion in aggregates forms (conformed by the corresponding SDS PAGE data showed in **Supplementary Fig.1**); (ii) after AuNPs synthesized on MBP-2AFP proteins, the major peak shifted from A7-A8 (MBP-2AFP) to A4-A5 (MBP-2AFP(AuNP)), but remained a narrow and sharp peak, which impied that MBP-2AFP (AuNP) might be mainly still in monomer forms. (b) Solution collected from tubes (A3 to A6) of the narrrow peak of MBP-2AFP(AuNP) as showed in (a), the featured brown color and clear solution of A4-A5 tubes indicated that the MBP-2AFP(AuNP) were stably dispersed in those tubes.

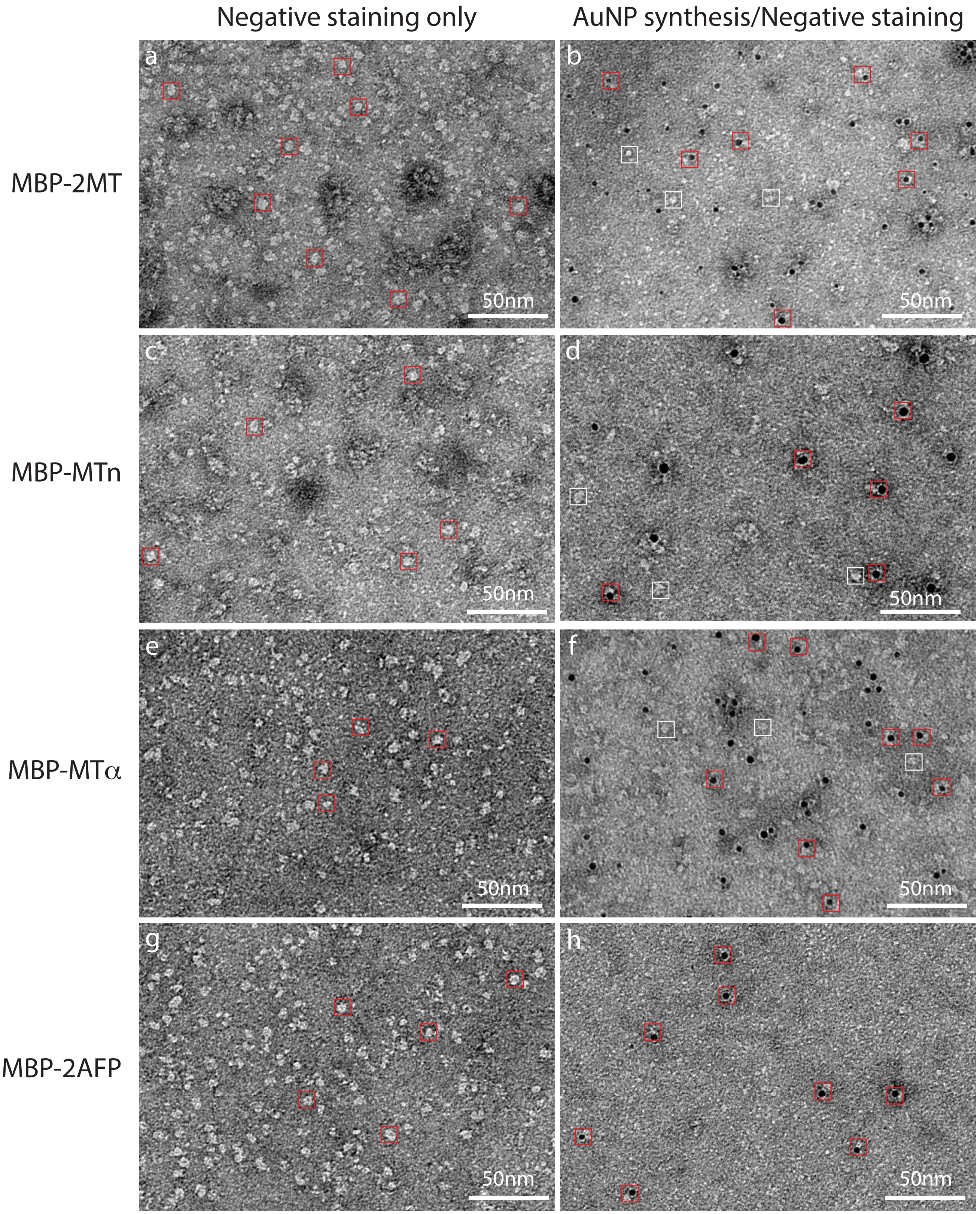


**Supplementary Figure 10B Single-molecule level EM imaging of MBP-tag fusion proteins (MBP-2MT, MBP-MTn, MBP-MTα and MBP-2AFP) subjected to AuNP synthesis followed by negative staining**

(**a,c,e,g**) EM images of negative stained MBP-2MT, MBP-MTn, MBP-MTα and MBP-2AFP molecules without AuNP synthesis, the individual molecules (~7nm in sizes) were marked with a 8nm x 8nm red boxes. (**b,d,f,h**) EM images of MBP-2MT, MBP-MTn, MBP-MTα and MBP-2AFP molecules underwent ANSM-based AuNP synthesis and negative staining, the 8nm x 8nm red boxes marked out these proteins forming AuNPs on the tags in 1:1 ratio, the 8nm x 8nm white boxes marked out those proteins without forming visible AuNPs which might be caused by the insufficient access of gold thiolate sources due to physical hinder (e.g., wrapped by other proteins). To avoid multiple AuNPs aggregation with the standard 2-ME/D-P AuNP syntheis protocol (**Fig.2**), here a stronger reducing reagent TCEP was used for reducing the tags, a TCEP/D-P AuNP synthesis protocol used for (b,d,f,h) (0.2 mM TCEP reducing 30 min, 3 mM D-P mixed with 1mM $HAuCl_4$ for 30 min, add extra 10mM D-P quickly mixed, then immediately reduced with 1mM $NaBH_4$ ). Note: all the proteins were subjected to negative stained with 2% UA.
