## Supplementary Figure 11-15 for "Direct synthesis of EM-visible gold nanoparticles on genetically encoded tags for single-molecule visualization in cells"

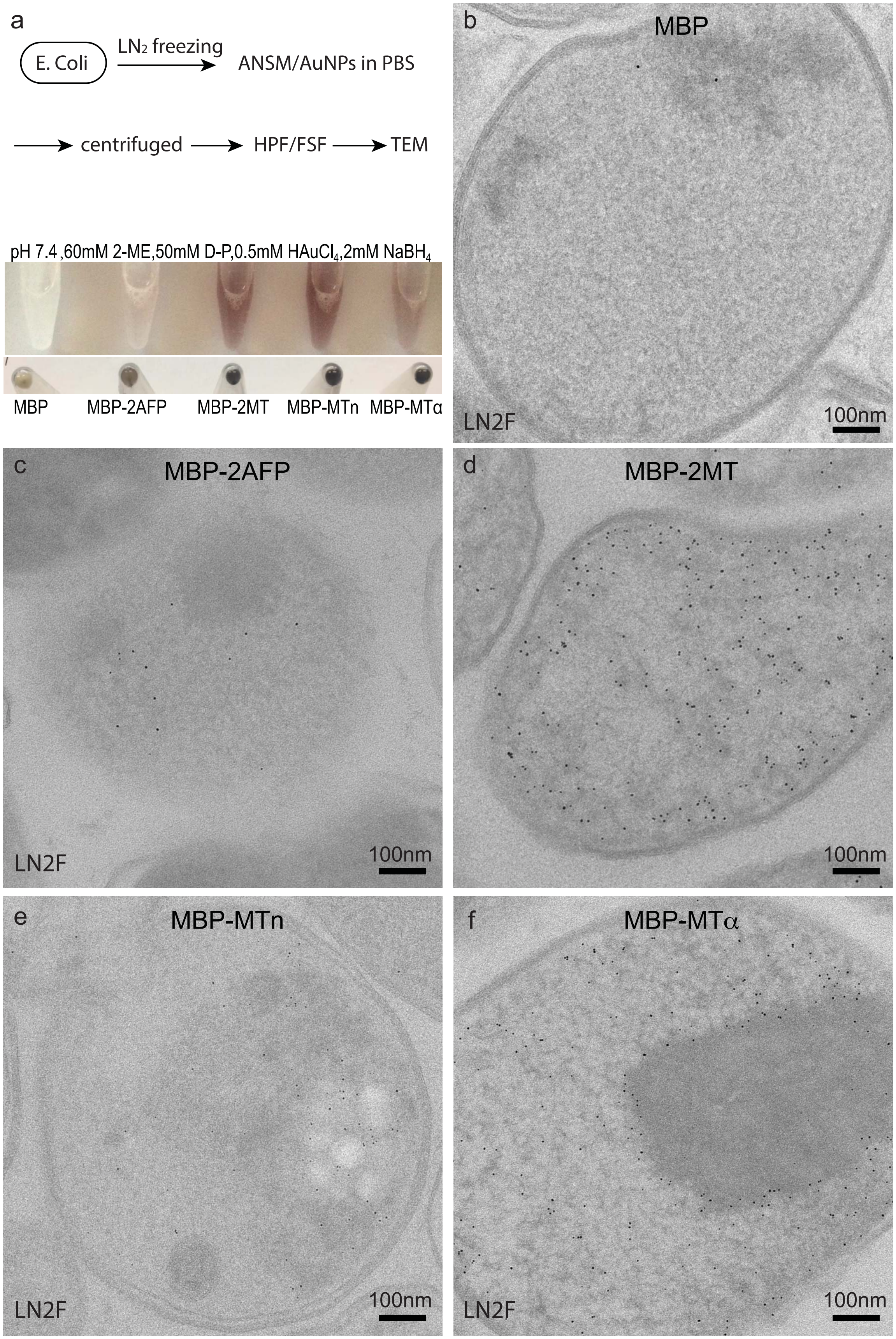


**Supplementary Figure 11 Optimization of the conditions for ANSM-based AuNPs synthesis in *E. coli* cells by the disruption of cell membranes with liquid nitrogen freezing (LN2F).** (a) Scheme for the screening condition for AuNPs synthesis, one of the optimal condition was used for ANSM-based AuNPs synthesis in *E. coli* cells (as shown in Fig.3b-c): the control (MBP) has no color change, the 4 tags (2AFP, 2MT,MTn, MTα ) are purple to brown in color, indicating the formation of AuNPs. (b-f) the corresponding EM images of the 5 specimens: almost no AuNPs detected in MBP expressed cell, in contrast, a lot of AuNPs observed in the rest cells expressing 4 tags (2AFP,2MT,MTn and MTα).

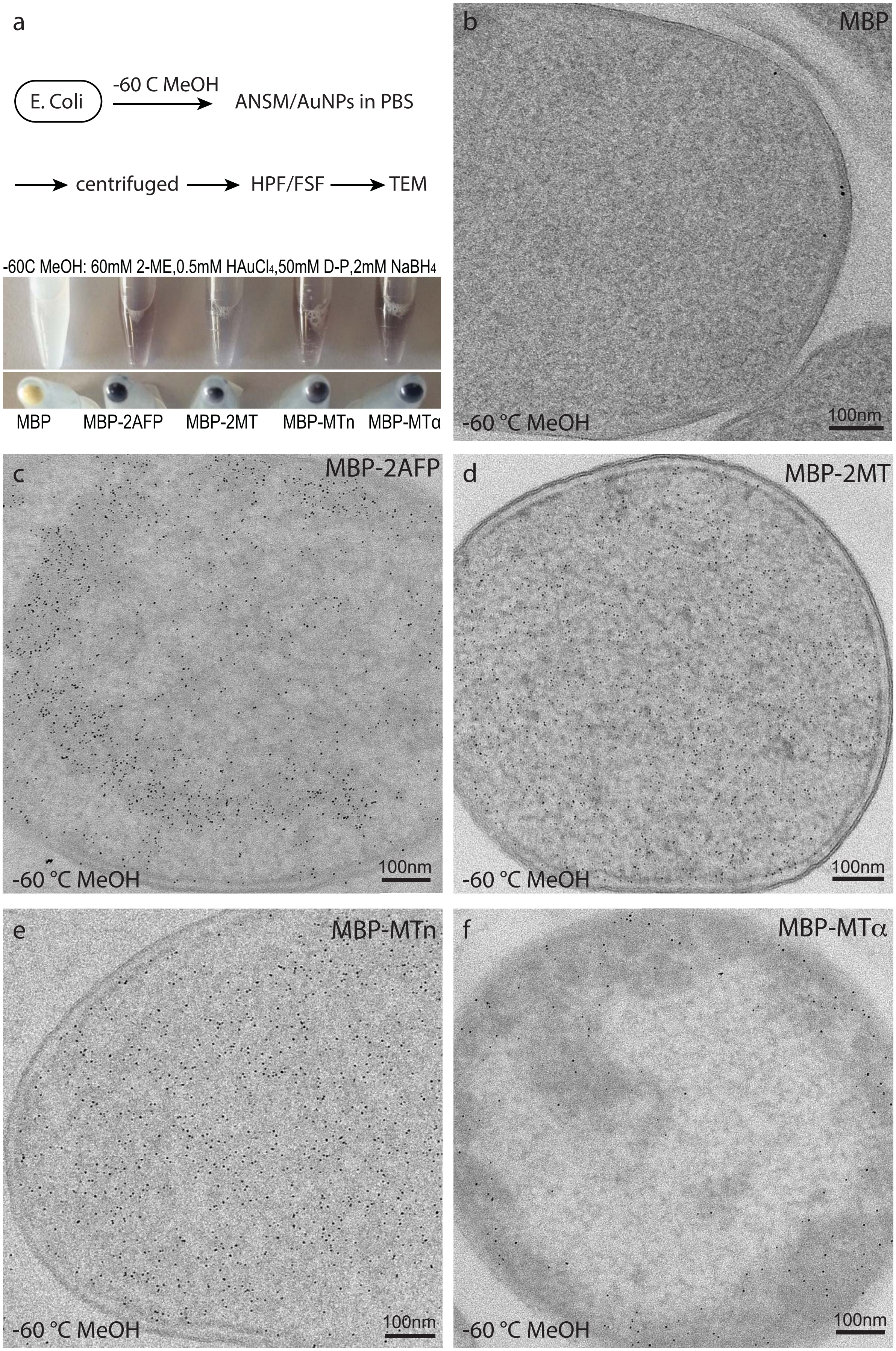


**Supplementary Figure 12 ANSM-based synthesis of AuNPs on cysteine-rich tags expressed in *E. coli* cells treated by -60 ˚C methanol**. (a) Scheme for prepare the specimen for AuNPs synthesis and the the colors of the cells in solutions and the corresponding centrifuged pellets. (b-f) EM images of the cells of 5 *E. coli* strains prepared by the scheme described in (a).

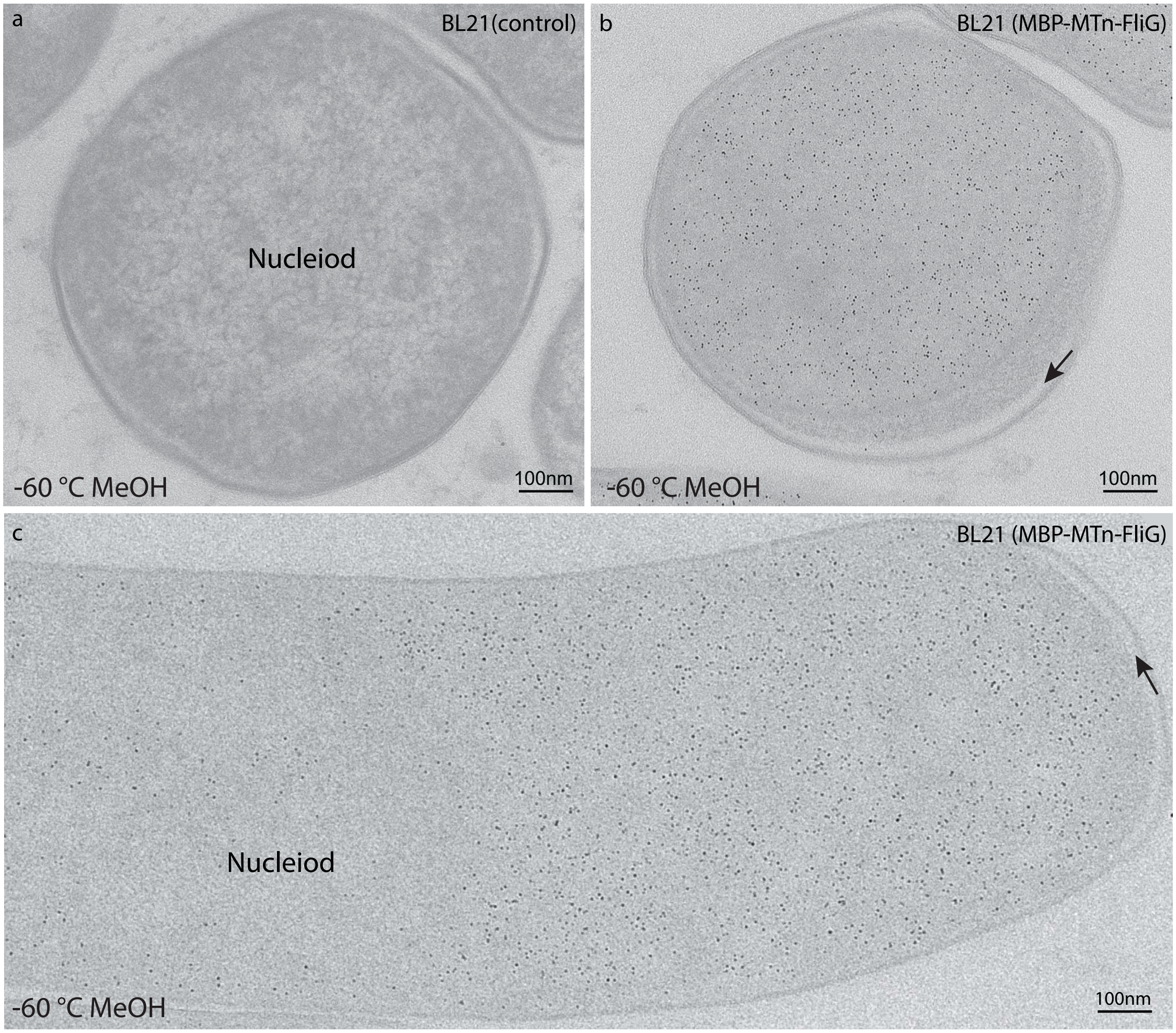


**Supplementary Figure 13A AuNPs specific distributions in the *E. coli* cells overexpressing MBP-MTn-FliG.** (a) Almost no AuNPs formed in the wild-type BL21(DE3) cells ; (b) AuNPs distributed relative unifrom in a cross-section of a E. coli cell expressing MBP-MTn-FliG; (c) AuNPs aaccumulated at the pole of a E. coli cell expressing MBP-MTn-FliG, only a few AuNPs found in the nuceiod region, but no AuNPs were found in the periplasmic spaces (pointed by the black arrows). Specimen prepared by -60˚C methanol freezing for 1-2 min, then centrifuged to remove the methanol and changed to 4˚C PBS-A for ANSM -based AuNPs synthesis with the 2-ME/D-P protocol.

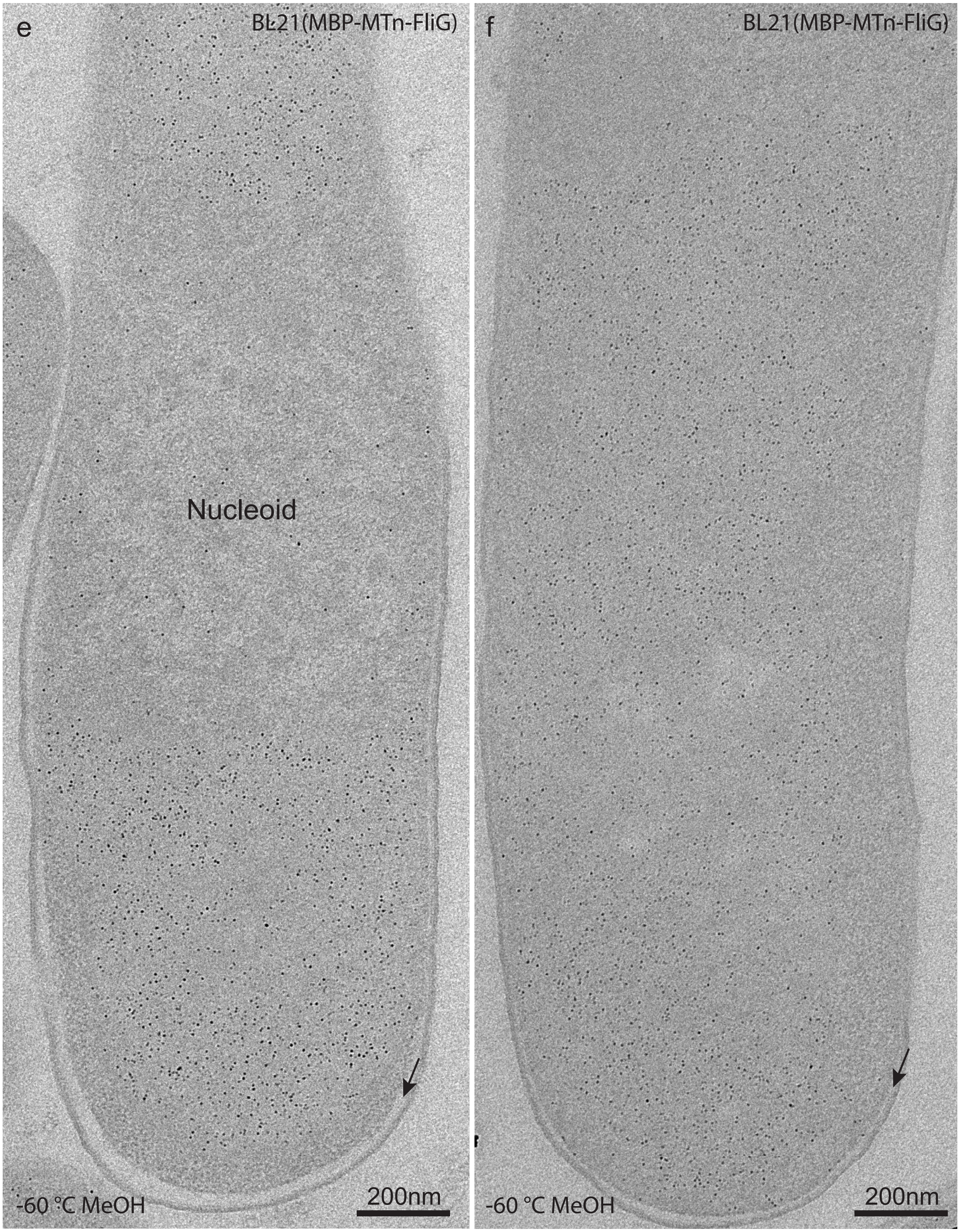


**Supplementary Figure 13B AuNPs specific distributions in the *E. coli* cells overexpressing MBP-MTn-FliG.** (e-f) AuNPs accumulated at two poles of the cell, but were seldom seen in the nucleoid region and no AuNPs in the periplasmic spaces (pointed by the black arrows); Specimen prepared by -60˚C methanol freezing for 1-2 min, then centrifuged to remove the methanol and changed to 4˚C PBS-A for ANSM -based AuNPs synthesis with the 2-ME/D-P protocol. The expressed MBP-MTn-FliG molecues tended to accumulate in two poles of the cell, but the AuNPs were well dispersed and the proteins could be purified from the supernant, therefore, it is uncertain whether those proteins were in inclusion bodies or not. Anyway, suchcharacteristic distributions indicated that the AuNPs sysnthesis protocol worked very effectively.

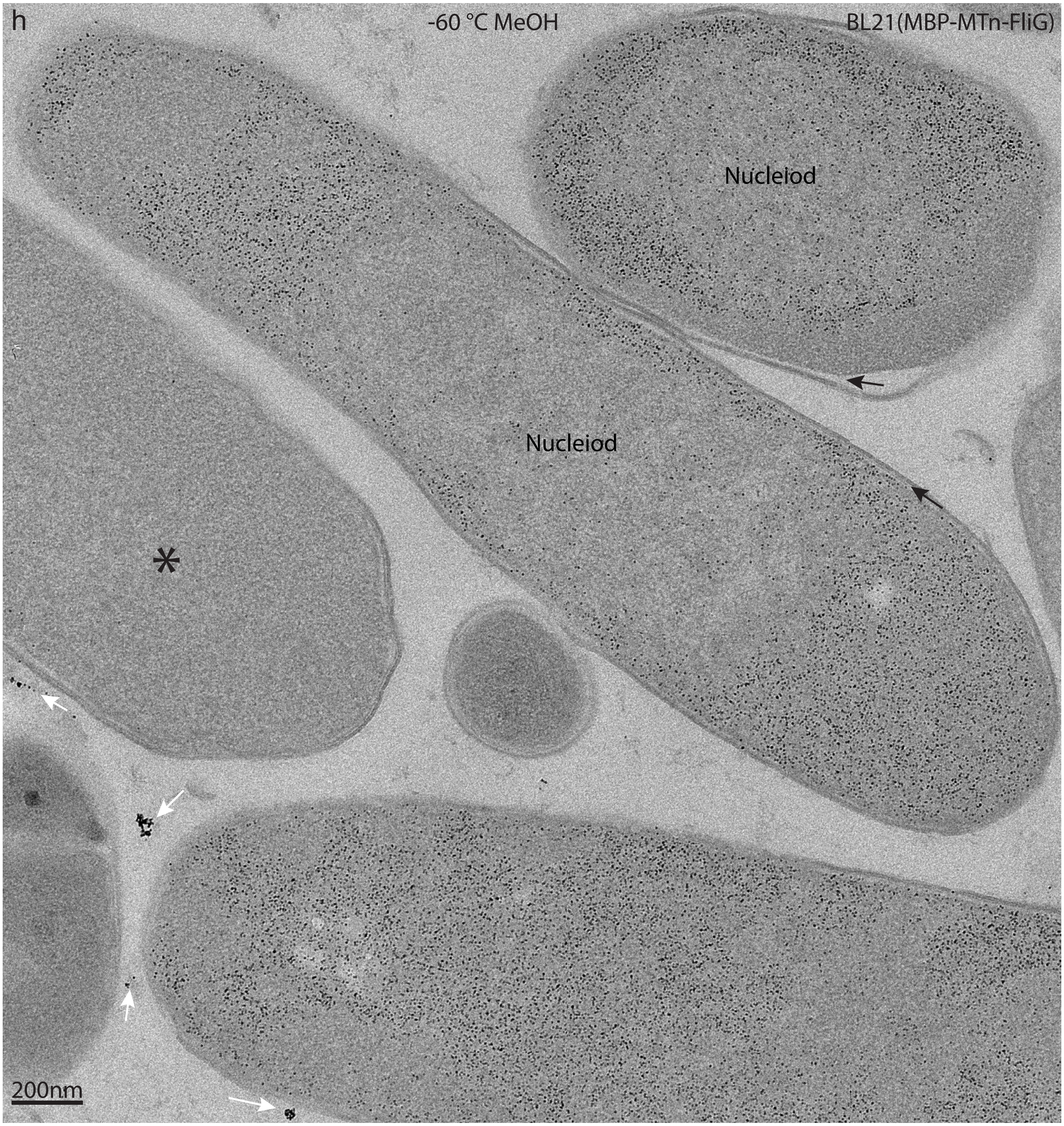


**Supplementary Figure 13C AuNPs specific distributions in the *E. coli* cells overexpressing MBP-MTn-FliG**
(h) AuNPs highly accumulated at two poles of the cell but not the nucleiod region, interestngly,also a lot of AuNPs surrounding the nucleiod of the E. coli cell, which implied that the MBP-MTn-FliG proteins were in normally distribution instead of in the possible inclusion body; Specimen prepared by -60˚C methanol freezing for 1 min, then centrifuged to remove the methanol and changed to 4˚C PBS-A for ANSM-based AuNPs synthesis with the 2-ME/D-P protocol. It should be noted that almost no AuNPs foundin the periplasmic spaces (pointed by black arrows), only a few AuNPs in the nucleiod regions, outside the cells looks quite clean excepta few dark aggregates (pointed by white arrows), which might be formed by the undissolved Au(I)(SR) polymers. However, such polymers usually are too big to penetrate into the cells, thus they are unlikely to interfere the tags inside the cells. There is almost no AuNPs found in one cell marked by a black asterisk, it might be interpreted by a cross-section of nucleiod region or a cell lost most of the plasmids of the MTn tags.

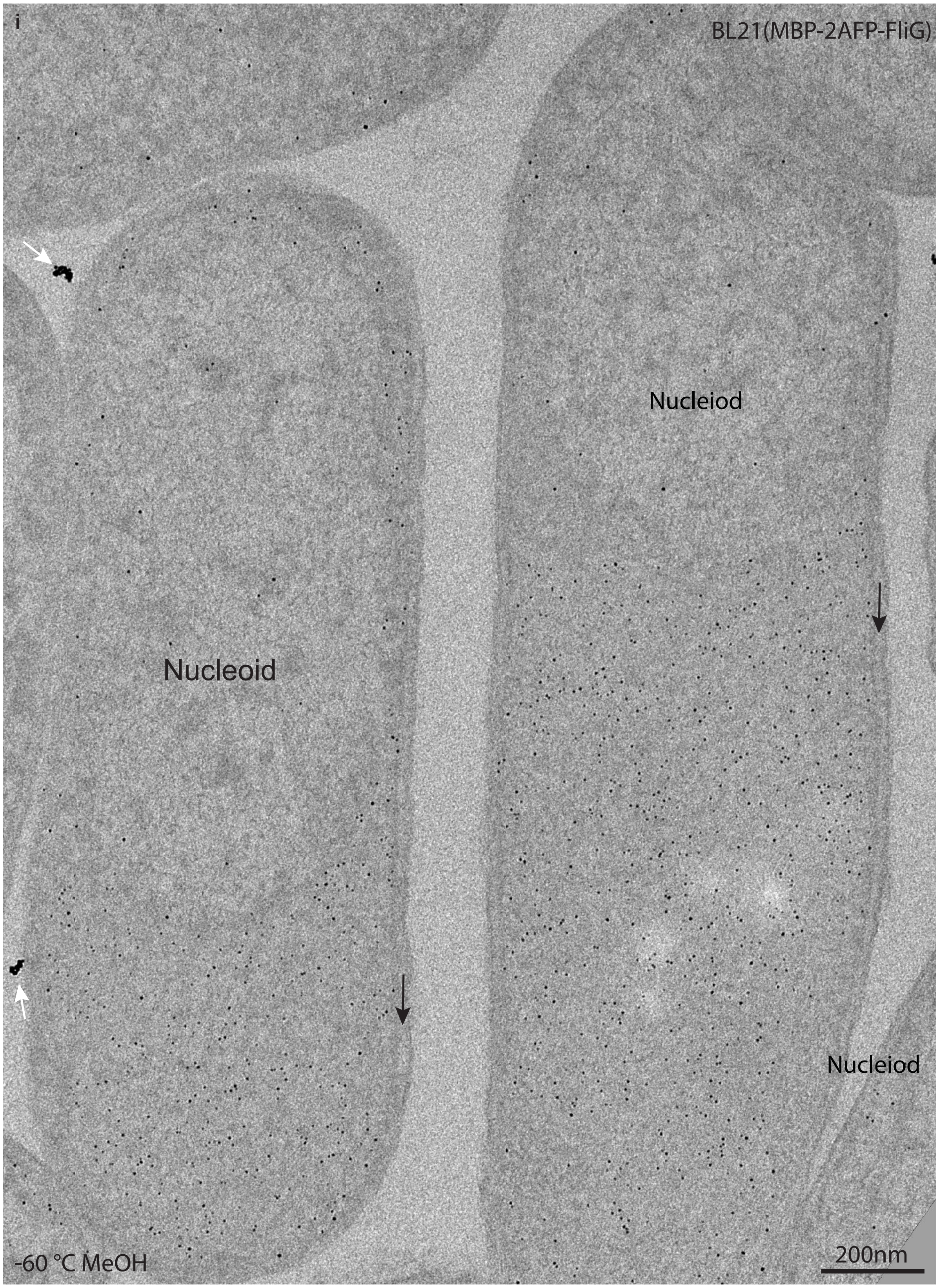


**Supplementary Figure 13D AuNPs specific distributions in the *E. coli* cells overexpressing MBP-2AFP-FliG.** (i) AuNPs accumulated at poles of the cells, but much less AuNPs in Nucleiod regions; Specimen prepared by -60˚C methanol freezing for 1 min, then centrifuged to remove the methanol and changed to 4˚C PBS-A for ANSM-based AuNPs synthesis with the 2-ME/D-P protocol. There is almost no AuNPs found in the periplasmic spaces (pointed by the black arrows). Outside the cells looks quite clean excepta few dark aggregates (pointed by white arrows), which might be formed by the undissolved Au(I)(SR) polymers.The overall feature of the MBP-2AFP-FliG distribution is quite similar to those of MBP-MTn-FliG (refer to Supplementary Figure 13C(h)) , except the densities of the AuNPs are less in MBP-2AFP-FliG expressing cells.

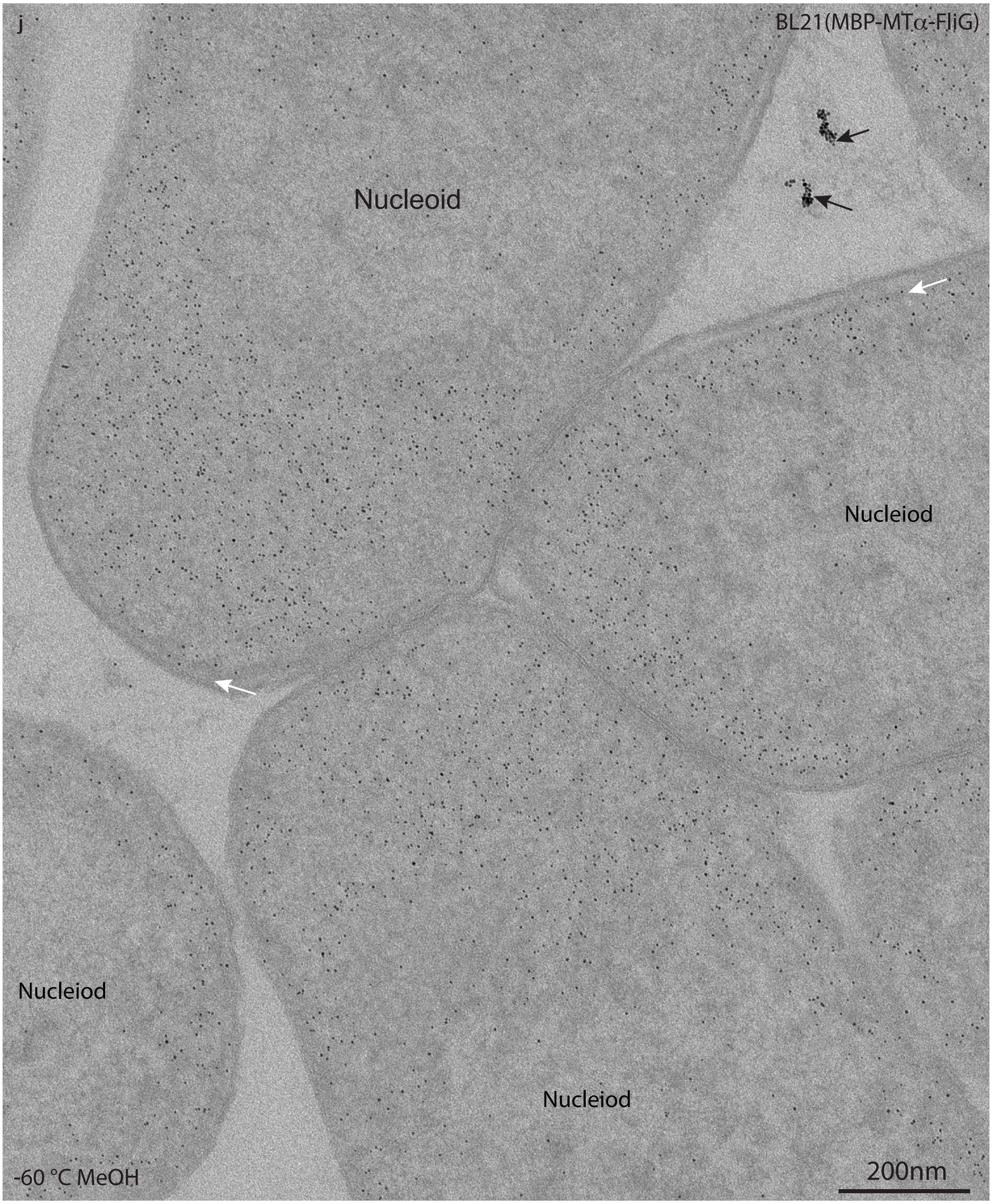


**Supplementary Figure 13E AuNPs specific distributions in the *E. coli* cells overexpressing MBP-MTα-FliG.** (j) AuNPs accumulated at poles of the cells, but much less AuNPs in Nucleiod regions; Specimen prepared by -60˚C methanol freezing for 1 min, then centrifuged to remove the methanol and changed to 4˚C PBS-A for ANSM-based AuNPs synthesis with the 2-ME/D-P protocol. There is almost no AuNPs found in the periplasmic spaces (pointed by the black arrows). Outside the cells looks quite clean excepta few dark aggregates (pointed by white arrows), which might be formed by the undissolved Au(I)(SR) polymers.The overall feature of the MBP-MTα-FliG distribution is quite similar to those of MBP-MTn-FliG (refer to Supplementary Figure 13C(h)) , except the densities of the AuNPs are less in MBP-MTα-FliG expressing cells.

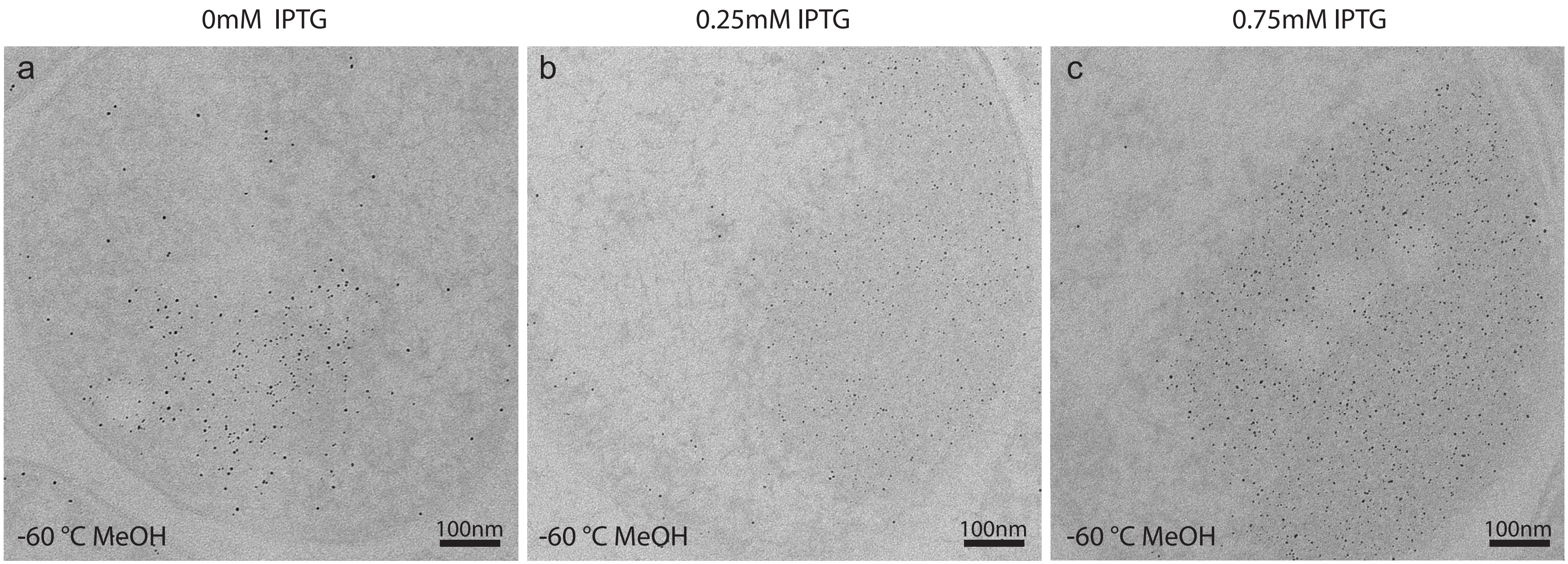


**Supplementary Figure 14 The number of AuNPs formed in cells expressing GBP-MT increasing with the induction concentration of IPTG for 4 hours.** The majority of GBP-MT fusion proteins located in the inclusion bodies of cells as indicated by the AuNPs. Compared with the induction with 0.25mM (**b**) or 0.75mM (**c**) IPTG induction, the inclusion body is smaller and contained less AuNPs without IPTG induction (**a**), the inclusion bodies for 0.25mM and 0.75mM IPTG inductions are quite similar except there are more AuNPs in 0.75mM case. Beyond the inclusion bodies, there are only a few AuNPs.

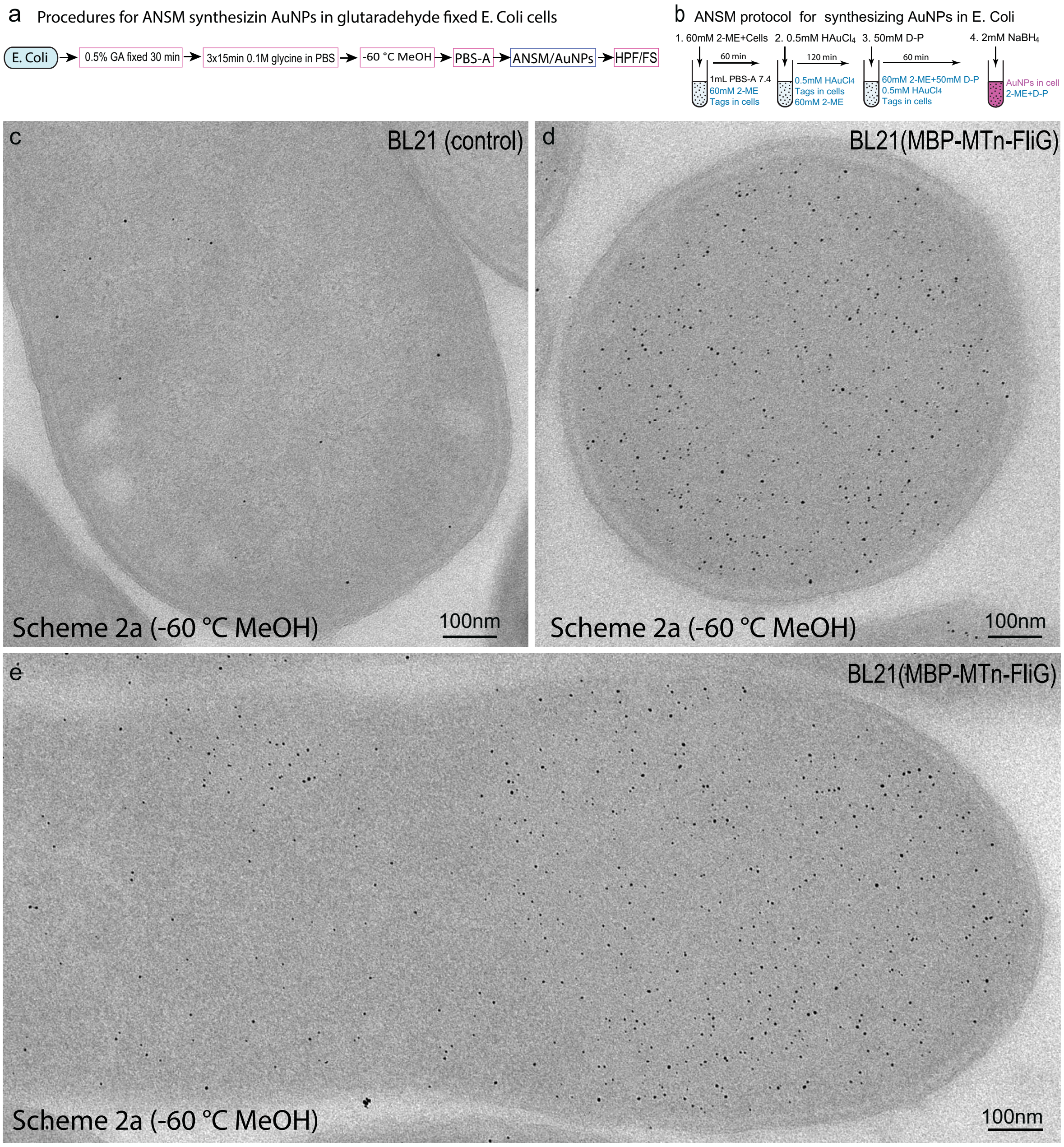


**Supplementary Figure 15 ANSM-based synthesis of AuNPs on 0.5% Glutaraldehyde fixed *E. coli* cells** (a) procedures for specimen preparation using a modified Scheme 2a by 1 min permeablization with -60 °C methanol (see **Supplementary Table 3**); (b) ANSM protocol for AuNPs synthesis with fixed *E. coli ;*(c) EM image of GA fixed wild-type *E. coli* (control), a few AuNPs firmed as background ; (d) Cross-section of *E. coli* expressing MBP- MTn-FliG which contained a lot of AuNPs; (e) Abundant AuNPs observed in the longitudinal section of an *E. coli* cell.
