## Supplementary Figure 16-20 for "Direct synthesis of EM-visible gold nanoparticles on genetically encoded tags for single-molecule visualization in cells"

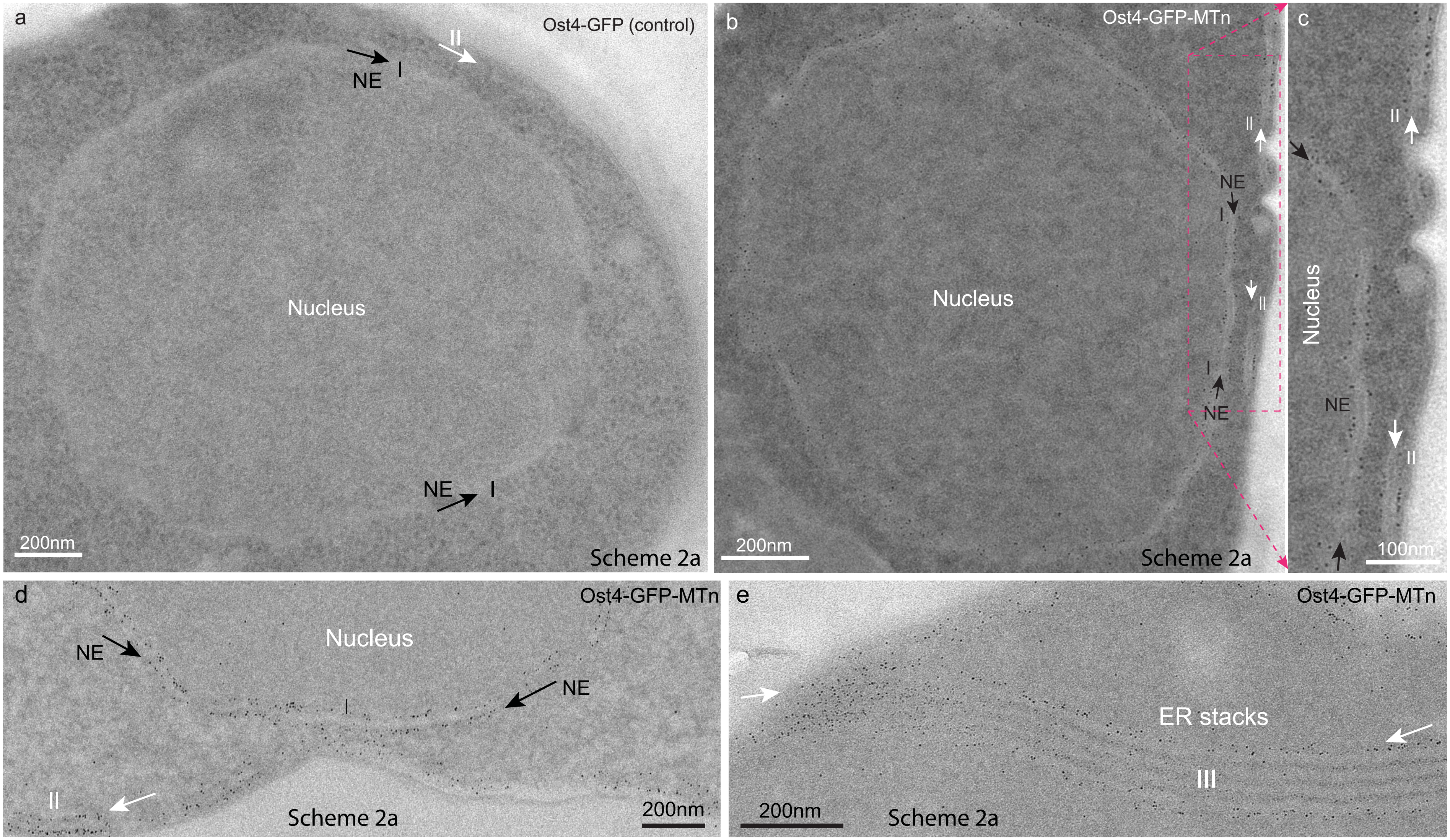


**Supplementary Figure 16 Synthesis of gold nanoparticles (AuNPs) in *S. pombe* cells processed with 3mM DTDPA oxidization, 0.5%GA fixation, 10% glycine neutralization, and 4°C methanol permeabilization (Scheme 2a, see Fig.4c and Supplementary Table 3)**
The yeast cells used for AuNPs synthesis were processed by following procedures: oxidized by 3 mM 3,3'-dithiodipropionic acid (DTDPA) in PBS for 30 min at 4°C, fixed with 0.5% GA for 15 min, then neutralized with 10% glycine in PBS for 30 min, permeabilized with 4°C cold methanol for 1-2 min. (**a**) No AuNPs formed in the Ost4-GFP expressing *S. pombe* cell (control); (**b-e**) a lot of 3-5nm AuNPs formed on the outer membrane surfaces of the nuclear envelop (NE, the black arrows pointed region I) or ER stack (III) of the Ost4-GFP-MTn expressed cells, only a few AuNPs not attached to the membranes. (**c**) An enlarged region marked with dash lines in (b) showing the AuNPs attached on the membranes. (**d**) AuNPs formed along the NE or ER outer membranes. (**e**) a lot of AuNPs formed along the ER stacks, the sizes of the AuNPs one the outer stacks are bigger than those on inner stacks due to the peneration barriers.

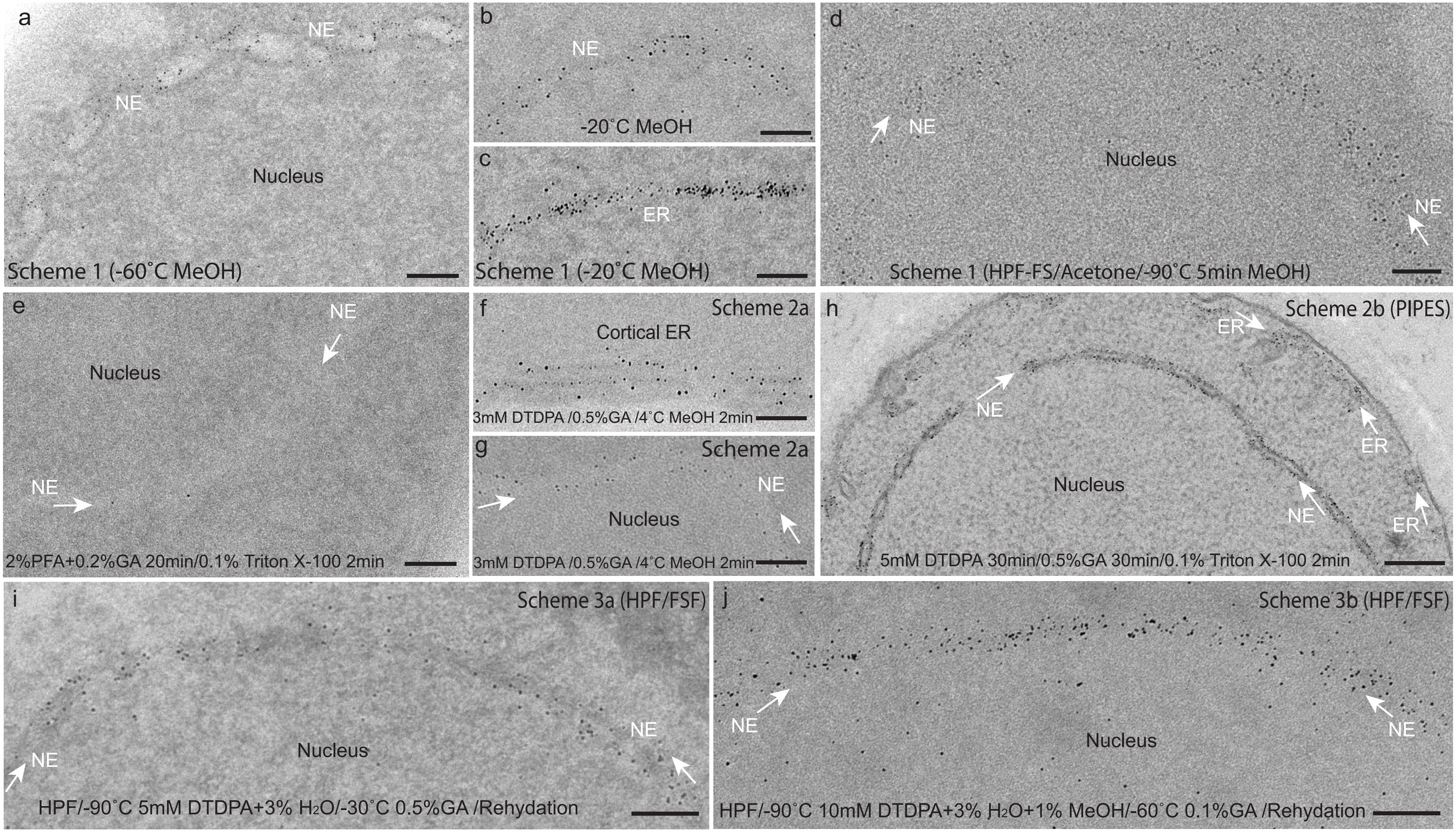


**Supplementary Figure 17A Optimization of specimen preparation protocols for visualizing the MTn tags and fine structures in *S. pombe* cells expressing Os4-GFP-MTn: thiol oxidization is essential for protecting the tags for AuNP synthesis.**

(**a-c**) Fixed and permeablized with -60˚C methanol or -20˚C methanol for 2 min; (**d**) Fixed by HPF, dehydrated in acetone at -90˚C 12h, treated by -90˚C methanol 5 min, then sucked the methanol away and placed on ice; (**e**) Fixed with 2% PFA+0.2% GA for 20 min, then permeabilized by Triton X-100 for 2min; (**f-g**) Oxidized by 3mM DTDPA in PBS for 30 min, fixed with 0.5% GA for 30 min, permeabilized with 4˚C methanol for 2 min; (**h**) Oxidized with 5 mM DTDPA in PIPES 30 min, then fixed with 0.5% GA 30 min in 0.1M PIPES and permeabilized with 0.1% Triton X-100 2 min (the procedures described as optimzed protocol for yeast cells in online methods) ; (**i**) Fixed by HPF, oxidized with 5 mM DTDPA with 3% H2O in acetone for 24 h, warm to -30˚C and fixed with 0.5% GA 5 h, then rehydated by adding 30% PBS-A at -35˚C and gradually increasing PBS-A to 100% at 4˚C; (**j**) Processed similar to (i) but with 10mM DTDPA and added additional 1% methanol, and fixed with 0.1% GA at 60˚C for 12 h, then gradually rehydrated. Those cells fixed with aldehyde fixatives were neutralized by 10% glycine in PBS for 60 min, and all the above treated cells were used for ANSM/AuNPs synthesis with the standard 60mM 2-ME+50 mM D-P protocol. The densities of AuNPs on the NE membranes in the non-aldehyde fixative treated cells (a-d) are relative higher than those of aldehyde fixatives fixed cells (e-j). Almost no AuNPs formed in cell directly fixed by aldehyde fixative without an oxidizing protection of the thiol groups on the MTn (e). The densities of AuNPs synthesized in those cells treated with oxdizing reagent, DTDPA, increaing with with the DTDPA concentrations, e.g., 3 mM, 5 mM or 10 mM, prior to the aldehyde fixation(f-j). The DTDPA oxidization is essential for protecting the tags for AuNP synthesis.

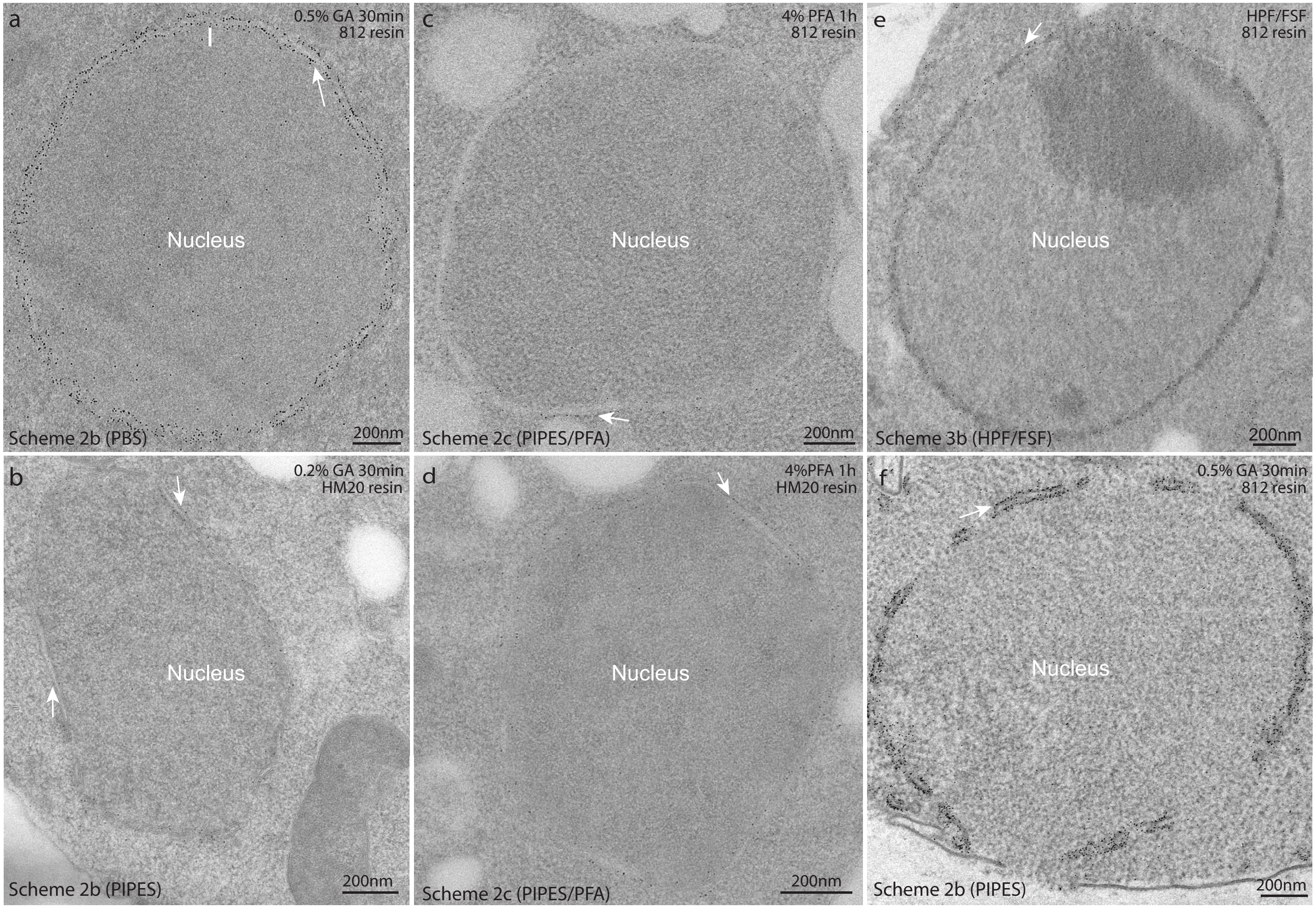


**Supplementary Figure 17B Optimization of specimen preparation protocols for visualizing the MTn tags and fine structures in *S. pombe* cells expressing Ost4-GFP-MTn: fixatives, fixation strength, buffers, resin types.**

(**a**) specimen was oxidized with 3 mM DTDPA 30 min in PBS at 4˚C, then was fixed with 0.5% GA for 30 min, was permeabilized with Triton X-100 for 2 min, was embedded in SPI-Pon 812 resin; (**b**) specimen was prepared similar to (a), except using PIPES buffer, 0.2% GA and HM20 resin; (**c**) specimen preparation similar to (a), except using 4% PFA in PIPES buffer for 1 h fixation; (**d**) specimen prepared was the same as for (c), except embedded in HM20 resin; (**e**) specimen prepared with Scheme 3b: fixed with HPF, oxidized with 3 mM DTDPA and fixed with 0.1% GA in acetone during freeze-substitution processes, then rehydrated for AuNPs synthesis; (**f**) specimen was prepared similar to (b), except using 0.5% GA and SPI-Pon 812 resin. All the above specimens were subjected to the standard protocol for ANSM-based AuNPs synthesis (60 mM 2-ME, 0.5 mM $HAuCl_4$ ,50 mM D-P). Both the GA and PFA can be used as fixatives and suitable for AuNP synthesis, it seems that PFA gave better fixation but less AuNPs (a-d). The fixation using PIPES buffer gave better membrane preservation by comparing (a) with (b,c,d,f ), indicated by white arrows pointed regions.Compared with Spomn 812 resin, the HM20 resin gaves better membrane contrast as indiated by (a) and (b), (c) and (d), the white pointed regions in (b) and (d) clearly demonstated the AuNPs are attached to the surfaces of the membranes of nuclear envelope. HPF/FSF protocol preserved much smooth curvature of nuclear envelope as indiated by (e) with the others, however, the rehydration processes disordered the fine structure of membrane and resulted in a diffused distrubution of AuNPs.

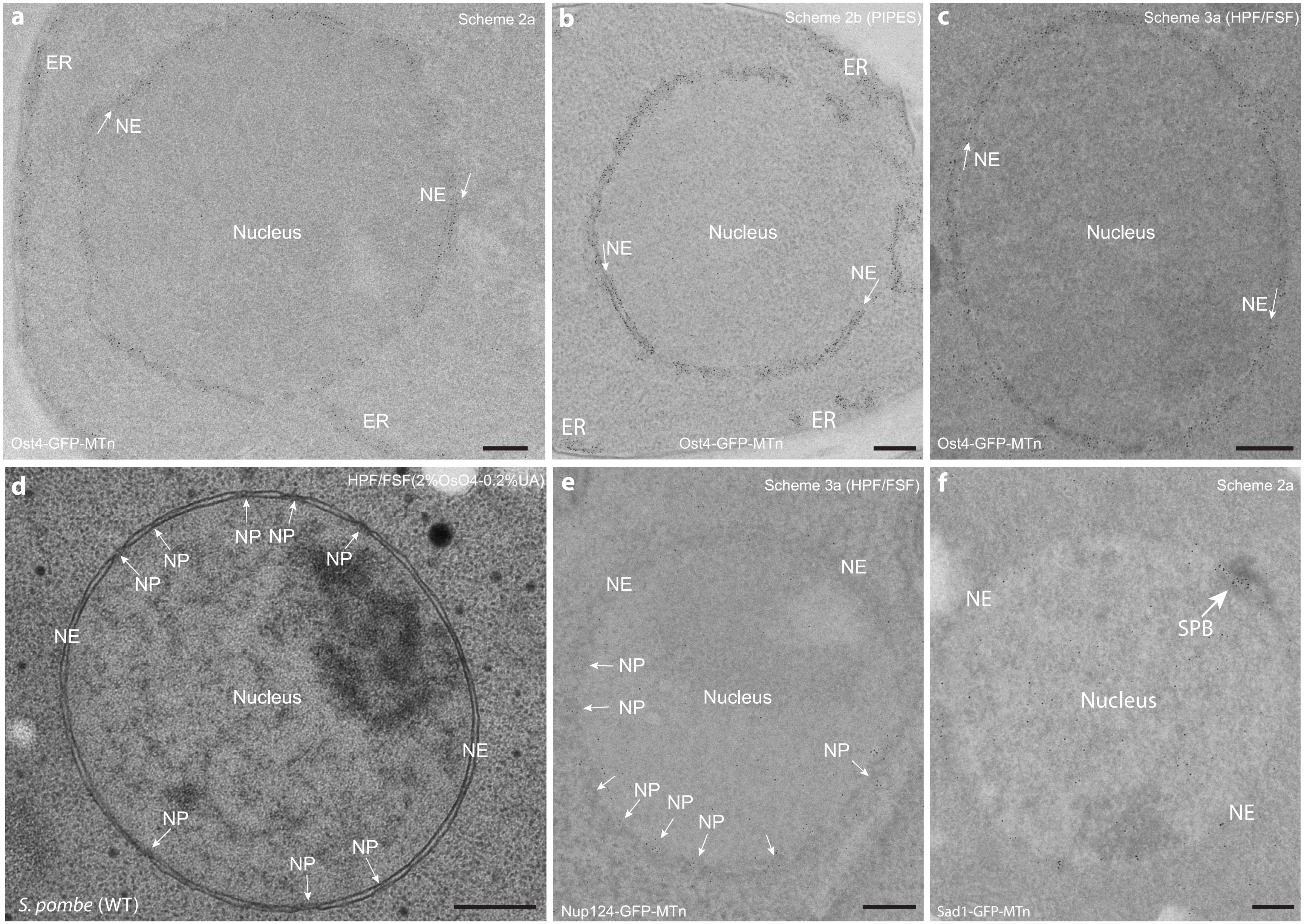


**Supplementary Figure 18 Localization of GFP-MTn tagged Ost4, Nup124, and Sad1 fusion proteins in *S. pombe* cells by synthesis of AuNPs on MTn tags**
(**a**) Ost4-GFP-MTn expressing *S. pombe* cell treated by Scheme 2a (**Fig.4c & Supplementary Table 3**): 3mM DTDPA oxidization for 30 min in PBS buffer , 0.5% GA fixaton for 30 min, 4˚C methanol permeabilization. (**b**) Ost4-GFP-MTn expressing cell treated with Scheme 2b (PIPES) : 5 mM DTDPA oxidization for 30 min in 0.1M pH 6.8 PIPES buffer with 1mM $MgCl_2$ and $CaCl_2$, and 0.1M sorbitol, 0.5% GA fixation for 30 min, 10% glycine neutralization overnight, 0.2 mg/ml zymolyase-20T cell wall digesting at RT for 30min, 0.1% Triton X-100 permeabilization for 2 min. (**c**) Ost4-GFP-MTn expressing cell treated by Scheme 3a : HPF, 5 mM DTDPA oxidized in cold acetone containing 3% $H_2O$, 0.5% GA fixed. (**d**) Wild-type *S. pombe* cell treated by standard HPF/FSF protocol with 2% $OsO_4$+0.2% UA,showing the nuclear pores (NPs). (**e**) Nup124-GFP-MTn expressing cells treated by the same protocol as used for (c), AuNPs formed at NPs. (**f**) A group of AuNPs formed at the spindle pole body (SPB) in a Sad1-GFP-MTn expressing cell treated with Scheme 2a. Most of the AuNPs were localized to the expected organelles: Ost4-GFP-MTn on membranses of nuclear envelope (NE) or ER, and Nup124-GFP-MTn on the nuclear pores (NPs) and Sad1-GFP-MTn on SPB. The scale bars correspond to 200nm.

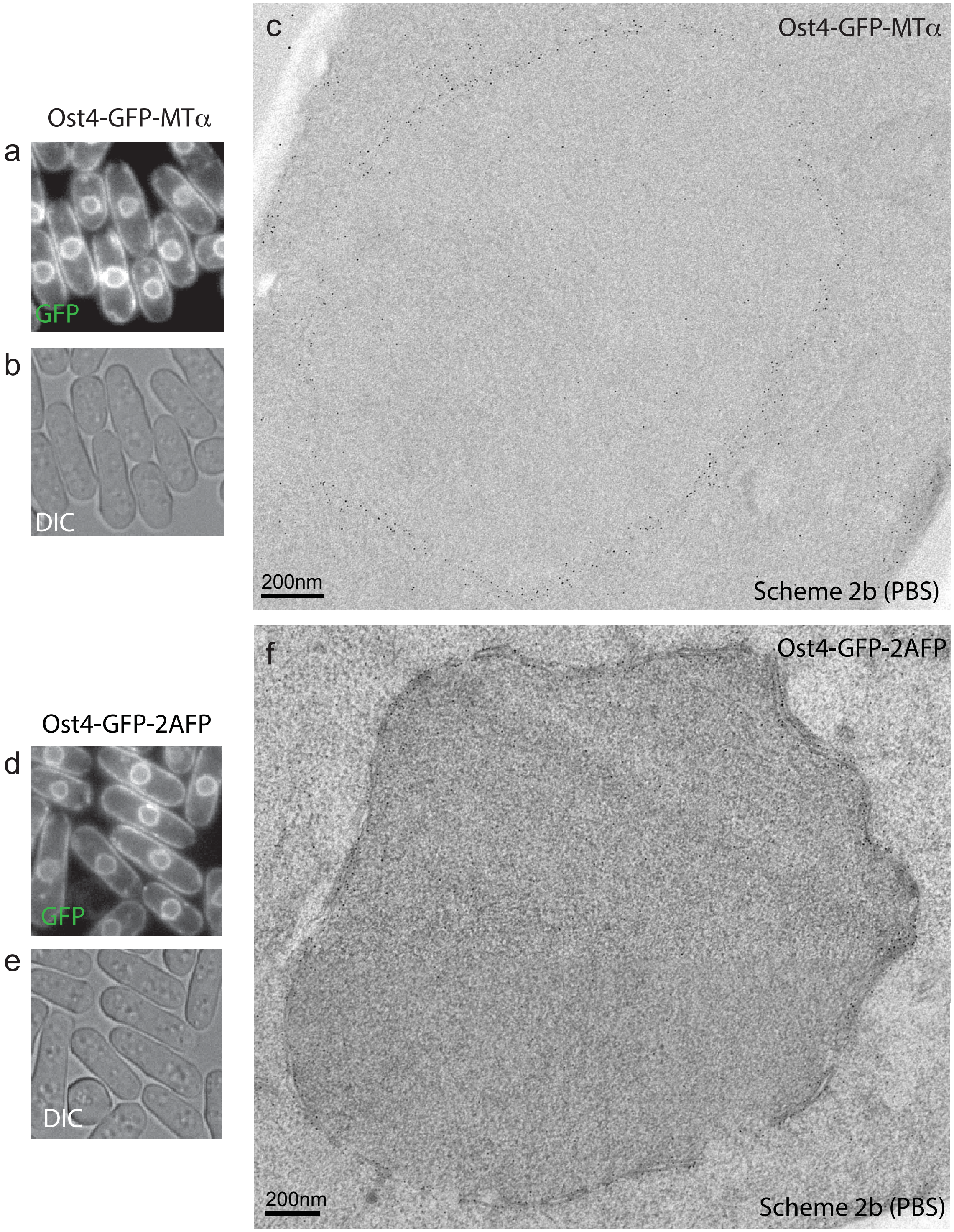


**Supplementary Figure 19A Expression patterns of *S. Pombe* strains expressing Ost4-GFP-MTα or Ost4-GFP-2AFP.** (**a**) Expression pattern of Ost4-GFP-MTα; (**b**) Differential interference contrast (DIC) image of the region corresponding to (a); (**c**) EM image of a typical nuclear envelop (NE) of an Ost4-GFP-MTα expressed cell showing the AuNPs distributed alonge the nuclear envleop as ring-shape; (**d**) Expression pattern of Ost4-GFP-2AFP (e) Differential interference contrast (DIC) image of the region corresponding to (d); (f) EM image of a typical nuclear envelop (NE) of an Ost4-GFP-2AFP expressed cell, showing the AuNPs attaced on the outer sufaces of NE membranes; The EM specimens were prepared with Scheme 2b (PBS) (see **Supplementary Table 3**).

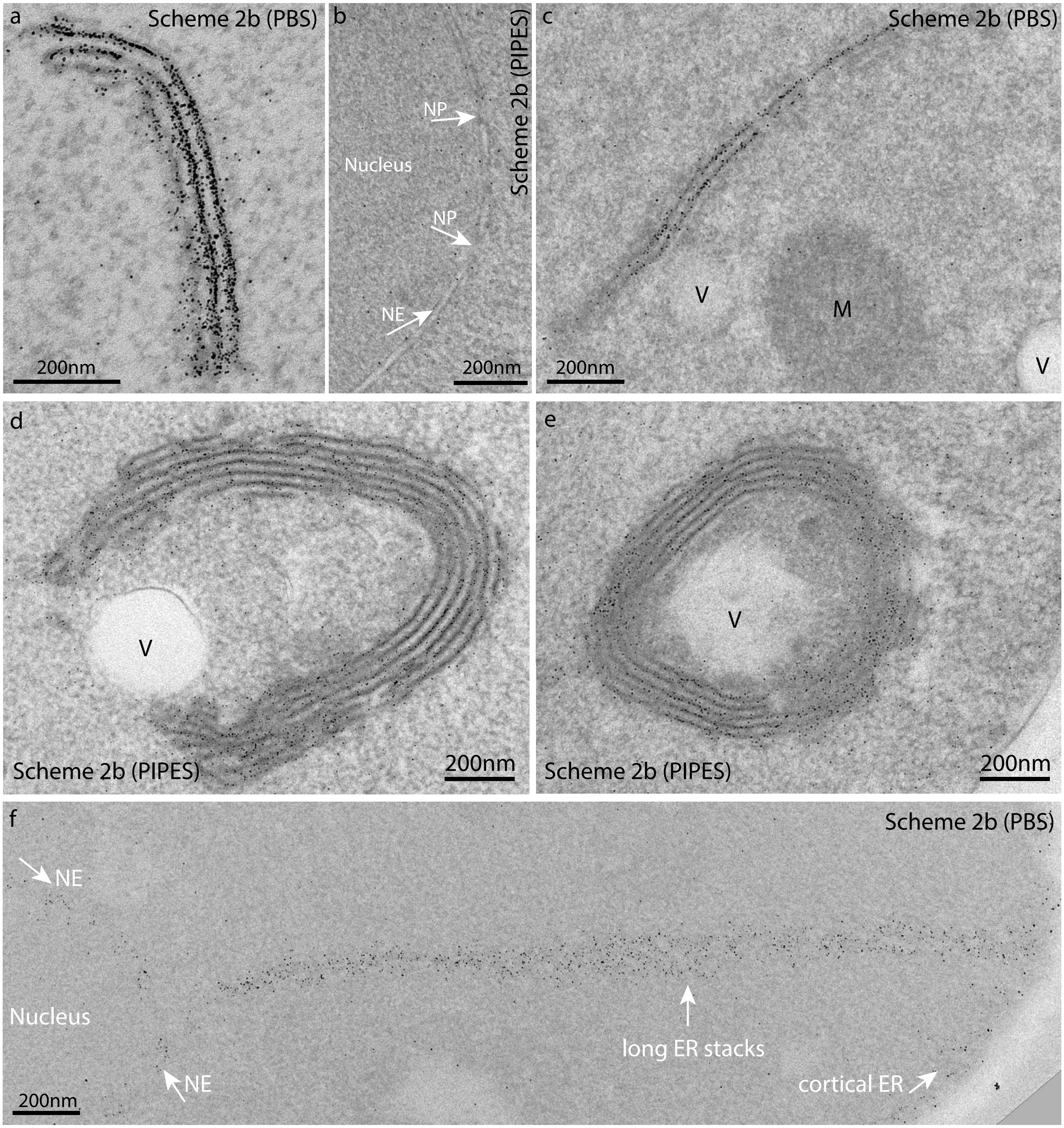


**Supplementary Figure 19B Expression patterns of *S. pombe* strains that expressed Ost4-GFP-MTα**
(a,c-f) EM image of a typical ER stack(s) demonstrated by AuNPs synthesized on Ost4-GFP-MTα expressing cells, almost no AuNPs in the mitochondrion (M) and the vaculo (V); The densities of AuNPs on nuclear envelope (NE) (b,f) were relative lower than those on the ER stacks (a,c-f). The EM specimens for (a,c,f) were prepared with Scheme 2b (PBS) (see Fig.4c & Supplementary Table 3): 3 mM DTDPA oxidization for 30 min PBS (pH 7.4) buffer, 0.5% GA fixation (Note: the vial containing 25% GA was not freshly opened, the actual diluted concentration might lower than 0.5%) for 30 min, 10% glycine neutralization overnight, 0.2 mg/ml zymolyase-20T cell wall digesting at RT for 30min, 0.1% Triton X-100 permeabilization for 2 min; the processed cells were used for standard ANSM/AuNPs synthesis; The Specmens for (b,d,e) were prepared with Scheme 2b (PIPES) similar to (a), except using 5 mM DTDPA in PIPES buffer (see **Supplementary Table 3**).

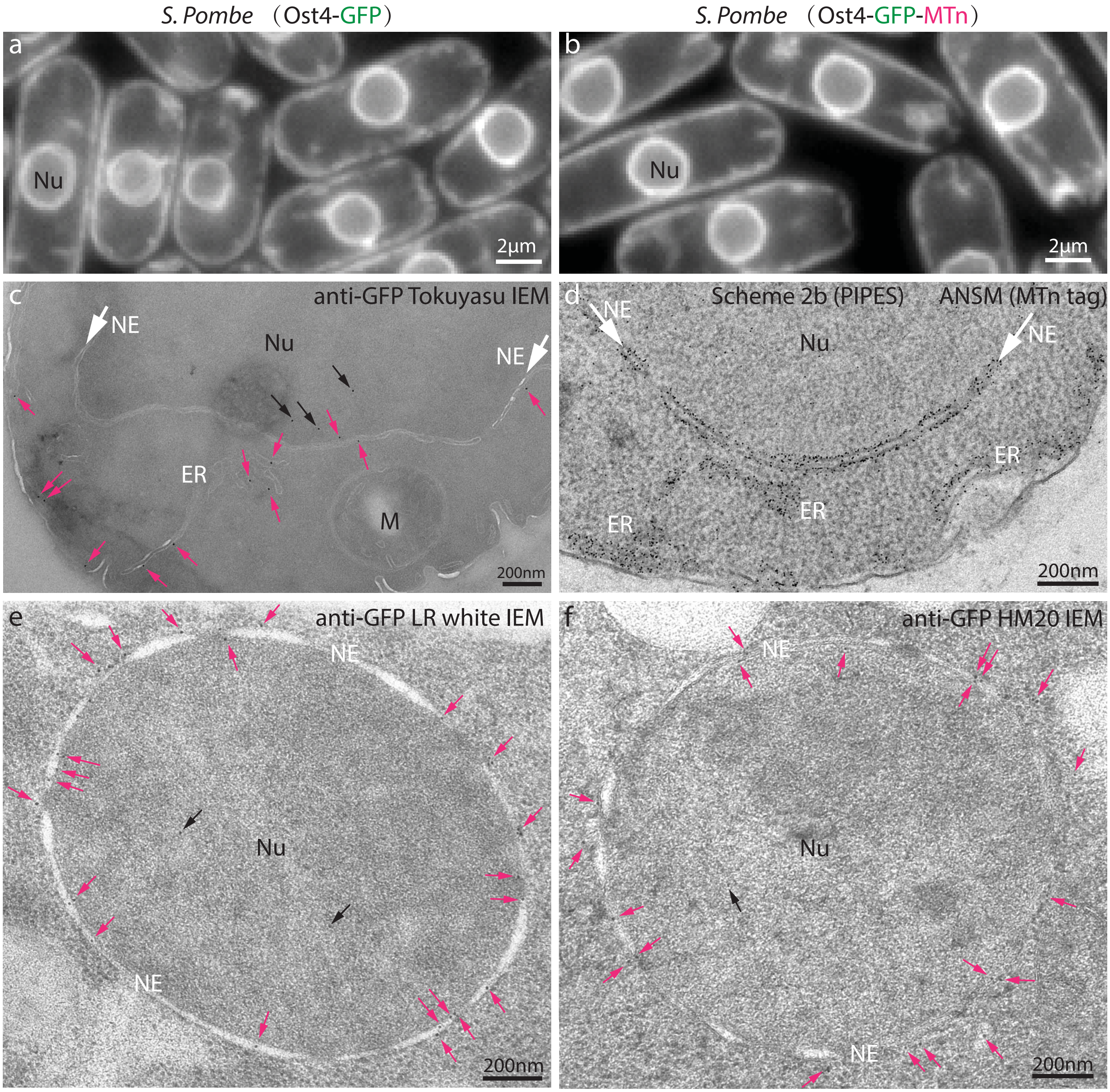


**Supplementary Figure 20 A comparison of the performance of metallothionein tag labeling (ANSM), immunogold staining with Tokuyasu technique and conventional immuno-EM methods**

(**a-b**) Typical expression patterns of the ER membrane-locating Ost4 demonstrated by the GFP reporters in the two *S. prombe* strains (Ost4-GFP and Ost4-GFP-MTn), indicated that the tagged Ost4 proteins were mainly localised on the ER membranes including the nuclear envelope (NE), and the two strains shared similar patterns. (**c**) Immunogold staining of Ost4-GFP by anti-GFP (goat) antibody (Cat. No.600-101-215, Rockland) and protein A-10 nm gold particles on cryosection of yeast cell expressing Ost4-GFP prepared with Tokuyasu technique (cells fixed with 2% PFA + 0.2% GA with 0.75M sorbitol in PHEM buffer, PH 6.9 for 1 h at 30˚C). (**d**) Direct synthesis of AuNPs on Ost4-GFP-MTn expressed in *S. pombe* cell by using the Scheme 2b (PIPES). (**e**) Immunogold staining of Ost4-GFP by anti-GFP (rabbit) antibody (Cat. No. ab6556, Abcam) and anti-rabbit IgG antibody-10nm gold particles on LR white embedded plastic section of yeast cell expressing Ost4-GFP (cells were prepared with HPF, FSF with 0.1% UA+5% $H_2O$ in acetone). (f) Immunogold staining of Ost4-GFP-MTn of yeast cells expressing Ost4-GFP-MTn (same conditions for fixation and immunogold staining as used for (e), except embedded in HM20 resin). Notably, the labeling density of ANSM AuNPs synthesis on Ost4-GFP-MTn (d) is ~10-20x higher than those in immunogold staining (c,e,f).
