## Supplementary Figure 21-24 for "Direct synthesis of EM-visible gold nanoparticles on genetically encoded tags for single-molecule visualization in cells"

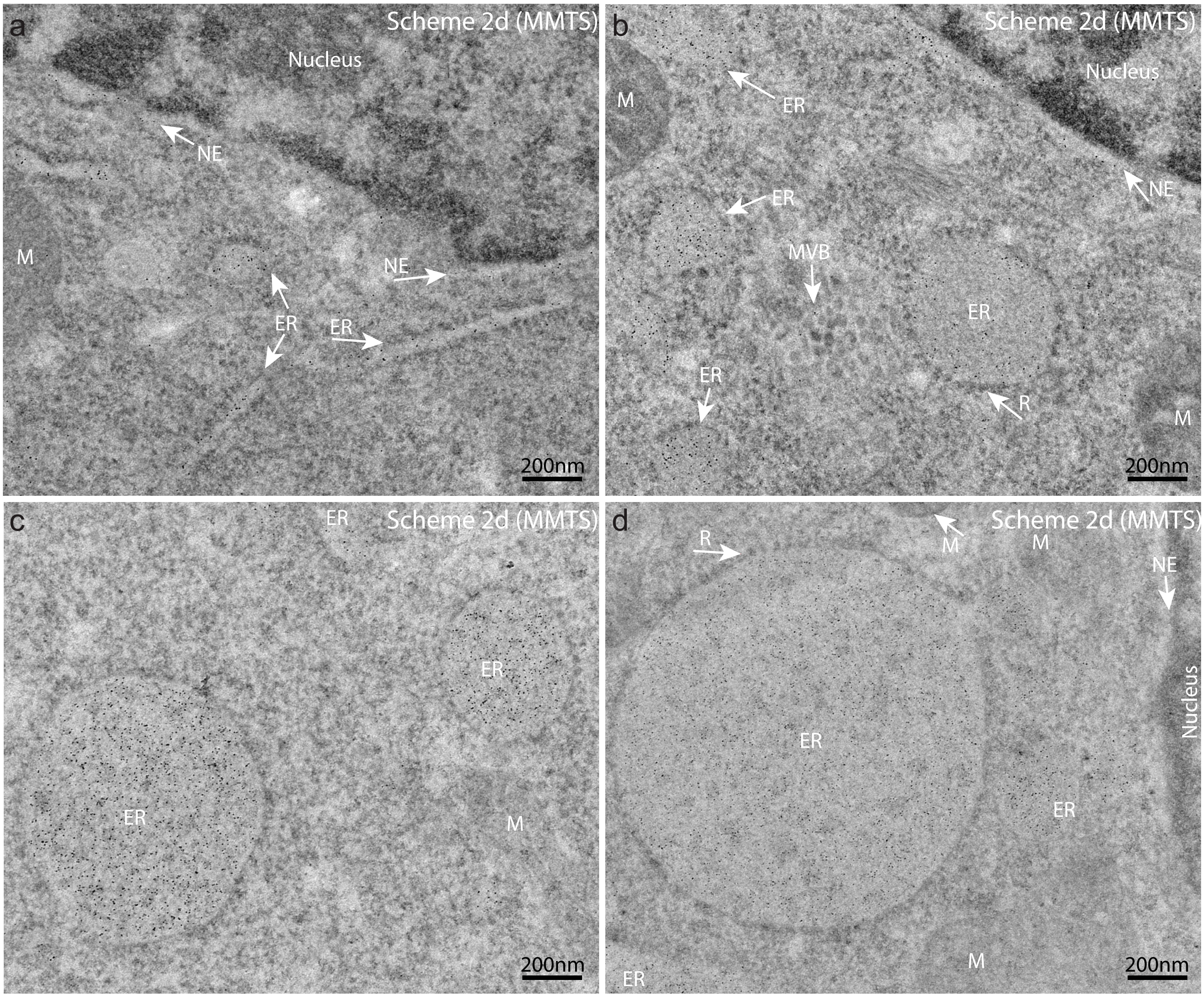


**Supplementary Figure 21A Distributions of AuNPs synthesized on GFP-MTn-KDEL that expressed in HeLa cells.** (**a**) AuNPs mainly distributed inside the lumen of long ER cisternae ,some in the expanded regions of ERs and nuclear envelope (NE), occassionally a few AuNPs scattered in cytosol or nucleus; (**b**) Majority of AuNPs were specifically located in expanded ER regions and some in NE, some inside the multivesicular body(MVB), no AuNPs in mitochondria (M);(**c-d**) A great many AuNPs accumulated in some huge expanded round ER blobs, some ribosomes (R) were attached to the outter membranes of the blobs,such huge ER blobs might be resulted from the overexpression of GFP-MTn-KDEL molecules (even in the fluorescent image can see the bright and huge blobs, refer to Fig.5a), in contrast, seldomly seen any AuNPs cytosols, mitochondria and nucleus. The specimens were prepared with **Scheme 2d (MMTS)** (See **Fig.4c** & **Supplementary Table 3**): the cells cultured on sapphire discs were oxidized by 25mM MMTS for 5 min at room temperature, then fixed with 0.1% GA (kept the MMTS during the fixation) for 5 min at RT, neutralized with 10% glycine overnight on ice, permeabilized with 0.1% Triton X-100 for 2 min,change to PBS-A buffer for ANSM/AuNPs synthesis (Refer to “Protocol for mammalian cellsattached to sapphire discs” in the online methods for details).

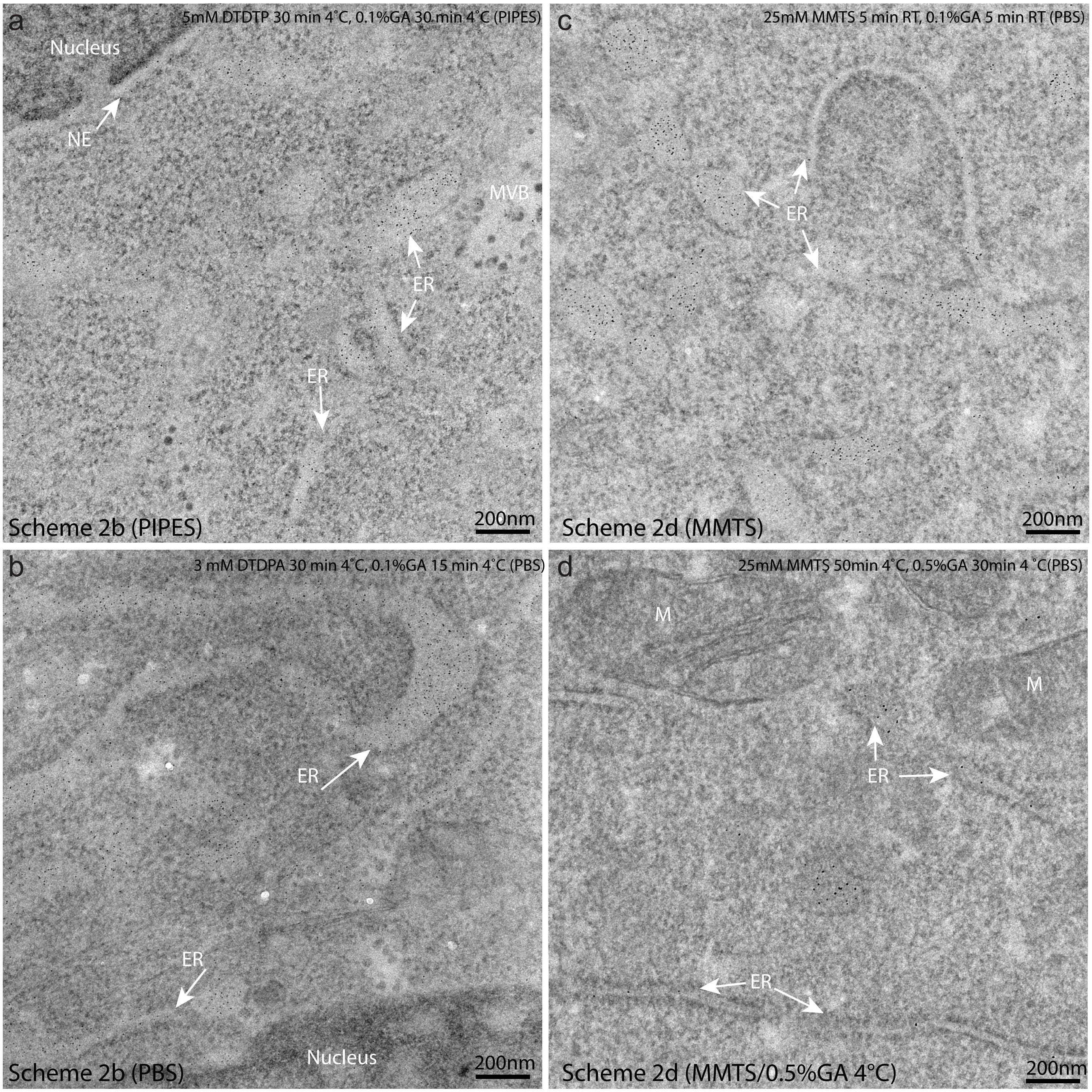


**Supplementary Figure 21B Evaluation of the performance of oxidants and the strength of glutaraldehyde fixation for preserving the activities of the MTn tags and fine structures in HeLa cells.**

(**a-b**) Specimens were prepared with Scheme 2 (see **Supplementary Table 3**): 3-5 mM DTDPA oxidizing for 30 min at 4˚C, followed with 15-30 min of 0.1% GA fixation at 4˚C and standard ANSM/AuNP synthesis, it demonstrated that 3-5mM DTDPA preserved the activity of MTn tags well as indicated by a lot of AuNPs specific localized in ER lumens, the less fixation duration achieved higher AuNPs density but got worse fine structures preservation (b). (**c-d**) Specimen were prepared with 25mM MMTS oxidization at either room temperature (RT) for 5 min or 4˚C for 50 min, then fixed with 0.1% GA for 5 min at RT or 0.5% GA for 30 min, followed with standard ANSM/AuNP synthesis, apparently 25mM MMTS oxidization at RT was efficient for protecting the activities of MTn tags but the fine structures preservation was compromised for weak strength of fixation (0.1% GA for 5 min at RT) (c); while the high strength of fxation 0.5% GA for 30 min preserved much better fine structure as indicated by the better membrane structures in ER and mitochondrion (M) but at the cost of lost of activities of MTn tags as indicated by lower density of AuNPs (d). In conclusion, the optimal oxidization could be achieved with either 3-5 mM DTDPA for 30 min at 4˚C or 25mM MMTS for 5 min at RT, the compromised fixation condition would be 0.1% GA for 30 min at 4˚C or for 5 min at RT. It should be pointed out that further optimization of the fixation conditions to achieve better ultrastructure preservation and reasonable labeling density is deserved to be explored.

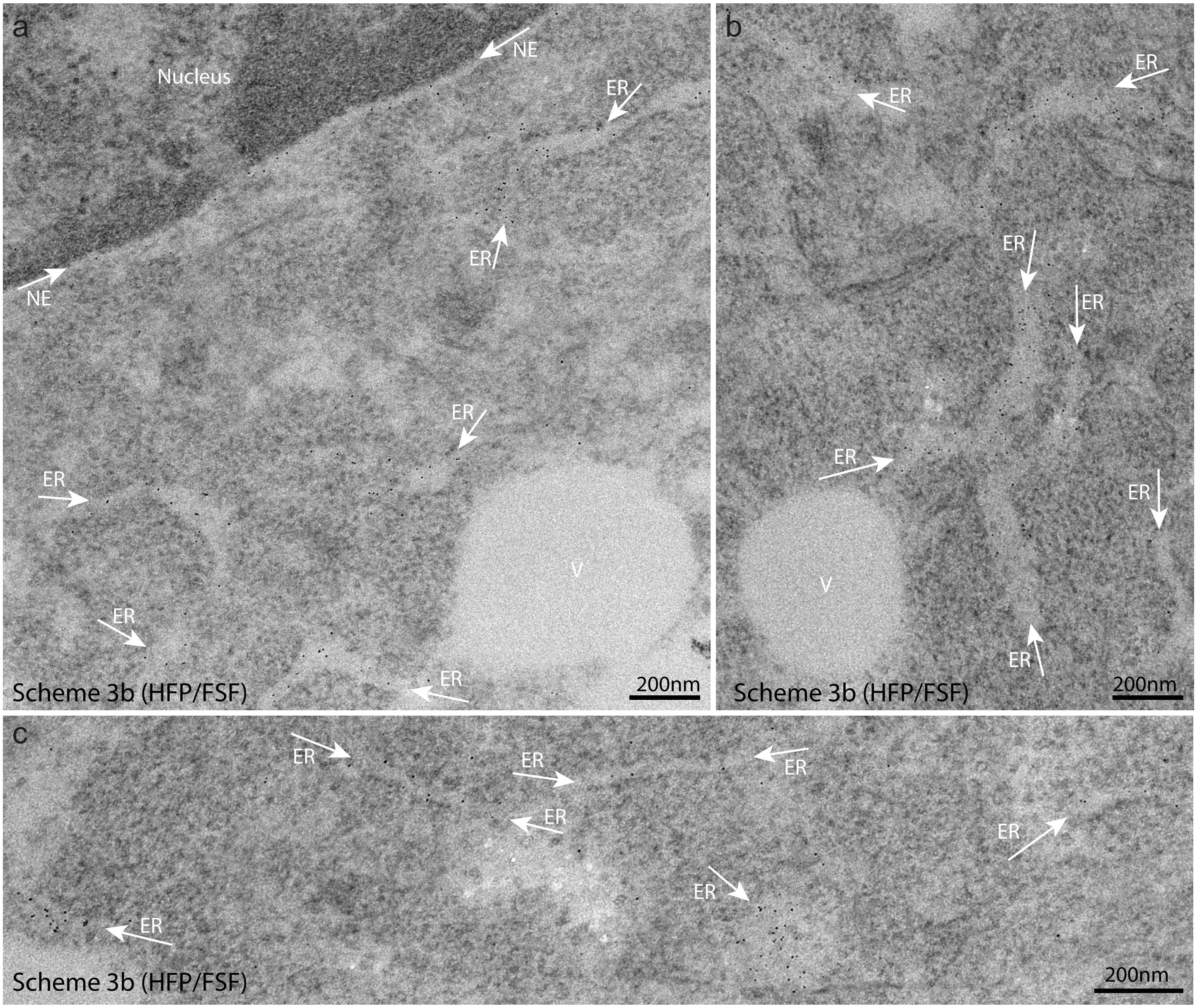


**Supplementary Figure 21C Distributions of AuNPs synthesized on GFP-MTn-KDEL that expressed in HeLa cells fixed with high pressure freezing**

(**a-c**) HeLa cells rapid fixed with high pressure freezing (HPF) and processed with a freeze-substitution based oxdization,GA fixation , rehydration, then used for standard AuNP synthesis (**Scheme 3b** as shown **Supplementary Table 3**) : the cells cultured on sapphire disc was fixed with HPF, then oxidized with 10mM DTDPA in 5% $H_2O$+1% MeOH in acetone at -90˚C for 12 h, then warm up to -20˚C within 8 h, after 2 h cooled back to -30˚C and add 0.1% GA for 5 h fixation, gradually rehydrated for ANSM/AuNPs synthesis (described in online methods section: "Procedures for FSF-based AuNP synthesis in mammalian cells"). Majority of AuNPs were localized to the lumens of long ER cisternae, nuclear envelope (NE), and expanded ER (all ponited by white arrows); however, the the membrane structures of ER, NE, vaculoes (V) were not well-preserved due to the rehydration processes (further refinement of the protocol might help to improve the preservation of ultrastructures).

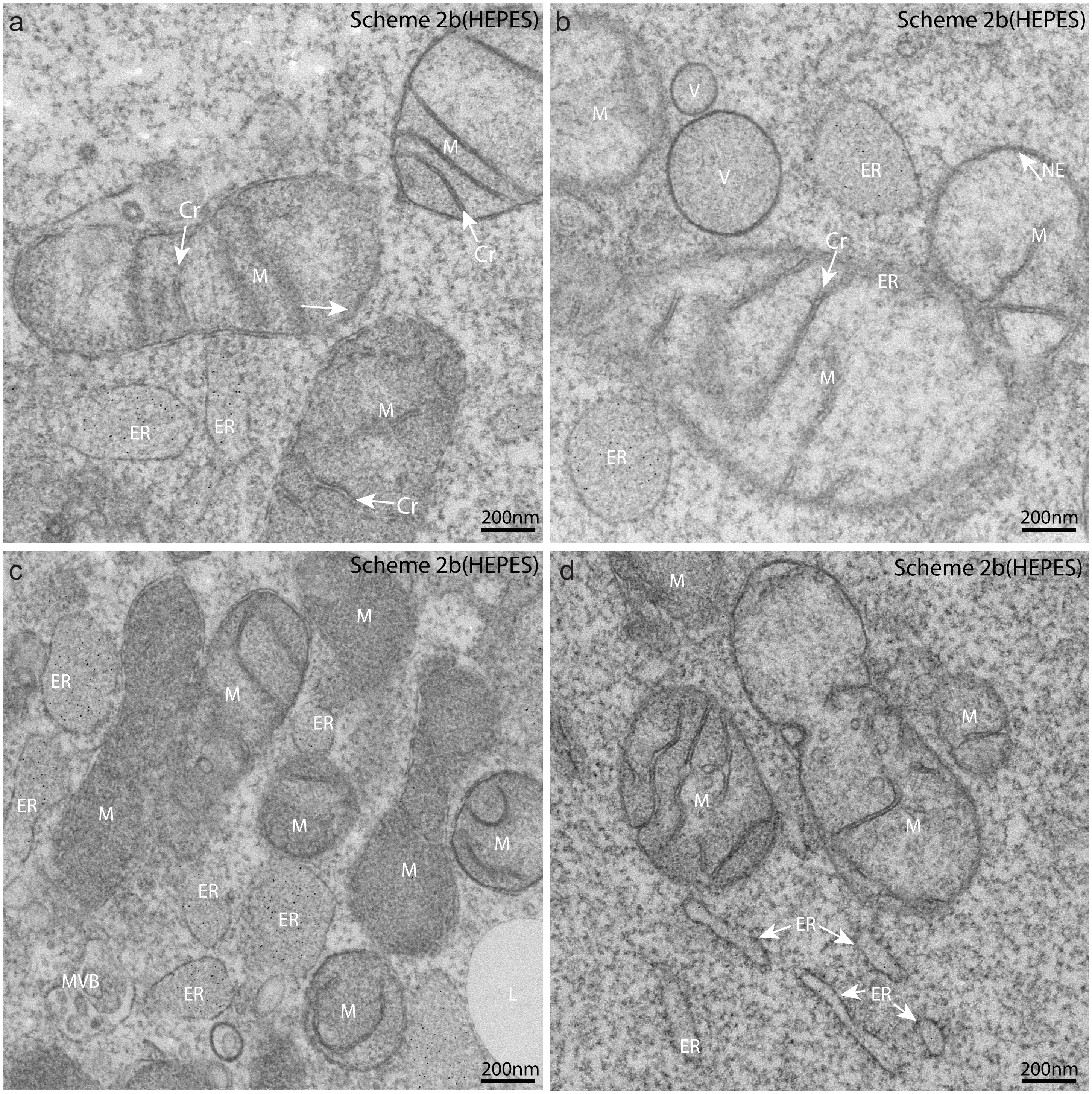


**Supplementary Figure 21D HeLa cells expressing GFP-MTn-KDEL oxidized and fixed in HEPES buffer preserved membrane structures**

AuNPs mainly distributed in the expanded regions of ERs (a-c), only a few AuNPs in tubular ER (d), almost no AuNPs could be detected in cytosol, mitochondria (M), vesicles (V) or lipid droplet (L); the membranes of cristae (Cr) , vesicles (V), and lipid droplet (L) (c) were well preserved. All the specmens (a-d) were prepared with Scheme 2b (HEPES) (see **Supplementary Table 3.**): oxidized with 5 mM DTDPA in HEPES buffer (0.2M HEPES buffer with 1mM $CaCl_2$ and 1 mM $MgCl_2$, pH 7.4) for 30 min at 4 °C; fixed with 0.05% GA for 5 min at 4 °C; 50mM glycine in HEPES buffer 2-9 h neutralization; 0.1 % Triton X-100 in HEPES buffer for 2 min; washed with 3x5 min PBS;standard AuNP synthesis for mammalian cells (60mM 2-ME 1 h at RT, 0.7mM $HAuCl_4$ and 40mM incubation 2 h at 4 °C, reduced with 1mM $NaBH_4$).Refer to “Protocol for mammalian cells attached to sapphire discs” in the online methods for details.

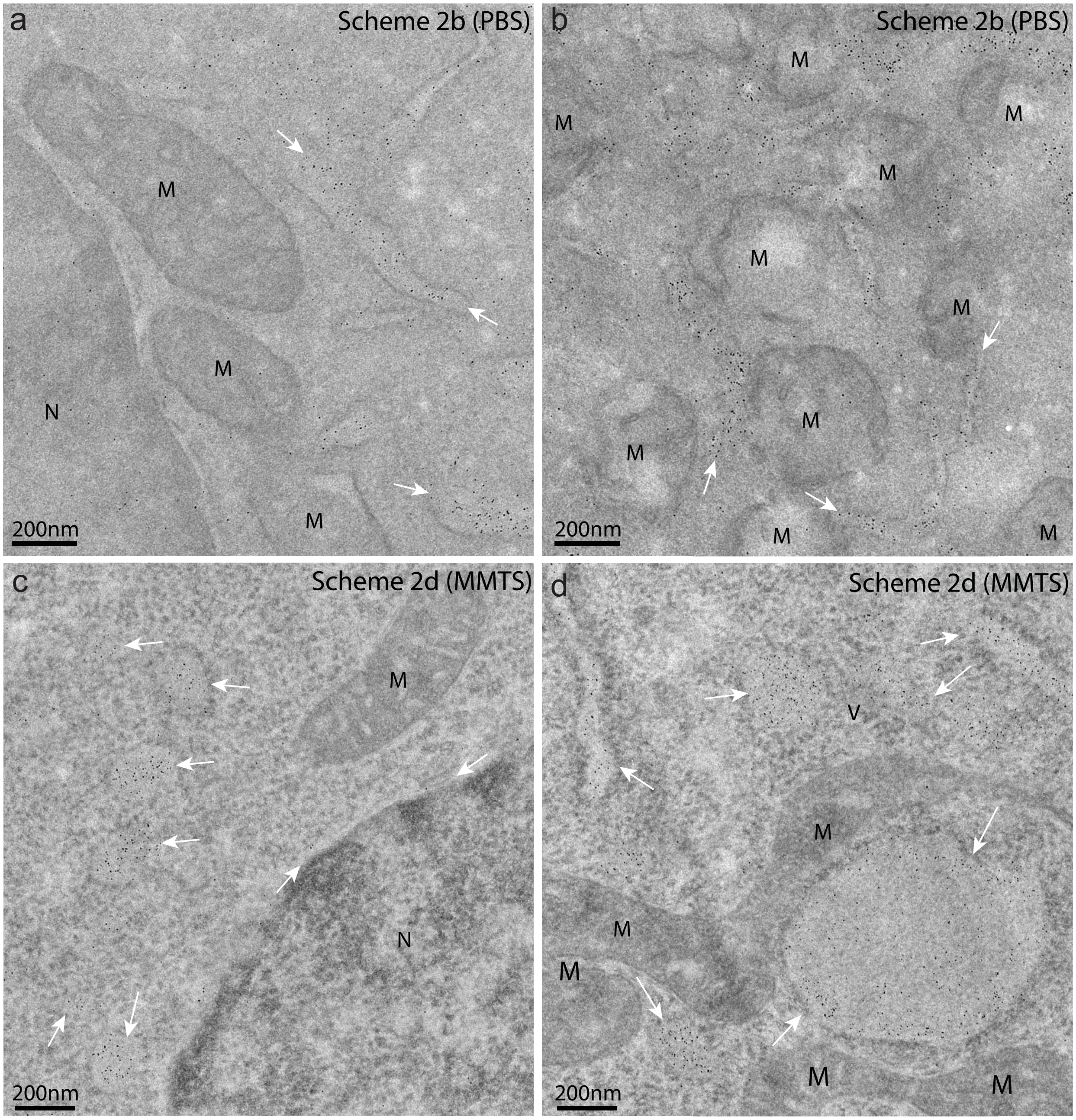


**Supplementary Figure 21E Distributions of AuNPs synthesized on GFP-MTn-KDEL that expressed in HeLa cells: the high nonspecific background signals of AuNPs might be caused by the centrifugation**

(**a-b**) AuNPs mainly distributed inside the lumen of tubular ERs and some in the expanded regions of ERs (ponited by white arrows), but apparently seen some nonspecific background signals of AuNPs in Nucleus (N), mitochondrion (M) and cytosol; the specimens were prepared with a modified Scheme 2b (PBS) (**Supplementary Table 3**): the cells cultured on cultured on 35mm dishes and digested with typsin-EDTA for cell pellets collection, the cell pellets were further processed by the procedures described in "Protocol for AuNP synthesis in mammalian cells in suspension" in online methods.
(**c-d**) AuNPs highly specific distributed in lumens of a tubular ERs or expanded ERs (pointed by white arrows), rarely seen nospecific signals of AuNPs in cytosol, mitochondrion (M), and nucleus (N); the cells cultured on sapphire discs were prepared with Scheme 2d (MMTS) (25mM MMTS oxidization in PBS for 5 min at RT, 0.1% GA fixed for 5 min at RT), the detail procedures described in "Protocol for AuNP synthesis in mammalian cells attached to sapphire discs" in online methods. Apparently, the random background signals of AuNPs for cells in suspension (subjected several times of centrifugation) are much higher than those of the cells directly cultured on sapphire discs without centrifugation.

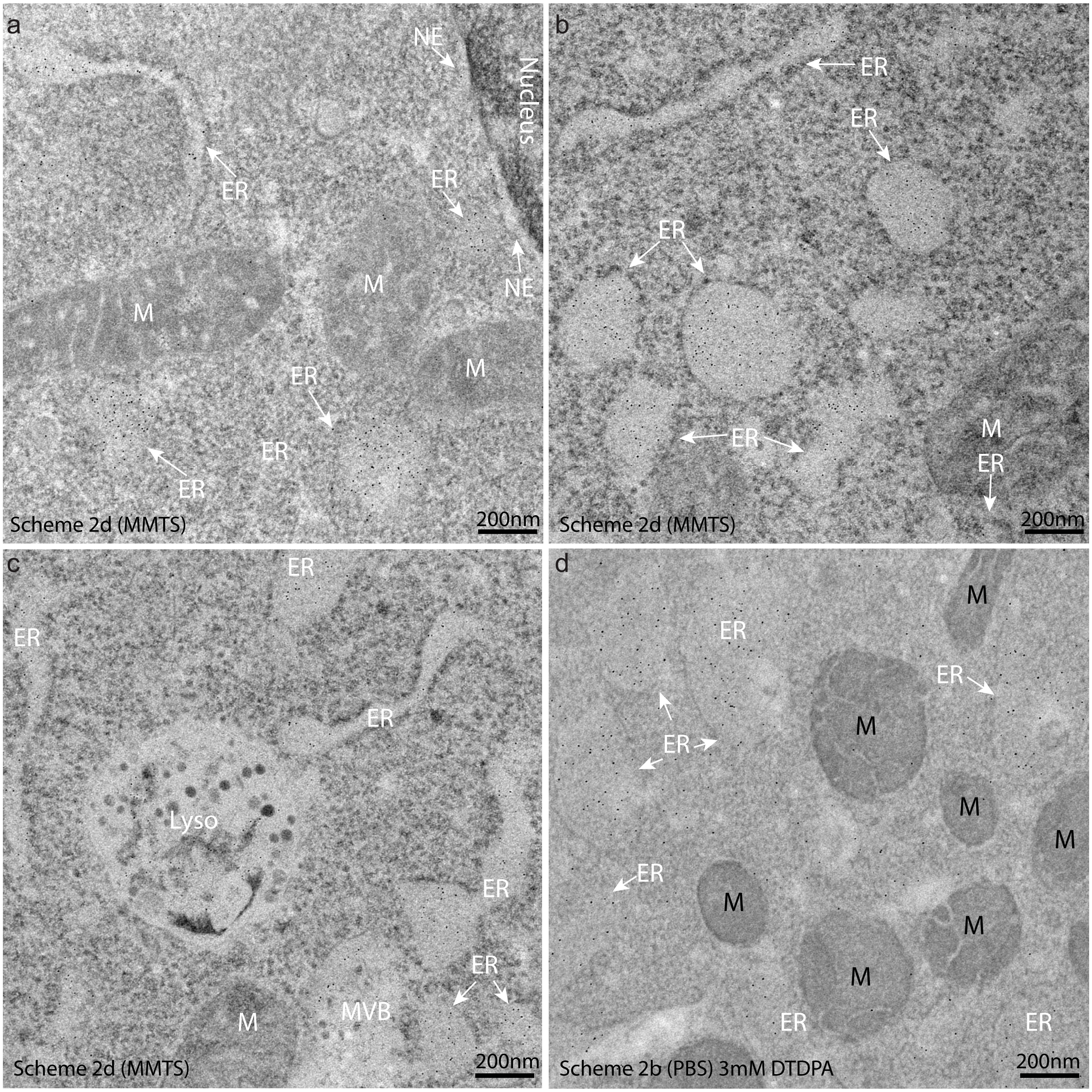


**Supplementary Figure 22 Distributions of AuNPs synthesized on GFP-2AFP-KDEL that expressed in HeLa cells.** (**a-c**) AuNPs mainly distributed inside the lumen of long cisternae of ERs and the expanded cisternae of ERs (ponited by white arrows); seldom found AuNPs in the mitochondria (M), but found a lot of AuNPs in the lysosome (Lyso) which might be sent to MVB-ysosome via the degradation pathway; the specimens were prepared with **Scheme 2d (MMTS)** (See **Fig.4c** & **Supplementary Table 3**): the cells cultured on sapphire discs were oxidized by 25mM MMTS for 5 min at room temperature, then fixed with 0.1% GA (kept the MMTS) for 5 min at RT, neutralized with 10% glycine overnight on ice, permeabilized with 0.1% Triton x-100 for 2 min,change to PBS-A buffer for ANSM/AuNPs synthesis (Refer to “Protocol for mammalian cells attached to sapphire discs” in the online methods for details). (**d**) a lot of AuNPs distributed in expanded cisternae of ERs (pointed by white arrows), but no AuNPs in the mitochondria (marked with ‘M’); The cells were grown on a 35mm diameter well of a Corning 6-well cell cuture plate, and digested with typsin-EDTA for oxidization with 3mM DTDPA at 4˚C for 30 min, then fixed with 0.1% GA at 4˚C for 30 min, neutralized with 10% glycine overnight, permeabilized with 0.1% Triton X-100 for 2 min,change to PBS-A buffer for ANSM/AuNPs synthesis (refer to “protocol for AuNP synthesis in mammalian cells in suspension”in online methods for details). Specimen in (d) was prepared with **Scheme 2b** (See **Fig.4c** & **Supplementary Table 3**).

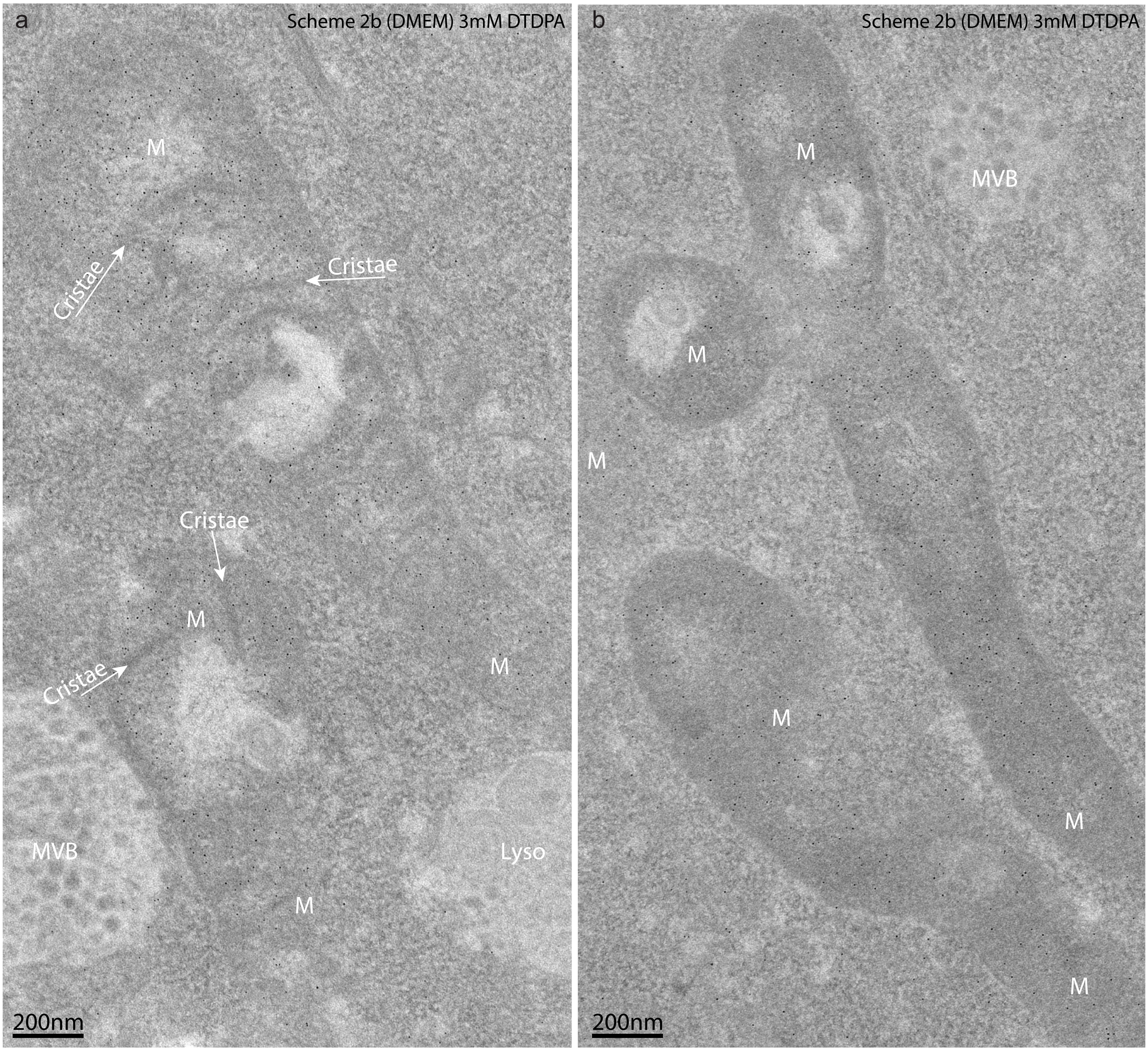


**Supplementary Figure 23A Distributions of AuNPs synthesized on mito-GFP-MTn that expressed in HeLa cells**
(a-b) AuNPs mainly distributed inside the matrix of mitochondria, but almost no AuNPs on the cristae of mitochondion (pointed by the white arrows); the AuNPs density in the cytosol, MVB, lysosome (Lyso) are relative low. The specimens were prepared with a modified **Scheme 2b (PBS)** (See Fig.4c & Supplementary Table 3): the cells cultured on sapphire discs were oxidized by 3mM DTDPA in DMEM for 30 min at 4°C, then further fixed with 0.1% GA in DMEM for 30 min at 4°C, neutralized with 10% glycine overnight on ice, permeabilized with 0.1% Triton X-100 for 2 min,change to PBS-A buffer for ANSM/AuNPs synthesis (Refer to “Protocol for mammalian cells attached to sapphire discs” in the online methods for details).

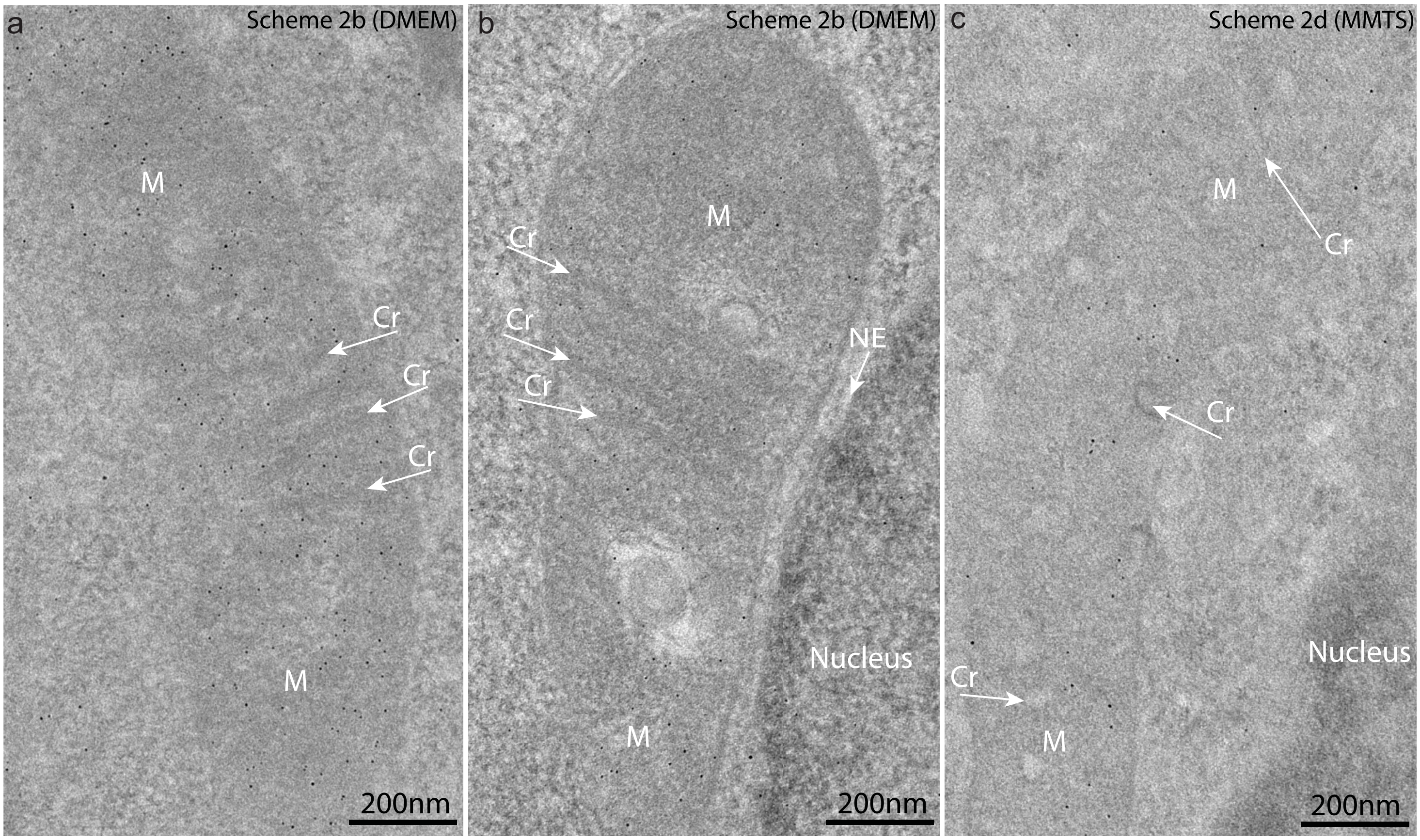


**Supplementary Figure 23B Distributions of AuNPs synthesized on mito-GFP-MTn that expressed in HeLa cells**
(**a-b**) AuNPs mainly distributed inside the matrix of mitochondria (M), but almost no AuNPs on the cristae (Cr) of mitochondion (pointed by the white arrows), clearly shown some AuNPs located to the matrix between the adjacent cristae; the AuNPs density in the cytosol, nucleus are relative low. The specimens were prepared with **Scheme 2b (DMEM)** (See **Supplementary Table 3**): the cells cultured on sapphire discs were oxidized by 3mM DTDPA in DMEM for 30 min at 4°C, then further fixed with 0.1% GA in DMEM for 30 min at 4°C, neutralized with 10% glycine overnight on ice, permeabilized with 0.1% Triton X-100 for 2 min, change to PBS-A buffer for ANSM/AuNPs synthesis (Refer to "Protocol for mammalian cells attached to sapphire discs" in the online methods for details). (**c**) AuNPs specifically locaized to matrix of mitochon of a specimen prepared with another Scheme 2d (MMTS) ( see Supplementary Table 3): the cells cultured on sapphire discs were oxidized by 25mM MMTS in PBS for 5 min at room temperature (RT), then further fixed with 0.1% GA for 5 min at RT, neutralized with 10% glycine overnight on ice, permeabilized with 0.1% Triton X-100 for 2 min,change to PBS-A buffer for ANSM/AuNPs synthesis (Refer to "Protocol for mammalian cells attached to sapphire discs" in the online methods for details).

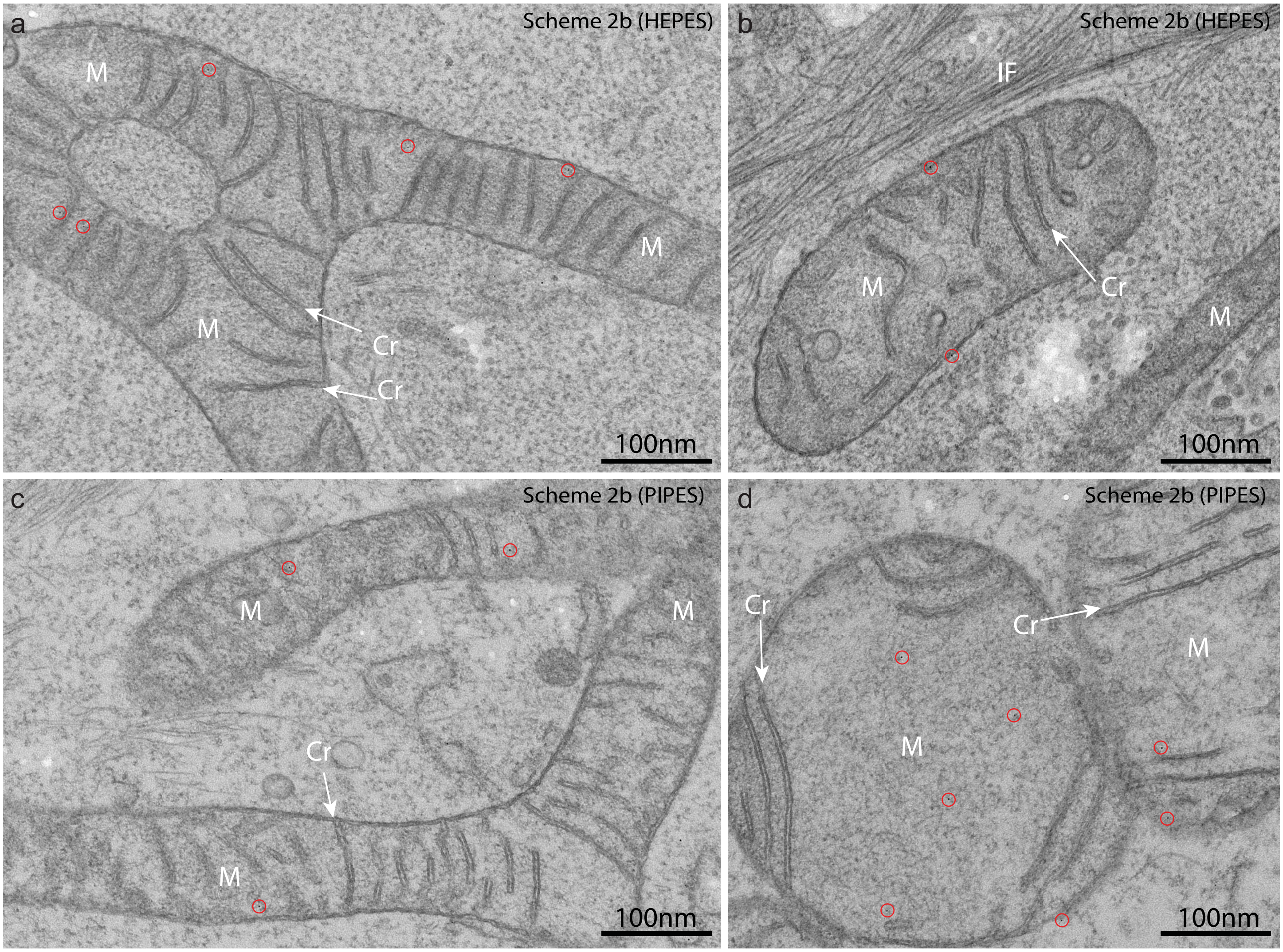


**Supplementary Figure 23C Distributions of AuNPs synthesized on mito-acGFP-MTn that expressed in HeLa cells** (**a-b**) EM images of HeLa cells expressed mito-acGFP-MTn, the specimens were prepared with Scheme 2b (HEPES) (See **Supplementary Table 3**): the cells cultured on sapphire discs were oxidized by 3 mM DTDPA in 0.2 M HEPES buffer with 1 mM $CaCl_2$ and 1 mM $MgCl_2$ (pH 7.4) for 30 min at 4°C, then fixed with additional 0.1% GA for 15min at 4°C. neutralized with 50mM glycine in HEPES buffer overnight on ice, permeabilized with 0.05% Triton X-100 in PBS for 2 min,change to PBS-A buffer for ANSM/AuNPs synthesis (Refer to "Protocol for mammalian cells attached to sapphire discs" in the online methods for details). (**c-d**) EM images of HeLa cells expressed mito-acGFP-MTn, the specimens were prepared with Scheme 2b (PIPES) (See **Supplementary Table 3**): the cells cultured on sapphire discs were oxidized by 3mM DTDPA in 0.1M PIPES buffer with 1m M $CaCl_2$ and 1 mM $MgCl_2$ (pH 7.2) for 30 min at 4°C, then fixed with additional 0.1% GA for 15 min at 4°C,neutralized with 10% glycine in PIPES buffer overnight on ice, permeabilized with 0.05% Triton X-100 in PBS for 2 min,change to PBS-A buffer for ANSM/AuNPs synthesis (Refer to "Protocol for mammalian cells attached to sapphire discs" in the online methods for details). It should be noted that using both HEPES and PIPES buffer with 1 mM $CaCl_2$ and 1 mM $MgCl_2$ were effective in stablizing the membrane structures of mitochondria (M), as indicated by the well-preserved cristae (Cr), however, the number of AuNPs (marked with red circles) in mitchondria were quite limited (it might be caused by the reduced permeability of the membranes, which resulted in bad thiol groups protection due to the insufficient DTDPA concentration in the mitochondrial matrix).

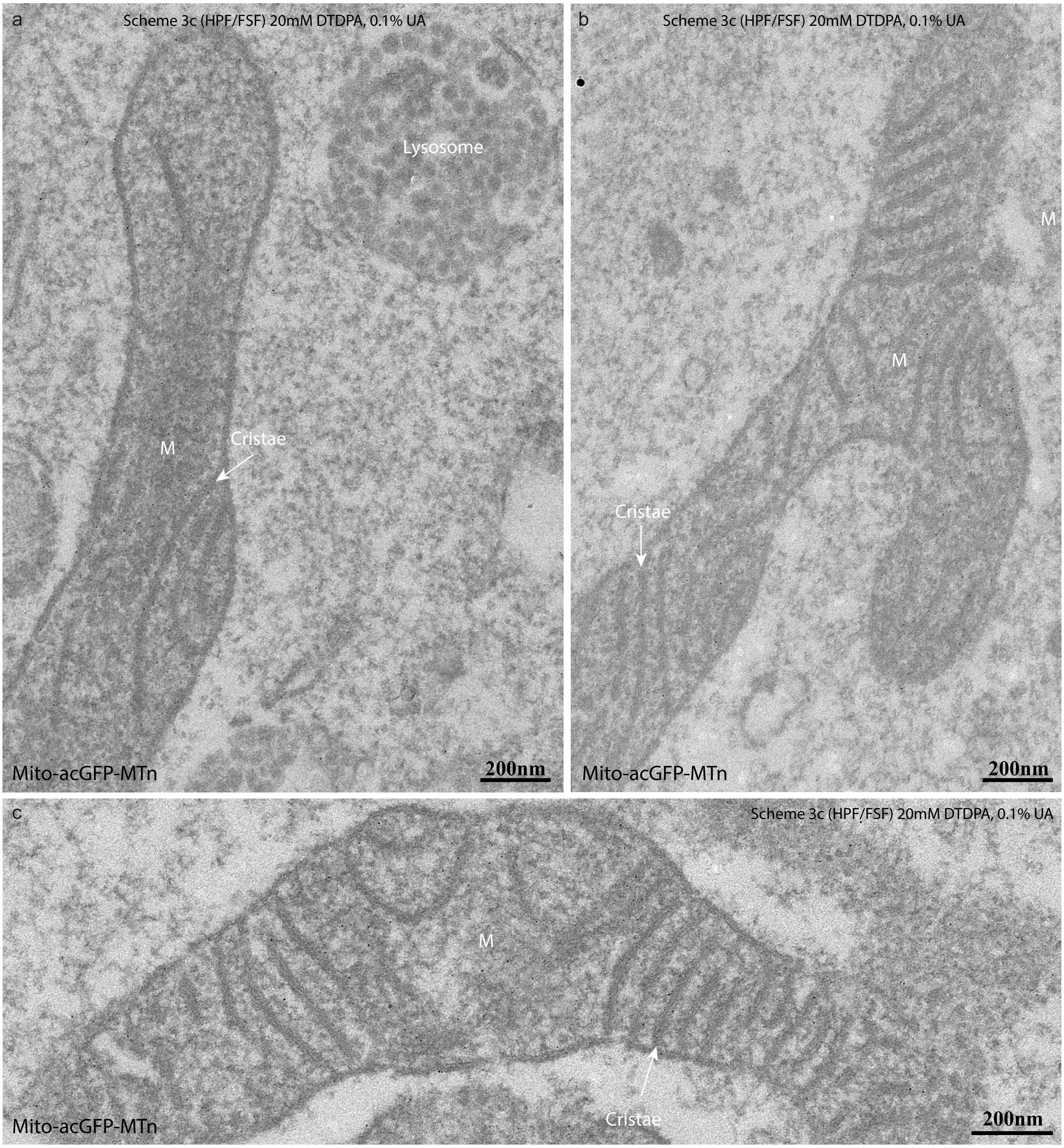


**Supplementary Figure 24A Distributions of AuNPs synthesized in HeLa cells expressing Mito-acGFP-MTn rapidly fixed with high pressure freezing**

(a-c) EM images of 90nm thick sections from Mito-acGFP-MTn expressed HeLa cells prepared by Scheme 3c (HPF/FSF) (**SupplementaryTable 3**). The HeLa cells cultured on sapphire discs, then rapid fixed with high pressure freezing, freeze-substitution and oxidization performed 20 mM DTDPA in acetone with 5% $H_2O$ and 1% methanol, added 0.1% UA for 5 h fixation at -30 °C, then sucked away the fixative medium and filled with 1 mL of ice-cooled 0.2M HEPES buffer with 1 mM $CaCl_2$ and 1 mM $MgCl_2$ for instant rehydration on ice, incubated with 10% glycine in HEPES buffer overnight on ice, changed to PBS-A buffer for AuNP synthesis (Refer to "Procedures for FSF-based AuNP synthesis in mammalian cells" in the online methods for details). There were higher densities of AuNPs formed in the matrix of mitochondrion (M), some AuNPs localized to the lysosome, a few AuNPs in cytosol (it is impossible to seperare background noises from the Mito-acGFP-MTn molecules synthesized in cytosol). The membrane structures were well preserved by using both UA and HEPES buffer although the cytosol was not well preserved.

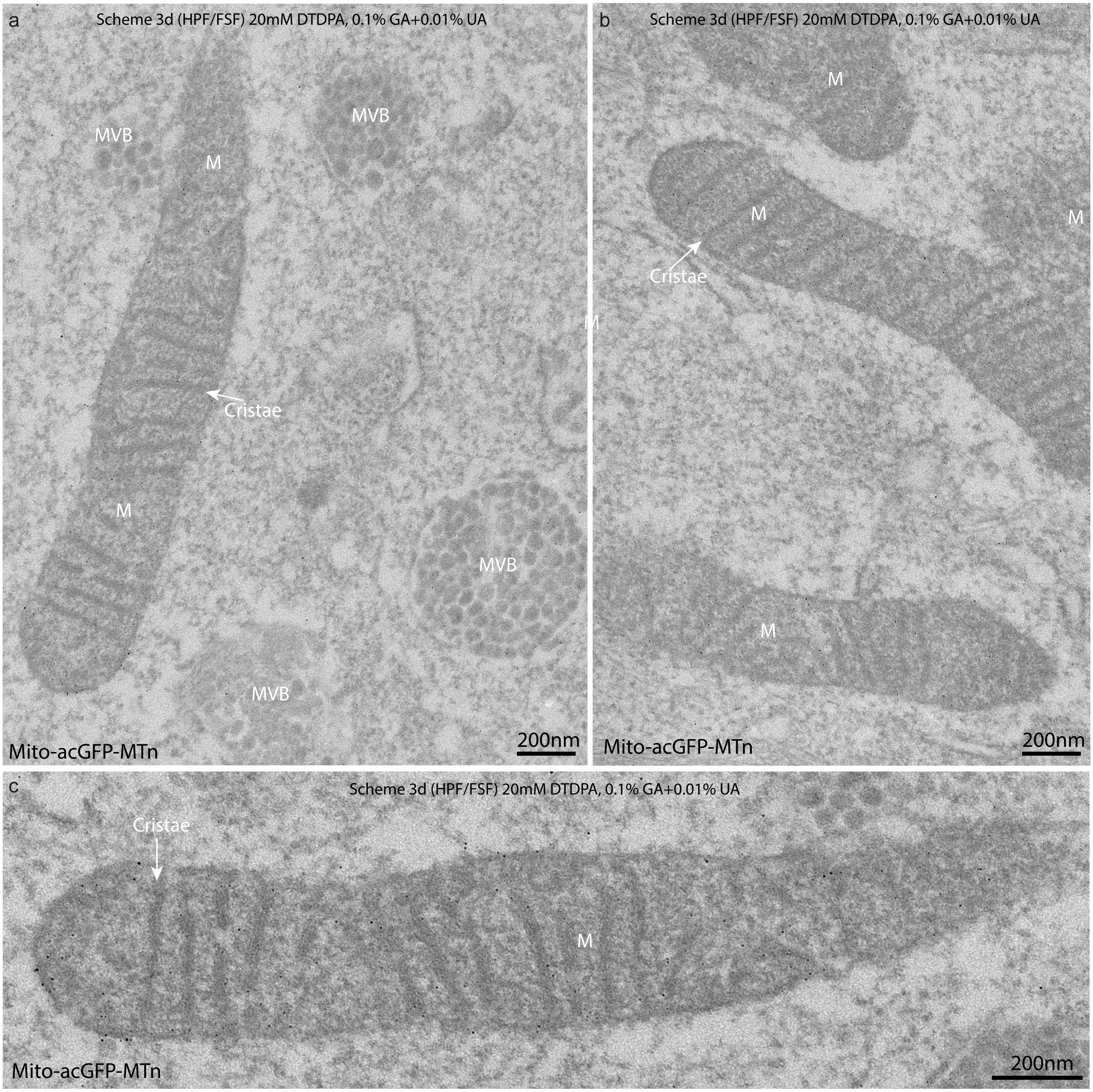


**Supplementary Figure 24B Distributions of AuNPs synthesized in HeLa cells expressing Mito-acGFP-MTn rapidly fixed with high pressure freezing**

(**a-c**) EM images of 90nm thick sections from HeLa cells expressing Mito-acGFP-MTn prepared by Scheme 3d (HPS/FSF) (**Supplementary Table 3**). The HeLa cells cultured on sapphire discs, then rapid fixed with high pressure freezing, freeze-substitution and oxidization performed 20 mM DTDPA in acetone with 5% $H_2O$ and 1% methanol, added 0.1% GA and 0.01% UA for 5 h fixation at -30 °C, then sucked away the fixative medium and filled with 1 mL of ice-cooled 0.2 M HEPES buffer with 1 mM $CaCl_2$ and 1 mM $MgCl_2$ for instant rehydration, incubated with 10% glycine in HEPES buffer overnight on ice, changed to PBS-A buffer for AuNP synthesis (Refer to “Procedures for FSF-based AuNP synthesis in mammalian cells” in the online methods for details). There were lots of AuNPs formed in the matrix of mitochondrion (M), some AuNPs localized to the MVBs (a-b). (**c**) The enlarged image of the mitochondrion shown in (a), AuNPs were scattered in the matrix.
